# Peroxisome-derived hydrogen peroxide can modulate the sulfenylation profiles of key redox signaling proteins

**DOI:** 10.1101/2021.10.08.463647

**Authors:** Celien Lismont, Iulia Revenco, Hongli Li, Cláudio F. Costa, Lisa Lenaerts, Mohamed A. F. Hussein, Jonas De Bie, Bernard Knoops, Paul P. Van Veldhoven, Rita Derua, Marc Fransen

**Affiliations:** Laboratory of Peroxisome Biology and Intracellular Communication, Department of Cellular and Molecular Medicine, KU Leuven, O&N1bis, Herestraat 49 box 901, B-3000 Leuven, Belgium; Laboratory of Protein Phosphorylation and Proteomics, Department of Cellular and Molecular Medicine, KU Leuven, O&N1, Herestraat 49 box 901, B-3000 Leuven, Belgium; Department of Biochemistry, Faculty of Pharmacy, Assiut University, 71515 Assiut, Egypt; Group of Animal Molecular and Cellular Biology, Institut des Sciences de la Vie (ISV), Université Catholique de Louvain, Louvain-la-Neuve, Belgium; SyBioMa, KU Leuven, O&N2, Herestraat 49, B-3000 Leuven, Belgium

**Keywords:** Peroxisome, hydrogen peroxide, cysteine thiol group, YAP1C-based sulfenome mining, peroxiredoxin, mitochondrion

## Abstract

Ever since the first characterization of peroxisomes, a central theme has been their involvement in cellular hydrogen peroxide (H_2_O_2_) metabolism. While the reputation of H_2_O_2_ drastically changed from an exclusively toxic molecule to a signaling messenger, the regulatory role of peroxisomes in these signaling events is still largely underappreciated. This is mainly because the number of known protein targets of peroxisome-derived H_2_O_2_ is rather limited and testing of specific targets is predominantly based on knowledge previously gathered in related fields of research. To gain a broader and more systematic insight into the role of peroxisomes in redox signaling, an unbiased approach is urgently needed. To accomplish this goal, we have combined a previously developed cell system in which peroxisomal H_2_O_2_ production can be modulated with a yeast AP-1-like-based sulfenome mining strategy to inventory protein thiol targets of peroxisome-derived H_2_O_2_ in different subcellular compartments. Using this unbiased approach, we were able to identify specific and common targets of peroxisome-derived and exogenous H_2_O_2_ in peroxisomes, the cytosol, and mitochondria. We also observed that the sulfenylation kinetics profiles of key targets belonging to different protein families can vary considerably. In addition, we obtained compelling but indirect evidence that peroxisome-derived H_2_O_2_ may oxidize at least some of its targets through a redox relay mechanism. In conclusion, given that sulfenic acids function as key intermediates in H_2_O_2_ signaling, the findings presented in this study provide initial but critical insight into how peroxisomes may be integrated in the cellular H_2_O_2_ signaling network.

**Highlights:**

- YAP1C-trapping is a robust tool to assess the peroxisomal H_2_O_2_-dependent sulfenome
- Exogenous and peroxisome-derived H_2_O_2_ have both common and distinct targets
- ANXA2, PRDX1, and SKP1 are major targets of peroxisome-derived H_2_O_2_
- The sulfenylation kinetics profiles of key redox-active proteins vary considerably
- Production of H_2_O_2_ inside peroxisomes directly impacts the mitochondrial sulfenome

## Introduction

Hydrogen peroxide (H_2_O_2_) has become recognized as one of the major physiological signaling agents [Sies and Jones, 2020]. Depending on the cellular context and its local concentration, this oxidant may exhibit antagonistic pleiotropic effects, ranging from cell proliferation, differentiation, and migration to stress adaptations, growth arrest, and even cell death [Lennicke et al., 2015; Sies and Jones, 2020]. A major mechanism by which H_2_O_2_ mediates its biological action is through protein thiol oxidation, a process that may trigger changes in protein structure, biochemical activity, subcellular localization, and/or binding affinity. A potential strategy to provide more insight into how temporary changes in local H_2_O_2_ levels can mediate signaling events is to inventory the oxidized proteins and cysteinyl residues involved. However, a key factor for successful implementation of such an approach is to have access to a robust model system in which H_2_O_2_ production and redox-active cysteine trapping can be strictly controlled in a spatiotemporal manner.

We recently developed a DD-DAO Flp-In T-REx 293 cell line-based approach that allows to modulate intracellular H_2_O_2_ production in a subcellular compartment-, dose-, and time-dependent manner [Lismont et al., 2019a]. These cells are characterized by the doxycycline (DOX)^1^-inducible expression of destabilization domain (DD)-tagged variants of D-amino acid oxidase (DAO), a peroxisomal flavoprotein that generates H_2_O_2_ while it oxidizes neutral and polar (but not acidic) D-amino acids to their corresponding imino acids. The subcellular localization of DD-DAO can be easily altered by inactivating its C-terminal peroxisomal targeting signal (PTS1) and/or appending other targeting signals. For example, we have generated stable cell lines in which DD-DAO, upon induction by DOX, is localized in the cytosol (c-DD-DAO) or the peroxisome lumen (po-DD-DAO) [Lismont et al., 2019a]. Importantly, to stabilize DD-DAO in the cytosol or to allow efficient post-translational import of this fusion protein into the organelle under study, the cells need to be cultured in the presence of both DOX and Shield1. The latter compound is a small cell-permeable molecule that binds to DD, thereby protecting cytosolic and nuclear DD-containing proteins from proteasomal degradation. To get rid of the not-yet-imported pool of organelle-targeted DD-DAO, which otherwise may complicate the interpretation of the results, the cells can – before the time of analysis – be chased in culture medium lacking DOX and Shield1. To control the amount and duration of H_2_O_2_ production, varying concentrations of D-amino acids can be added to or withdrawn from the assay medium.

The primary messenger action of H_2_O_2_ depends on its ability to react with deprotonated cysteine residues (Cys-S^-^), a process that in first instance leads to the formation of (unstable) sulfenic acid (-SOH) intermediates [Lismont et al., 2019b]. Interestingly, to gain more insight into the H_2_O_2_-dependent sulfenome *in cellulo,* a C-terminal region of yeast AP-1-like transcription factor (YAP1C)-based strategy was developed to trap, visualize, and enrich proteins that are sulfenylated in response to external H_2_O_2_ treatment in *Escherichia coli* [Takanishi et al., 2007] and *Arabidopsis thaliana* [Waszczak et al., 2014]. Here, we adopted this approach to trap, visualize, and enrich peroxisomal, cytosolic, or mitochondrial proteins that are sulfenylated in human cells in response to peroxisome-derived or externally added H_2_O_2_.

Peroxisomal respiration may be responsible for up to 20% of total oxygen consumption and 35% of total H_2_O_2_ production, at least in some mammalian tissues such as liver [de Duve and Baudhuin, 1966; Boveris et al., 1972]. In addition, disturbances in peroxisomal H_2_O_2_ metabolism have been associated with aging and age-associated disease [Fransen and Lismont, 2019]. Despite this, very little is known about how peroxisomes are embedded in H_2_O_2_-mediated signaling networks. In this study, we were able for the first time to map the potential impact of peroxisome-derived H_2_O_2_ on cellular redox signaling networks. This opens new perspectives for research on how perturbations in peroxisomal H_2_O_2_ metabolism may contribute to the initiation and development of oxidative stress-related diseases.

## Materials and Methods

### Plasmids

The cDNA coding for a human codon-optimized variant of IBD-SBP-YAP1C (Fig. S1) was synthesized by Integrated DNA Technologies and provided in the pUCIDT (Kan) cloning vector (pMF1986). A mammalian expression plasmid encoding IBD-SBP-YAP1C (pMF1987) was generated by transferring the EcoRI/NotI-restricted fragment of pMF1986 into the EcoRI/NotI-restricted backbone fragment of pMF1839 [Walton et al., 2017]. Mammalian expression vectors encoding mitochondrial (pMF1991), peroxisomal (pMF1992), or cytosolic (pMF2029) variants of IBD-SBP-YAP1C were generated by amplifying the IBD-SBP-YAP1C template via PCR (forward oligo: 5’-gggggatcccatggcatcaatgcagaagctg-3’; reverse oligos: 5’-ccgggggcggccgctcagttcatatgtt-3’ (for pMF1991 and pMF2029) and 5’-ccgggggcggccgctcaaagcttacttttgttcatatgtttattcaatgca-3’ (for pMF1992)) and subcloning the BamHI/NotI-restricted PCR products into the BamHI/NotI-restricted backbone fragments of pKillerRed-dMito (Evrogen) (for pMF1991) or pEGFP-N1 (Clontech) (for pMF1992 and pMF2029). The correctness of all plasmids was verified by DNA sequencing (LGC Genomics). The plasmids encoding EGFP-HsPEX11B (pTW110) [Fransen et al, 2001], c-roGFP2 (pMF1707) [Ivashchenko et al., 2011], po-roGFP2 (pMF1706) [Ivashchenko et al., 2011], or mt-roGFP2 (pMF1762) [Ivashchenko et al., 2011] have been described elsewhere. The plasmid encoding EGFP-HSPB1 was kindly provided by Prof. Dr. Ludo Van Den Bosch (KU Leuven, Belgium).

### Cell culture and transfections

Cell culture was essentially performed as previously described [Lismont et al., 2019a]. Briefly, all cells were cultured at 37°C in a humidified 5% CO2 incubator in minimum essential medium Eagle α (Lonza; BE12-169F) supplemented with 10% (v/v) fetal bovine serum (Biowest; S181B), 2 mM UltraGlutamine I (Lonza; BE17-605E/U1), and 0.2% (v/v) MycoZap (Lonza; VZA-2012). Transfections were performed by using the Neon Transfection System (Thermo Fisher Scientific; 1150 V, 20-ms pulse width, 2 pulses).

### Generation and manipulation of DD-DAO/IBD-SBP-YAP1C Flp-In T-REx 293 cell lines

The po-DD-DAO and c-DD-DAO Flp-In T-REx 293 cell lines have been detailed elsewhere [Lismont et al., 2019a]. To generate po- or c-DD-DAO Flp-In T-REx 293 cell lines constitutively expressing c-IBD-SBP-YAP1C, po-IBD-SBP-YAP1C, or mt-IBD-SBP-YAP1C, the corresponding Flp-In T-REx 293 cells were transfected with pMF2029, pMF1992, or pMF1991, respectively. Starting from two days later, the cells were routinely cultured in medium supplemented with (i) 10 μg/ml blasticidin (InvivoGen; ant-bl) and 100 μg/ml hygromycin B Gold (InvivoGen, ant-hg) to maintain the properties of the po- or c-DD-DAO Flp-In T-REx 293 stable cell lines, and (ii) 200 μg/ml of G418 (Acros Organics, BP673-5) to select for cells carrying the neomycin resistance cassette (with a minimum period of 3 weeks). To modulate the expression levels of DD-DAO in these cells, they were incubated in the absence or presence of 1 μg/ml doxycycline (DOX) (Sigma, D9891) and 500 nM Shield1 (Clontech, 632189). Treatments were always followed by a one-day chase period (no DOX, no Shield1) in order to remove the pool of residual cytosolic po-DD-DAO [Lismont et al., 2019a], unless specified otherwise.

### Fluorescence microscopy

Fluorescence microscopy was carried out as described previously [Ramazani et al., 2021]. The following excitation filters (Ex), dichromatic mirrors (Dm), and emission filters (Em) were chosen to match the fluorescent probe specifications: DAPI (Ex: BP360-370; Dm: 400 nm cut-off; Em: BA420-460); EGFP (Ex: BP470-495; Dm: 505 nm cut-off; Em: BA510-550); and Texas Red (Ex: BP545-580; Dm: 600 nm cut-off; Em: BA610IF). The cellSens software (Olympus Belgium) was used for image analysis. Samples for immunofluorescence microscopy were fixed, counterstained with 0.5 μg/ml DAPI (Sigma, D-9542) in Dulbecco’s phosphate-buffered saline (DPBS) for 1 min, and processed as described [Passmore et al., 2020].

### Redox proteomics sample preparation and analysis

Cells were grown to 60-80% confluency, trypsinized, collected in cell culture medium, pelleted (150 x g, 5 min), and washed once with DPBS without calcium and magnesium (BioWest, L0615). Cell density was determined by Bürker chamber counting and adjusted to 10^6^ cells/ml. After being subjected to different treatments (for specifications, see Results section), N-ethylmaleimide (NEM) (TCI, E0136) dissolved in methanol (Fisher Scientific, M/4062/17) was added to a final concentration of 10 mM. Next, the cells were pelleted (150 x g, 5 min), resuspended in lysis buffer (50 mM Tris-HCl pH 7.5, 150 mM NaCl, 1% (v/v) Triton X-100, 10% (v/v) glycerol) containing 10 mM NEM and a protease inhibitor mix (Sigma-Aldrich, P2714)) at a density of 10^7^ cells/ml, and lysed on ice for 10 min. Thereafter, a cleared lysate was produced by double centrifugation (20,000 x g, 10 min), each time discarding the pellet.

The cleared lysates were mixed with 300 μl of (prewashed) high capacity streptavidin agarose beads (Thermo Scientific, 20359) and incubated at 4°C on a rotation mixer. After 2 h, the bead suspensions were transferred to Micro-Spin columns (Thermo Scientific, 89879) and consecutively washed 5 times with lysis buffer, 5 times with wash buffer (50 mM Tris-HCl pH 7.5, 150 mM NaCl, 1% (v/v) Triton X-100), and 5 times with 50 mM Tris-HCl pH 8.0. Finally, the IBD-SBP-YAP1C-trapped proteins were eluted by incubating the columns 3 times for 15 min with 200 μl of elution buffer (10 mM DTT in 50 mM Tris-HCl pH 8.0). The eluates were pooled and subsequently processed for proteomics analysis. To analyze comparable amounts of IBD-SBP-YAP1C-containing protein complexes, the ratio of cleared lysate to streptavidin agarose beads were chosen such that the affinity matrix was slightly oversaturated, as determined by detection of residual IBD-SBP-YAP1C in the non-bound fraction.

Eluates were incubated for 30 min at 37°C to allow the DTT to fully reduce all proteins. Thereafter, the samples were alkylated (37°C, 30 min) with 25 mM iodoacetamide (Sigma, I-6125). Excess iodoacetamide was quenched with 25 mM DTT (AppliChem, A2948) (37°C, 30 min). Subsequently, the proteins were precipitated as described [Wessel and Flügge, 1984] and digested overnight with modified trypsin (Pierce, 90057) in the presence of 50 mM ammonium bicarbonate (Sigma-Aldrich, A6141), 5% acetonitrile (Applied Biosystems, 400315), and 0.01% ProteaseMAX surfactant (Promega, V2072). Trypsin was inactivated by addition of 0.5% (v/v) trifluoroacetic acid (Applied Biosystems, 400028). The resulting peptides were desalted with C18 ZipTip pipette tips (Merck Millipore, ZTC18S960) and loaded onto an Ultimate 3000 UPLC system (Dionex, Thermo Fisher Scientific) equipped with an Acclaim PepMap100 pre-column (C18; particle size: 3 μm; pore size 100 Å; diameter: 75 μm; length: 20 mm; Thermo Fisher Scientific) and a C18 PepMap analytical column (particle size: 2 μm; pore size: 100 Å; diameter: 50 μm; length: 150 mm; Thermo Fisher Scientific) using a 40 min linear gradient (300 nL/min) coupled to a Q Exactive Orbitrap mass spectrometer (Thermo Fisher Scientific) operated in data-dependent acquisition mode. After the initial pilot experiment, the mass spectrometry (MS) method was adapted, essentially doubling the maximum injection time for MS/MS. Peptides were identified by Mascot (Matrix Science) using Uniprot *Homo sapiens* as a database (# entries: 194619). S-carbamidomethylation (C), N-ethylmaleimide (C), and oxidation (M) were included as variable modifications. Two missed cleavages were allowed, peptide tolerance was set at 10 ppm and 20 mmu for MS and MS/MS, respectively. Progenesis QI software (Nonlinear Dynamics) was used for relative quantification of proteins based on peptides validated by the Proteome Discoverer 2.2 Percolator node (FDR < 1%). Only proteins that were identified by at least 2 exclusive peptides or by 1 exclusive peptide with a posterior error probability (PEP) smaller than 0.05 were taken into account.

Proteins that could not be unambiguously identified or were identified as keratins, extracellular proteins, or proteins that – according to the Human Protein Atlas database (http://www.proteinatlas.org) – are not expressed in HEK-293 cells, were manually removed. Proteins enriched at least two-fold upon H_2_O_2_ exposure were retained for further analysis. Only exclusive peptides were taken into account for quantification.

### Antibodies

The pre-immune serum was collected from a rabbit before immunization with the 21 kDa subunit of rat palmitoyl-CoA oxidase [Baumgart et al., 1996]; the rabbit polyclonal antisera against EGFP [Fransen et al., 2001], PEX13 [Fransen et al., 2001], PRDX1 [Goemaere and Knoops, 2012], PRDX3 [Goemaere and Knoops, 2012], or PRDX5 [Goemaere and Knoops, 2012] have been described elsewhere; and the rabbit polyclonal antisera against TUBA (Santa Cruz Biotechnology; sc-5546) and the goat anti-rabbit secondary antibodies, conjugated to Texas Red (Calbiochem, 401355) or alkaline phosphatase (Sigma, A3687), were commercially obtained.

### Mobility shift electrophoresis

Electrophoretic mobility shift assays (EMSAs) were performed as described previously [Lismont et al., 2019a]. A slightly modified protocol was used for samples taken during the proteomics sample preparation. Specifically, for the “input” and “non-bound” samples, 50 μl of cleared cell lysates and bead supernatants were respectively mixed with 2 X SDS-PAGE sample buffer without reducing agent and heated to 65°C for 10 min. For the “bound” and “bound after elution” samples, 10 μl of bead volume was mixed with non-reducing 2 X SDS-PAGE sample buffer and heated to 100°C for 10 min.

### Data availability

The mass spectrometry raw data and search result files will be deposited in the PRIDE repository (www.ebi.ac.uk/pride/) upon acceptance of the manuscript.

### Other

Subcellular fractionations of rat liver [Anthonio et al., 2009] and HEK-293 cells [Lismont et al., 2019c] were carried out as described elsewhere. Functional enrichment analysis was performed using g:GOST (https://biit.cs.ut.ee/gprofiler/gost) within the g:Profiler tool [Raudvere et al., 2019]. Heat maps were generated with Graphpad Prism version 9.0.0 for Windows (GraphPad Software). The iCysMod (http://icysmod.omicsbio.info/index.php) [Wang et al., 2021] and TF2DNA (http://fiserlab.org/tf2dna_db//index.html) [Pujato et al., 2014] databases were used as resources to search for known protein cysteine oxidation sites and transcription factors in the human dataset, respectively.

## Results

### Validation of the po-DD-DAO/IBD-SBP-YAP1C Flp-In T-REx 293 cell lines

To profile peroxisome-derived H_2_O_2_-mediated protein thiol oxidation in a spatiotemporal manner in mammalian cells, po-DD-DAO Flp-In T-REx 293 cells [Lismont et al., 2019a] were first electroporated with a plasmid encoding a peroxisomal (po-), cytosolic (c-), or mitochondrial (mt-) variant of human codon-optimized IBD-SBP-YAP1C and subsequently cultured for at least 3 weeks in regular medium supplemented with 200 μg/ml of G418 to enrich cells that have taken up the plasmid. IBD-SBP-YAP1C is a hybrid protein in which (i) the YAP1C moiety can react with and trap protein sulfenic acids, (ii) the SBP domain contains a high-affinity streptavidin-binding peptide that can be used to enrich IBD-SBP-YAP1C complexes on streptavidin matrices, and (iii) the IBD domain consists of two streptococcal protein G IgG-binding domains that enable visualization of IBD-SBP-YAP1C complexes in IgG overlay assays. To confirm the correct localization of the different IBD-SBP-YAP1C fusion proteins in the G418-enriched cell lines, cells were transfected with a plasmid encoding a relevant fluorescent subcellular localization marker and, three days later, processed for immunofluorescence microscopy by using a pre-immune rabbit serum and a goat anti-rabbit secondary antibody conjugated to Texas Red (Fig. 1). From this experiment, it is clear that the majority of the cells express IBD-SBP-YAP1C. However, it is also evident that, at least in some cells, varying portions of po-IBD-SBP-YAP1C and, to a certain extent, also mt-IBD-SBP-YAP1C still reside in the cytosol. Given that (i) the functionality of peroxisomal and mitochondrial targeting signals fused to a heterologous protein strongly depends on the protein context [Yogev and Pines, 2011; Kunze et al., 2018], (ii) the priority of protein import into mitochondria and peroxisomes is governed by competition for binding to limiting amounts of import receptor [Weidberg and Amon, 2018; Rosenthal et al., 2020], and (iii) expression of the YAP1C fusion proteins is driven by the cytomegalovirus promoter, one of the strongest naturally occurring promoters [Even et al., 2016], this observation may not be that surprising. Although this observation may complicate the interpretation of downstream results, this knowledge also allows us to correctly anticipate this shortcoming.

**Figure 1.**
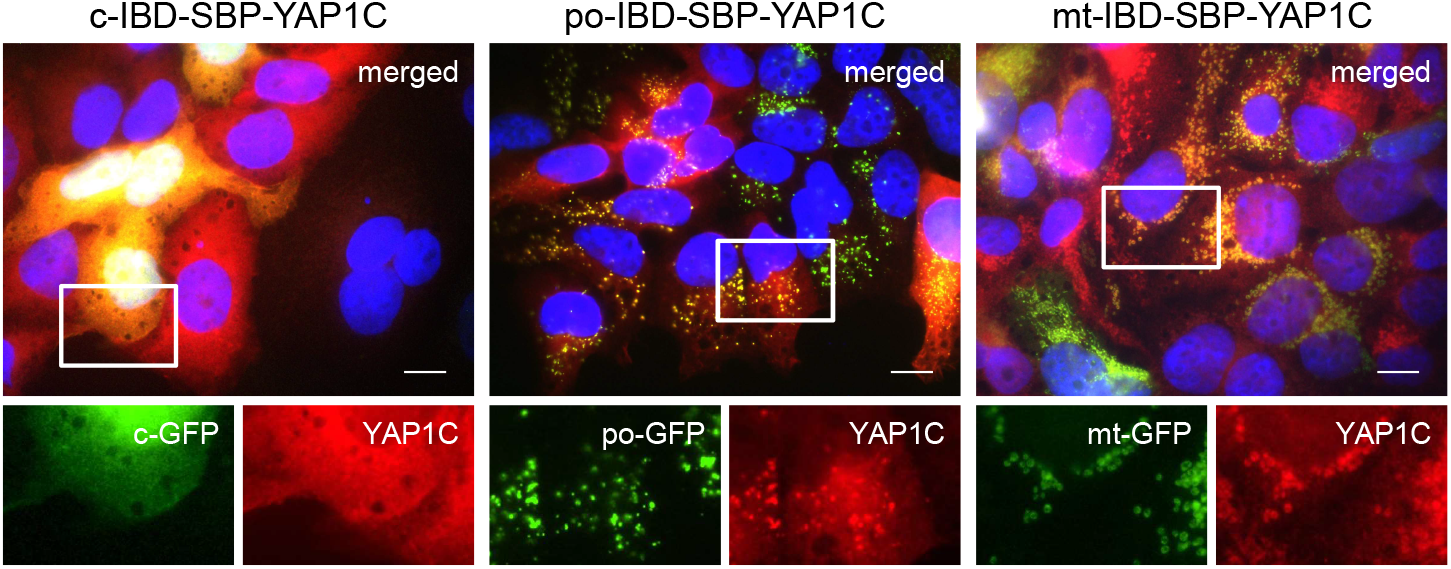
Validation of the subcellular localization of different IBD-SBP-YAP1C fusion proteins in po-DD-DAO Flp-In T-REx 293 cells. Po-DD-DAO Flp-In T-REx 293 cells enriched for expression of c-, po-, or mt-IBD-SBP-YAP1C were transfected with plasmids encoding c-, po-, or mt-roGFP2 (GFP) as marker for the respective cell compartment. After three days, the cells were processed for immunofluorescence microscopy using rabbit pre-immune serum and goat anti-rabbit secondary antibody conjugated to Texas Red. Nuclei were counterstained with DAPI. Scale bars, 10 μm. Representative images are shown. The boxed areas in the upper panels are enlarged in the lower panels. IBD, IgG-binding domain; SBP, streptavidin-binding domain.

### Differentially-localized YAP1C proteins form different protein complexes upon exposure of cells to exogenous or peroxisome-derived H_2_O_2_

To validate the *in cellulo* trapping strategy for sulfenylated proteins, we first investigated IBD-SBP-YAP1C complex formation in different subcellular compartments upon exposure of cells to exogenous or peroxisome-derived H_2_O_2_. An outline of the experimental workflow is depicted and detailed in Fig. 2A. In a first experiment, po-DD-DAO Flp-In T-REx 293 cells expressing c-IBD-SBP-YAP1C were exposed or not to 1 mM H_2_O_2_ for 10 min and subsequently processed as detailed in the legend to Fig. 2A. IgG blot overlay analysis of the input fractions (I), the non-bound fractions (NB), the streptavidin-bound fractions (B), and the streptavidin-bound fractions after elution with a reducing agent (BE) confirmed that exposure of the cells to external H_2_O_2_ triggered the formation of many c-IBD-SBP-YAP1C-containing higher molecular weight complexes that can be enriched on streptavidin beads and are sensitive to the reducing agent dithiothreitol (DTT) (Fig. 2B). The latter feature is important to allow selective elution of target proteins from the affinity matrix. Next, a similar experiment was performed with po-DD-DAO Flp-In T-REx 293 cells expressing a compartment-specific variant of IBD-SBP-YAP1C and in which peroxisomal H_2_O_2_ production was induced or not by supplementing the assay medium (DPBS) with 10 mM D- or L-Ala, respectively. Multiple IBD-SBP-YAP1C-containing protein complexes could be detected in all conditions in which peroxisomal H_2_O_2_ was produced (Fig. 2C). Interestingly, a direct comparison of the staining patterns of the streptavidin-enriched fractions clearly shows that the interaction profiles of IBD-SBP-YAP1C with sulfenylated proteins differed considerably depending on the source of H_2_O_2_ as well as on the subcellular location of the YAP1C fusion protein (Fig. 2D).

**Figure 2.**
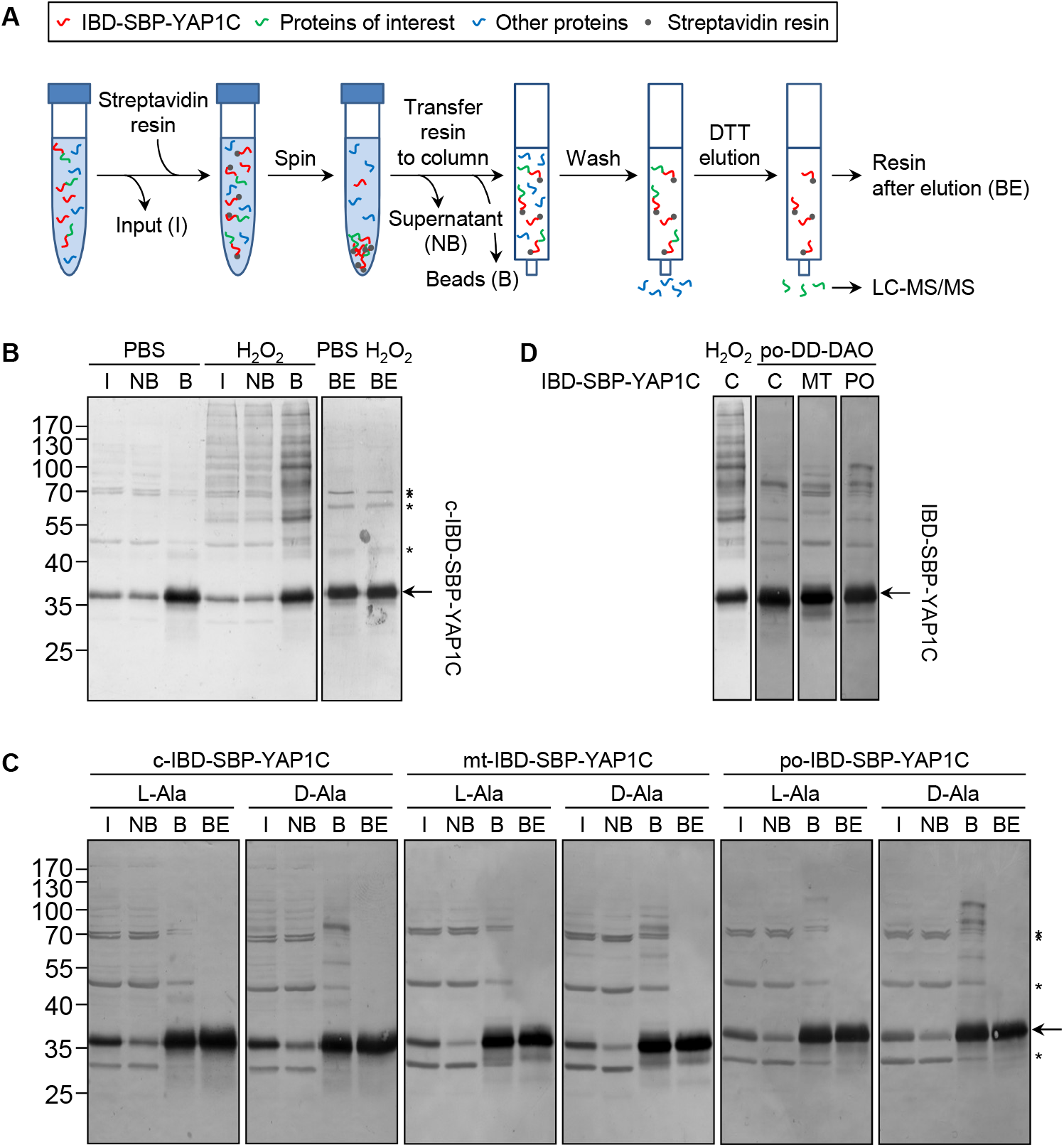
The interaction profile of IBD-SBP-YAP1C with sulfenylated proteins differs considerably depending on the source of H_2_O_2_ as well as its subcellular location. **(A)** Outline of the experimental workflow. After cell treatment, a cleared lysate (= input, I) was prepared. Subsequently, streptavidin agarose beads were added, thereby enriching IBD-SBP-YAP1C complexes on the affinity resin. After a brief spin, the non-bound proteins (NB) present in the supernatant were removed, and the resin containing the bound IBD-SBP-YAP1C complexes (B) was transferred to a Micro-Spin column and thoroughly washed. Finally, the disulfide-bonded interaction partners of IBD-SBP-YAP1C were eluted with reducing agent (DTT) and the eluate was further processed for LC-MS/MS analysis. At each step, small aliquots were saved for immunoblot analysis. BE, beads after elution. **(B)** C-IBD-SBP-YAP1C complex formation upon external H_2_O_2_ treatment. Po-DD-DAO Flp-In T-REx 293 cells expressing c-IBD-SBP-YAP1C were incubated in DPBS containing 10 mM 3-AT and supplemented or not with 10 mM H_2_O_2_. After 10 min, the cells were treated with 10 mM NEM to block free thiols. Next, the samples were further processed as depicted in panel A, and I, NB, B, and BE aliquots were subjected to immunoblotting using rabbit pre-immune serum and goat anti-rabbit secondary antibody coupled to alkaline phosphatase. **(C)** Peroxisomal H_2_O_2_ production triggers IBD-SBP-YAP1C complex formation in the cytosol, mitochondria, and peroxisomes. Po-DD-DAO Flp-In T-REx 293 cells expressing c-, mt-, or po-IBD-SBP-YAP1C were cultured in the presence of DOX (1 μg/ml) and Shield1 (500 nM) for 3 or 4 days, followed by a chase for one day in medium without DOX/Shield1. Next, the cells were incubated in DPBS containing 10 mM 3-AT and supplemented with either 10 mM L- or D-Ala. After 10 min, the cells were treated with 10 mM NEM to block free thiols. The samples were processed as depicted in panel A, and I, NB, B, and BE aliquots were subjected to immunoblotting as detailed in the legend of panel B. **(D)** Comparison of IBD-SBP-YAP1C complexes formed under different experimental conditions. In the two left lanes, the staining patterns of c-IBD-SBP-YAP1C formed upon external H_2_O_2_ addition or peroxisomal H_2_O_2_ production are compared; in the three right lanes, differentially located IBD-SBP-YAP1C complexes formed upon peroxisomal H_2_O_2_ production are compared. The different lanes are the aligned bound samples of those shown in **B** and **C**. The arrows and asterisks mark IBD-SBP-YAP1C and non-specific immunoreactive bands, respectively. IBD, IgG-binding domain; SBP, streptavidin-binding domain.

### The subcellular sulfenome upon exogenous H_2_O_2_ treatment: a pilot experiment

To corroborate the sulfenome mining strategy, we carried out an exploratory experiment in which cells expressing c-, mt-, or po-IBD-SBP-YAP1C were treated or not with 1 mM H_2_O_2_ for 10 minutes. After ensuring that the protocol was carried out correctly (Fig. S2), the DTT eluates were processed for LC-MS/MS analysis. After validation of the identified peptides, probable contaminating proteins (e.g., keratins, IgGs, and extracellular proteins) as well as proteins that could not be unambiguously identified were manually removed. A set of 50 proteins that were enriched two-fold or more in at least one of the H_2_O_2_-treated conditions could be listed (Table S1).

Gene ontology analysis of the hits revealed a significant enrichment for proteins primarily implicated in cell redox homeostasis and cellular oxidant detoxification (Fig. S3). These include GSR, PRDX1, PRDX2, PRDX3, PRDX4, PRDX5, PRDX6, TXN, and TXNRD2. Interestingly, while 18 out of 50 proteins were enriched in all H_2_O_2_-treated samples, each IBD-SBP-YAP1c fusion protein also retained unique interactors (Fig. 3). This finding agrees with our previous results showing that differentially-localized IBD-SBP-YAP1C proteins form different complexes upon treatment of cells with external H_2_O_2_ (Fig. 2D).

**Figure 3.**
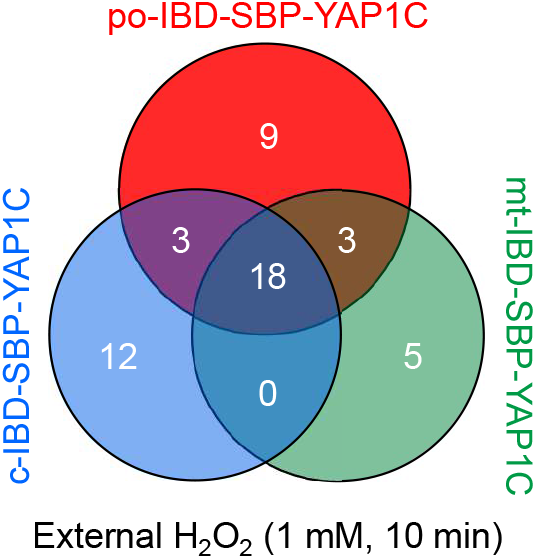
Venn diagram showing the number and overlap of po-, c-, and mt-IBD-SBP-YAP1C interactors upon treatment of po-DD-DAO Flp-In T-REx 293 cells with external H_2_O_2_.

To gain more insight into the binding selectivity of each IBD-SBP-YAP1C, we calculated the percentage distribution of each interactor with each of the YAP1C-fusion proteins (Table S2). In line with expectations, this analysis revealed that (i) po-IBD-SBP-YAP1C predominantly interacts with *bona fide* peroxisomal matrix proteins (e.g., ACOX1 [Yifrach et al., 2018]) or proteins that show a partial peroxisomal localization (e.g., LDHB [Schueren et al., 2014] and MDH1 [Hofhuis et al., 2016]), (ii) mt-IBD-SBP-YAP1C mainly interacts with genuine mitochondrial proteins (e.g., TXNRD2, ME2, ECI1, and HSD17B10 [Rath et al., 2021]), (iii) c-IBD-SBP-YAP1C preferentially interacts with proteins that are predominantly located in the cytosol (e.g., DNPEP, UCHL3, and TUBB1 [Thul et al., 2017]), and (iv) interactors located simultaneously in peroxisomes, mitochondria, and the cytosol (e.g., HSPA9 [Jo et al., 2020] and PRDX5 [Knoops et al., 2011]) are trapped by all YAP1C fusion proteins. However, diverse interactors known to be located in the cytosol and/or nucleus (e.g., BOLA2B, CSTB, HPRT1, PRDX1, PRDX2, PRDX6, PRMT5, SKP1, WDR77, and YWHAZ [Thul et al., 2017]) also bound to po- and mt-IBD-SBP-YAP1C, at least to a certain extent. This phenomenon can most likely be explained by the fact that a small but significant portion (rough estimation: 5-25%) of these IBD-SBP-YAP1C-fusion proteins still resides in the cytosol (Fig. 1). A similar reasoning can be applied with regard to the observation that a small portion of ACOX1 interacts with c-IBD-SBP-YAP1C. Indeed, it can be expected that the c-IBD-SBP-YAP1C-interacting portion of ACOX1 represents the protein pool that has not yet been imported into peroxisomes. Here, it is also important to note that the percentage distribution of some interactors displayed an unexpected behavior. For example, some cytosolically located target proteins preferentially bound to po-IBD-SBP-YAP1C (e.g., GSR, MARCKSL1, RPL13, RPL27A, SET, TXN, YWHAB, YWHAE, and YWHAQ) or mt-IBD-SBP-YAP1C (e.g., EPPK1, HSPB1, S100B, SERPINB4, and TXN). A comprehensive explanation is currently lacking, but it cannot be excluded that portions of these proteins are partially associated with the respective organelle. However, this remains to be further investigated.

Finally, this experiment clearly shows that some interactors (e.g., PRDX1, PRDX2, S100B, and TXN) are already partially trapped by their respective IBD-SBP-YAP1Cs even in the absence of H_2_O_2_ treatment. On one hand, this may point to background contamination. However, a more likely explanation is that these interactors are extremely sensitive to sulfenic acid formation, a hypothesis supported by their recognized roles in localized, rapid, specific, and reversible redox-regulated signaling events [Zhukova et al., 2004; Hanschmann et al., 2013].

### The cytosolic, mitochondrial, and peroxisomal sulfenome undergo time-dependent changes upon peroxisomal H_2_O_2_ generation

A similar experiment was performed with po-DD-DAO Flp-In T-REx 293 cells expressing compartment-specific variants of IBD-SBP-YAP1C and in which peroxisomal H_2_O_2_ production was induced or not by supplementing the assay medium with 10 mM D- or L-Ala, respectively. However, this time we included different time points (0, 2, 5, 15, 30, and 60 minutes) (Fig. S4) and adapted the LC-MS/MS method providing a higher sensitivity in MS/MS. After validation of the MS results, a total of 450 unique proteins that were two-fold or more enriched in at least one time point upon peroxisomal H_2_O_2_ production were retained. The number and overlap of interactors identified with po-, c-, or mt-IBD-SBP-YAP1C at each time point are visualized in Fig. 4. From this overview, it is clear that the number of individual and common interactors of different IBD-SBP-YAP1Cs changed in function of time. One may argue that these differences are due to variations in experimental handling. However, given that peroxisomal H_2_O_2_ generation resulted in different but consistent response profiles for distinct interactor classes, this assumption is unlikely to hold. A specific example is shown for c-IBP-SBP-YAP1C, which traps multiple peroxiredoxins (PRDXs) and 14-3-3 proteins upon such treatment (Fig. 5A,B). More examples can be found in Tables S3, S4, and S5, which respectively provide raw abundance heat maps of proteins trapped at different times by c-, po-, and mt-IBD-SBP-YAP1C in response to peroxisome-derived H_2_O_2_.

**Figure 4.**
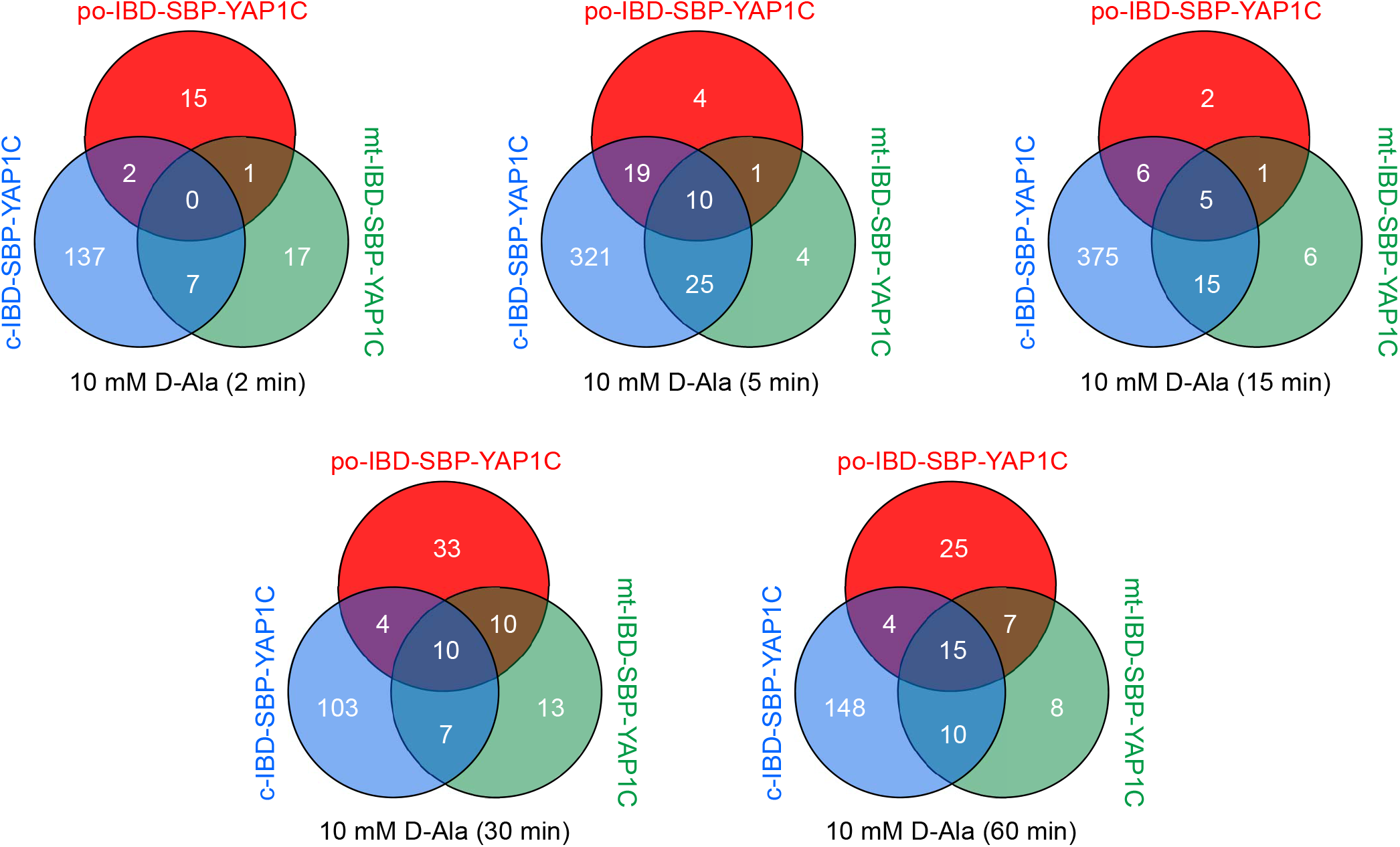
Venn diagrams showing the number and overlap of po-, c-, and mt-IBD-SBP-YAP1C interactors at different time points after treatment of po-DD-DAO Flp-In T-REx 293 cells with 10 mM D-Ala.

**Figure 5.**
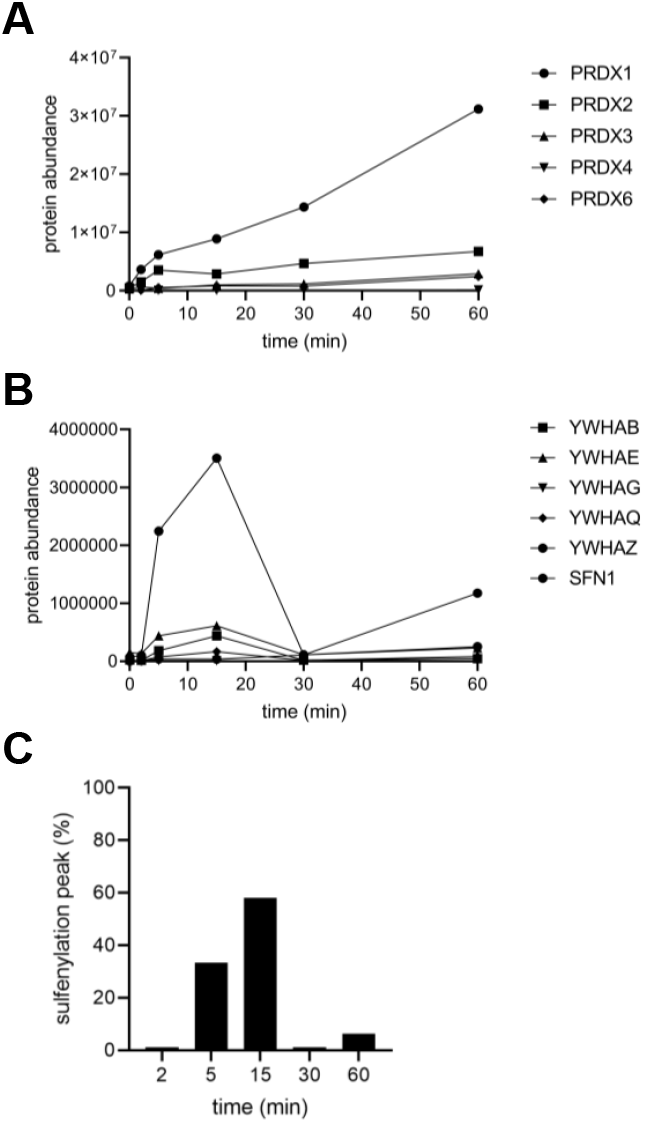
Sulfenylation kinetics of various c-IBD-SBP-YAPC1 interactors in response to peroxisomal H_2_O_2_ production. Po-DD-DAO Flp-In T-REx 293 cells expressing c-IBD-SBP-YAP1C were cultured in medium containing 1 μg/ml of DOX and 500 nM Shieldl. After 3 days, the medium was replaced with medium lacking DOX/Shield1. One day later, the cells were incubated in DPBS containing 10 mM 3-AT and 10 mM D-alanine. At selected time points (0, 2, 5, 15, 30, and 60 min), free thiol groups were blocked with NEM. Next, the YAP1C-containing protein complexes were affinity purified, and the c-IBD-SBP-YAP1C interaction partners were eluted with reducing agent and processed for LC-MS/MS analysis. **(A)** PRDX trapping by c-IBD-SBP-YAP1C (PRDX5 was not identified as hit). **(B)** 14-3-3 trapping by c-IBD-SBP-YAP1C (YWHAH was not identified as hit). **(C)** Percentage distribution of the sulfenylation peak time of all c-IBD-SBP-YAP1C interactors (n, 442).

Manual inspection of the 84 different po-IBD-SBP-YAP1C interactors (Table S4) surprisingly revealed the presence of only a few proteins that are frequently (e.g., CAT, HSD17B4, LDHA, and LDHB) [Yifrach et al., 2018] or sporadically (e.g., HSPA9) [Jo et al., 2020] detected in peroxisomes. This observation can potentially be explained in different ways. For example, given that the peroxisomal H_2_O_2_ sensor po-roGFP-ORP1 is already almost fully oxidized in Flp-In T-REx 293 cells under basal conditions [Lismont et al., 2019a], it may well be that no redox-sensitive protein thiol groups were left as target for oxidation by newly formed H_2_O_2_. On the other hand, it cannot be ruled out that the peroxisomal proteins trapped by po-IBD-SBP-YAP1C represent the cytosolic protein pools that have not yet been imported into peroxisomes. To gain more insight into this problem, we compared the response kinetics of CAT, HSD17B4, LDHA, LDHB, and HSPA9 to peroxisome-derived H_2_O_2_ in cells expressing po- or c-IBD-SBP-YAP1C (Fig. S5). From this figure, it is clear that (i) depending on the interactor and time point, the amount of protein trapped by po-IBD-SBP-YAP1C is up to 1.5-fold higher (e.g., CAT, 30 min) or up to 200-fold lower (e.g., HSD17B4, 15 min) than the amount trapped by c-IBD-SBP-YAP1C, and (ii) the sulfenylation profiles of all po-IBD-SBP-YAP1C (left panels) and some c-IBD-SBP-YAP1C (middle panels) interactors exhibit bimodal patterns, thereby reflecting a heterogeneous character of protein thiol oxidation (see Discussion). The observation that the capturing ratios of the peroxisomal targets by po- and c-IBD-SBP-YAP1C vary between different interactors (e.g., compare LDHA and LDHB at 15 min) and time points (e.g., compare CAT at 5 and 30 min), demonstrates that at least a portion of these complexes were formed inside peroxisomes. However, it is also clear that, at least for a number of these interactors (e.g., HSD17B4, LDHA, and HSPA9), complex formation inside peroxisomes is a rather negligible phenomenon compared to complex formation in the cytosol.

Unlike external H_2_O_2_ addition, peroxisomal H_2_O_2_ production did not result in a po-IBD-SBP-YAP1C-mediated trapping of ACOX1 or MDH1. Nonetheless, such treatment did result in the trapping of various cytosolic, cytoskeletal, and plasma membrane-associated proteins (Table S4), many of which are known to be sulfenylated on at least one cysteine residue [Wang et al., 2021]. Importantly, given that (i) we have previously demonstrated that peroxisome-derived H_2_O_2_ can efficiently permeate across the peroxisomal membrane [Lismont et al., 2019a], (ii) a small portion of po-IBD-SBP-YAP1C is mislocalized to the cytosol (Fig. 1), and (iii) the fraction of interactor bound to po-IBD-SBP-YAP1C at the peak of protein sulfenylation is on average less than 10% of the amount of interactor bound to c-IBD-SBP-YAP1C, it is safe to conclude that the majority of cytosolic (e.g., ENO1), cytoskeletal (e.g., POF1B), and plasma membrane-associated (e.g., ANXA2) interactors were trapped by the residual cytosolic fraction of po-IBD-SBP-YAP1C (Fig. S6). However, for some cytosolic interactors (e.g., APOA1, PRDX1, PRDX2, SKP1, and TXN), the amount of po-IBD-SBP-YAP1C-bound protein was much higher than what one would expect from the estimated percentage of mislocalized po-IBD-SBP-YAP1C (Fig. S7). Although it was not the scope of this work to dig into the subcellular localization of all these and other targets of peroxisome-derived H_2_O_2_, we decided to explore this intriguing observation in more detail for PRDX1. Upon immunoblot analysis of subcellular fractions derived from HEK-293 cells or rat liver, a small but significant portion of PRDX1 appears to be associated with peroxisomes (Fig. 6), thereby strengthening our working hypothesis.

**Figure 6.**
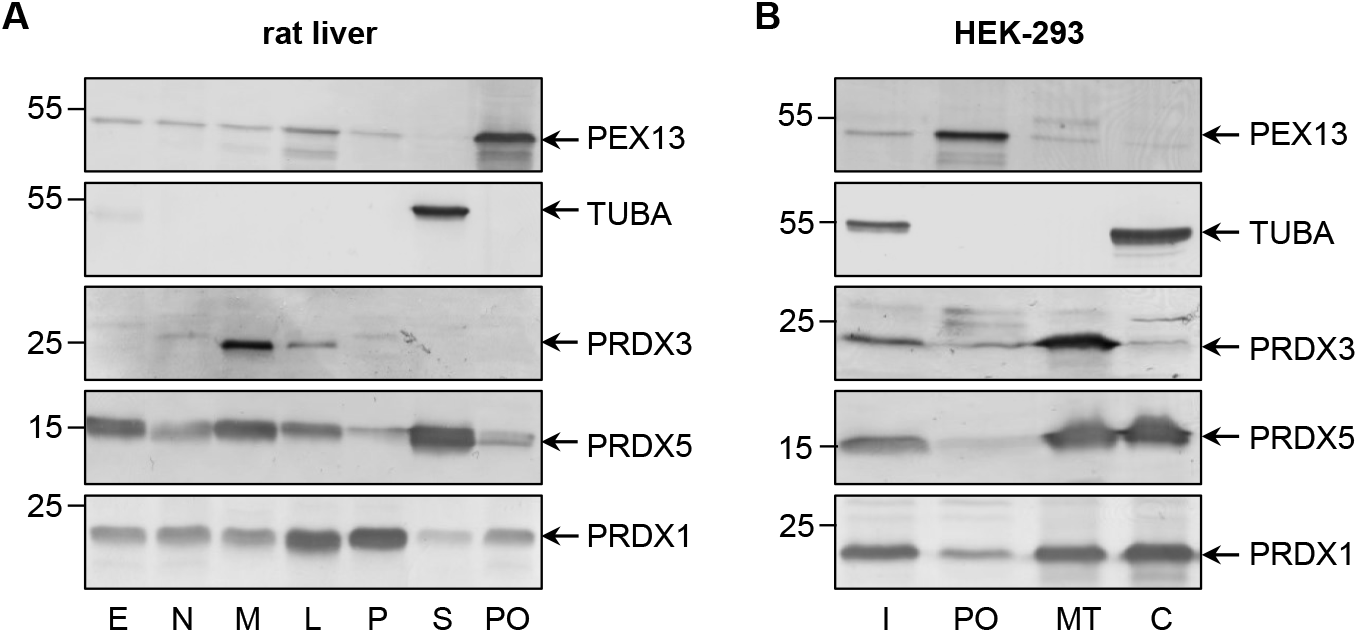
Subcellular distribution of PRDX1 in rat liver and HEK-293 cells. **(A)** Rat liver protein (20 μg) present in a post-nuclear fraction (E), a nuclear fraction (N), a heavy mitochondrial fraction (M), a light mitochondrial fraction (L), a microsomal fraction (P), the cytosol (S), or purified peroxisomes (PO) were subjected to SDS-PAGE and processed for immunoblotting with antibodies directed against PEX13 (peroxisomes), TUBA (cytosol), PRDX3 (mitochondria), PRDX5 (cytosol, mitochondria, nucleus, and peroxisomes), or PRDX1. **(B)** Total cell homogenates (T) from Flp-In T-REx 293 cells were fractionated by differential centrifugation to yield a 1500 x g supernatant (Input, I) that was subsequently subjected to Nycodenz gradient centrifugation [Lismont et al., 2019a]. The I fraction (8% of the amount loaded onto the gradient) and equal volumes of gradient fractions enriched in peroxisomes (PO), mitochondria (MT), or the cytosol (C) were processed for immunoblot analysis as described for the rat liver fractions. The migration points of relevant molecular mass markers (expressed in kDa) are shown on the left. Specific proteins are marked by arrows.

A careful examination of the 56 different mt-IBD-SBP-YAP1C interactors (Table S5) revealed the presence of 12 proteins (ATP5F1A, GSR, HSDB4, HSP90AA1, HSPA9, HSPD1, PC, PCCA, PCCB, PPP2R2B, PRDX3, and VDAC1) with a *bona fide* (partial) mitochondrial localization [Rath et al., 2021], thereby providing direct molecular evidence for the previously established peroxisome-mitochondria redox connection [Lismont et al., 2015]. As observed for po-IBD-SBP-YAP1C (Table S4; Fig. S6), mt-IBD-SBP-YAP1C also captured some cytosolic (e.g., ENO1), cytoskeletal (e.g., POF1B), and plasma membrane-associated (e.g., ANXA2) interactors (Fig. S8). This observation strengthens our prior interpretation (Table S2) that also a small portion of mt-IBD-SBP-YAP1C is not yet imported into mitochondria. For comparison, we also included PRDX3, a mitochondrial member of the PRDX family (Fig. S8).

We also identified a subset of 442 different c-IBD-SBP-YAP1C interactors (Table S3). KEGG pathway analysis revealed an enrichment for proteins implicated, among others, in ribosome biology, carbon metabolism, biosynthesis of amino acids, and proteasome functioning (the -log10(padj) values are 4.118 x 10^-28^, 3.754 x 10^-9^, 4.792 x 10^-6^, and 2.294 x 10^-5^, respectively). Surprisingly, despite the fact that it is well known that H_2_O_2_ can, directly or indirectly, oxidatively modify different classes of proteins involved in signal transduction (e.g., kinases, phosphatases, proteases, antioxidant enzymes, transcription factors, etc.), no transcription factors were found to interact with c-IBD-SBP-YAP1C. One potential explanation for this finding is that members belonging to this group of proteins are oxidatively modified via redox relay, and not through direct oxidation of redox-sensitive cysteines to sulfenic acid (see Discussion). Another interesting observation is that, with the exception of proteins involved in the maintenance of cellular redox homeostasis (e.g., GSR, PRDX1, PRDX2, PRDX3, PRDX4, PRDX6, TXN, and ERP44), virtually all other interactors display a sulfenylation peak time at 5 or 15 min (Fig. 5C). In addition, proteins belonging to the same protein family share in general a common pattern (e.g., compare the peak time values in Figs. 5 and S9). Protein families that, besides the PRDXs (Fig. 5A) and 14-3-3 (Fig. 5B), are abundantly modified by peroxisome-derived H_2_O_2_ include constituents of the cytoskeleton (Fig. S9A,B), annexins (Fig. S9C), protein chaperones (Fig. S9D), S100 proteins (Fig. S9E), and negative regulators of endopeptidase activity (Fig. S9F). Finally, it is worth noting that some c-IBD-SBP-YAP1C interactors (e.g., ANXA2, DSG1, DSP1, FLG2, GAPDH, and JUP) already appear to be considerably sulfenylated under basal conditions (Table S3). Once again, this observation strongly supports a role for these proteins in redox-regulated housekeeping signaling pathways.

### The cytosolic sulfenome responds to exogenous H_2_O_2_ in a dose-dependent manner

In a laboratory setting, (patho)physiological scenarios of how H_2_O_2_ can drive cellular signaling events are often mimicked by external addition of this oxidant to cultured cells. In order to assess how H_2_O_2_ levels and sulfenylation responses in our DD-DAO-based approach compare to this often used strategy, we treated cells with different concentrations of H_2_O_2_, ranging between 10 μM and 1 mM, for 10 minutes (Fig. S10). Using cells expressing c-IBD-SBP-YAP1C, we identified 375 proteins that were two-fold or more enriched in at least one of the treated conditions (Table S6). Note that, due to the increased sensitivity settings in this LC-MS/MS run, the amount of hits greatly exceeded the number of targets identified in the initial validation experiment (Table S1).

The number and overlap of interactors identified upon treatment of the cells with different H_2_O_2_ concentrations are visualized in Fig. 7A. From these data, it can be deduced that treatment with 100 μM H_2_O_2_ yields the highest number of sulfenylated proteins. This can be explained by the fact that, at this concentration of H_2_O_2_, 60% of all targets reach their sulfenylation peak (Fig. 7B), including the antioxidant defense enzymes (Table S6) that did not even reach their peak within 1 h of peroxisomal H_2_O_2_ production (Tables S3). Interestingly, PRDX5 is the only peroxiredoxin that was exclusively sulfenylated by externally added H_2_O_2_ (Table S6). This may be explained because (i) PRDX5 has the lowest expression level of all PRDXs in HEK-293 cells [Geiger et al., 2012] and (ii) its preferred substrates are lipid peroxides instead of hydrogen peroxide [Knoops et al., 2011]. Given that PRDX5 is already more than two-fold enriched from 10 μM H_2_O_2_ onwards, peroxisomal H_2_O_2_ production by DD-DAO is far less stringent than external addition of 10 μM H_2_O_2_. Taking into consideration that (i) data in literature suggest a 390-to 650-fold concentration difference between the extra- and intracellular H_2_O_2_ levels [Huang et al., 2014; Lyublinska and Antunes, 2019], which would result in 15 to 26 nM intracellular H_2_O_2_ upon external addition of 10 μM H_2_O_2_, (ii) within the time frame of the experiment (60 min), the equilibrium between peroxisomal H_2_O_2_ production by DD-DAO and the cell’s antioxidant defense mechanisms has not yet been reached (as demonstrated by the fact that antioxidant enzymes did not reach their sulfenylation peak), and (iii) in our experiments, the intracellular H_2_O_2_ levels may be slightly higher due to the presence of the catalase inhibitor 3-amino-1,2,4-triazole (3-AT), we estimate that DD-DAO activity results in (cytosolic) H_2_O_2_ concentrations ranging between 5 and 50 nM. Here, it is also worth mentioning that there is relatively little overlap between the protein targets of peroxisome-derived and externally added H_2_O_2_ (Fig. 8A). When comparing the sulfenylation profiles upon addition of external H_2_O_2_ (concentration curve) or peroxisomal H_2_O_2_ production (time curve), it is clear that depending on the H_2_O_2_ source, different proteins show distinct responses. A set of examples is shown in Fig. 8B-J. All together, these findings highlight that data obtained with external H_2_O_2_ cannot simply be extrapolated to internally produced H_2_O_2_, which reflects a more physiological condition.

**Figure 7.**
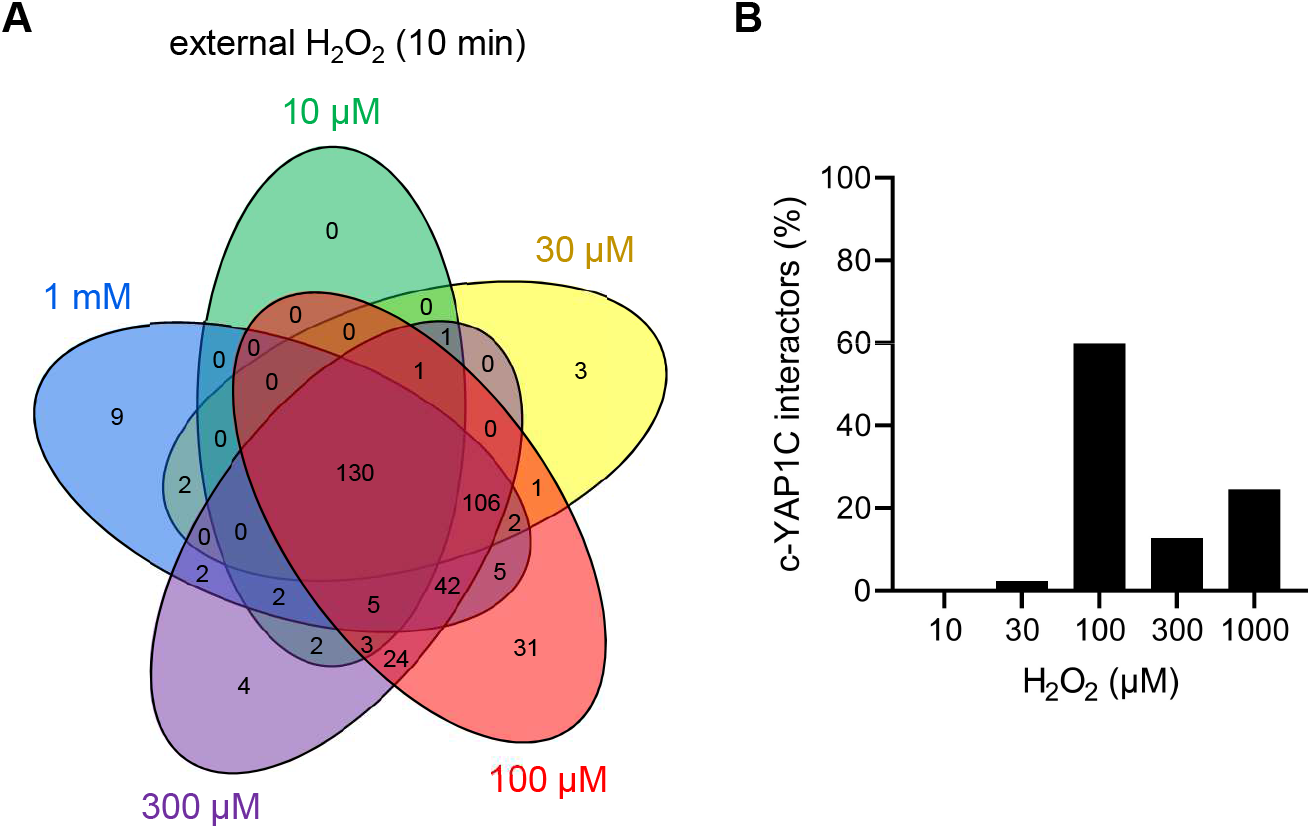
The cytosolic sulfenome in response to different concentrations of extracellular H_2_O_2_. **(A)** Venn diagram showing the number and overlap of significantly enriched interactors of c-IBD-SBP-YAP1C upon treatment of po-DD-DAO Flp-In T-REx 293 cells with different concentrations of external H_2_O_2_ for 10 min. **(B)** Percentage distribution of total c-IBD-SBP-YAP1C (c-YAP1C) interactors (n, 375) with a sulfenylation peak at the indicated H_2_O_2_ concentration.

Strikingly, whereas upon peroxisomal H_2_O_2_ production the sulfenylation peak rapidly decreases to baseline levels for most proteins, treatment with external H_2_O_2_ most often results in a slower and rather modest decrease (Fig. 8B-J, compare the blue and red profiles). Given that we have previously shown that po-DD-DAO-mediated H_2_O_2_ production results in steadily increasing levels of cytosolic H_2_O_2_ and disulfide bond formation over time [Lismont et al., 2019a], a potential explanation is that the fast decrease in sulfenylation observed under this condition represents a combined effect of (i) a thioredoxin (TXN)-mediated reduction of the disulfide bond between IBD-SBP-YAP1C and its interactors [Izawa et al., 1999], and (ii) a continuous depletion of freely available redox-sensitive cysteine thiols (e.g., due to overoxidation or disulfide bond formation with other proteins). Importantly, our observation that under conditions of oxidative stress transient disulfide bond formation between HSPB1 and c-IBD-SBP-YAP1C (Fig. 9A) precedes previously reported [Zavialov et al., 1998] disulfide-mediated changes in the oligomeric state of HSPB1 (Fig. 9B), is in line with this view. On the other hand, upon external H_2_O_2_ addition, the modest decrease in sulfenylation after peaking may be explained by reduced availability of TXN and other thiol-disulfide reductases as a consequence of their overoxidation, a phenomenon supported by the observation that also these enzymes themselves are less sulfenylated at these H_2_O_2_ concentrations (Fig. 8H-J). Finally, the observation that proteins can be released from c-IBD-SBP-YAP1C, underscores the transient nature of these disulfide bonds, even in an oxidizing environment.

**Figure 8.**
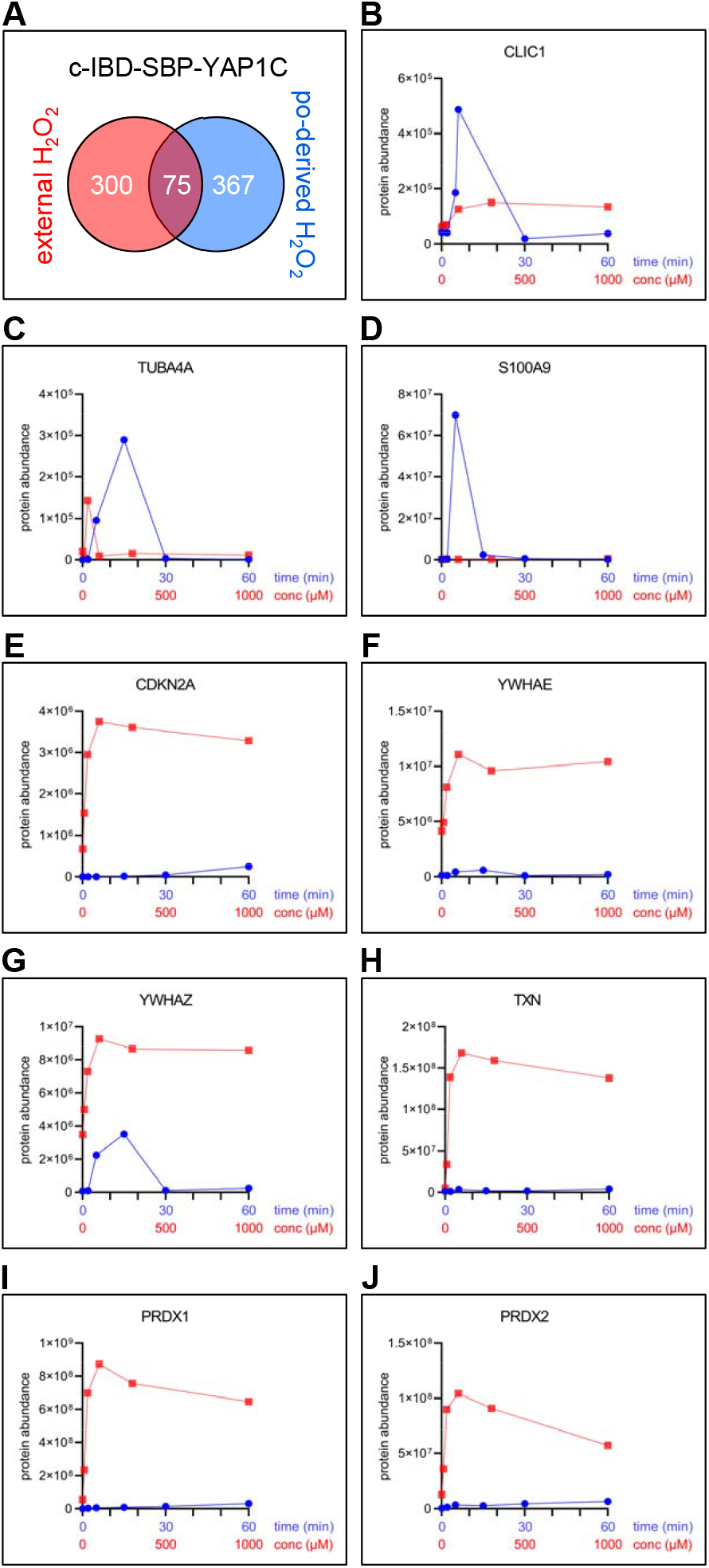
The pool and sulfenylation profiles of c-IBD-SBP-YAP1C interactors differ depending on the H_2_O_2_ source. **(A)** Venn diagram showing the number and overlap of significantly enriched interactors of c-IBD-SBP-YAP1C upon treatment of po-DD-DAO FlpIn T-REx 293 cells with 10 mM D-Ala (Table S3) or 1 mM H_2_O_2_ for 10 min (Table S6). **(B-J)** Sulfenylation profiles of a selected set of c-IBD-SBP-YAP1C interactors upon peroxisomal H_2_O_2_ production (time curve; in blue) or addition of external H_2_O_2_ for 10 min (concentration curve; in red). Protein abundances are based on peptides with a PEP < 0.05 that were commonly retrieved in the experiments shown.

**Figure 9.**
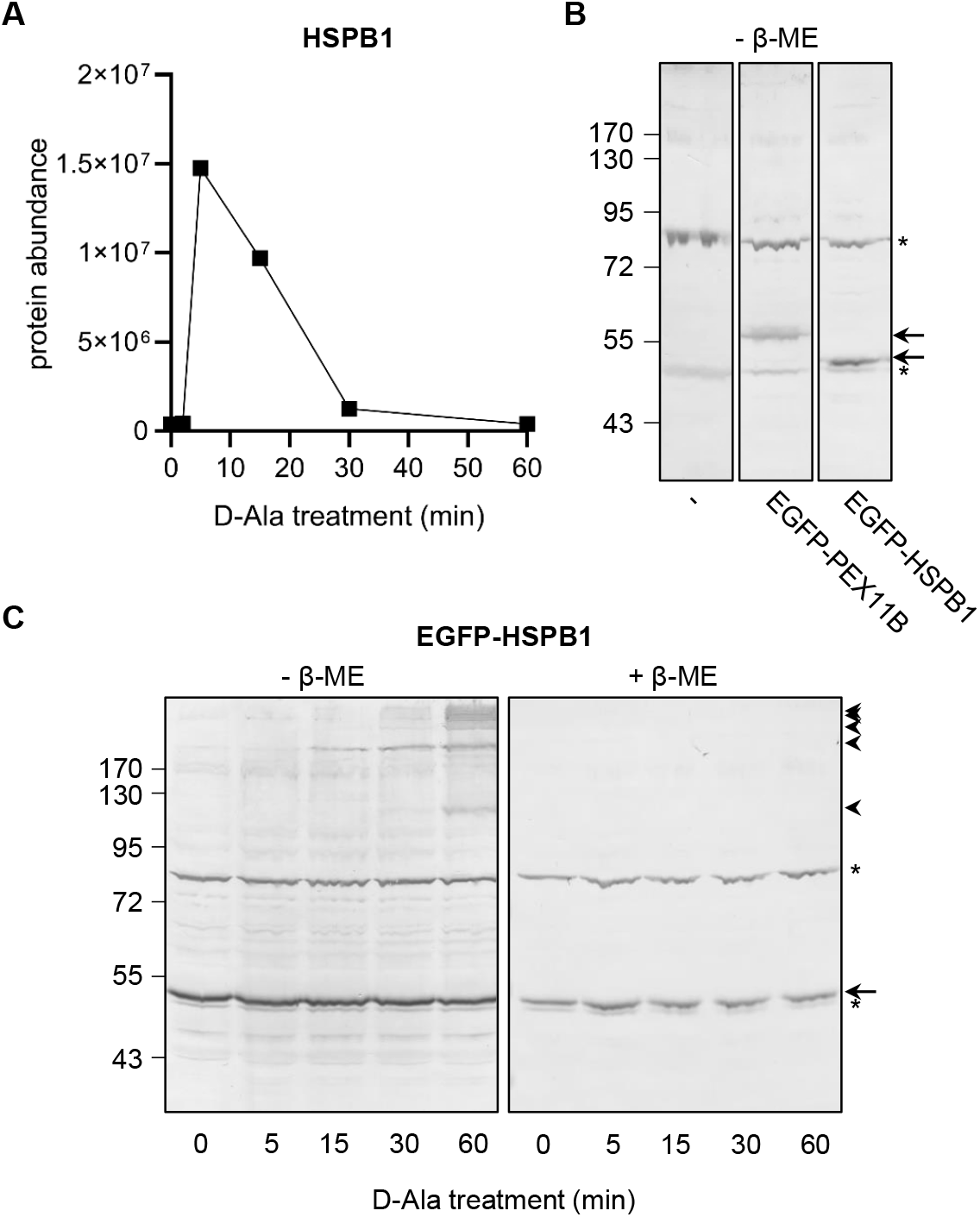
Sulfenylation and disulfide bond formation kinetics of HSPB1 in response to peroxisomal H_2_O_2_ production. Po-DD-DAO Flp-In T-REx 293 cells expressing c-IBD-SBP-YAP1C **(A)** or not **(B, C)** were cultured in medium containing 1 μg/ml of DOX and 500 nM Shield1. After 3 days, the medium was replaced with medium lacking DOX/Shield1, and the cells were transfected **(B, C)** or not **(A)** with a plasmid encoding no EGFP-fusion protein (-), EGFP-PEX11B, or EGFP-HSPB1. One day later, the cells were incubated in DPBS containing 10 mM 3-AT and 10 mM D-alanine (D-Ala). At the indicated time points, free thiol groups were blocked with NEM, and **(A)** the YAP1C-containing protein complexes were affinity purified and processed for LC-MS/MS, or **(B, C)** the cells were processed for SDS-PAGE under non-reducing (-β-ME) and reducing (+β-ME) conditions and subsequently subjected to immunoblot analysis with antibodies specific for EGFP. The migration points of relevant molecular mass markers (expressed in kDa) are shown on the left. The arrows and arrowheads mark the non-modified and oxidatively modified proteins, respectively. Note that panel B was included to document the non-specific immunoreactive bands (marked by asterisks) of the anti-EGFP antiserum.

### Assessment of potential pitfalls

The findings presented thus far clearly demonstrate that the YAP1C-based sulfenome mining approach is a very powerful and efficient tool to identify targets of peroxisome-derived H_2_O_2_. As mentioned above and described elsewhere [Lismont et al., 2019a], we used a po-DD-DAO/IBD-SBP-YAP1C Flp-In T-REx 293 cell line in which the expression levels and stability of DD-DAO can be strictly controlled to overcome possible interfering effects of H_2_O_2_ produced by newly synthesized DD-DAO that has not yet been imported into peroxisomes. To document the validity of this assumption at the proteome level, we also performed a sulfenome mining experiment with c-DD-DAO Flp-In T-REx 293 cells expressing c-IBD-SBP-YAP1C (Fig. S11). Importantly, even in a condition where initially 100 % of DD-DAO was located in the cytosol, only 16 out of the 274 identified targets were enriched in the chase condition (Fig. 10A; Table S7). In addition, the enrichment values of these 16 hits only slightly exceeded the threshold value of 2 (avg/stdev: 2.95/1.28), which is much lower than the enrichment values (avg/stdev: 12.00/29.67) of the targets identified in the no-chase condition. Furthermore, to gain more insight into target proteins that may be indirectly retained on the affinity matrix through electrostatic interaction with truly sulfenylated proteins [Liebthal et al., 2020], Flp-In T-REx 293 cells expressing c-IBD-SBP-YAP1C were treated with 1 mM H_2_O_2_ for 10 minutes and, after enrichment of the corresponding c-IBD-SBP-YAP1C complexes and completing the normal washing procedure, the streptavidin column was first 3 times eluted with high salt (1 M NaCl) and subsequently 3 times with DTT (this experiment was done in parallel with the experiment shown in Fig. S10). LC-MS/MS analysis of the eluates revealed the presence of 262 distinct interactors, of which only 6 predominantly eluted in a NaCl-dependent manner (Fig. 10B; Table S8). In summary, these findings confirm that the vast majority of targets identified are *bona fide* sulfenylated proteins, thereby demonstrating that the employed procedure is very robust.

**Figure 10.**
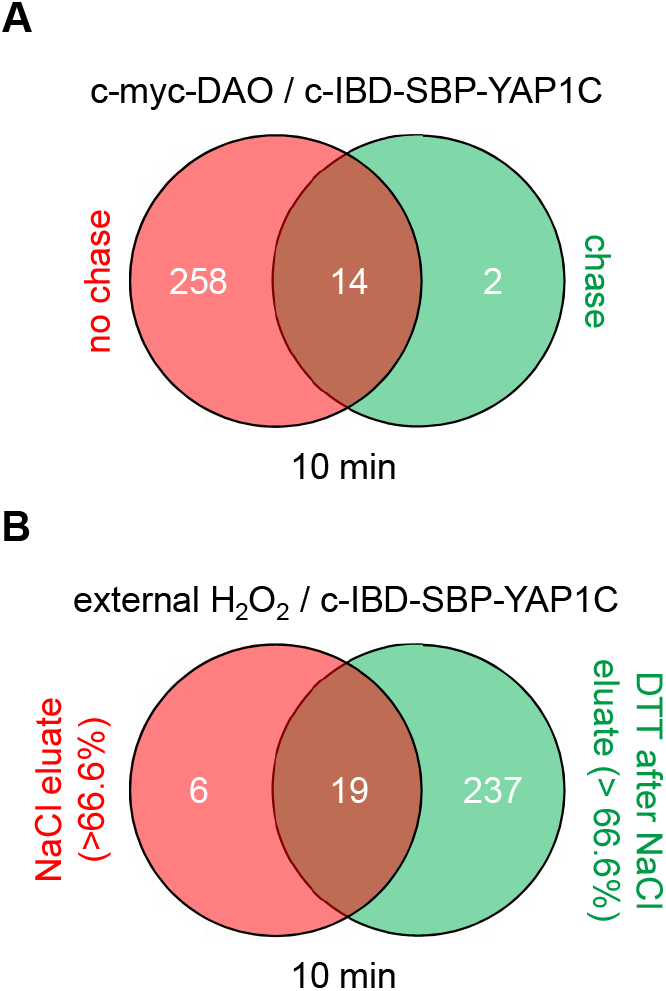
Venn diagrams showing the number and overlap of c-IBD-SBP-YAP1C interactors under different experimental conditions. **(A)** c-DD-DAO-driven H_2_O_2_ production in the absence or presence of a 24-h chase period. **(B)** External H_2_O_2_ addition and elution of the interactors first with 1 M NaCl and subsequently with 10 mM DTT. The threshold values to belong to a specific group reflect the percentage of protein recovered in each fraction relative to the total protein abundance (red: > 66.6% in the NaCl eluate; green: > 66.6% in the DTT after NaCl eluate; brown: less than 66.6% in both eluates).

### Identification of oxidatively modified cysteines

In our protocol, free and oxidatively modified cysteines are differentially alkylated (Fig. 11), providing more information on which cysteines are oxidized. In short, after cell treatment, free thiols are irreversibly blocked with NEM, cells are lysed, and IBD-SBP-YAP1C-containing protein complexes are enriched on the streptavidin affinity matrix. Upon DTT elution, the originally sulfenylated cysteines captured by IBD-SBP-YAP1C are reduced and become available for iodoacetamide alkylation, thereby resulting in the formation of an S-carbamidomethyl cysteine. As LC-MS/MS analysis can distinguish between different cysteine modifications, it is possible to determine at what stage in the protocol the cysteines of the identified peptides were alkylated. Here, it is essential to point out that S-carbamidomethylation does not necessarily indicate that the cysteine was indeed sulfenylated, since S-carbamidomethylated cysteines can also result from pre-existing disulfide bridges or cysteines that are inaccessible to NEM. In addition, as LC-MS/MS analysis does not detect every peptide of a protein, the detection of redox-sensitive cysteine-containing peptides is no absolute requirement to categorize a hit as an authentically sulfenylated protein. In the 841 IBD-SBP-YAP1C-interactors that were identified during the time course of our experiments, we could detect 167 proteins containing at least one S-carbamidomethyl cysteine (Table S9). As could be expected, the number of detected S-carbamidomethylated cysteines is determined by the sensitivity settings of the LC-MS/MS run as well as by the stringency of the treatment which intrinsically correlates with peptide abundance (e.g., treatment of cells with external H_2_O_2_ resulted in more S-carbamidomethyl cysteines than H_2_O_2_ production inside peroxisomes). A comparative analysis of our data with the list of proteins in the iCysMod database revealed that, of the 319 redox-sensitive cysteines identified, 89 cysteines have already been reported to be sulfenylated by others, 163 cysteines were already shown to undergo other oxidative thiol modifications, and 67 cysteines represent novel redox-regulated cysteines (Table S9).

**Figure 11.**
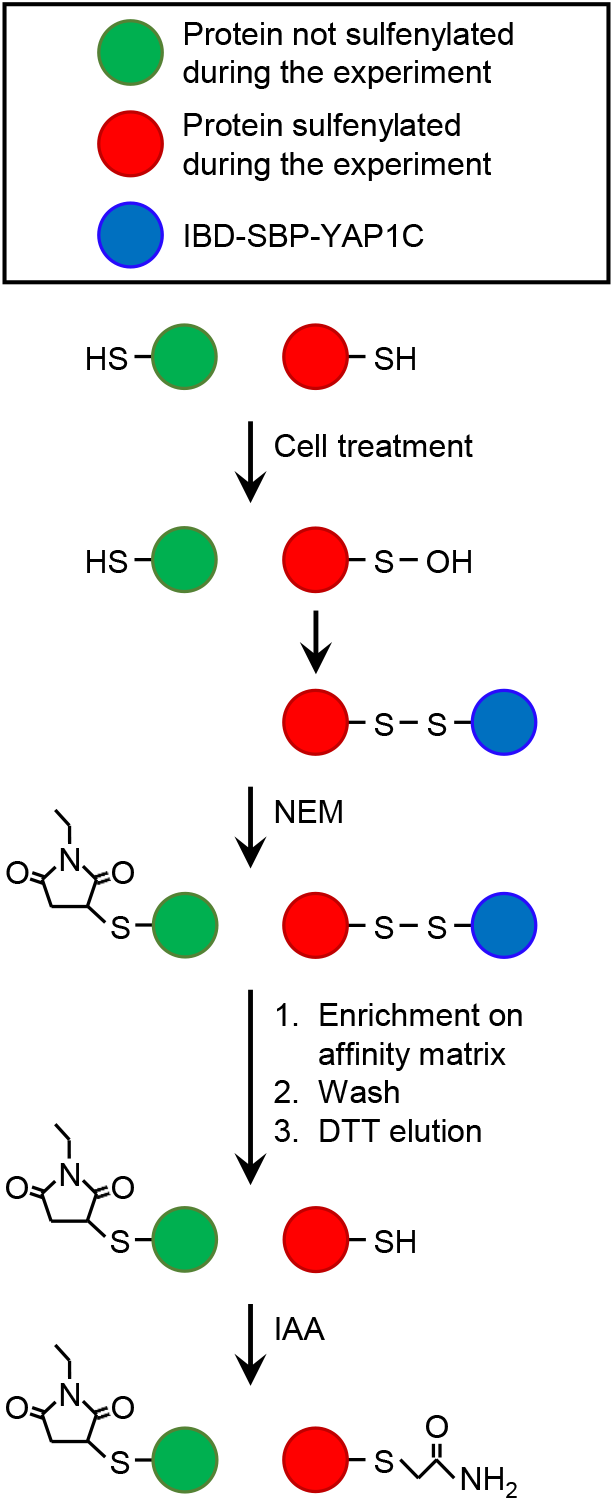
Differential alkylation of cysteine thiols and sulfenic acids. After cell treatment, free thiols are immediately blocked with N-ethyl maleimide (NEM). Proteins containing at least one sulfenylated cysteine that forms a disulfide bridge with IBD-SBP-YAP1C are enriched on the streptavidin affinity matrix, extensively washed, eluted with dithiothreitol (DTT), and alkylated with iodoacetamide (IAA). This results in a differential alkylation of originally reduced or sulfenylated cysteines within the affinity-purified proteins. For more details about the experimental procedure, see Materials and Methods.

## Discussion

Currently, it is widely accepted that peroxisomes can act as H_2_O_2_ signaling platforms, thereby conveying metabolic information into redox signaling events [Fransen and Lismont, 2019; He et al., 2021]. However, little is known about the molecular targets of peroxisomal H_2_O_2_ and, to address this gap, we designed an efficient and unique sulfenome mining approach to capture and identify such targets in a dynamic manner. From our results, it is clear that peroxisome-derived H_2_O_2_ can trigger cysteine oxidation in multiple members of various protein families, including but not limited to antioxidant enzymes, constituents of the cytoskeleton, protein chaperones, annexins, and 14-3-3 and S100 proteins (Tables S3-5). Given that many of these proteins are on the crossroads of key cellular processes [Rubio et al., 2004; Wu et al., 2021; Méndez-Barbero et al., 2021], such as carbon metabolism, protein synthesis and folding, proteasome functioning, and calcium signaling, it can be expected that genetic-, age-, and environment-related changes in peroxisomal H_2_O_2_ metabolism also contribute to disease pathogenesis.

This is perhaps best exemplified by the observations that (i) inherited catalase deficiency is associated with oxidative stress-related disorders, such as neoplasms, atherosclerosis, and diabetes [Góth and Nagy, 2013], and (ii) a gain-of-function mutation (N237S) in acyl-CoA oxidase 1 (ACOX1), a peroxisomal enzyme that oxidizes very-long-chain fatty acids and produces H_2_O_2_ as a byproduct, causes oxidative damage associated with severe Schwann cell loss and myelination defects in humans [Chung et al., 2020].

Our data also show that the IBD-SBP-YAP1C interactome differs considerably depending on the subcellular location of the YAP1C fusion protein (Fig. 3), the duration of the oxidative insult (Fig. 4), the H_2_O_2_ concentration (Fig. 7A), and the source of the oxidant (Fig. 8A). In addition, it is important to keep in mind that the capturing rate of sulfenylated proteins by specific YAP1C-fusion proteins will be influenced by other factors, such as (i) the local concentrations of the bait and target proteins, (ii) the import efficiencies of these protein into their organelle of destination, (iii) the local H_2_O_2_ levels, and (iv) the basal oxidation state of the redox-sensitive cysteines within the target protein. Notably, given that IBD-SBP-YAP1C captures only S-sulfenylated proteins, our findings do only shed light on the peroxisome sulfenome, but not on the full sulfur redoxome. That is, proteins that are oxidatively modified via redox relay, and not through direct oxidation by H_2_O_2_, will remain undetected. Whether or not this underlies the absence of transcription factors in our target list, remains to be investigated. Another limitation of our approach is that the molecular mass of IBD-SBP-YAP1C is relatively high (30 kDa) (Fig. S1). Indeed, given that most cysteines are buried inside proteins or protein complexes [Poole, 2015], the corresponding sulfenic acids may not be accessible for IBD-SBP-YAP1C due to steric hindrance, thereby resulting in an underestimation of targets. Finally, another consideration is that IBD-SBP-YAP1C complexes can be reduced *in cellulo* (Fig. 5C). Although this may seem surprising at first sight, it makes sense given that otherwise IBD-SBP-YAP1C expression would cause severe toxicity due to irreversible trapping of thiol redox signaling proteins (e.g., TXN and PRDX1) that are already significantly sulfenylated under basal conditions.

To provide a deeper understanding of how intracellular cysteine redox networks are regulated in response to various H_2_O_2_ insults, it is crucial to monitor the sulfenylation dynamics at the level of individual proteins. From our experiments, it is evident that the cytosolic and mitochondrial sulfenome undergo time-dependent changes upon peroxisomal H_2_O_2_ production (Fig. 4). Importantly, the latter observation provides direct molecular evidence for the previously established peroxisome-mitochondria redox connection [Lismont et al., 2019b]. In addition, it is crystal clear that the sulfenylation profiles can vary considerably between proteins (Fig. S7) and the type of H_2_O_2_ treatment (Fig. 8). However, in general, members of the same protein family exhibit a similar response behavior (Figs. 5 and S9). Also, as pointed out in the Results section, the sulfenylation profiles of all (partially) peroxisomal po-IBD-SBP-YAP1C interactors exhibited a bimodal pattern (Fig. S5). This heterogeneous character likely reflects a combination of factors, including (i) sulfenylation of multiple cysteine residues with distinct redox sensitivity, (ii) the bimodal localization of po-IBD-SBP-YAP1C, and/or (iii) time-dependent changes in local H_2_O_2_ levels. Finally, from the multiple lists of IBD-SBP-YAP1C interactors (Tables S1,S3-S7) and the sulfenylation profiles of individual targets (Fig. 8), it can be concluded that findings obtained with external H_2_O_2_, even at concentrations as low as 10 μM, cannot simply be extrapolated to conditions in which this oxidant is produced endogenously (e.g., inside peroxisomes or the cytosol). Here, it is important to mention that we routinely included 10 mM 3-AT in our assay buffer to inhibit catalase activity. However, during the course of our experiments, we obtained immunoblot and proteomics data documenting that a 15 min D-Ala treatment of po-DD-DAO expressing Flp-in T-REx 293 cells in the absence of 3-AT yields a result comparable to a 2-5 min D-Ala treatment in the presence of 3-AT (data not shown).

Another intriguing aspect of this study is that the employed sulfenome mining approach can apparently also provide more insight into the potential subcellular localization of the IBD-SBP-YAP1C interactors. One example that we studied in more detail is PRDX1, a predominantly cytosolic and nuclear protein that functions as antioxidant enzyme and protein chaperone under oxidative distress conditions [Rhee and Woo, 2020]. Given that the amount of PRDX1 captured by po-IBD-SBP-YAP1C was much higher than what one would expect from the estimated amount of the mislocalized bait protein (Fig. S7), our data suggested that this protein was also present in peroxisomes, a finding strengthened by subcellular fraction studies on HEK-293 cells and rat liver (Fig. 6). Note that, in contrast to mitochondria, typical 2-Cys PRDXs and TXNs have not yet been identified in mammalian peroxisomes [Lismont et al., 2015]. In this context, it is worth noting that, at least according to our sulfenome mining data (Fig. S7), also PRDX2 and TXN are partially located inside peroxisomes. However, it is not the scope of this work to dig into the subcellular localization of all these and other targets of peroxisome-derived H_2_O_2_, but this information may stimulate research efforts on how peroxisomes regulate their intraorganellar redox state.

Finally, some interesting open questions that arise from this work but need to be addressed in more detail in future studies include: Why are some peroxisomal proteins (e.g., HSD17B4) more efficiently enriched by c-IBD-SBP-YAP1C than by po-IBD-SBP-YAP1C, and what is the molecular explanation for this observation and the finding that only very few *bona fide* peroxisomal proteins are captured by po-IBD-SBP-YAP1C? Why are transcription factors lacking in our sulfenome lists? Are non-peroxisomal proteins that are captured by po-IBD-SBP-YAP1C in aberrantly high levels partially localized in peroxisomes, and if so how are they targeted to this organelle? Are the mitochondrial proteins captured by mt-IBD-SBP-YAP1C directly sulfenylated by H_2_O_2_ that diffuses out of peroxisomes or does peroxisome-derived H_2_O_2_ function as a second messenger that activates ROS-induced ROS release in neighboring mitochondria?

## Conclusions

In this study, we provide a first snapshot view of the HEK-293 sulfenome in response to peroxisome-derived or externally added H_2_O_2_. Our approach distinguishes between targets in different cellular locations and allows, under ideal conditions, to identify the transiently sulfenylated cysteines within target proteins. The outcome of these experiments revealed a previously unexplored role of peroxisomes in diverse redox-regulated processes, including but not limited to cytoskeletal remodeling, calcium signaling, and protein synthesis and turnover. As such, this study opens new perspectives for research on how perturbations in peroxisomal H_2_O_2_ metabolism may contribute to the initiation and development of redox stress-related diseases.

## Acknowledgments

We thank Dr. Sebastien Carpentier for interesting discussions about MS data analysis, and Prof. Dr. Joris Messens (VUB, Belgium), Dr. Daria Ezerina (VUB, Belgium), and Dr. Jesalyn Bolduc (VUB, Belgium) for their valuable discussions and comments regarding the YAP1C-based sulfenome mining data. We are also grateful to Prof. Dr. Ludo Van Den Bosch for providing us with the EGFP-HSPB1 plasmid.

## Legends to Supplementary Figures

**Figure S1. DNA and protein sequence of human codon-optimized IBD-SBP-YAP1C.** The start and stop codons are indicated in bold. The restriction sites for EcoRI (gaattc) and NotI (gcggccgc) are shaded in grey. The IgG-binding domains (IBD), the streptavidin-binding domain (SBP), and the YAP1 C-terminal cysteine-rich domain (YAP1C) are shaded in yellow, blue, and black, respectively. The redox-active cysteine is shaded in green, and the amino acids shaded in red indicate positions where cysteines were replaced by alanine and threonine residues, respectively.

**Figure S2. Differentially-localized IBD-SBP-YAP1C proteins form different complexes upon treatment of cells with external H_2_O_2_.** Po-DD-DAO Flp-In T-REx 293 cells expressing c-, mt-, or po-IBD-SBP-YAP1C were harvested and resuspended in DPBS containing 10 mM 3-AT, in combination (H_2_O_2_) or not (DPBS) with 1 mM H_2_O_2_. After 10 minutes, free thiol groups were blocked with NEM. Next, the YAP1C-containing protein complexes were enriched on streptavidin beads and the c-YAP1C interaction partners were eluted with DTT. The samples were processed for SDS-PAGE under non-reducing conditions and then subjected to immunoblot analysis with rabbit pre-immune serum. The migration points of relevant molecular mass markers (expressed in kDa) are shown on the left. The arrow indicates the non-oxidatively modified YAP1C fusion proteins. I, input; NB, not bound to beads; B, bound to beads; BE, bound to beads after DTT elution. The DTT eluates were further processed for LC-MS/MS analysis (see Table S1).

**Figure S3. Functional enrichment analysis of IBD-SBP-YAP1C interactors upon treatment of po-DD-DAO Flp-In T-REx 293 cells with 1 mM H_2_O_2_ for 10 min**. The analysis was performed using g:GOST (https://biit.cs.ut.ee/gprofiler/gost) within the g:Profiler tool. The colored blocks below the interactors refer to different evidence codes. BP, biological process.

**Figure S4. Formation kinetics of c-, mt-, and po-IBD-SBP-YAP1C complexes in response to peroxisomal H_2_O_2_ production.** Po-DD-DAO Flp-In T-REx 293 cells expressing c-, mt-, or po-IBD-SBP-YAP1C were cultured in medium containing 1μ g/ml of DOX and 500 nM Shield1. After 3 days, the medium was replaced with medium lacking DOX/Shield1. One day later, the cells were incubated in DPBS containing 10 mM 3-AT and 10 mM D-alanine. At selected time points (0, 2, 5, 15, 30, and 60 min), free thiol groups were blocked with NEM. Next, the IBD-SBP-YAP1C-containing protein complexes were affinity purified and the IBD-SBP-YAP1C interaction partners were eluted with DTT. The samples were processed for SDS-PAGE under non-reducing conditions and then subjected to immunoblot analysis with rabbit pre-immune serum. The migration points of relevant molecular mass markers (expressed in kDa) are shown on the left. The arrow and arrowheads mark the non-modified and oxidatively modified IBD-SBP-YAP1C complexes, respectively. Non-specific bands are indicated with asterisks. I, input; NB, not bound to beads; B, bound to beads; BE, bound to beads after DTT elution.

**Figure S5. Sulfenome profiles of peroxisome-targeted po-IBD-SBP-YAP1C interactors in response to peroxisomal H_2_O_2_ production.** Po-DD-DAO Flp-In T-REx 293 cells expressing po- or c-IBD-SBP-YAP1C were cultured in medium containing 1 μg/ml of DOX and 500 nM Shield1. After 3 days, the medium was replaced with medium lacking DOX/Shield1. One day later, the cells were incubated in DPBS containing 10 mM 3-AT and 10 mM D-alanine. At selected time points (0, 2, 5, 15, 30, and 60 min), free thiol groups were blocked with NEM. Next, the YAP1C-containing protein complexes were affinity purified, the IBD-SBP-YAP1C interaction partners were eluted with DTT, and the eluates were processed for LC-MS/MS analysis. The raw protein abundances are plotted over time. Protein abundances are based on peptides with a PEP < 0.05 that were commonly retrieved in the experiments shown, with the exception of HSD17B4 and HSPA9 (for these proteins, no common peptides were identified in the po-IBD-SBP-YAP1C and c-IBD-SBP-YAP1C samples).

**Figure S6. Sulfenome profiles of a selected set of cytosolically located po-IBD-SBP-YAP1C interactors in response to peroxisomal H_2_O_2_ production.** Po-DD-DAO Flp-In T-REx 293 cells expressing po- or c-IBD-SBP-YAP1C were cultured in medium containing 1 μg/ml of DOX and 500 nM Shield1. After 3 days, the medium was replaced with medium lacking DOX/Shield1. One day later, the cells were incubated in DPBS containing 10 mM 3-AT and 10 mM D-alanine. At selected time points (0, 2, 5, 15, 30, and 60 min), free thiol groups were blocked with NEM. Next, the YAP1C-containing protein complexes were affinity purified, the IBD-SBP-YAP1C interaction partners were eluted with DTT, and the eluates were processed for LC-MS/MS analysis. The raw protein abundances are plotted over time. Protein abundances are based on peptides with a PEP < 0.05 that were commonly retrieved in the experiments shown.

**Figure S7. Sulfenome profiles of aberrantly behaving po-IBD-SBP-YAP1C interactors in response to peroxisomal H_2_O_2_ production.** Po-DD-DAO Flp-In T-REx 293 cells expressing po- or c-IBD-SBP-YAP1C were cultured in medium containing 1 μg/ml of DOX and 500 nM Shield1. After 3 days, the medium was replaced with medium lacking DOX/Shield1. One day later, the cells were incubated in DPBS containing 10 mM 3-AT and 10 mM D-alanine. At selected time points (0, 2, 5, 15, 30, and 60 min), free thiol groups were blocked with NEM. Next, the YAP1C-containing protein complexes were affinity purified, the IBD-SBP-YAP1C interaction partners were eluted with DTT, and the eluates were processed for LC-MS/MS analysis. The raw protein abundances are plotted over time. Protein abundances are based on peptides with a PEP < 0.05 that were commonly retrieved in the experiments shown.

**Figure S8. Sulfenome profiles of cytosolically located mt-IBD-SBP-YAPC1 interactors in response to peroxisomal H_2_O_2_ production.** Po-DD-DAO Flp-In T-REx 293 cells expressing mt- or c-IBD-SBP-YAP1C were cultured in medium containing 1 μg/ml of DOX and 500 nM Shield1. After 3 days, the medium was replaced with medium lacking DOX/Shield1. One day later, the cells were incubated in DPBS containing 10 mM 3-AT and 10 mM D-alanine. At selected time points (0, 2, 5, 15, 30, and 60 min), free thiol groups were blocked with NEM. Next, the YAP1C-containing protein complexes were affinity purified, the IBD-SBP-YAP1C interaction partners were eluted with DTT, and the eluates were processed for LC-MS/MS analysis. The raw protein abundances are plotted over time. Note that PRDX3 was included as reference profile of a typical mitochondrial protein. Protein abundances are based on peptides with a PEP < 0.05 that were commonly retrieved in the experiments shown.

**Figure S9. Sulfenome profiles of various protein families of c-IBD-SBP-YAPC1 interactors in response to peroxisomal H_2_O_2_ production.** Po-DD-DAO Flp-In T-REx 293 cells expressing c-IBD-SBP-YAP1C were cultured in medium containing 1 μg/ml of DOX and 500 nM Shield1. After 3 days, the medium was replaced with medium lacking DOX/Shield1. One day later, the cells were incubated in DPBS containing 10 mM 3-AT and 10 mM D-alanine. At selected time points (0, 2, 5, 15, 30, and 60 min), free thiol groups were blocked with NEM. Next, the YAP1C-containing protein complexes were affinity purified, the c-IBD-SBP-YAP1C interaction partners were eluted with DTT, and the eluates were processed for LC-MS/MS analysis. **(A)** Actin-related proteins, **(B)** tubulin-related proteins, **(C)** annexins, **(D)** protein chaperones, **(E)** S100 proteins, and **(F)** negative regulators of endopeptidase activity.

**Figure S10. Formation of c-IBD-SBP-YAP1C complexes in response to different concentrations of external H_2_O_2_.** Po-DD-DAO Flp-In T-REx 293 cells expressing c-IBD-SBP-YAP1C were harvested and resuspended in DPBS containing 10 mM 3-AT and different concentrations of H_2_O_2_ (0, 10, 30, 100, 300, or 1000 μM). After 10 min, free thiol groups were blocked with NEM. Next, the c-IBD-SBP-YAP1C-containing protein complexes were affinity purified and the c-IBD-SBP-YAP1C interaction partners were eluted with DTT. The samples were processed for SDS-PAGE under non-reducing conditions and then subjected to immunoblot analysis with rabbit pre-immune serum. The migration points of relevant molecular mass markers (expressed in kDa) are shown on the left. The arrow and arrowheads mark the non-modified and oxidatively modified c-YAP1C complexes, respectively. Non-specific bands are indicated with asterisks. I, input; NB, not bound to beads; B, bound to beads; BE, bound to beads after DTT elution.

**Figure S11. A 24-h chase period is sufficient to counteract c-DD-DAO activity-induced c-IBD-SBP-YAP1C complex formation.** C-DD-DAO Flp-In T-REx 293 cells expressing c-IBD-SBP-YAP1C were cultured in medium containing 1 μg/ml of DOX and 500nM Shield1. After 3 days, the medium was replaced with medium containing (I, induction) or lacking (I+C, induction + chase) DOX/Shield1. One day later, the cells were incubated in DPBS containing 10 mM 3-AT and 10 mM L- or D-alanine. After 10 min, free thiol groups were blocked with NEM. Next, the c-IBD-SBP-YAP1C-containing protein complexes were affinity purified and its interaction partners were eluted with DTT. The samples were processed for SDS-PAGE under non-reducing conditions and then subjected to immunoblot analysis with rabbit pre-immune serum. The migration points of relevant molecular mass markers (expressed in kDa) are shown on the left. The arrow and arrowheads mark the non-modified and oxidatively modified c-IBD-SBP-YAP1C complexes, respectively. Non-specific bands are indicated with asterisks. I, input; NB, not bound to beads; B, bound to beads; BE, bound to beads after DTT elution.

## Legends to Supplementary Tables

**Table S1. Heat map summarizing the raw abundances of proteins trapped by po-, mt-, or c-IBD-SBP-YAP1C in response to treatment of po-DD-DAO Flp-In T-REx 293 cells with 1 mM H_2_O_2_ for 10 min.** DTT eluates of the experiment shown in Fig. S2 were processed for LC-MS/MS analysis. After validation, proteins enriched two-fold or more in at least one of the H_2_O_2_-treated conditions were retrieved, named according to their UniProtKB gene name, and sorted in alphabetical order. The primary subcellular localizations of the proteins (according to the same database) are indicated between brackets. The values represent raw abundances and are color-coded (with white and blue being low and high, respectively). C, cytosol; CS, cytoskeleton; EC, extracellular; EE, early endosome; ER, endoplasmic reticulum; GA, Golgi apparatus; LY, lysosome; MT, mitochondria; NU, nucleus; PM, plasma membrane; PO, peroxisome.

**Table S2. Heat map summarizing the percentage distribution of proteins trapped by po-, mt-, or c-IBD-SBP-YAP1C upon treatment of po-DD-DAO Flp-In T-REx 293 cells with 1 mM H_2_O_2_ for 10 min.** The percentage distributions of the raw abundances of proteins trapped by po-, mt, or c-IBD-SBP-YAP1C (denoted as po-YAP, mt-YAP, and c-YAP, respectively) after H_2_O_2_ treatment were calculated and presented in a heat map (with white and blue intensities representing low and high percentages of complex formation). The UniProtKB gene name was used to name the interactors, and their primary subcellular localizations (according to the same database) are indicated between brackets. C, cytosol; CS, cytoskeleton; EC, extracellular; EE, early endosome; ER, endoplasmic reticulum; GA, Golgi apparatus; LY, lysosome; MT, mitochondria; NU, nucleus; PM, plasma membrane; PO, peroxisome.

**Table S3. Heat map summarizing the raw abundances of proteins trapped by c-IBD-SBP-YAP1C in response to treatment of po-DD-DAO Flp-In T-REx 293 cells with D-Ala.** DTT eluates of the experiment shown in Fig. S4A were processed for LC-MS/MS analysis. After validation, proteins enriched two-fold or more in at least one of the H_2_O_2_-treated conditions were retrieved, named according to their UniProtKB gene name, and sorted in alphabetical order. The primary subcellular localizations of the proteins (according to the same database) are indicated between brackets. The values represent raw abundances and are color-coded (with white and blue being low and high, respectively). C, cytosol; CS, cytoskeleton; EC, extracellular; EE, early endosome; ER, endoplasmic reticulum; GA, Golgi apparatus; LY, lysosome; MT, mitochondria; NU, nucleus; PM, plasma membrane; PO, peroxisome.

**Table S4. Heat map summarizing the raw abundances of proteins trapped by po-IBD-SBP-YAP1C in response to treatment of po-DD-DAO Flp-In T-REx 293 cells with D-Ala.** DTT eluates of the experiment shown in Fig. S4C were processed for LC-MS/MS analysis. After validation, proteins enriched two-fold or more in at least one of the H_2_O_2_-treated conditions were retrieved, named according to their UniProtKB gene name, and sorted in alphabetical order. The primary subcellular localizations of the proteins (according to the same database) are indicated between brackets. The values represent raw abundances and are color-coded (with white and blue being low and high, respectively). C, cytosol; CS, cytoskeleton; EC, extracellular; EE, early endosome; GA, Golgi apparatus; LY, lysosome; MT, mitochondria; NU, nucleus; PM, plasma membrane; PO, peroxisome.

**Table S5. Heat map summarizing the raw abundances of proteins trapped by mt-IBD-SBP-YAP1C in response to treatment of po-DD-DAO Flp-In T-REx 293 cells with D-Ala.** DTT eluates of the experiment shown in Fig. S4B were processed for LC-MS/MS analysis. After validation, proteins enriched two-fold or more in at least one of the H_2_O_2_-treated conditions were retrieved, named according to their UniProtKB gene name, and sorted in alphabetical order. The primary subcellular localizations of the proteins (according to the same database) are indicated between brackets. The values represent raw abundances and are color-coded (with white and blue being low and high, respectively). C, cytosol; CS, cytoskeleton; EC, extracellular; EE, early endosome; ER, endoplasmic reticulum; GA, Golgi apparatus; LY, lysosome; MT, mitochondria; NU, nucleus; PM, plasma membrane.

**Table S6. Heat map summarizing the raw abundances of proteins trapped by c-IBD-SBP-YAP1C upon treatment of po-DD-DAO Flp-In T-REx 293 cells with different H_2_O_2_ concentrations for 10 min.** DTT eluates of the experiment shown in Fig. S10 were processed for LC-MS/MS analysis. After validation, proteins enriched two-fold or more in at least one of the H_2_O_2_-treated conditions were retrieved, named according to their UniProtKB gene name, and sorted in alphabetical order. The primary subcellular localizations of the protein (according to the same database) are indicated between brackets. The values represent raw abundances and are color-coded (with white and blue being low and high, respectively). ???, no specified localization; C, cytosol; CS, cytoskeleton; EC, extracellular; EE, early endosome; ER, endoplasmic reticulum; ES, endosomes; GA, Golgi apparatus; LY, lysosome; MT, mitochondria; NU, nucleus; PM, plasma membrane; RE, recycling endosome.

**Table S7. Heat map summarizing the relative abundances of proteins trapped by c-IBD-SBP-YAP1C in response to L- or D-Ala treatment of chased versus non-chased c-DD-DAO Flp-In T-REx 293 cells.** DTT eluates of the experiment shown in Fig. S11 were processed for LC-MS/MS analysis. After validation, proteins enriched two-fold or more in at least one of the H_2_O_2_-treated conditions were retrieved, named according to their UniProtKB gene name, and sorted in alphabetical order. The primary subcellular localizations of the protein (according to the same database) are indicated between brackets. The values of the no chase (NC) conditions represent the raw protein abundances. Given that DOX significantly reduces the proliferation rate of HEK-293 cells [Ahler et al., 2013], the raw abundances in the chase (C) conditions were first corrected for DOX-induced protein level differences observed as differences in total peptide abundance between the L-Ala-treated no chase and chase conditions. Protein abundances are color-coded (with white and blue being low and high, respectively). ???, no specified localization; C, cytosol; CS, cytoskeleton; EC, extracellular; ER, endoplasmic reticulum; GA, Golgi apparatus; LY, lysosome; MT, mitochondria; NU, nucleus; PM, plasma membrane.

**Table S8. Heat map summarizing the percentage distribution of NaCl- and DTT-eluted proteins trapped by c-IBD-SBP-YAP1C upon treatment of po-DD-DAO Flp-In T-REx 293 cells with 1 mM H_2_O_2_ for 10 min.** The percentage distribution of proteins trapped by c-IBD-SBP-YAP1C after H_2_O_2_ treatment and sequentially eluted with 1 M NaCl and 10 mM DTT was calculated and presented in a heat map (with red and green intensities representing low and high percentages of elution). The UniProtKB gene name was used to name the interactors, and their primary subcellular localizations (according to the same database) are indicated between brackets. ???, no specified localization; C, cytosol; CS, cytoskeleton; EC, extracellular; ER, endoplasmic reticulum; ES, endosome; GA, Golgi apparatus; LY, lysosome; MT, mitochondria; NU, nucleus; PM, plasma membrane; PO, peroxisome.

**Table S9. Overview of S-carbamidomethylated cysteines.** The S-carbamidomethylated cysteines identified in proteins that were at least two-fold enriched in an (external or endogenous) H_2_O_2_-treated condition were compared to oxidative cysteine modifications reported in the iCysMod database and color coded: green, S-sulfenylation is has already been reported by others; blue, the iCysMod database lists other oxidative cysteine thiol modifications; red, the cysteine residue is not yet listed in the iCysMod database (based on the information available on 23 September 2021).

## Footnotes

1 Abbreviations: 3-AT, 3-amino-1,2,4-triazole; c-, cytosolic; DAO, D-amino acid oxidase; DD, destabilization domain; DTT, dithiothreitol; DOX, doxycycline; DPBS, Dulbecco’s phosphate-buffered saline; EMSA, electrophoretic mobility shift assay; IBD, IgG-binding domain; MS, mass spectrometry; mt-, mitochondrial; NEM, N-ethylmaleimide; PEP, posterior error probability; po-, peroxisomal; PRDX, peroxiredoxin; PTS1, C-terminal peroxisomal targeting signal; SBP, streptavidin-binding protein; YAP1C, C-terminal region of yeast AP-1-like transcription factor. Given the extensive list of proteins identified in the sulfenome mining experiments, we refer the reader to the UniProtKB database for details on the protein acronyms used in the manuscript.

## References

Ahler E, Sullivan WJ, Cass A, Braas D, York AG, Bensinger SJ, Graeber TG, Christofk HR. Doxycycline alters metabolism and proliferation of human cell lines. PLoS One 2013; 8: e64561. https://doi.org/10.1371/journal.pone.0064561

Anthonio EA, Brees C, Baumgart-Vogt E, Hongu T, Huybrechts SJ, Van Dijck P, Mannaerts GP, Kanaho Y, Van Veldhoven PP, Fransen M. Small G proteins in peroxisome biogenesis: the potential involvement of ADP-ribosylation factor 6. BMC Cell Biol 2009; 10: 58. https://doi.org/10.1186/1471-2121-10-58

Baumgart E, Vanhooren JC, Fransen, M, Van Leuven F, Fahimi HD, Van Veldhoven PP, Mannaerts GP. Molecular cloning and further characterization of rat peroxisomal trihydroxycoprostanoyl-CoA oxidase. Biochem J 1996; 320: 115–121. https://doi.org/10.1042/bj3200115

Boveris A, Oshino N, Chance B. The cellular production of hydrogen peroxide. Biochem J 1972; 128: 617–630. https://doi.org/10.1042/bj1280617

Chung HL, Wangler MF, Marcogliese PC, et al. Loss-or gain-of-function mutations in ACOX1 cause axonal loss via different mechanisms. Neuron 2020; 106: 589–606. https://doi.org/10.1016/j.neuron.2020.02.021

De Duve C, Baudhuin P. Peroxisomes (microbodies and related particles). Physiol Rev 1966; 46: 323–357. https://doi.org/10.1152/physrev.1966.46.2.323

Even DY, Kedmi A, Basch-Barzilay S, Ideses D, Tikotzki R, Shir-Shapira H, Shefi O, Juven-Gershon T. Engineered promoters for potent transient overexpression. PLoS One 2016; 11: e0148918. https://doi.org/10.1371/journal.pone.0148918

Fransen M, Lismont C. Redox signaling from and to peroxisomes: progress, challenges, and prospects. Antioxid Redox Signal 2019; 30: 95–112. https://doi.org/10.1089/ars.2018.7515

Fransen M, Wylin T, Brees C, Mannaerts GP, Van Veldhoven PP. Human Pex19p binds peroxisomal integral membrane proteins at regions distinct from their sorting sequences. Mol Cell Biol 2001; 21: 4413–4424. https://doi.org/10.1128/MCB.21.13.4413-4424.2001

Geiger T, Wehner A, Schaab C, Cox J, Mann M. Comparative proteomic analysis of eleven common cell lines reveals ubiquitous but varying expression of most proteins. Mol Cell Proteomics 2012; 11: M111.014050. https://doi.org/10.1074/mcp.M111.014050

Goemaere J, Knoops B. Peroxiredoxin distribution in the mouse brain with emphasis on neuronal populations affected in neurodegenerative disorders. J Comp Neurol 2012; 520: 258–280. https://doi.org/10.1002/cne.22689

Góth L, Nagy T. Inherited catalase deficiency: is it benign or a factor in various age-related disorders? Mutat Res 2013; 753: 147–154. https://doi.org/10.1016/j.mrrev.2013.08.002

Hanschmann EM, Godoy JR, Berndt C, Hudemann C, Lillig CH. Thioredoxins, glutaredoxins, and peroxiredoxins--molecular mechanisms and health significance: from cofactors to antioxidants to redox signaling. Antioxid Redox Signal 2013; 19: 1539–1605. https://doi.org/10.1089/ars.2012.4599

He A, Dean JM, Lodhi IJ. Peroxisomes as cellular adaptors to metabolic and environmental stress. Trends Cell Biol 2021; 31: 656–670. https://doi.org/10.1016/j.tcb.2021.02.005

Hofhuis J, Schueren F, Nötzel C, Lingner T, Gärtner J, Jahn O, Thoms S. The functional readthrough extension of malate dehydrogenase reveals a modification of the genetic code. Open Biol 2016; 6: 160246. https://doi.org/10.1098/rsob.160246

Huang B, Sikes H. Quantifying intracellular hydrogen peroxide perturbations in terms of concentration. Redox Biol 2014; 2: 955–962. https://doi.org/10.1016/j.redox.2014.08.001

Ivashchenko O, Van Veldhoven PP, Brees C, Ho YS, Terlecky SR, Fransen M. Intraperoxisomal redox balance in mammalian cells: oxidative stress and interorganellar crosstalk. Mol Biol Cell 2011; 22: 1440–1451. https://doi.org/10.1091/mbc.e10-11-0919

Izawa S, Maeda K, Sugiyama K, Mano J, Inoue Y, Kimura A. Thioredoxin deficiency causes the constitutive activation of Yap1, an AP-1-like transcription factor in *Saccharomyces cerevisiae*. J Biol Chem 1999; 274: 28459–28465. https://doi.org/10.1074/jbc.274.40.28459

Jo DS, Park SJ, Kim AK, et al. Loss of HSPA9 induces peroxisomal degradation by increasing pexophagy. Autophagy 2020; 16: 1989–2003. https://doi.org/10.1080/15548627.2020.1712812

Knoops B, Goemaere J, Van der Eecken V, Declercq JP. Peroxiredoxin 5: structure, mechanism, and function of the mammalian atypical 2-Cys peroxiredoxin. Antioxid Redox Signal 2011; 15: 817–829. https://doi.org/10.1089/ars.2010.3584

Kunze M. Predicting peroxisomal targeting signals to elucidate the peroxisomal proteome of mammals. Subcell Biochem 2018; 89: 157–199. https://doi.org/10.1007/978-981-13-2233-4_7

Lennicke C, Rahn J, Lichtenfels R, Wessjohann LA, Seliger B. Hydrogen peroxide – production, fate and role in redox signaling of tumor cells. Cell Commun Signal 2015; 13: 39. https://doi.org/10.1186/s12964-015-0118-6

Liebthal M, Schuetze J, Dreyer A, Mock HP, Dietz KJ. Redox conformation-specific protein-protein interactions of the 2-cysteine peroxiredoxin in *Arabidopsis*. Antioxidants 2020; 9: 515. https://doi.org/10.3390/antiox9060515

Lismont C, Koster J, Provost S, Baes M, Van Veldhoven PP, Waterham HR, Fransen M. Deciphering the potential involvement of PXMP2 and PEX11B in hydrogen peroxide permeation across the peroxisomal membrane reveals a role for PEX11B in protein sorting. Biochim Biophys Acta Biomembr 2019c; 1861: 182991. https://doi.org/10.1016/j.bbamem.2019.05.013

Lismont C, Nordgren M, Brees C, Knoops B, Van Veldhoven PP, Fransen M. Peroxisomes as modulators of cellular protein thiol oxidation: a new model system. Antioxid Redox Signal 2019a; 30: 22–39. https://doi.org/10.1089/ars.2017.6997

Lismont C, Nordgren M, Van Veldhoven PP, Fransen M. Redox interplay between mitochondria and peroxisomes. Front Cell Dev Biol 2015; 3: 35. https://doi.org/10.3389/fcell.2015.00035

Lismont C, Revenco I, Fransen M. Peroxisomal hydrogen peroxide metabolism and signaling in health and disease. Int J Mol Sci 2019b; 20: 3673. https://doi.org/10.3390/ijms20153673

Lyublinskaya O, Antunes F. Measuring intracellular concentration of hydrogen peroxide with the use of genetically encoded H2O2 biosensor HyPer. Redox Biol 2019; 24: 101200. https://doi.org/10.1016/j.redox.2019.101200

Méndez-Barbero N, Gutiérrez-Muñoz C, Blázquez-Serra R, Martín-Ventura JL, Blanco-Colio LM. Annexins: Involvement in cholesterol homeostasis, inflammatory response and atherosclerosis. Clin Investig Arterioscler 2021; 33: 206–216. https://doi.org/10.1016/j.arteri.2020.12.010

Passmore JB, Carmichael RE, Schrader TA, Godinho LF, Ferdinandusse S, Lismont C, Wang Y, Hacker C, Islinger M, Fransen M, Richards DM, Freisinger P, Schrader M. Mitochondrial fission factor (MFF) is a critical regulator of peroxisome maturation. Biochim Biophys Acta Mol Cell Res 2020; 1867: 118709. https://doi.org/10.1016/j.bbamcr.2020.118709

Poole LB. The basics of thiols and cysteines in redox biology and chemistry. Free Radic Biol Med 2015; 80: 148–157. https://doi.org/10.1016/j.freeradbiomed.2014.11.013

Pujato M, Kieken F, Skiles AA, Tapinos N, Fiser A. Prediction of DNA binding motifs from 3D models of transcription factors; identifying TLX3 regulated genes. Nucleic Acids Res 2014; 42: 13500–132512. https://doi.org/10.1093/nar/gku1228

Ramazani Y, Knops N, Berlingerio SP, Adebayo OC, Lismont C, Kuypers DJ, Levtchenko E, van den Heuvel LP, Fransen M. Therapeutic concentrations of calcineurin inhibitors do not deregulate glutathione redox balance in human renal proximal tubule cells. PLoS One 2021; 16: e0250996. https://doi.org/10.1371/journal.pone.0250996

Rath S, Sharma R, Gupta R, et al. MitoCarta3.0: an updated mitochondrial proteome now with sub-organelle localization and pathway annotations. Nucleic Acids Res 2021; 49: D1541–D1547. https://doi.org/10.1093/nar/gkaa1011

Raudvere U, Kolberg L, Kuzmin I, Arak T, Adler P, Peterson H, Vilo J. g:Profiler: a web server for functional enrichment analysis and conversions of gene lists (2019 update). Nucleic Acids Res 2019; 38: W71–W77. https://doi.org/10.1093/nar/gkq329

Rhee SG, Woo HA. Multiple functions of 2-Cys peroxiredoxins, I and II, and their regulations via post-translational modifications. Free Radic Biol Med 2020; 152: 107–115. https://doi.org/10.1016/j.freeradbiomed.2020.02.028

Rosenthal M, Metzl-Raz E, Bürgi J, Yifrach E, Drwesh L, Fadel A, Peleg Y, Rapaport D, Wilmanns M, Barkai N, Schuldiner M, Zalckvar E. Uncovering targeting priority to yeast peroxisomes using an in-cell competition assay. Proc Natl Acad Sci USA 2020; 117: 21432–21440. https://doi.org/10.1073/pnas.1920078117

Rubio MP, Geraghty KM, Wong BHC, Wood NT, Campbell DG, Morrice N, Mackintosh C. 14-3-3-affinity purification of over 200 human phosphoproteins reveals new links to regulation of cellular metabolism, proliferation and trafficking. Biochem J 2004; 379: 395–408. https://doi.org/10.1042/bi20031797

Schueren F, Lingner T, George R, Hofhuis J, Dickel C, Gärtner J, Thoms S. Peroxisomal lactate dehydrogenase is generated by translational readthrough in mammals. Elife 2014; 3: e03640. https://doi.org/10.7554/eLife.03640.001

Sies H, Jones DP. Reactive oxygen species (ROS) as pleiotropic physiological signalling agents. Nat Rev Mol Cell Biol 2020; 21: 363–383. https://doi.org/10.1038/s41580-020-0230-3

Takanishi CL, Ma LH, Wood MJ. A genetically encoded probe for cysteine sulfenic acid protein modification *in vivo*. Biochemistry 2007; 46: 14725–14732. https://doi.org/10.1021/bi701625s

Thul PJ, Åkesson L, Wiking M et al. A subcellular map of the human proteome. Science 2017; 356: eaa13321. https://doi.org/10.1126/science.aal3321

Walton PA, Brees C, Lismont C, Apanasets O, Fransen M. The peroxisomal import receptor PEX5 functions as a stress sensor, retaining catalase in the cytosol in times of oxidative stress. Biochim Biophys Acta Mol Cell Res 2017; 1864: 1833–1843. https://doi.org/10.1016/j.bbamcr.2017.07.013

Wang P, Zhang Q, Li S, Cheng B, Xue H, Wei Z, Shao T, Liu ZX, Cheng H, Wang Z. iCysMod: an integrative database for protein cysteine modifications in eukaryotes. Brief Bioinform 2021; 22: bbaa400. https://doi.org/10.1093/bib/bbaa400

Waszczak C, Akter S, Eeckhout D, Persiau G, Wahni K, Bodra N, Van Molle I, De Smet B, Vertommen D, Gevaert K, De Jaeger G, Van Montagu M, Messens J, Van Breusegem F. Sulfenome mining in *Arabidopsis thaliana*. Proc Natl Acad Sci USA 2014; 111: 11545–11550. https://doi.org/10.1073/pnas.1411607111

Weidberg H, Amon A. MitoCPR-A surveillance pathway that protects mitochondria in response to protein import stress. Science 2018; 360: eaan4146. https://doi.org/10.1126/science.aan4146

Wessel D, Flügge U. A method for the quantitative recovery of protein in dilute solution in the presence of detergents and lipids. Anal Biochem 1984; 138: 141–143. https://doi.org/10.1016/0003-2697(84)90782-6

Wu Y, Zhou Q, Guo F, Chen M, Tao X, Dong D. S100 Proteins in pancreatic cancer: current knowledge and future perspectives. Front Oncol 2021; 11: 711180. https://doi.org/10.3389/fonc.2021.711180

Yifrach E, Fischer S, Oeljeklaus S, Schuldiner M, Zalckvar E, Warscheid B. Defining the mammalian peroxisomal proteome. Subcell Biochem 2018; 89: 47–66. https://doi.org/10.1007/978-981-13-2233-4_2

Yogev O, Pines O. Dual targeting of mitochondrial proteins: mechanism, regulation and function. Biochim Biophys Acta 2011; 1808: 1012–1020. https://doi.org/10.1016/j.bbamem.2010.07.004

Zavialov A, Benndorf R, Ehrnsperger M, Zav’yalov V, Dudich I, Buchner J, Gaestel M. The effect of the intersubunit disulfide bond on the structural and functional properties of the small heat shock protein Hsp25. Int J Biol Macromol 1998; 22: 163–173. https://doi.org/10.1016/S0141-8130(98)00014-2

Zhukova L, Zhukov I, Bal W, Wyslouch-Cieszynska A. Redox modifications of the C-terminal cysteine residue cause structural changes in S100A1 and S100B proteins. Biochim Biophys Acta 2004; 1742: 191–201. https://doi.org/10.1016/j.bbamcr.2004.10.002

