## Supplementary Figures for "Peroxisome-derived hydrogen peroxide can modulate the sulfenylation profiles of key redox signaling proteins"

#### Fig. S1

```
gaattcccacc**atg**gggactccggcagtcacaacctacaaactggtgattaacgggaagaca
              M  G  T  P  A  V  T  T  Y  K  L  V  I  N  G  K  T
  ctcaaaggtgaaacgaccacaaaagcagttgatgcggaaactgccgaaaaggcatttaag
   L  K  G  E  T  T  T  K  A  V  D  A  E  T  A  E  K  A  F  K
  caatatgccaacgataatggggtggacggggtgtggacgtatgacgatgccacgaaaaca
   Q  Y  A  N  D  N  G  V  D  G  V  W  T  Y  D  D  A  T  K  T
  ttcaccgtgactgaagtgaatacccccgccgttacaacttataaattggttattaacggc
   F  T  V  T  E  V  N  T  P  A  V  T  T  Y  K  L  V  I  N  G
  aaaacactcaaggggggagaccaccactaaggcagtggatgcagagactgcagagaaggca
   K  T  L  K  G  E  T  T  T  K  A  V  D  A  E  T  A  E  K  A
  ttcaagcaatacgcgaatgataacggcgtcgacggggtttggacttatgatgacgccact
   F  K  Q  Y  A  N  D  N  G  V  D  G  V  W  T  Y  D  D  A  T
  aagacttttacggtcacagagatcgatgagaacctgtactttcaaggcggaagcatggac
   K  T  F  T  V  T  E  I  D  E  N  L  Y  F  Q  G  G  S  M  D
  gaaaagacgacagggtggcggggaggtcacgttgttgagggtttggcaggtgaacttgaa
   E  K  T  T  G  W  R  G  G  H  V  V  E  G  L  A  G  E  L  E
  cagctccgggcgaggctggaacaccatcctcaaggtcaacgagaacccgggggaggagga
   Q  L  R  A  R  L  E  H  H  P  Q  G  Q  R  E  P  G  G  G  G
  tcactggagattacatctctttacaagaaggctggaagcacaaatggatcaagcctgcag
   S  L  E  I  T  S  L  Y  K  K  A  G  S  T  N  G  S  S  L  Q
  aatgctgacaagattaataatggaaatgacaacgataatgacaatgatgtggtaccttca
   N  A  D  K  I  N  N  G  N  D  N  D  N  D  N  D  V  V  P  S
  aaggagggctcactgctccgctgtagcgagatttgggaccgaataacaactcatccgaaa
   K  E  G  S  L  L  R  C  S  E  I  W  D  R  I  T  T  H  P  K
  tactctgacattgatgtagatgggctcgcgtccgaattgatggccaaggccaaaacaagc
   Y  S  D  I  D  V  D  G  L  A  S  E  L  M  A  K  A  K  T  S
  gaacgaggagtcgtgataaacgcggaagatgtccagcttgcattgaataaacatatgaac
   E  R  G  V  V  I  N  A  E  D  V  Q  L  A  L  N  K  H  M  N
  **tga**gcggccgc
```

**Fig. S2**

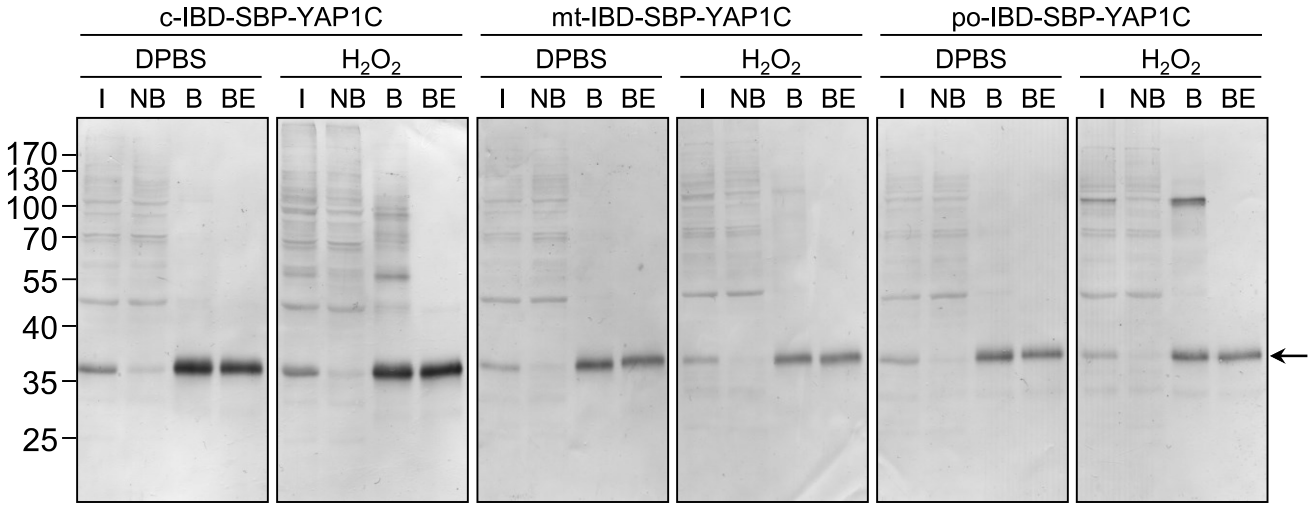
c-IBD-SBP-YAP1C
mt-IBD-SBP-YAP1C
po-IBD-SBP-YAP1C
DPBS
$H_2O_2$
I
NB
B
BE
170
130
100
70
55
40
35
25

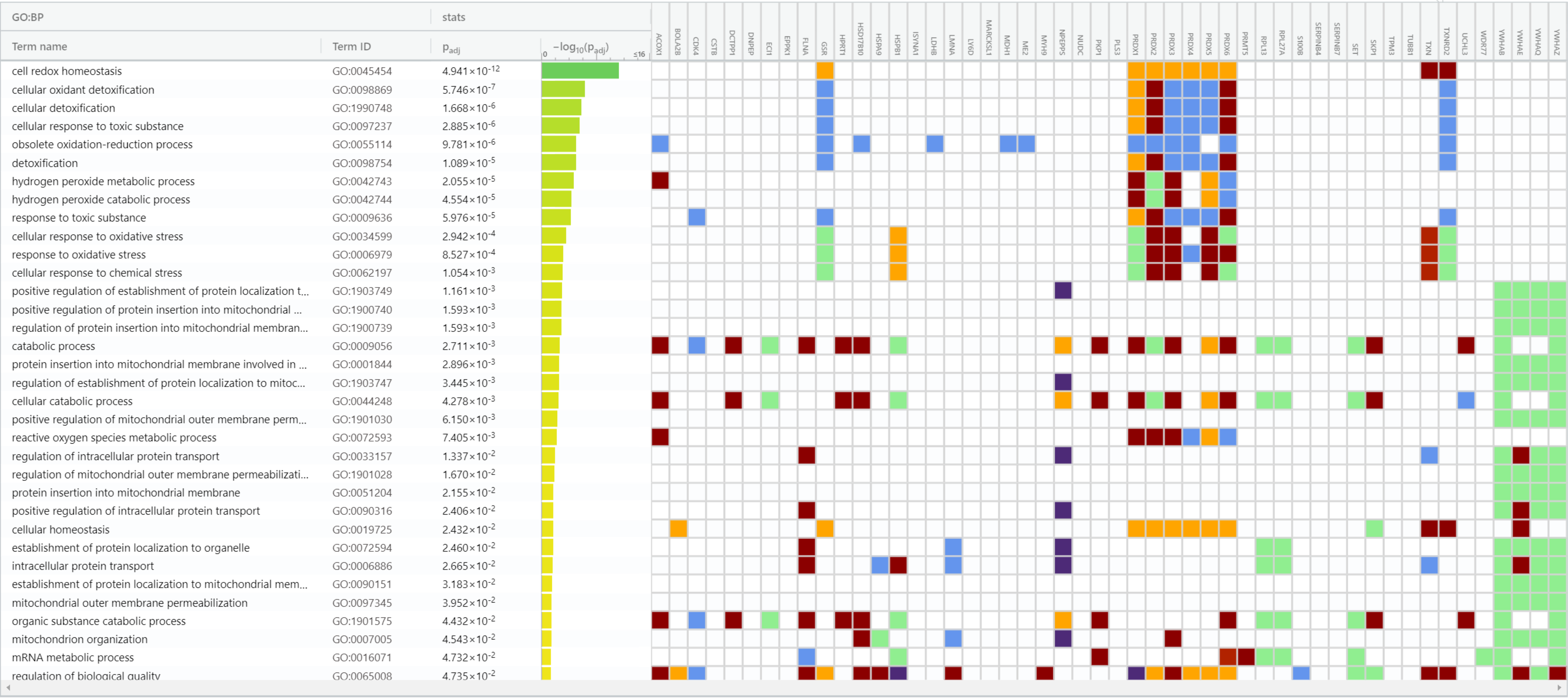
GO:BP
stats
Term name
Term ID
Padj
−log10(Padj)
0
≤16
ACOX1
BOLA2B
CDK4
CSTB
DCTPP1
DNPEP
ECI1
EPPK1
FLNA
GSR
HPRT1
HSD17B10
HSPA9
HSPB1
ISYNA1
LDHB
LMNA
LY6D
MARCKSL1
MDH1
ME2
MYH9
NPEPPS
NUDC
PKP1
PLS3
PRDX1
PRDX2
PRDX3
PRDX4
PRDX5
PRDX6
PRMT5
RPL13
RPL27A
S100B
SERPINB4
SERPINB7
SET
SKP1
TPM3
TUBB1
TXN
TXNRD2
UCHL3
WDR77
YWHAB
YWHAE
YWHAQ
YWHAZ
cell redox homeostasis GO:0045454 4.941×10-12
cellular oxidant detoxification GO:0098869 5.746×10-7
cellular detoxification GO:1990748 1.668×10-6
cellular response to toxic substance GO:0097237 2.885×10-6
obsolete oxidation-reduction process GO:0055114 9.781×10-6
detoxification GO:0098754 1.089×10-5
hydrogen peroxide metabolic process GO:0042743 2.055×10-5
hydrogen peroxide catabolic process GO:0042744 4.554×10-5
response to toxic substance GO:0009636 5.976×10-5
cellular response to oxidative stress GO:0034599 2.942×10-4
response to oxidative stress GO:0006979 8.527×10-4
cellular response to chemical stress GO:0062197 1.054×10-3
positive regulation of establishment of protein localization t... GO:1903749 1.161×10-3
positive regulation of protein insertion into mitochondrial ... GO:1900740 1.593×10-3
regulation of protein insertion into mitochondrial membran... GO:1900739 1.593×10-3
catabolic process GO:0009056 2.711×10-3
protein insertion into mitochondrial membrane involved in ... GO:0001844 2.896×10-3
regulation of establishment of protein localization to mitoc... GO:1903747 3.445×10-3
cellular catabolic process GO:0044248 4.278×10-3
positive regulation of mitochondrial outer membrane perm... GO:1901030 6.150×10-3
reactive oxygen species metabolic process GO:0072593 7.405×10-3
regulation of intracellular protein transport GO:0033157 1.337×10-2
regulation of mitochondrial outer membrane permeabilizati... GO:1901028 1.670×10-2
protein insertion into mitochondrial membrane GO:0051204 2.155×10-2
positive regulation of intracellular protein transport GO:0090316 2.406×10-2
cellular homeostasis GO:0019725 2.432×10-2
establishment of protein localization to organelle GO:0072594 2.460×10-2
intracellular protein transport GO:0006886 2.665×10-2
establishment of protein localization to mitochondrial mem... GO:0090151 3.183×10-2
mitochondrial outer membrane permeabilization GO:0097345 3.952×10-2
organic substance catabolic process GO:1901575 4.432×10-2
mitochondrion organization GO:0007005 4.543×10-2
mRNA metabolic process GO:0016071 4.732×10-2
regulation of biological quality GO:0065008 4.735×10-2

Fig. S4

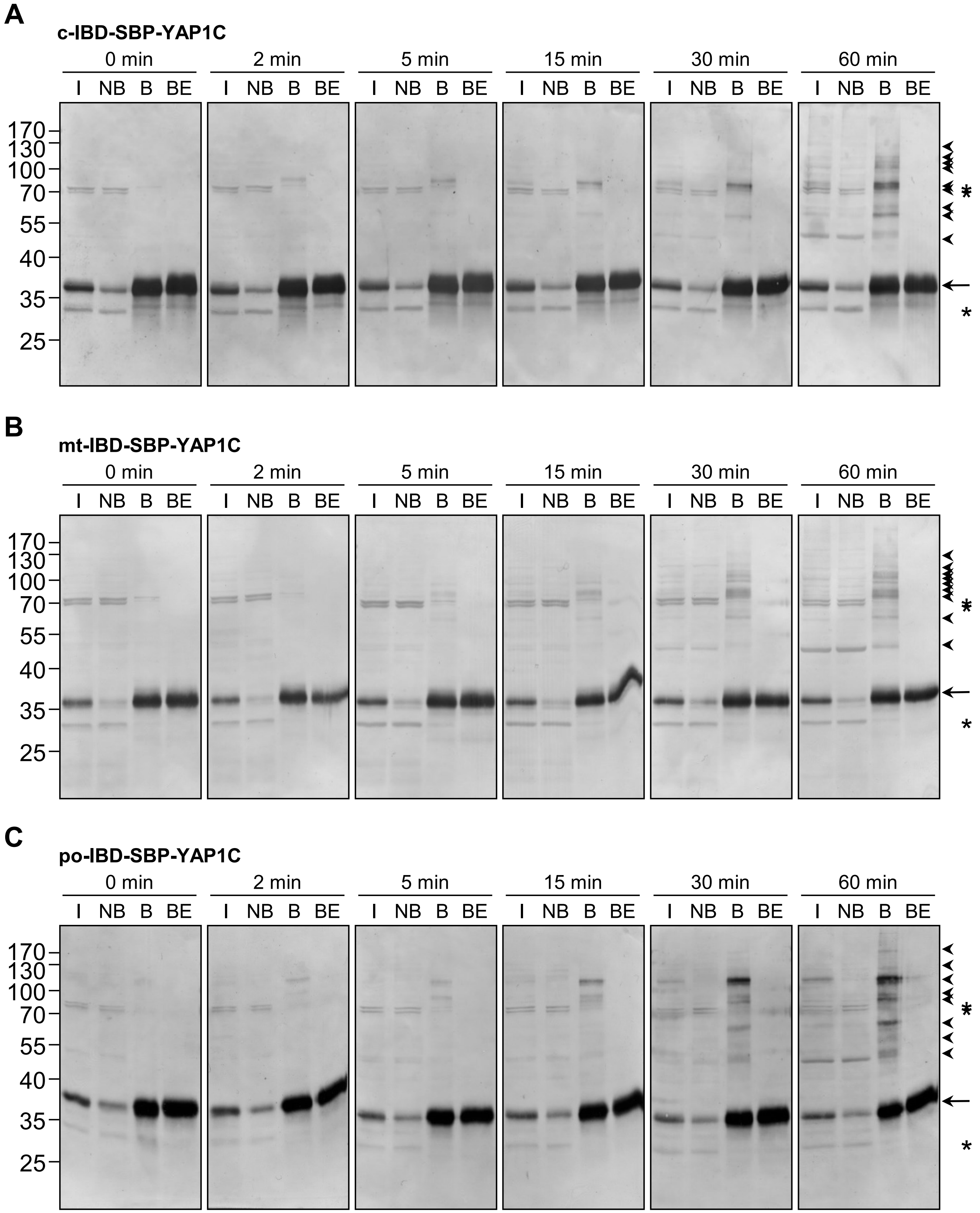
A
c-IBD-SBP-YAP1C
0 min
2 min
5 min
15 min
30 min
60 min
I NB B BE
170
130
100
70
55
40
35
25
*
*
B
mt-IBD-SBP-YAP1C
C
po-IBD-SBP-YAP1C

Fig. S5

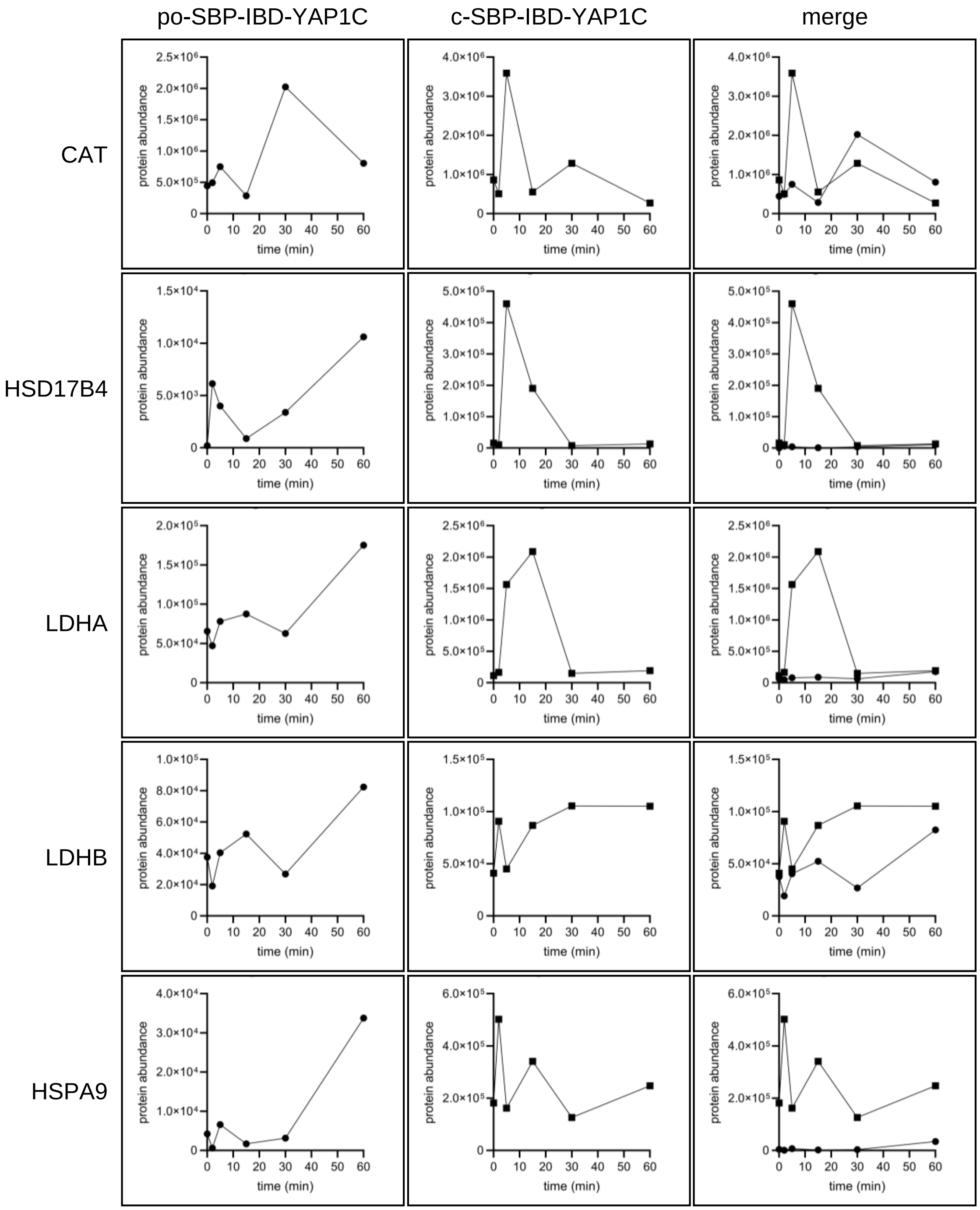
po-SBP-IBD-YAP1C
c-SBP-IBD-YAP1C
merge
CAT
HSD17B4
LDHA
LDHB
HSPA9
protein abundance
time (min)

### Fig. S6

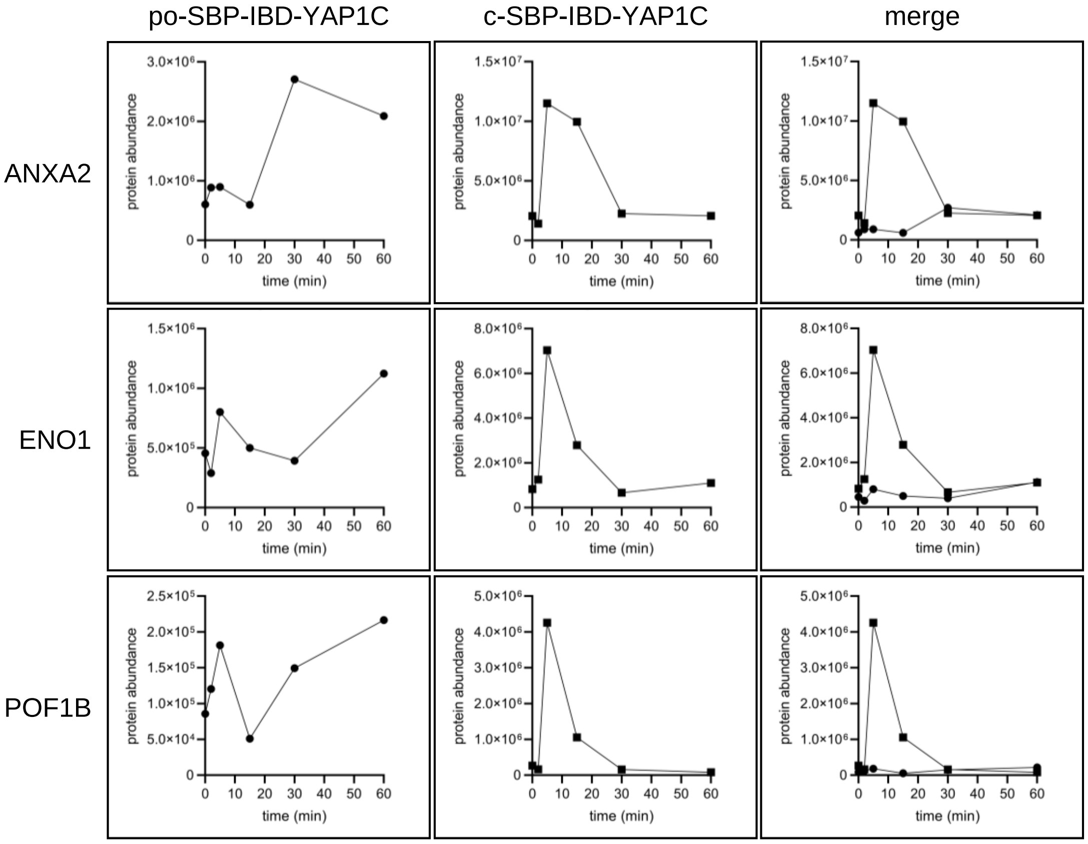

po-SBP-IBD-YAP1C
c-SBP-IBD-YAP1C
merge
ANXA2
ENO1
POF1B
protein abundance
time (min)

### Fig. S7

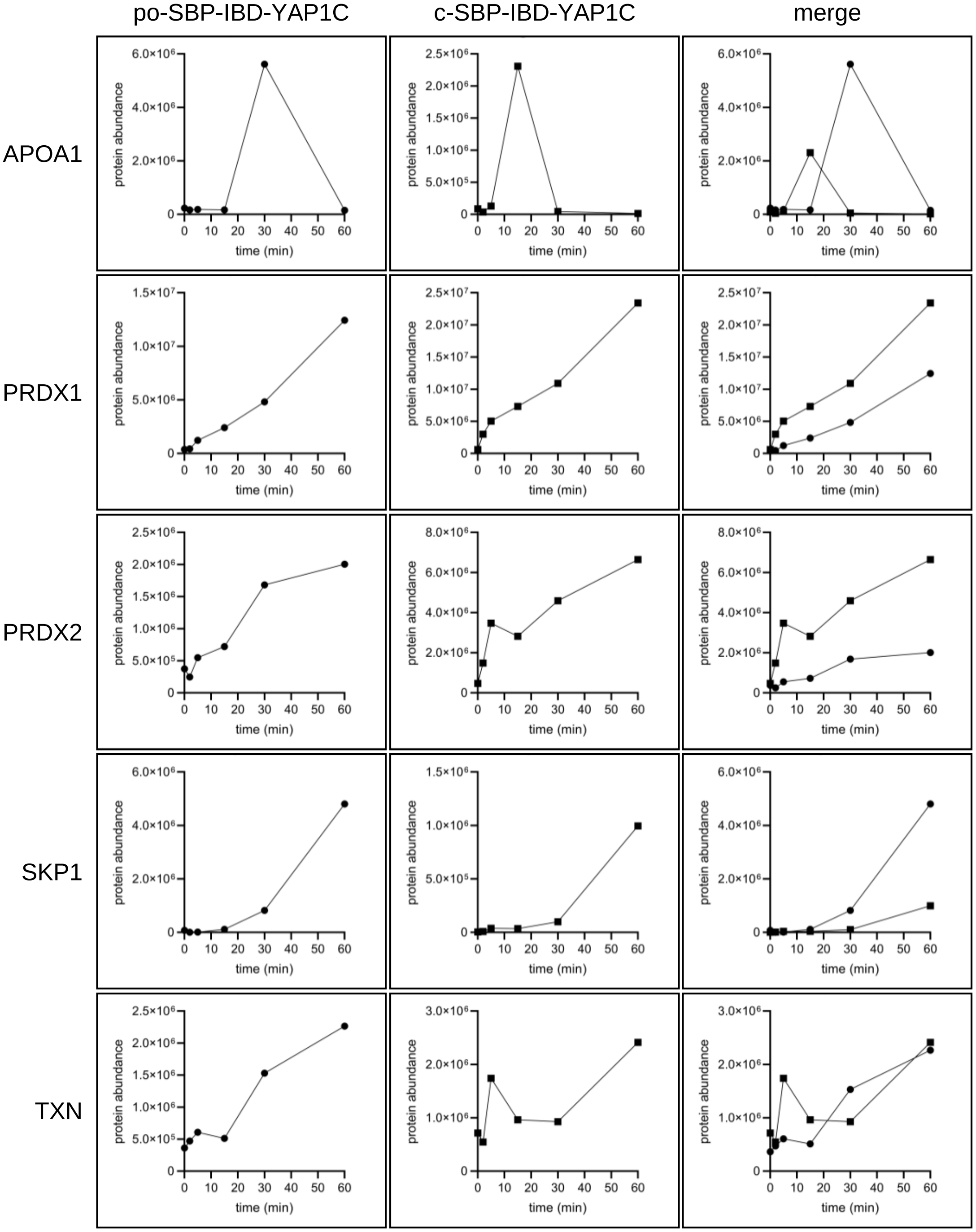
po-SBP-IBD-YAP1C
c-SBP-IBD-YAP1C
merge
APOA1
PRDX1
PRDX2
SKP1
TXN
protein abundance
time (min)

**Fig. S8**

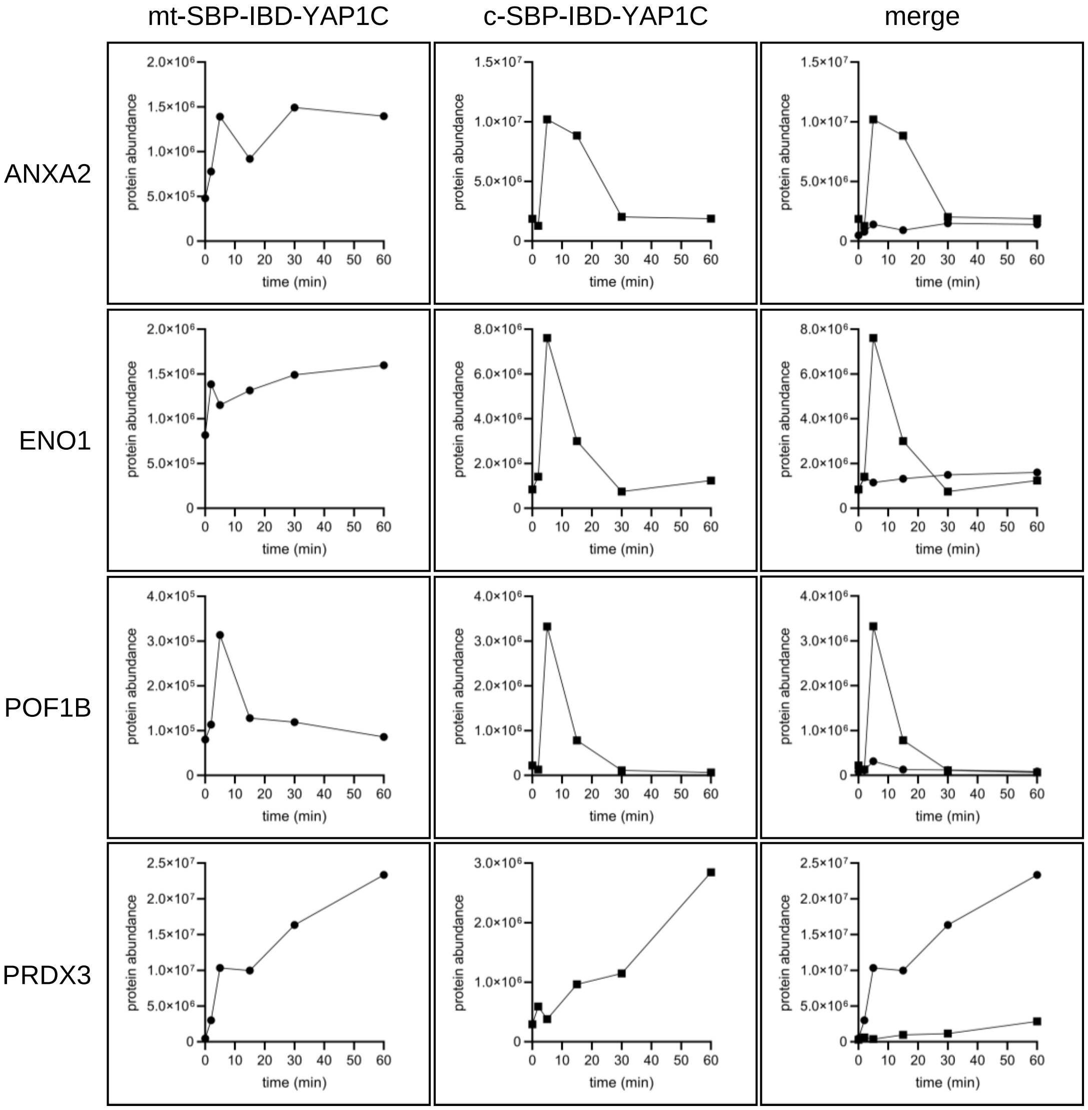

mt-SBP-IBD-YAP1C
c-SBP-IBD-YAP1C
merge
ANXA2
ENO1
POF1B
PRDX3
protein abundance
time (min)

**Fig. S9**

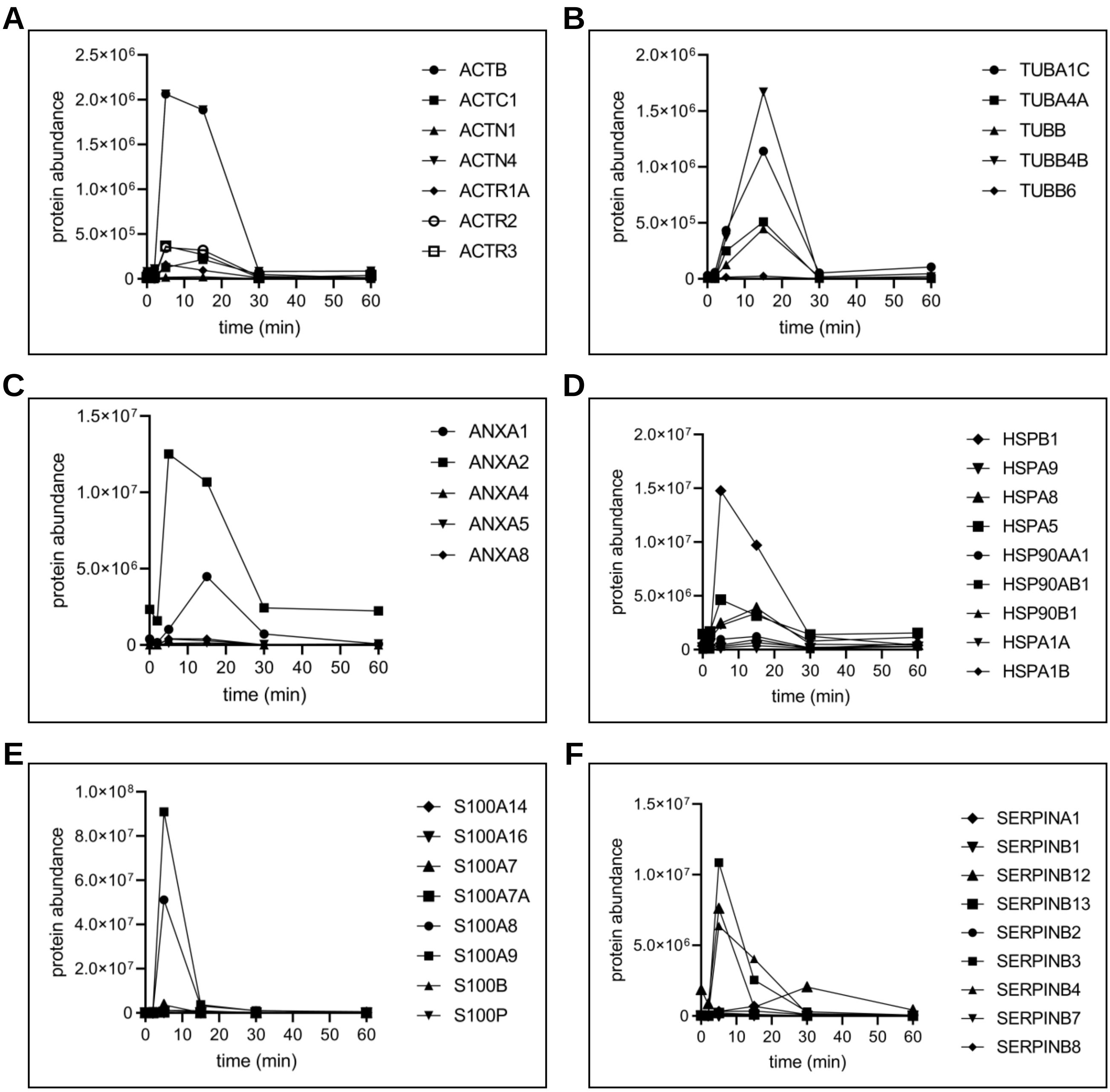
A
protein abundance
2.5×10⁶
2.0×10⁶
1.5×10⁶
1.0×10⁶
5.0×10⁵
0
0 10 20 30 40 50 60
time (min)
ACTB
ACTC1
ACTN1
ACTN4
ACTR1A
ACTR2
ACTR3
B
protein abundance
2.0×10⁶
1.5×10⁶
1.0×10⁶
5.0×10⁵
0
0 10 20 30 40 50 60
time (min)
TUBA1C
TUBA4A
TUBB
TUBB4B
TUBB6
C
protein abundance
1.5×10⁷
1.0×10⁷
5.0×10⁶
0
0 10 20 30 40 50 60
time (min)
ANXA1
ANXA2
ANXA4
ANXA5
ANXA8
D
protein abundance
2.0×10⁷
1.5×10⁷
1.0×10⁷
5.0×10⁶
0
0 10 20 30 40 50 60
time (min)
HSPB1
HSPA9
HSPA8
HSPA5
HSP90AA1
HSP90AB1
HSP90B1
HSPA1A
HSPA1B
E
protein abundance
1.0×10⁸
8.0×10⁷
6.0×10⁷
4.0×10⁷
2.0×10⁷
0
0 10 20 30 40 50 60
time (min)
S100A14
S100A16
S100A7
S100A7A
S100A8
S100A9
S100B
S100P
F
protein abundance
1.5×10⁷
1.0×10⁷
5.0×10⁶
0
0 10 20 30 40 50 60
time (min)
SERPINA1
SERPINB1
SERPINB12
SERPINB13
SERPINB2
SERPINB3
SERPINB4
SERPINB7
SERPINB8

**Fig. S10**

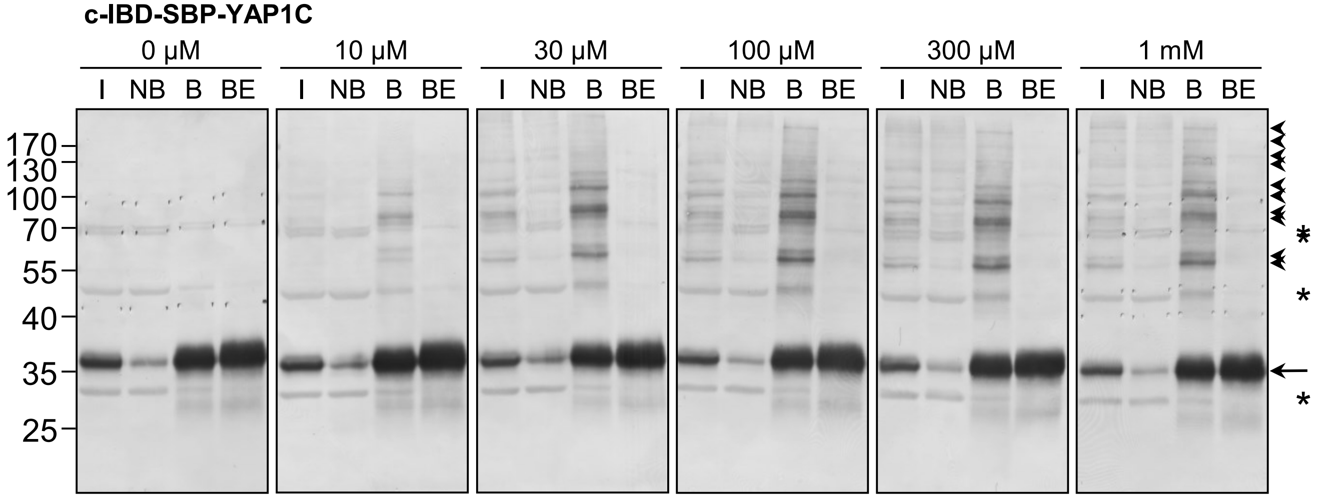

c-IBD-SBP-YAP1C
0 µM
10 µM
30 µM
100 µM
300 µM
1 mM
I NB B BE
170
130
100
70
55
40
35
25
*
*
*

**Fig. S11**

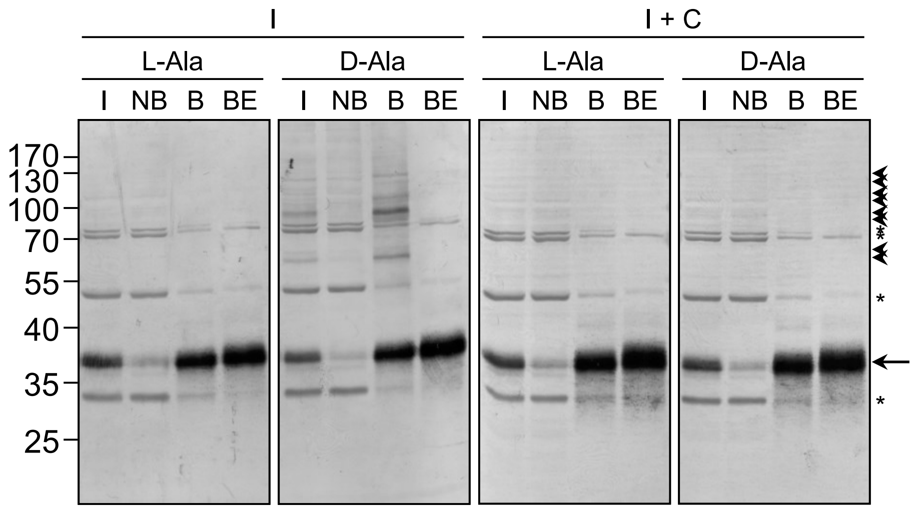

I
I + C
L-Ala
D-Ala
L-Ala
D-Ala
I NB B BE
I NB B BE
I NB B BE
I NB B BE
170
130
100
70
55
40
35
25
*
*
*
