## Supplementary Tables for "Peroxisome-derived hydrogen peroxide can modulate the sulfenylation profiles of key redox signaling proteins"

### Table S1

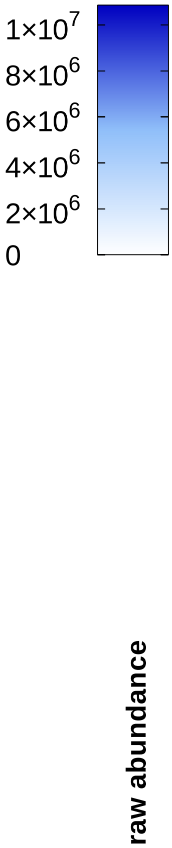


| Protein | po-IBD-SBP-YAP 0 mM $H_2O_2$ | po-IBD-SBP-YAP 1 mM $H_2O_2$ | mt-IBD-SBP-YAP 0 mM $H_2O_2$ | mt-IBD-SBP-YAP 1 mM $H_2O_2$ | c-IBD-SBP-YAP 0 mM $H_2O_2$ | c-IBD-SBP-YAP 1 mM $H_2O_2$ |
|---|---|---|---|---|---|---|
| ACOX1 (PO) | $1.51\times10^{5}$ | $2.45\times10^{6}$ | $1.61\times10^{5}$ | $1.35\times10^{5}$ | $1.38\times10^{5}$ | $2.83\times10^{5}$ |
| BOLA2B (C/NU) | $8.38\times10^{3}$ | $1.24\times10^{5}$ | $2.74\times10^{4}$ | $9.07\times10^{4}$ | $3.05\times10^{4}$ | $2.11\times10^{5}$ |
| CDK4 (C/NU) | $5.41\times10^{3}$ | $5.80\times10^{4}$ | $2.86\times10^{3}$ | $2.87\times10^{4}$ | $1.02\times10^{4}$ | $3.12\times10^{5}$ |
| CSTB (C/NU) | $1.93\times10^{5}$ | $1.24\times10^{6}$ | $2.46\times10^{5}$ | $9.35\times10^{5}$ | $4.31\times10^{5}$ | $2.02\times10^{6}$ |
| DCTPP1 (C/NU/MT) | $3.62\times10^{3}$ | $5.95\times10^{4}$ | $1.34\times10^{3}$ | $2.94\times10^{4}$ | $1.19\times10^{4}$ | $1.56\times10^{5}$ |
| DNPEP (C) | $1.67\times10^{4}$ | $1.03\times10^{5}$ | $3.28\times10^{4}$ | $6.53\times10^{4}$ | $3.35\times10^{4}$ | $1.19\times10^{6}$ |
| ECI1 (MT) | $2.08\times10^{3}$ | $1.86\times10^{4}$ | $8.69\times10^{3}$ | $1.89\times10^{5}$ | $1.99\times10^{4}$ | $2.13\times10^{4}$ |
| EPPK1 (CS) | $5.86\times10^{5}$ | $1.71\times10^{5}$ | $6.65\times10^{5}$ | $6.14\times10^{5}$ | $2.07\times10^{5}$ | $4.63\times10^{5}$ |
| FLNA (CS) | $1.00\times10^{0}$ | $4.09\times10^{3}$ | $1.83\times10^{4}$ | $1.19\times10^{4}$ | $4.38\times10^{3}$ | $3.12\times10^{4}$ |
| GSR (C/MT) | $1.32\times10^{5}$ | $1.39\times10^{6}$ | $1.78\times10^{5}$ | $1.20\times10^{6}$ | $1.68\times10^{5}$ | $1.61\times10^{6}$ |
| HPRT1 (C) | $2.36\times10^{4}$ | $8.25\times10^{5}$ | $1.12\times10^{5}$ | $7.81\times10^{5}$ | $1.54\times10^{5}$ | $2.12\times10^{6}$ |
| HSD17B10 (MT) | $2.26\times10^{4}$ | $3.42\times10^{5}$ | $1.31\times10^{5}$ | $2.31\times10^{6}$ | $8.43\times10^{4}$ | $2.31\times10^{5}$ |
| HSPA9 (MT) | $7.57\times10^{5}$ | $1.38\times10^{6}$ | $9.05\times10^{5}$ | $2.26\times10^{6}$ | $1.35\times10^{6}$ | $1.24\times10^{6}$ |
| HSPB1 (C/NU/CS) | $1.09\times10^{6}$ | $5.39\times10^{5}$ | $1.21\times10^{6}$ | $1.98\times10^{6}$ | $7.64\times10^{5}$ | $1.54\times10^{6}$ |
| ISYNA1 (C) | $4.51\times10^{4}$ | $2.50\times10^{5}$ | $4.35\times10^{4}$ | $3.50\times10^{5}$ | $1.03\times10^{5}$ | $8.04\times10^{5}$ |
| LDHB (C/MT) | $1.49\times10^{5}$ | $5.61\times10^{5}$ | $3.48\times10^{5}$ | $1.34\times10^{5}$ | $1.03\times10^{6}$ | $1.43\times10^{5}$ |
| LMNA (NU) | $3.91\times10^{5}$ | $1.48\times10^{5}$ | $1.05\times10^{6}$ | $2.97\times10^{5}$ | $1.90\times10^{5}$ | $4.80\times10^{5}$ |
| LY6D (PM) | $2.99\times10^{4}$ | $3.62\times10^{4}$ | $4.99\times10^{4}$ | $4.79\times10^{4}$ | $2.66\times10^{4}$ | $7.30\times10^{4}$ |
| MARCKSL1(SC/PM) | $1.00\times10^{0}$ | $2.21\times10^{4}$ | $1.60\times10^{3}$ | $3.83\times10^{3}$ | $2.68\times10^{4}$ | $7.52\times10^{3}$ |
| MDH1 (C) | $4.83\times10^{4}$ | $2.59\times10^{5}$ | $1.06\times10^{5}$ | $3.41\times10^{4}$ | $9.89\times10^{4}$ | $4.91\times10^{4}$ |
| ME2 (MT) | $2.04\times10^{2}$ | $9.56\times10^{2}$ | $2.19\times10^{2}$ | $1.68\times10^{5}$ | $2.91\times10^{3}$ | $8.27\times10^{2}$ |
| MYH9 (CS) | $1.32\times10^{5}$ | $9.95\times10^{4}$ | $1.66\times10^{5}$ | $1.60\times10^{5}$ | $1.49\times10^{5}$ | $3.30\times10^{5}$ |
| NPEPPS (C/NU) | $1.34\times10^{5}$ | $2.45\times10^{5}$ | $3.03\times10^{5}$ | $4.06\times10^{5}$ | $1.93\times10^{5}$ | $9.54\times10^{5}$ |
| NUDC (CS/NU) | $3.45\times10^{4}$ | $1.18\times10^{5}$ | $2.69\times10^{4}$ | $1.16\times10^{5}$ | $8.20\times10^{4}$ | $6.18\times10^{5}$ |
| PKP1 (NU) | $1.36\times10^{6}$ | $6.65\times10^{5}$ | $1.39\times10^{6}$ | $1.39\times10^{6}$ | $7.37\times10^{5}$ | $1.85\times10^{6}$ |
| PLS3 (C) | $3.57\times10^{4}$ | $1.11\times10^{5}$ | $3.82\times10^{4}$ | $8.61\times10^{4}$ | $4.92\times10^{4}$ | $3.08\times10^{5}$ |
| PRDX1 (C) | $5.17\times10^{6}$ | $1.63\times10^{7}$ | $7.47\times10^{6}$ | $1.71\times10^{7}$ | $5.58\times10^{6}$ | $5.09\times10^{7}$ |
| PRDX2 (C) | $2.02\times10^{6}$ | $2.48\times10^{6}$ | $2.93\times10^{6}$ | $3.79\times10^{6}$ | $1.88\times10^{6}$ | $6.70\times10^{6}$ |
| PRDX3 (MT/C/EE) | $1.27\times10^{5}$ | $7.89\times10^{5}$ | $3.75\times10^{5}$ | $2.42\times10^{6}$ | $2.51\times10^{5}$ | $3.08\times10^{5}$ |
| PRDX4 (C/ER) | $1.79\times10^{4}$ | $6.47\times10^{4}$ | $3.19\times10^{4}$ | $5.49\times10^{4}$ | $5.70\times10^{4}$ | $8.75\times10^{4}$ |
| PRDX5 (C/NU/MT/PO) | $2.19\times10^{3}$ | $4.48\times10^{4}$ | $4.33\times10^{3}$ | $5.38\times10^{4}$ | $5.15\times10^{3}$ | $3.34\times10^{4}$ |
| PRDX6 (C/LY) | $3.15\times10^{4}$ | $4.64\times10^{5}$ | $6.49\times10^{4}$ | $4.98\times10^{5}$ | $2.02\times10^{5}$ | $1.38\times10^{6}$ |
| PRMT5 (C/NU/GA) | $3.15\times10^{4}$ | $2.93\times10^{5}$ | $3.78\times10^{4}$ | $2.72\times10^{5}$ | $4.36\times10^{4}$ | $6.38\times10^{5}$ |
| RPL13 (C) | $1.37\times10^{4}$ | $3.53\times10^{4}$ | $1.52\times10^{4}$ | $4.31\times10^{3}$ | $8.62\times10^{4}$ | $1.44\times10^{4}$ |
| RPL27A (C/ER) | $6.83\times10^{3}$ | $1.55\times10^{4}$ | $6.91\times10^{3}$ | $3.14\times10^{3}$ | $4.13\times10^{4}$ | $1.05\times10^{4}$ |
| S100B (C/NU) | $1.94\times10^{6}$ | $1.17\times10^{6}$ | $2.50\times10^{6}$ | $5.16\times10^{6}$ | $2.34\times10^{6}$ | $1.44\times10^{6}$ |
| SERPINB4 (C) | $5.62\times10^{4}$ | $4.89\times10^{4}$ | $5.41\times10^{4}$ | $1.30\times10^{5}$ | $3.94\times10^{4}$ | $4.31\times10^{4}$ |
| SERPINB7 (C) | $3.65\times10^{4}$ | $9.41\times10^{3}$ | $9.92\times10^{4}$ | $1.86\times10^{4}$ | $2.15\times10^{4}$ | $4.33\times10^{4}$ |
| SET (C/NU) | $3.72\times10^{3}$ | $1.63\times10^{4}$ | $2.51\times10^{4}$ | $8.70\times10^{3}$ | $5.27\times10^{4}$ | $2.12\times10^{3}$ |
| SKP1 (C/NU) | $4.16\times10^{4}$ | $1.91\times10^{6}$ | $2.55\times10^{4}$ | $1.05\times10^{6}$ | $3.01\times10^{4}$ | $2.90\times10^{6}$ |
| TPM3 (CS) | $1.27\times10^{5}$ | $4.22\times10^{5}$ | $2.43\times10^{5}$ | $4.41\times10^{5}$ | $2.40\times10^{5}$ | $1.12\times10^{6}$ |
| TUBB1 (CS) | $4.34\times10^{4}$ | $4.82\times10^{4}$ | $4.92\times10^{4}$ | $3.63\times10^{4}$ | $1.26\times10^{5}$ | $4.51\times10^{5}$ |
| TXN (C/NU/EC) | $3.07\times10^{6}$ | $4.75\times10^{6}$ | $4.52\times10^{6}$ | $5.42\times10^{6}$ | $2.55\times10^{6}$ | $7.96\times10^{6}$ |
| TXNRD2 (MT) | $1.00\times10^{0}$ | $1.00\times10^{0}$ | $5.93\times10^{4}$ | $3.82\times10^{6}$ | $1.00\times10^{0}$ | $1.00\times10^{0}$ |
| UCHL3 (C) | $6.47\times10^{3}$ | $3.39\times10^{4}$ | $1.14\times10^{4}$ | $4.26\times10^{4}$ | $1.46\times10^{4}$ | $3.94\times10^{5}$ |
| WDR77 (C/NU) | $2.57\times10^{4}$ | $2.79\times10^{5}$ | $4.21\times10^{4}$ | $2.42\times10^{5}$ | $2.98\times10^{4}$ | $7.54\times10^{5}$ |
| YWHAB (C) | $4.41\times10^{4}$ | $1.56\times10^{5}$ | $6.83\times10^{4}$ | $1.01\times10^{5}$ | $1.25\times10^{5}$ | $2.07\times10^{5}$ |
| YWHAE (C/NU) | $2.22\times10^{4}$ | $1.15\times10^{5}$ | $4.40\times10^{4}$ | $8.15\times10^{4}$ | $1.33\times10^{5}$ | $2.28\times10^{5}$ |
| YWHAQ (C) | $1.49\times10^{4}$ | $2.53\times10^{5}$ | $3.05\times10^{4}$ | $1.26\times10^{5}$ | $7.34\times10^{4}$ | $2.76\times10^{5}$ |
| YWHAZ (C) | $2.08\times10^{5}$ | $4.22\times10^{5}$ | $3.01\times10^{5}$ | $2.96\times10^{5}$ | $4.92\times10^{5}$ | $7.67\times10^{5}$ |

**mM $H_2O_2$**
**(10 min)**

#### Table S2

| Protein | po-YAP | mt-YAP | c-YAP |
|---|---|---|---|
| ACOX1 (PO) | 85 | 5 | 10 |
| BOLA2B (C/NU) | 29 | 21 | 50 |
| CDK4 (C/NU) | 15 | 7 | 78 |
| CSTB (C/NU) | 30 | 22 | 48 |
| DCTPP1 (C/NU/MT) | 24 | 12 | 64 |
| DNPEP (C) | 8 | 5 | 88 |
| ECI1 (MT) | 8 | 83 | 9 |
| EPPK1 (CS) | 14 | 49 | 37 |
| FLNA (CS) | 9 | 25 | 66 |
| GSR (C/MT) | 33 | 29 | 38 |
| HPRT1 (C) | 22 | 21 | 57 |
| HSD17B10 (MT) | 12 | 80 | 8 |
| HSPA9 (MT) | 28 | 46 | 25 |
| HSPB1 (C/NU/CS) | 13 | 49 | 38 |
| ISYNA1 (C) | 18 | 25 | 57 |
| LDHB (C/MT) | 67 | 16 | 17 |
| LMNA (NU) | 16 | 32 | 52 |
| LY6D (PM) | 23 | 30 | 46 |
| MARCKSL1(SC/PM) | 66 | 11 | 22 |
| MDH1 (C) | 76 | 10 | 14 |
| ME2 (MT) | 1 | 99 | 0 |
| MYH9 (CS) | 17 | 27 | 56 |
| NPEPPS (C/NU) | 15 | 25 | 59 |
| NUDC (CS/NU) | 14 | 14 | 73 |
| PKP1 (NU) | 17 | 36 | 47 |
| PLS3 (C) | 22 | 17 | 61 |
| PRDX1 (C) | 19 | 20 | 60 |
| PRDX2 (C) | 19 | 29 | 52 |
| PRDX3 (MT/C/EE) | 22 | 69 | 9 |
| PRDX4 (C/ER) | 31 | 27 | 42 |
| PRDX5 (C/NU/MT/PO) | 34 | 41 | 25 |
| PRDX6 (C/LY) | 20 | 21 | 59 |
| PRMT5 (C/NU/GA) | 24 | 23 | 53 |
| RPL13 (C) | 65 | 8 | 27 |
| RPL27A (C/ER) | 53 | 11 | 36 |
| S100B (C/NU) | 15 | 66 | 19 |
| SERPINB4 (C) | 22 | 59 | 19 |
| SERPINB7 (C) | 13 | 26 | 61 |
| SET (C/NU) | 60 | 32 | 8 |
| SKP1 (C/NU) | 33 | 18 | 50 |
| TPM3 (CS) | 21 | 22 | 56 |
| TUBB1 (CS) | 9 | 7 | 84 |
| TXN (C/NU/EC) | 26 | 30 | 44 |
| TXNRD2 (MT) | 0 | 100 | 0 |
| UCHL3 (C) | 7 | 9 | 84 |
| WDR77 (C/NU) | 22 | 19 | 59 |
| YWHAB (C) | 34 | 22 | 45 |
| YWHAE (C/NU) | 27 | 19 | 54 |
| YWHAQ (C) | 39 | 19 | 42 |
| YWHAZ (C) | 28 | 20 | 52 |

percentage distribution (color scale 0–100)

1 mM $H_2O_2$ (10 min)

### Table S3 (page 1)

#### po-$H_2O_2$ / c-YAP1C

raw abundance: $1\times10^{7}$, $8\times10^{6}$, $6\times10^{6}$, $4\times10^{6}$, $2\times10^{6}$, 0

| 0 | 2 | 5 | 15 | 30 | 60 | |
|---|---|---|---|---|---|---|
| $6.81\times10^{2}$ | $1.02\times10^{3}$ | $5.81\times10^{4}$ | $5.93\times10^{4}$ | $3.68\times10^{2}$ | $1.70\times10^{3}$ | ACADVL (MT) |
| 0 | $1.63\times10^{2}$ | $3.95\times10^{4}$ | $2.15\times10^{3}$ | $1.52\times10^{2}$ | 0 | ACP3 (NU/C/PM/LY/EC) |
| $1.08\times10^{6}$ | $1.36\times10^{6}$ | $1.08\times10^{7}$ | $1.45\times10^{7}$ | $2.79\times10^{6}$ | $2.03\times10^{6}$ | ACTB (NU/CS) |
| $1.24\times10^{4}$ | $1.61\times10^{4}$ | $1.27\times10^{5}$ | $2.15\times10^{5}$ | $4.79\times10^{4}$ | $1.00\times10^{4}$ | ACTC1 (CS) |
| $2.86\times10^{2}$ | $3.14\times10^{2}$ | $1.75\times10^{4}$ | $2.17\times10^{4}$ | $5.69\times10^{2}$ | $3.82\times10^{3}$ | ACTN1 (C/PM) |
| $8.05\times10^{4}$ | $1.11\times10^{5}$ | $2.06\times10^{6}$ | $1.89\times10^{6}$ | $8.26\times10^{4}$ | $8.71\times10^{4}$ | ACTN4 (C/NU/CS) |
| 0 | $1.84\times10^{3}$ | $1.58\times10^{5}$ | $9.38\times10^{4}$ | $1.23\times10^{4}$ | $1.82\times10^{3}$ | ACTR1A (CS) |
| $2.49\times10^{4}$ | $9.61\times10^{3}$ | $3.51\times10^{5}$ | $3.20\times10^{5}$ | $2.52\times10^{4}$ | $1.34\times10^{4}$ | ACTR2 (NU/CS) |
| $1.62\times10^{4}$ | $2.37\times10^{4}$ | $3.66\times10^{5}$ | $2.70\times10^{5}$ | $1.30\times10^{4}$ | $3.83\times10^{4}$ | ACTR3 (NU/C/CS) |
| $2.48\times10^{4}$ | $3.14\times10^{4}$ | $2.09\times10^{5}$ | $7.38\times10^{4}$ | $4.42\times10^{4}$ | $7.06\times10^{4}$ | AHCY (C) |
| $2.63\times10^{5}$ | $2.11\times10^{5}$ | $2.29\times10^{6}$ | $5.37\times10^{6}$ | $3.19\times10^{5}$ | $1.05\times10^{5}$ | AHNAK (NU) |
| $1.92\times10^{3}$ | $4.02\times10^{3}$ | $4.47\times10^{4}$ | $2.47\times10^{5}$ | $2.83\times10^{3}$ | $2.79\times10^{3}$ | AHNAK2 (NU) |
| $6.08\times10^{2}$ | $1.68\times10^{2}$ | $2.13\times10^{4}$ | $1.77\times10^{4}$ | $5.07\times10^{2}$ | $1.32\times10^{2}$ | AIMP1 (C/NU/ER/GA/PM) |
| $1.26\times10^{3}$ | $4.68\times10^{3}$ | $1.16\times10^{5}$ | $1.13\times10^{4}$ | $2.63\times10^{3}$ | $5.56\times10^{3}$ | AKR1B10 (LY/EC) |
| $5.15\times10^{3}$ | $1.76\times10^{4}$ | $2.10\times10^{5}$ | $3.82\times10^{5}$ | $9.54\times10^{3}$ | $5.58\times10^{3}$ | ALDH2 (MT) |
| $4.26\times10^{2}$ | $1.53\times10^{3}$ | $6.39\times10^{4}$ | $2.05\times10^{5}$ | $3.80\times10^{3}$ | $1.67\times10^{3}$ | ALDH3A1 (C) |
| $6.69\times10^{3}$ | $1.58\times10^{4}$ | $1.83\times10^{5}$ | $1.97\times10^{5}$ | $5.87\times10^{3}$ | $2.08\times10^{4}$ | ALDH3A2 (ER) |
| $4.62\times10^{4}$ | $4.44\times10^{4}$ | $4.54\times10^{5}$ | $2.18\times10^{5}$ | $4.55\times10^{4}$ | $4.60\times10^{4}$ | ALDH9A1 (C) |
| $1.63\times10^{5}$ | $2.84\times10^{5}$ | $2.81\times10^{6}$ | $2.37\times10^{6}$ | $2.54\times10^{5}$ | $2.94\times10^{5}$ | ALDOA (C/NU) |
| $6.40\times10^{3}$ | $1.94\times10^{4}$ | $5.20\times10^{5}$ | $2.64\times10^{5}$ | $1.10\times10^{4}$ | $2.03\times10^{4}$ | ALDOC (C/CS/EC) |
| $2.47\times10^{5}$ | $1.45\times10^{5}$ | $1.44\times10^{6}$ | $3.95\times10^{5}$ | $1.64\times10^{5}$ | $6.05\times10^{4}$ | ALOX12B (C) |
| $9.33\times10^{3}$ | $1.27\times10^{4}$ | $9.55\times10^{4}$ | $2.91\times10^{4}$ | $1.78\times10^{4}$ | $6.32\times10^{3}$ | ALOXE3 (C) |
| $8.37\times10^{4}$ | $1.74\times10^{5}$ | $9.43\times10^{4}$ | $1.54\times10^{5}$ | $1.76\times10^{5}$ | $2.47\times10^{5}$ | ALYREF (C/NU) |
| $3.96\times10^{5}$ | $1.52\times10^{5}$ | $1.02\times10^{6}$ | $4.48\times10^{6}$ | $7.13\times10^{5}$ | $6.74\times10^{4}$ | ANXA1 (C/NU/PM/EC) |
| $2.34\times10^{6}$ | $1.58\times10^{6}$ | $1.25\times10^{7}$ | $1.07\times10^{7}$ | $2.43\times10^{6}$ | $2.23\times10^{6}$ | ANXA2 (PM/EC) |
| $3.82\times10^{3}$ | $6.58\times10^{3}$ | $3.75\times10^{5}$ | $2.82\times10^{5}$ | $5.59\times10^{3}$ | $5.81\times10^{3}$ | ANXA4 (C/NU/PM/EC) |
| $1.61\times10^{4}$ | $1.52\times10^{4}$ | $1.01\times10^{5}$ | $1.30\times10^{5}$ | $3.52\times10^{4}$ | $7.02\times10^{4}$ | ANXA5 (C/EC) |
| $3.00\times10^{3}$ | $3.15\times10^{3}$ | $4.03\times10^{5}$ | $3.96\times10^{5}$ | $5.98\times10^{3}$ | $5.87\times10^{3}$ | ANXA8 (PM, ES) |
| $1.02\times10^{5}$ | $4.89\times10^{4}$ | $1.53\times10^{5}$ | $2.39\times10^{6}$ | $4.92\times10^{4}$ | $1.97\times10^{4}$ | APOA1 (C/PM/EC) |
| $5.94\times10^{2}$ | $3.89\times10^{2}$ | $1.23\times10^{2}$ | $3.51\times10^{4}$ | 0 | $4.88\times10^{0}$ | APOBEC3B (NU) |
| $1.38\times10^{4}$ | $4.44\times10^{4}$ | $5.05\times10^{4}$ | $1.03\times10^{5}$ | $7.16\times10^{3}$ | $1.79\times10^{4}$ | ARF3 (GA/C) |
| $1.53\times10^{3}$ | $5.10\times10^{2}$ | $2.18\times10^{4}$ | $4.64\times10^{4}$ | 0 | $7.06\times10^{2}$ | ARF6 (C/PM/GA/ES) |
| $1.91\times10^{6}$ | $1.34\times10^{6}$ | $8.59\times10^{6}$ | $7.93\times10^{5}$ | $1.95\times10^{6}$ | $5.59\times10^{5}$ | ARG1 (C) |
| $6.48\times10^{3}$ | $1.63\times10^{4}$ | $1.62\times10^{5}$ | $1.49\times10^{5}$ | $7.18\times10^{3}$ | $1.43\times10^{5}$ | ARHGDIA (C) |
| $2.06\times10^{3}$ | $9.92\times10^{2}$ | $1.28\times10^{5}$ | $1.28\times10^{5}$ | $6.54\times10^{2}$ | $9.61\times10^{2}$ | ARPC3 (NU/CS) |
| $1.80\times10^{4}$ | $1.48\times10^{4}$ | $9.49\times10^{4}$ | $3.93\times10^{4}$ | $2.80\times10^{4}$ | $5.34\times10^{3}$ | ASAH1 (C/LY/EC) |
| $5.03\times10^{0}$ | $5.20\times10^{2}$ | $1.67\times10^{5}$ | $1.24\times10^{4}$ | $8.28\times10^{2}$ | $4.48\times10^{2}$ | ASPRV1 (C/NU) |
| $1.28\times10^{3}$ | $7.75\times10^{2}$ | $1.09\times10^{4}$ | $7.26\times10^{4}$ | $1.12\times10^{3}$ | $1.89\times10^{4}$ | ATP2A2 (ER) |
| $8.85\times10^{4}$ | $1.15\times10^{5}$ | $4.24\times10^{5}$ | $1.15\times10^{6}$ | $1.42\times10^{5}$ | $1.37\times10^{5}$ | ATP5F1A (MT) |
| $2.13\times10^{5}$ | $3.00\times10^{5}$ | $5.90\times10^{5}$ | $1.63\times10^{6}$ | $2.78\times10^{5}$ | $2.83\times10^{5}$ | ATP5F1B (MT) |
| $1.05\times10^{3}$ | $8.45\times10^{3}$ | $1.15\times10^{4}$ | $5.26\times10^{4}$ | $7.16\times10^{3}$ | $8.41\times10^{3}$ | ATP5F1C (MT) |
| $1.15\times10^{3}$ | $1.16\times10^{3}$ | $1.21\times10^{4}$ | $1.29\times10^{5}$ | $1.29\times10^{3}$ | $5.42\times10^{3}$ | ATP5PB (MT) |
| 0 | 0 | $3.64\times10^{2}$ | $7.72\times10^{3}$ | 0 | 0 | ATP5PD (MT) |
| $7.67\times10^{3}$ | $2.33\times10^{4}$ | $1.22\times10^{4}$ | $1.98\times10^{5}$ | $1.64\times10^{4}$ | $3.48\times10^{4}$ | ATP5PO (MT) |
| $2.80\times10^{3}$ | $1.96\times10^{3}$ | $1.99\times10^{5}$ | $1.68\times10^{5}$ | $1.89\times10^{3}$ | $5.09\times10^{3}$ | ATP6V1A (C) |
| $3.75\times10^{3}$ | $9.53\times10^{2}$ | $2.70\times10^{4}$ | $2.23\times10^{4}$ | $8.53\times10^{3}$ | $1.47\times10^{3}$ | ATP6V1G1 (PM) |
| $3.24\times10^{5}$ | $3.77\times10^{5}$ | $3.93\times10^{6}$ | $5.96\times10^{5}$ | $7.93\times10^{5}$ | $1.88\times10^{5}$ | BLMH (C) |
| $2.74\times10^{3}$ | $4.52\times10^{3}$ | $5.29\times10^{4}$ | $2.11\times10^{5}$ | $3.15\times10^{3}$ | $3.02\times10^{3}$ | CA2 (C/PM) |
| $2.81\times10^{4}$ | $7.90\times10^{4}$ | $7.40\times10^{5}$ | $2.08\times10^{5}$ | $2.29\times10^{4}$ | $2.54\times10^{4}$ | CALM2 (C) |
| $1.62\times10^{4}$ | $4.18\times10^{4}$ | $1.74\times10^{5}$ | $1.83\times10^{5}$ | $7.71\times10^{4}$ | $9.49\times10^{4}$ | CALR (C/ER/LY/EC) |

#### Table S3
#### (page 2)

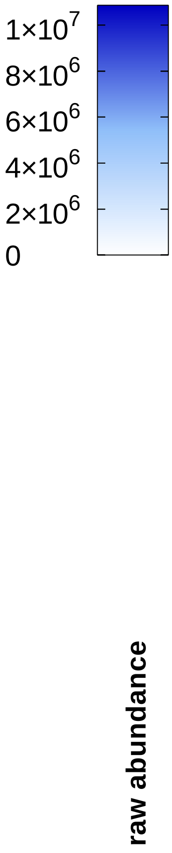


**po-$H_2O_2$ / c-YAP1C**

| Protein | 0 | 2 | 5 | 15 | 30 | 60 |
|---|---|---|---|---|---|---|
| CAND1 (C/NU) | $4.24\times10^{2}$ | $1.66\times10^{2}$ | $3.33\times10^{3}$ | $3.53\times10^{4}$ | $6.82\times10^{2}$ | $1.50\times10^{2}$ |
| CANX (ER) | $3.72\times10^{3}$ | $1.05\times10^{4}$ | $5.26\times10^{4}$ | $1.26\times10^{5}$ | $1.66\times10^{4}$ | $3.32\times10^{4}$ |
| CAP1 (PM) | $4.51\times10^{3}$ | $2.42\times10^{3}$ | $1.70\times10^{5}$ | $3.56\times10^{5}$ | $1.12\times10^{3}$ | $5.58\times10^{3}$ |
| CAPG (C/NU) | $5.62\times10^{3}$ | $4.02\times10^{3}$ | $3.18\times10^{5}$ | $3.57\times10^{5}$ | $8.72\times10^{2}$ | $2.99\times10^{4}$ |
| CAPN1 (C/PM) | $9.82\times10^{4}$ | $1.04\times10^{5}$ | $8.54\times10^{5}$ | $5.05\times10^{5}$ | $8.27\times10^{4}$ | $6.97\times10^{4}$ |
| CAPZA1 (CS) | $2.30\times10^{4}$ | $2.62\times10^{4}$ | $7.35\times10^{5}$ | $3.61\times10^{5}$ | $2.51\times10^{4}$ | $2.80\times10^{4}$ |
| CAPZB (CS) | $6.74\times10^{4}$ | $5.01\times10^{4}$ | $5.04\times10^{5}$ | $2.81\times10^{5}$ | $3.85\times10^{4}$ | $6.45\times10^{4}$ |
| CARD18 (???) | $6.04\times10^{2}$ | $3.25\times10^{2}$ | $4.03\times10^{4}$ | $6.92\times10^{3}$ | 0 | 0 |
| CASP14 (C/NU) | $8.14\times10^{5}$ | $8.07\times10^{5}$ | $7.24\times10^{6}$ | $1.25\times10^{6}$ | $1.14\times10^{6}$ | $2.51\times10^{5}$ |
| CAT (PO) | $8.83\times10^{5}$ | $5.19\times10^{5}$ | $3.67\times10^{6}$ | $5.77\times10^{5}$ | $1.30\times10^{6}$ | $2.75\times10^{5}$ |
| CBR1 (C) | $1.65\times10^{3}$ | $5.15\times10^{3}$ | $1.89\times10^{5}$ | $2.38\times10^{5}$ | $1.12\times10^{4}$ | $1.62\times10^{4}$ |
| CBX1 (NU) | $9.35\times10^{3}$ | $2.10\times10^{4}$ | $3.41\times10^{4}$ | $3.54\times10^{4}$ | $1.79\times10^{4}$ | $1.86\times10^{4}$ |
| CCT2 (C) | $6.28\times10^{3}$ | $5.20\times10^{3}$ | $2.60\times10^{4}$ | $1.19\times10^{5}$ | $5.52\times10^{3}$ | $8.19\times10^{3}$ |
| CCT4 (C/CS) | $1.46\times10^{3}$ | $9.07\times10^{4}$ | $6.46\times10^{4}$ | $1.66\times10^{5}$ | $3.12\times10^{4}$ | $2.50\times10^{4}$ |
| CCT5 (C/CS) | $3.66\times10^{3}$ | $3.54\times10^{3}$ | $3.49\times10^{4}$ | $3.71\times10^{4}$ | $3.14\times10^{3}$ | $3.72\times10^{3}$ |
| CCT6A (C) | $1.09\times10^{2}$ | $1.06\times10^{3}$ | $3.72\times10^{4}$ | $1.28\times10^{5}$ | $1.37\times10^{2}$ | $2.90\times10^{3}$ |
| CCT8 (C/CS) | $6.78\times10^{3}$ | $4.40\times10^{3}$ | $1.27\times10^{5}$ | $2.06\times10^{5}$ | $2.62\times10^{3}$ | $9.72\times10^{3}$ |
| CD44 (PM) | 0 | 0 | 0 | $4.88\times10^{4}$ | 0 | 0 |
| CDK4 (C/NU) | $6.53\times10^{3}$ | $1.28\times10^{3}$ | $2.85\times10^{4}$ | $9.11\times10^{4}$ | $4.26\times10^{4}$ | $2.47\times10^{5}$ |
| CDKN2A (C/NU) | $4.87\times10^{2}$ | $9.10\times10^{2}$ | $3.03\times10^{3}$ | $1.73\times10^{4}$ | $4.67\times10^{4}$ | $2.48\times10^{5}$ |
| CFL1 (C/NU/CS) | $9.27\times10^{4}$ | $2.98\times10^{5}$ | $5.77\times10^{5}$ | $1.24\times10^{6}$ | $1.69\times10^{5}$ | $4.07\times10^{5}$ |
| CHMP4B (C/NU/ES) | 0 | $6.73\times10^{2}$ | $2.03\times10^{4}$ | $2.89\times10^{4}$ | 0 | $2.00\times10^{2}$ |
| CHTOP (NU) | $3.18\times10^{4}$ | $5.16\times10^{4}$ | $8.04\times10^{4}$ | $9.06\times10^{4}$ | $1.30\times10^{5}$ | $1.08\times10^{5}$ |
| CKB (C) | $8.29\times10^{4}$ | $9.00\times10^{4}$ | $1.55\times10^{5}$ | $2.76\times10^{5}$ | $3.26\times10^{5}$ | $3.75\times10^{5}$ |
| CKMT1A (MT) | $8.28\times10^{3}$ | $7.21\times10^{3}$ | $7.90\times10^{4}$ | $1.22\times10^{5}$ | $1.55\times10^{4}$ | $6.21\times10^{3}$ |
| CLIC1 (C/NU/PL) | $4.16\times10^{4}$ | $4.06\times10^{4}$ | $1.86\times10^{5}$ | $5.26\times10^{5}$ | $1.92\times10^{4}$ | $3.79\times10^{4}$ |
| CLTC (CS/VS) | $4.91\times10^{4}$ | $2.71\times10^{4}$ | $4.98\times10^{5}$ | $1.26\times10^{6}$ | $1.82\times10^{4}$ | $3.22\times10^{4}$ |
| CMPK1 (C/NU) | $3.25\times10^{2}$ | $2.02\times10^{3}$ | $1.16\times10^{5}$ | $5.81\times10^{4}$ | $7.59\times10^{3}$ | $7.55\times10^{2}$ |
| COX7A2 (MT) | $1.03\times10^{4}$ | $1.82\times10^{4}$ | $3.00\times10^{4}$ | $6.15\times10^{4}$ | $2.22\times10^{4}$ | $1.56\times10^{4}$ |
| CRYZL1 (C) | $1.72\times10^{2}$ | 0 | $2.61\times10^{3}$ | $6.10\times10^{2}$ | $7.77\times10^{1}$ | $2.01\times10^{4}$ |
| CS (MT) | $1.58\times10^{4}$ | $4.27\times10^{3}$ | $1.71\times10^{4}$ | $1.54\times10^{5}$ | $1.02\times10^{4}$ | $8.86\times10^{3}$ |
| CSNK1A1 (C/NU/CS) | $5.46\times10^{2}$ | $1.81\times10^{3}$ | $2.32\times10^{4}$ | $1.09\times10^{5}$ | $7.43\times10^{3}$ | $4.19\times10^{3}$ |
| CSTA (C) | $1.36\times10^{6}$ | $3.42\times10^{5}$ | $4.04\times10^{6}$ | $4.45\times10^{5}$ | $1.32\times10^{6}$ | $2.76\times10^{5}$ |
| CSTB (C/NU) | $1.14\times10^{5}$ | $9.89\times10^{4}$ | $4.32\times10^{5}$ | $9.56\times10^{5}$ | $4.05\times10^{5}$ | $8.33\times10^{5}$ |
| CTNNA1 (CS/PM) | $2.22\times10^{3}$ | $1.98\times10^{3}$ | $2.48\times10^{4}$ | $6.40\times10^{4}$ | $1.03\times10^{3}$ | $6.92\times10^{2}$ |
| CTNND1 (C/NU/PM) | $5.99\times10^{2}$ | $2.27\times10^{2}$ | $6.26\times10^{3}$ | $4.47\times10^{4}$ | $1.84\times10^{2}$ | $4.20\times10^{2}$ |
| CTSA (LY) | $3.24\times10^{4}$ | $2.01\times10^{4}$ | $2.79\times10^{5}$ | $2.15\times10^{4}$ | $3.76\times10^{4}$ | $7.44\times10^{3}$ |
| CTSD (LY/EC) | $2.66\times10^{5}$ | $4.52\times10^{5}$ | $5.55\times10^{6}$ | $7.30\times10^{5}$ | $5.17\times10^{5}$ | $1.38\times10^{5}$ |
| CYB5R3 (C/MT/ER) | $1.16\times10^{4}$ | $3.85\times10^{3}$ | $1.32\times10^{4}$ | $6.85\times10^{4}$ | $6.11\times10^{3}$ | $1.98\times10^{4}$ |
| CYCS (MT) | $5.57\times10^{3}$ | $1.22\times10^{4}$ | $1.96\times10^{4}$ | $2.42\times10^{4}$ | $3.32\times10^{2}$ | $4.55\times10^{2}$ |
| CYFIP2 (C/NU) | $7.98\times10^{2}$ | $1.01\times10^{3}$ | $1.91\times10^{4}$ | $3.88\times10^{4}$ | $1.39\times10^{2}$ | $2.72\times10^{2}$ |
| DCTN1 (C/NU/CS) | $6.29\times10^{3}$ | $7.88\times10^{3}$ | $4.92\times10^{4}$ | $9.52\times10^{4}$ | $5.60\times10^{3}$ | $4.65\times10^{3}$ |
| DCTN2 (CS) | 0 | 0 | 0 | $1.61\times10^{4}$ | $5.12\times10^{2}$ | $1.89\times10^{2}$ |
| DCTPP1 (C/NU/MT) | $2.07\times10^{2}$ | $2.00\times10^{3}$ | $1.60\times10^{3}$ | $3.64\times10^{3}$ | $3.81\times10^{3}$ | $2.88\times10^{4}$ |
| DDOST (ER) | $9.14\times10^{3}$ | $7.99\times10^{3}$ | $9.11\times10^{4}$ | $6.61\times10^{4}$ | $9.83\times10^{3}$ | $1.23\times10^{4}$ |
| DDX39B (C/NU) | $3.84\times10^{3}$ | $1.06\times10^{4}$ | $5.73\times10^{4}$ | $1.21\times10^{5}$ | $1.11\times10^{4}$ | $5.69\times10^{4}$ |
| DDX3X (C/CS/NU/PM) | $5.97\times10^{3}$ | $5.50\times10^{3}$ | $1.62\times10^{5}$ | $2.87\times10^{5}$ | $7.34\times10^{3}$ | $5.79\times10^{3}$ |
| DHX9 (C/NU/CS) | $7.35\times10^{2}$ | $1.02\times10^{3}$ | $3.17\times10^{4}$ | $4.27\times10^{4}$ | $1.09\times10^{4}$ | $5.66\times10^{1}$ |
| DLD (NU/MT) | $4.60\times10^{4}$ | $7.77\times10^{4}$ | $2.99\times10^{5}$ | $1.13\times10^{5}$ | $1.23\times10^{5}$ | $8.00\times10^{4}$ |
| DNAJB1 (C/NU) | $1.74\times10^{2}$ | $3.66\times10^{2}$ | $3.83\times10^{3}$ | $3.81\times10^{4}$ | $8.09\times10^{2}$ | $1.18\times10^{3}$ |

### Table S3 (page 3)

**po-$H_2O_2$ / c-YAP1C**

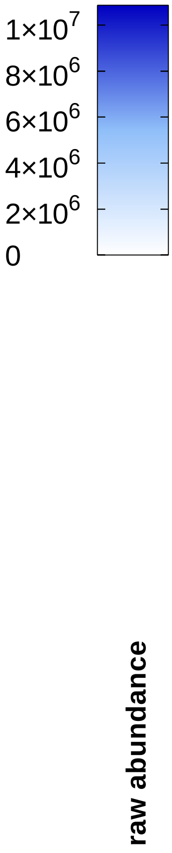


| Protein | 0 | 2 | 5 | 15 | 30 | 60 |
|---|---|---|---|---|---|---|
| DSG1 (PM) | $1.02\times10^{7}$ | $6.30\times10^{6}$ | $5.22\times10^{7}$ | $1.00\times10^{7}$ | $1.07\times10^{7}$ | $3.47\times10^{6}$ |
| DSG3 (PM) | $3.52\times10^{2}$ | $7.61\times10^{2}$ | $2.15\times10^{3}$ | $1.73\times10^{5}$ | $4.16\times10^{2}$ | 0 |
| DSP (CS/PM) | $1.99\times10^{7}$ | $9.66\times10^{6}$ | $9.57\times10^{7}$ | $4.26\times10^{7}$ | $1.16\times10^{7}$ | $5.37\times10^{6}$ |
| DSTN (C/CS/EC) | $1.24\times10^{4}$ | $1.50\times10^{4}$ | $8.89\times10^{4}$ | $1.90\times10^{5}$ | $2.21\times10^{4}$ | $1.21\times10^{4}$ |
| DYNC1H1 (CS) | $5.90\times10^{3}$ | $4.80\times10^{3}$ | $7.82\times10^{4}$ | $2.28\times10^{5}$ | $7.26\times10^{3}$ | $3.11\times10^{3}$ |
| DYNLL1 (NU/MF/CS) | $3.23\times10^{2}$ | $1.88\times10^{3}$ | $3.47\times10^{4}$ | $6.23\times10^{4}$ | $7.74\times10^{2}$ | $7.69\times10^{2}$ |
| EEF1A1 (C/NU/PM) | $3.37\times10^{5}$ | $4.52\times10^{5}$ | $4.96\times10^{6}$ | $8.75\times10^{6}$ | $4.65\times10^{5}$ | $1.75\times10^{6}$ |
| EEF1B2 (C) | $1.46\times10^{4}$ | $2.45\times10^{4}$ | $8.07\times10^{4}$ | $1.38\times10^{5}$ | $1.34\times10^{4}$ | $5.63\times10^{4}$ |
| EEF1G (C/NU/EC) | $6.61\times10^{4}$ | $9.77\times10^{4}$ | $8.17\times10^{5}$ | $9.82\times10^{5}$ | $7.20\times10^{4}$ | $1.74\times10^{5}$ |
| EEF2 (C/NU) | $2.20\times10^{5}$ | $1.79\times10^{5}$ | $2.77\times10^{6}$ | $2.29\times10^{6}$ | $1.96\times10^{5}$ | $5.64\times10^{5}$ |
| EIF3H (C) | $1.08\times10^{4}$ | $5.31\times10^{2}$ | $2.70\times10^{4}$ | $2.10\times10^{4}$ | $4.26\times10^{3}$ | $2.42\times10^{3}$ |
| EIF3L (C) | $1.67\times10^{3}$ | $1.34\times10^{3}$ | $1.88\times10^{3}$ | $1.74\times10^{4}$ | $8.15\times10^{2}$ | $3.89\times10^{3}$ |
| EIF4A1 (C/EC) | $3.72\times10^{4}$ | $2.72\times10^{4}$ | $8.92\times10^{5}$ | $8.42\times10^{5}$ | $2.25\times10^{4}$ | $6.95\times10^{4}$ |
| EIF5A (C/NU/ER) | $3.72\times10^{3}$ | $9.85\times10^{3}$ | $8.93\times10^{4}$ | $2.12\times10^{5}$ | $2.98\times10^{3}$ | $2.57\times10^{4}$ |
| EIF6 (C/NU) | $8.19\times10^{4}$ | $1.26\times10^{5}$ | $5.39\times10^{5}$ | $2.56\times10^{5}$ | $1.33\times10^{5}$ | $6.57\times10^{4}$ |
| ENO1 (C/NU) | $9.15\times10^{5}$ | $1.56\times10^{6}$ | $9.21\times10^{6}$ | $3.77\times10^{6}$ | $8.61\times10^{5}$ | $1.44\times10^{6}$ |
| EPPK1 (CS) | $1.75\times10^{5}$ | $1.28\times10^{5}$ | $4.16\times10^{6}$ | $5.75\times10^{6}$ | $1.83\times10^{5}$ | $6.95\times10^{4}$ |
| EPRS1 (C/PM) | $3.94\times10^{3}$ | $1.24\times10^{3}$ | $8.37\times10^{4}$ | $5.45\times10^{4}$ | $1.86\times10^{3}$ | $8.58\times10^{2}$ |
| ERP29 (ER) | $7.84\times10^{1}$ | $8.17\times10^{0}$ | 0 | $1.74\times10^{4}$ | 0 | $2.17\times10^{1}$ |
| ERP44 (ER) | $2.89\times10^{4}$ | $3.89\times10^{4}$ | $3.69\times10^{4}$ | $2.25\times10^{4}$ | $5.42\times10^{4}$ | $7.44\times10^{4}$ |
| ETF1 (C) | 0 | 0 | 0 | $9.51\times10^{3}$ | 0 | 0 |
| ETFA (MT) | 0 | $9.27\times10^{2}$ | 0 | $1.54\times10^{4}$ | 0 | $2.64\times10^{3}$ |
| EVPL (CS) | $4.77\times10^{4}$ | $7.50\times10^{4}$ | $8.12\times10^{5}$ | $1.83\times10^{6}$ | $2.82\times10^{4}$ | $3.62\times10^{4}$ |
| EWSR1 (C/NU/PM) | $9.50\times10^{3}$ | $4.14\times10^{4}$ | $3.70\times10^{4}$ | $8.79\times10^{4}$ | $3.33\times10^{4}$ | $2.63\times10^{4}$ |
| EZR (PM/CS) | $6.67\times10^{4}$ | $4.11\times10^{4}$ | $2.58\times10^{5}$ | $5.98\times10^{5}$ | $2.27\times10^{4}$ | $7.86\times10^{4}$ |
| FABP5 (C/NU/EC) | $3.32\times10^{5}$ | $3.27\times10^{5}$ | $4.90\times10^{6}$ | $8.18\times10^{5}$ | $1.43\times10^{5}$ | $1.31\times10^{5}$ |
| FASN (C) | $1.61\times10^{3}$ | $2.35\times10^{3}$ | $1.69\times10^{5}$ | $1.26\times10^{5}$ | $4.20\times10^{3}$ | $1.41\times10^{3}$ |
| FLG (C/PM) | $8.89\times10^{5}$ | $1.42\times10^{6}$ | $1.48\times10^{7}$ | $2.41\times10^{6}$ | $9.03\times10^{5}$ | $4.57\times10^{5}$ |
| FLG2 (C) | $6.52\times10^{6}$ | $5.44\times10^{6}$ | $5.65\times10^{7}$ | $3.06\times10^{6}$ | $6.87\times10^{6}$ | $2.47\times10^{6}$ |
| FLNA (CS) | $9.04\times10^{3}$ | $9.83\times10^{3}$ | $2.45\times10^{5}$ | $3.91\times10^{5}$ | $9.12\times10^{3}$ | $2.74\times10^{4}$ |
| FLNB (CS) | $1.53\times10^{5}$ | $1.03\times10^{5}$ | $8.42\times10^{5}$ | $1.21\times10^{6}$ | $1.30\times10^{5}$ | $8.55\times10^{4}$ |
| FUS (NU) | $4.42\times10^{4}$ | $1.04\times10^{5}$ | $1.21\times10^{5}$ | $2.17\times10^{5}$ | $3.11\times10^{5}$ | $5.81\times10^{4}$ |
| GANAB (GA/ER) | $5.89\times10^{3}$ | $9.88\times10^{3}$ | $6.01\times10^{3}$ | $4.38\times10^{4}$ | $1.02\times10^{4}$ | $3.99\times10^{4}$ |
| GAPDH (C/NU/CS) | $2.06\times10^{6}$ | $1.73\times10^{6}$ | $1.05\times10^{7}$ | $4.52\times10^{6}$ | $2.29\times10^{6}$ | $1.86\times10^{6}$ |
| GARS1 (C/EC) | $4.08\times10^{3}$ | $5.58\times10^{3}$ | $2.51\times10^{4}$ | $4.62\times10^{4}$ | $3.56\times10^{3}$ | $1.05\times10^{4}$ |
| GDI1 (C/GA) | $1.11\times10^{5}$ | $1.20\times10^{5}$ | $7.22\times10^{5}$ | $4.14\times10^{5}$ | $7.77\times10^{4}$ | $1.00\times10^{5}$ |
| GFUS (C/EC) | 0 | $4.61\times10^{2}$ | $1.43\times10^{4}$ | $6.37\times10^{4}$ | $9.60\times10^{1}$ | $1.68\times10^{3}$ |
| GGCT(C/EC) | $4.35\times10^{5}$ | $3.20\times10^{5}$ | $2.19\times10^{6}$ | $3.32\times10^{5}$ | $5.29\times10^{5}$ | $1.40\times10^{5}$ |
| GGH (LY /EC) | $8.21\times10^{4}$ | $1.39\times10^{5}$ | $3.51\times10^{5}$ | $1.38\times10^{5}$ | $1.03\times10^{5}$ | $2.67\times10^{4}$ |
| GLOD4 (MT) | 0 | $2.16\times10^{3}$ | $5.34\times10^{4}$ | $3.64\times10^{4}$ | $3.88\times10^{3}$ | $1.38\times10^{4}$ |
| GLUD1 (ER/MT) | 0 | $3.01\times10^{2}$ | $1.83\times10^{3}$ | $2.43\times10^{4}$ | $7.61\times10^{2}$ | $8.21\times10^{2}$ |
| GLUL(C/MT/PM/ER) | $9.22\times10^{3}$ | $1.89\times10^{4}$ | $9.11\times10^{5}$ | $5.75\times10^{4}$ | $9.47\times10^{3}$ | $1.91\times10^{4}$ |
| GM2A (LY) | $3.49\times10^{4}$ | $2.63\times10^{4}$ | $5.59\times10^{5}$ | $1.75\times10^{5}$ | $1.63\times10^{4}$ | $7.64\times10^{3}$ |
| GOT2 (MT/PM) | $9.28\times10^{3}$ | $1.51\times10^{4}$ | $2.58\times10^{4}$ | $7.67\times10^{4}$ | $3.83\times10^{3}$ | $4.28\times10^{4}$ |
| GPHN (C/PM) | $7.43\times10^{3}$ | $3.78\times10^{3}$ | 0 | $4.33\times10^{3}$ | $5.49\times10^{3}$ | $2.60\times10^{4}$ |
| GPI (C/ER) | $1.42\times10^{3}$ | $6.15\times10^{3}$ | $5.86\times10^{4}$ | $8.74\times10^{4}$ | $3.84\times10^{3}$ | $3.92\times10^{3}$ |
| GSDMA (C/PM) | $2.60\times10^{5}$ | $1.88\times10^{5}$ | $2.23\times10^{6}$ | $4.23\times10^{5}$ | $1.96\times10^{5}$ | $6.86\times10^{4}$ |
| GSN (CS/EC) | $1.34\times10^{5}$ | $8.35\times10^{4}$ | $9.37\times10^{5}$ | $4.48\times10^{6}$ | $7.44\times10^{4}$ | $2.29\times10^{5}$ |
| GSR (C/MT) | $6.82\times10^{3}$ | $3.04\times10^{4}$ | $1.93\times10^{4}$ | $7.11\times10^{4}$ | $1.68\times10^{5}$ | $5.22\times10^{5}$ |
| GSTP1 (C/NU/MT) | $2.34\times10^{4}$ | $6.14\times10^{4}$ | $1.74\times10^{6}$ | $1.44\times10^{6}$ | $4.76\times10^{4}$ | $3.27\times10^{4}$ |

### Table S3
### (page 4)

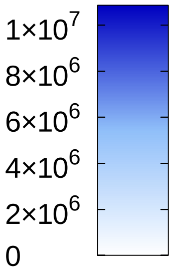


**po-$H_2O_2$ / c-YAP1C**

| | 0 | 2 | 5 | 15 | 30 | 60 |
|---|---|---|---|---|---|---|
| H1-0 (NU) | $1.21\times10^{4}$ | $1.11\times10^{4}$ | $5.58\times10^{4}$ | $2.22\times10^{5}$ | $1.08\times10^{4}$ | $3.94\times10^{3}$ |
| H1-2 (NU) | $1.76\times10^{5}$ | $5.00\times10^{5}$ | $1.97\times10^{5}$ | $1.87\times10^{6}$ | $3.32\times10^{5}$ | $2.25\times10^{5}$ |
| H1-5 (NU) | $9.26\times10^{3}$ | $1.11\times10^{4}$ | $6.64\times10^{4}$ | $3.88\times10^{5}$ | $1.03\times10^{4}$ | $9.51\times10^{2}$ |
| H2AC4 (NU) | $7.37\times10^{2}$ | $4.65\times10^{3}$ | $2.55\times10^{3}$ | $1.36\times10^{5}$ | 0 | $6.30\times10^{2}$ |
| H2BC11 (NU) | $1.05\times10^{4}$ | $2.84\times10^{4}$ | $4.36\times10^{4}$ | $3.96\times10^{5}$ | $5.62\times10^{3}$ | $3.40\times10^{3}$ |
| H3-3A (NU) | $1.63\times10^{5}$ | $3.85\times10^{4}$ | $3.50\times10^{5}$ | $1.44\times10^{6}$ | $2.28\times10^{4}$ | $1.51\times10^{4}$ |
| H4C1 (NU) | $9.29\times10^{5}$ | $3.67\times10^{5}$ | $3.15\times10^{6}$ | $9.81\times10^{6}$ | $1.68\times10^{5}$ | $1.05\times10^{5}$ |
| HADHA (MT) | $8.06\times10^{3}$ | $4.37\times10^{3}$ | $6.37\times10^{4}$ | $1.84\times10^{5}$ | $1.18\times10^{4}$ | $2.04\times10^{3}$ |
| HAL (C) | $1.45\times10^{5}$ | $1.47\times10^{5}$ | $1.24\times10^{6}$ | $3.03\times10^{5}$ | $1.39\times10^{5}$ | $5.33\times10^{4}$ |
| HARS1 (C) | $2.15\times10^{4}$ | $1.09\times10^{4}$ | $1.14\times10^{5}$ | $3.03\times10^{4}$ | $1.11\times10^{4}$ | $6.79\times10^{3}$ |
| HK1 (C/MT) | $6.18\times10^{3}$ | $5.56\times10^{2}$ | $4.56\times10^{3}$ | $7.69\times10^{4}$ | $7.38\times10^{2}$ | $1.09\times10^{4}$ |
| HLA-DRA (ER/LY/ES/PM)) | $2.31\times10^{2}$ | $1.35\times10^{2}$ | $1.20\times10^{3}$ | $4.38\times10^{4}$ | $5.31\times10^{2}$ | 0 |
| HNRNPA1 (NU/C) | $3.39\times10^{4}$ | $7.11\times10^{4}$ | $1.01\times10^{5}$ | $3.53\times10^{5}$ | $7.25\times10^{4}$ | $1.54\times10^{5}$ |
| HNRNPA2B1 (C/NU/EC) | $1.14\times10^{4}$ | $1.99\times10^{4}$ | $6.42\times10^{4}$ | $1.07\times10^{5}$ | $1.27\times10^{4}$ | $1.48\times10^{4}$ |
| HNRNPA3 (NU) | $8.17\times10^{3}$ | $1.08\times10^{4}$ | $7.30\times10^{4}$ | $1.89\times10^{5}$ | $1.31\times10^{4}$ | $9.60\times10^{3}$ |
| HNRNPAB (C/NU) | $3.56\times10^{4}$ | $2.12\times10^{4}$ | $5.07\times10^{4}$ | $9.90\times10^{4}$ | $1.11\times10^{4}$ | $1.86\times10^{4}$ |
| HNRNPC (NU) | $1.24\times10^{4}$ | $4.70\times10^{4}$ | $3.80\times10^{4}$ | $1.90\times10^{5}$ | $9.34\times10^{2}$ | $1.71\times10^{4}$ |
| HNRNPD (C/NU) | $4.09\times10^{3}$ | $3.45\times10^{3}$ | $1.13\times10^{4}$ | $8.81\times10^{4}$ | $4.08\times10^{2}$ | $8.52\times10^{3}$ |
| HNRNPF (NU) | $5.78\times10^{2}$ | 0 | $5.87\times10^{3}$ | $2.53\times10^{4}$ | $5.74\times10^{2}$ | $4.30\times10^{1}$ |
| HNRNPH2 (NU) | $2.31\times10^{2}$ | $2.52\times10^{3}$ | $1.23\times10^{5}$ | $1.29\times10^{5}$ | $4.45\times10^{3}$ | $4.30\times10^{3}$ |
| HNRNPK (C/NU) | $5.86\times10^{3}$ | $1.58\times10^{4}$ | $1.42\times10^{5}$ | $5.61\times10^{5}$ | $3.07\times10^{3}$ | $2.22\times10^{4}$ |
| HNRNPM (NU) | $1.07\times10^{4}$ | $1.51\times10^{4}$ | $1.05\times10^{5}$ | $2.28\times10^{5}$ | $5.26\times10^{3}$ | $6.65\times10^{3}$ |
| HNRNPU (C/NU/CS) | $1.25\times10^{4}$ | $5.03\times10^{4}$ | $6.51\times10^{4}$ | $2.31\times10^{5}$ | $6.97\times10^{3}$ | $1.33\times10^{4}$ |
| HPRT1 (C) | $8.61\times10^{3}$ | $2.35\times10^{4}$ | $1.09\times10^{5}$ | $1.74\times10^{5}$ | $1.27\times10^{5}$ | $7.44\times10^{5}$ |
| HSD17B10 (MT) | 0 | 0 | 0 | $1.53\times10^{4}$ | 0 | $2.00\times10^{3}$ |
| HSD17B4 (PO) | $1.58\times10^{4}$ | $1.07\times10^{4}$ | $4.74\times10^{5}$ | $2.01\times10^{5}$ | $7.75\times10^{3}$ | $1.34\times10^{4}$ |
| HSP90AA1 (C/NU/MT/PM) | $7.71\times10^{4}$ | $1.08\times10^{5}$ | $9.52\times10^{5}$ | $1.22\times10^{6}$ | $1.36\times10^{5}$ | $5.38\times10^{5}$ |
| HSP90AB1 (C/NU/PM/EC) | $1.37\times10^{4}$ | $3.07\times10^{4}$ | $4.02\times10^{5}$ | $9.35\times10^{5}$ | $5.31\times10^{4}$ | $3.12\times10^{5}$ |
| HSP90B1 (ER) | $5.25\times10^{4}$ | $1.02\times10^{5}$ | $2.84\times10^{5}$ | $6.42\times10^{5}$ | $2.07\times10^{5}$ | $3.16\times10^{5}$ |
| HSPA1A (C/NU/EC) | $2.40\times10^{4}$ | $6.00\times10^{4}$ | $1.72\times10^{4}$ | $3.24\times10^{4}$ | $1.48\times10^{4}$ | $3.30\times10^{4}$ |
| HSPA1B (C/CS) | $6.44\times10^{5}$ | $9.67\times10^{5}$ | $2.25\times10^{6}$ | $3.40\times10^{6}$ | $7.97\times10^{5}$ | $1.15\times10^{6}$ |
| HSPA5 (C/ER) | $1.45\times10^{6}$ | $1.67\times10^{6}$ | $4.65\times10^{6}$ | $3.15\times10^{6}$ | $1.40\times10^{6}$ | $1.54\times10^{6}$ |
| HSPA8 (C/PM/NU) | $7.39\times10^{5}$ | $1.48\times10^{6}$ | $2.43\times10^{6}$ | $3.87\times10^{6}$ | $5.00\times10^{5}$ | $4.43\times10^{5}$ |
| HSPA9 (MT) | $1.91\times10^{5}$ | $5.29\times10^{5}$ | $1.80\times10^{5}$ | $3.62\times10^{5}$ | $1.31\times10^{5}$ | $2.61\times10^{5}$ |
| HSPB1 (C/NU/CS) | $3.71\times10^{5}$ | $4.09\times10^{5}$ | $1.48\times10^{7}$ | $9.71\times10^{6}$ | $1.25\times10^{6}$ | $3.84\times10^{5}$ |
| IDE (C/PM/EC) | $1.59\times10^{5}$ | $1.84\times10^{5}$ | $5.91\times10^{6}$ | $8.25\times10^{5}$ | $1.17\times10^{5}$ | $9.05\times10^{4}$ |
| IL1RN (C/EC) | $7.46\times10^{3}$ | $4.08\times10^{3}$ | $1.09\times10^{5}$ | $8.29\times10^{4}$ | $5.61\times10^{3}$ | $2.75\times10^{2}$ |
| IL36G (C/EC) | $4.69\times10^{4}$ | $2.08\times10^{4}$ | $4.42\times10^{5}$ | $1.21\times10^{5}$ | $4.55\times10^{4}$ | $8.63\times10^{3}$ |
| IL36RN (C/EC) | $2.06\times10^{3}$ | $1.82\times10^{3}$ | $9.71\times10^{4}$ | $2.05\times10^{5}$ | $2.89\times10^{2}$ | $3.90\times10^{2}$ |
| IQGAP1 (C/NU/PM) | $1.21\times10^{4}$ | $1.90\times10^{4}$ | $1.95\times10^{5}$ | $5.32\times10^{5}$ | $3.55\times10^{4}$ | $1.65\times10^{4}$ |
| ISYNA1 (C) | $1.58\times10^{6}$ | $5.23\times10^{5}$ | $8.00\times10^{6}$ | $7.33\times10^{5}$ | $2.62\times10^{6}$ | $8.73\times10^{5}$ |
| IVL (C) | $1.13\times10^{5}$ | $8.45\times10^{4}$ | $3.20\times10^{6}$ | $5.53\times10^{6}$ | $1.27\times10^{5}$ | $5.73\times10^{4}$ |
| JUP (CS/PM) | $7.51\times10^{6}$ | $4.91\times10^{6}$ | $3.85\times10^{7}$ | $1.12\times10^{7}$ | $6.07\times10^{6}$ | $2.76\times10^{6}$ |
| LAMP1 (LY) | $4.87\times10^{4}$ | $3.27\times10^{4}$ | $3.54\times10^{5}$ | $1.82\times10^{5}$ | $3.72\times10^{4}$ | $2.25\times10^{4}$ |
| LCN2 (EC/VE) | $8.93\times10^{2}$ | $1.37\times10^{3}$ | $4.45\times10^{5}$ | $7.98\times10^{3}$ | $6.62\times10^{3}$ | $3.31\times10^{2}$ |
| LCP1 (PM/CS) | $5.37\times10^{2}$ | $6.38\times10^{3}$ | $1.41\times10^{4}$ | $8.21\times10^{4}$ | $4.02\times10^{3}$ | $3.12\times10^{4}$ |
| LDHA (C) | $1.30\times10^{5}$ | $1.90\times10^{5}$ | $1.79\times10^{6}$ | $2.60\times10^{6}$ | $1.66\times10^{5}$ | $2.19\times10^{5}$ |
| LDHB (C/MT) | $5.61\times10^{4}$ | $1.45\times10^{5}$ | $7.39\times10^{4}$ | $1.37\times10^{5}$ | $1.87\times10^{5}$ | $1.84\times10^{5}$ |
| LGALS3 (C/NU/EC) | $5.09\times10^{4}$ | $9.35\times10^{4}$ | $5.17\times10^{5}$ | $1.37\times10^{6}$ | $3.49\times10^{4}$ | $7.65\times10^{4}$ |
| LGALS7 (C/NU/EC) | $3.27\times10^{5}$ | $5.47\times10^{5}$ | $3.09\times10^{7}$ | $3.16\times10^{7}$ | $4.54\times10^{5}$ | $3.57\times10^{5}$ |

**Table S3**
**(page 5)**

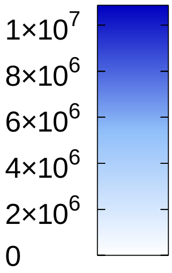


**po-$H_2O_2$ / c-YAP1C**

| 0 | 2 | 5 | 15 | 30 | 60 | |
|---|---|---|---|---|---|---|
| $1.72\times10^{4}$ | $1.03\times10^{4}$ | $1.96\times10^{5}$ | $3.97\times10^{4}$ | $4.13\times10^{3}$ | $2.38\times10^{3}$ | LGALSL (?) |
| $1.28\times10^{5}$ | $1.45\times10^{5}$ | $5.67\times10^{6}$ | $4.84\times10^{6}$ | $1.00\times10^{5}$ | $6.73\times10^{4}$ | LMNA (NU) |
| $2.80\times10^{3}$ | $1.01\times10^{4}$ | $3.16\times10^{4}$ | $3.30\times10^{4}$ | $2.07\times10^{2}$ | $3.09\times10^{3}$ | LMNB1 (NU) |
| $5.58\times10^{2}$ | $3.32\times10^{3}$ | $1.76\times10^{4}$ | $1.66\times10^{4}$ | 0 | 0 | LMNB2 (NU) |
| $2.41\times10^{2}$ | $6.19\times10^{2}$ | $8.14\times10^{4}$ | $4.77\times10^{4}$ | $1.08\times10^{3}$ | $1.35\times10^{4}$ | LTA4H (C) |
| $5.01\times10^{3}$ | $3.88\times10^{3}$ | $1.64\times10^{5}$ | $1.73\times10^{5}$ | $3.30\times10^{3}$ | $3.77\times10^{3}$ | LY6D (PM) |
| $4.14\times10^{2}$ | $5.26\times10^{3}$ | $3.94\times10^{3}$ | $2.17\times10^{5}$ | $1.94\times10^{3}$ | $2.24\times10^{3}$ | MACROH2A1 (NU) |
| $1.82\times10^{3}$ | $4.57\times10^{3}$ | $4.98\times10^{3}$ | $6.06\times10^{4}$ | $3.90\times10^{3}$ | $8.33\times10^{3}$ | MAP4 (CS) |
| $2.17\times10^{4}$ | $2.31\times10^{4}$ | $2.82\times10^{5}$ | $2.11\times10^{5}$ | $8.61\times10^{3}$ | $1.95\times10^{4}$ | MDH1 (C) |
| $5.11\times10^{4}$ | $9.72\times10^{4}$ | $3.69\times10^{5}$ | $5.95\times10^{5}$ | $9.92\times10^{4}$ | $7.51\times10^{4}$ | MDH2 (MT) |
| $3.28\times10^{3}$ | $7.57\times10^{3}$ | $8.56\times10^{4}$ | $1.90\times10^{5}$ | $6.24\times10^{3}$ | $2.03\times10^{4}$ | MSN (CS/PM) |
| $1.02\times10^{3}$ | $2.76\times10^{3}$ | $1.96\times10^{4}$ | $3.74\times10^{3}$ | $4.60\times10^{3}$ | $2.69\times10^{3}$ | MTAP (C/NU) |
| 0 | $1.21\times10^{0}$ | 0 | $1.14\times10^{4}$ | $1.58\times10^{3}$ | $1.51\times10^{4}$ | MTHFD1 (C) |
| $2.61\times10^{3}$ | $6.61\times10^{3}$ | $2.42\times10^{2}$ | $6.13\times10^{4}$ | $3.22\times10^{2}$ | $1.67\times10^{4}$ | MTPN (C/NU) |
| $1.34\times10^{3}$ | $2.24\times10^{3}$ | $2.63\times10^{5}$ | $2.29\times10^{5}$ | $2.79\times10^{3}$ | $5.67\times10^{3}$ | MYH14 (C/CS/EC) |
| $1.06\times10^{5}$ | $9.86\times10^{4}$ | $2.30\times10^{6}$ | $2.76\times10^{6}$ | $1.36\times10^{5}$ | $1.12\times10^{5}$ | MYH9 (CS) |
| $1.28\times10^{3}$ | $4.88\times10^{2}$ | $3.60\times10^{4}$ | $1.39\times10^{5}$ | $6.43\times10^{1}$ | $6.28\times10^{3}$ | MYL12A (C/CS/EC) |
| $1.95\times10^{4}$ | $3.23\times10^{4}$ | $2.25\times10^{5}$ | $6.62\times10^{5}$ | $1.14\times10^{4}$ | $3.07\times10^{4}$ | MYL6 (C/CS/EC) |
| $2.60\times10^{4}$ | $1.64\times10^{4}$ | $2.73\times10^{5}$ | $1.78\times10^{5}$ | $2.98\times10^{4}$ | $3.12\times10^{4}$ | NAGK (C/EC) |
| $1.63\times10^{3}$ | $6.64\times10^{2}$ | $2.53\times10^{5}$ | $1.24\times10^{5}$ | $1.33\times10^{3}$ | $7.68\times10^{2}$ | NAMPT (C/NU/EC) |
| $7.12\times10^{0}$ | $1.34\times10^{2}$ | $3.63\times10^{4}$ | $3.08\times10^{4}$ | $1.16\times10^{2}$ | $6.00\times10^{2}$ | NAPA (PM) |
| $1.13\times10^{5}$ | $7.36\times10^{4}$ | $1.96\times10^{6}$ | $4.40\times10^{5}$ | $4.67\times10^{5}$ | $1.53\times10^{5}$ | NCCRP1 (C/NU/MT) |
| $5.53\times10^{3}$ | $1.30\times10^{4}$ | $5.60\times10^{3}$ | $1.26\times10^{5}$ | $5.44\times10^{3}$ | $1.44\times10^{4}$ | NCL (C/NU) |
| 0 | 0 | $1.59\times10^{3}$ | $4.25\times10^{4}$ | 0 | 0 | NDRG1 (C/NU/PM/CS) |
| $3.96\times10^{2}$ | 0 | $9.41\times10^{3}$ | $2.49\times10^{4}$ | 0 | $3.79\times10^{2}$ | NDRG2 (C) |
| $1.99\times10^{4}$ | $1.22\times10^{4}$ | $2.02\times10^{5}$ | $1.43\times10^{5}$ | $1.39\times10^{4}$ | $4.14\times10^{3}$ | NEU2 (C) |
| $4.18\times10^{3}$ | $2.93\times10^{2}$ | $5.47\times10^{3}$ | $2.57\times10^{4}$ | $8.44\times10^{2}$ | $1.20\times10^{4}$ | NONO (NU) |
| $4.46\times10^{4}$ | $4.14\times10^{4}$ | $5.18\times10^{5}$ | $1.97\times10^{5}$ | $8.01\times10^{4}$ | $1.85\times10^{5}$ | NPEPPS (C/NU) |
| $6.30\times10^{4}$ | $7.80\times10^{4}$ | $4.74\times10^{4}$ | $1.46\times10^{5}$ | $7.26\times10^{4}$ | $8.69\times10^{4}$ | NPM1 (NU) |
| $1.95\times10^{0}$ | 0 | $7.67\times10^{4}$ | $9.17\times10^{3}$ | 0 | $2.19\times10^{2}$ | NT5C3A (C/ER) |
| $4.63\times10^{3}$ | $9.34\times10^{3}$ | $6.86\times10^{3}$ | $1.76\times10^{4}$ | $1.54\times10^{4}$ | $1.68\times10^{5}$ | NUDC (CS/NU) |
| $1.72\times10^{3}$ | $7.88\times10^{2}$ | $7.43\times10^{3}$ | $4.63\times10^{4}$ | 0 | $1.29\times10^{2}$ | OTUB1 (C) |
| $1.00\times10^{5}$ | $1.65\times10^{5}$ | $7.58\times10^{5}$ | $1.06\times10^{6}$ | $2.20\times10^{5}$ | $2.02\times10^{5}$ | P4HB (PM/ER) |
| $2.05\times10^{4}$ | $2.04\times10^{4}$ | $2.80\times10^{5}$ | $2.55\times10^{5}$ | $2.29\times10^{4}$ | $2.21\times10^{4}$ | PABPC1 (C/NU) |
| $3.96\times10^{3}$ | $5.30\times10^{2}$ | $6.25\times10^{4}$ | $2.78\times10^{4}$ | $1.71\times10^{3}$ | $4.04\times10^{2}$ | PAFAH1B1 (NU/CS/PM) |
| $9.87\times10^{2}$ | $7.78\times10^{3}$ | $3.75\times10^{4}$ | $3.18\times10^{4}$ | $1.80\times10^{4}$ | $4.11\times10^{4}$ | PAICS (C/EC) |
| $4.34\times10^{3}$ | $1.05\times10^{3}$ | $1.09\times10^{4}$ | $3.65\times10^{4}$ | $1.25\times10^{3}$ | $9.52\times10^{4}$ | PC (MT) |
| $5.92\times10^{2}$ | $6.74\times10^{2}$ | $1.36\times10^{4}$ | $5.98\times10^{4}$ | $6.48\times10^{2}$ | $3.04\times10^{3}$ | PCBP1 (C/NU) |
| $1.80\times10^{3}$ | $7.61\times10^{2}$ | $1.94\times10^{3}$ | $2.36\times10^{4}$ | $3.84\times10^{2}$ | $3.36\times10^{3}$ | PCBP2 (C/NU) |
| $4.05\times10^{5}$ | $7.97\times10^{5}$ | $1.63\times10^{5}$ | $3.42\times10^{5}$ | $5.00\times10^{5}$ | $1.49\times10^{6}$ | PCCA (MT) |
| $1.35\times10^{6}$ | $2.24\times10^{6}$ | $4.60\times10^{5}$ | $6.26\times10^{5}$ | $1.27\times10^{6}$ | $3.07\times10^{6}$ | PCCB (MT) |
| $1.65\times10^{3}$ | $6.23\times10^{2}$ | $5.58\times10^{4}$ | $1.16\times10^{4}$ | $3.03\times10^{3}$ | $9.91\times10^{1}$ | PCYOX1 (LY) |
| $1.21\times10^{3}$ | $1.25\times10^{3}$ | $9.02\times10^{4}$ | $1.74\times10^{5}$ | $2.89\times10^{3}$ | $6.85\times10^{2}$ | PDCD6IP (C/CS) |
| $4.34\times10^{4}$ | $6.60\times10^{4}$ | $3.17\times10^{5}$ | $7.03\times10^{5}$ | $7.02\times10^{4}$ | $1.04\times10^{5}$ | PDIA3 (ER) |
| $1.85\times10^{3}$ | $7.93\times10^{3}$ | $4.77\times10^{4}$ | $5.94\times10^{4}$ | $3.44\times10^{3}$ | $1.51\times10^{4}$ | PDIA4 (ER) |
| $1.23\times10^{5}$ | $7.05\times10^{4}$ | $5.97\times10^{5}$ | $2.87\times10^{5}$ | $6.50\times10^{4}$ | $3.65\times10^{4}$ | PEBP1 (C) |
| $6.77\times10^{4}$ | $6.90\times10^{4}$ | $3.56\times10^{5}$ | $1.01\times10^{6}$ | $3.65\times10^{4}$ | $2.20\times10^{5}$ | PFN1 (CS) |
| $8.40\times10^{4}$ | $6.58\times10^{4}$ | $1.53\times10^{6}$ | $1.24\times10^{6}$ | $3.87\times10^{4}$ | $1.84\times10^{5}$ | PGK1 (C) |
| $4.36\times10^{3}$ | $7.47\times10^{3}$ | $3.02\times10^{4}$ | $1.47\times10^{5}$ | $2.58\times10^{3}$ | $3.31\times10^{3}$ | PHB2 (C/NU/MT/PM) |
| $3.21\times10^{4}$ | $1.19\times10^{4}$ | $1.60\times10^{5}$ | $6.10\times10^{5}$ | $9.84\times10^{3}$ | $1.62\times10^{4}$ | PHGDH (C/EC) |

### Table S3
### (page 6)

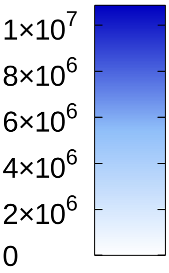


**po-$H_2O_2$ / c-YAP1C**

| 0 | 2 | 5 | 15 | 30 | 60 | |
|---|---|---|---|---|---|---|
| $1.56\times10^{5}$ | $2.22\times10^{5}$ | $3.92\times10^{6}$ | $3.28\times10^{6}$ | $2.83\times10^{5}$ | $2.85\times10^{5}$ | PKM (C/NU) |
| $1.10\times10^{6}$ | $3.59\times10^{5}$ | $4.74\times10^{6}$ | $4.58\times10^{6}$ | $1.17\times10^{6}$ | $2.76\times10^{5}$ | PKP1 (NU) |
| $4.59\times10^{3}$ | $4.45\times10^{3}$ | $9.72\times10^{4}$ | $6.33\times10^{5}$ | $7.63\times10^{3}$ | $1.45\times10^{3}$ | PKP3 (NU) |
| $9.89\times10^{2}$ | $4.65\times10^{2}$ | $2.89\times10^{4}$ | $7.12\times10^{3}$ | $8.26\times10^{2}$ | $2.34\times10^{3}$ | PLA2G4E (C/PM/LY/ES) |
| $9.03\times10^{4}$ | $7.12\times10^{4}$ | $9.28\times10^{5}$ | $1.81\times10^{6}$ | $4.25\times10^{4}$ | $5.92\times10^{4}$ | PLEC (CS) |
| $5.24\times10^{3}$ | $9.22\times10^{3}$ | $4.32\times10^{4}$ | $2.52\times10^{5}$ | $1.60\times10^{4}$ | $8.07\times10^{2}$ | PLP2 (ER/PM) |
| $1.33\times10^{4}$ | $1.71\times10^{4}$ | $5.15\times10^{5}$ | $5.38\times10^{5}$ | $5.05\times10^{4}$ | $2.67\times10^{5}$ | PLS3 (C) |
| $7.63\times10^{4}$ | $2.88\times10^{4}$ | $3.87\times10^{5}$ | $1.20\times10^{5}$ | $8.95\times10^{4}$ | $3.99\times10^{4}$ | PNP (C) |
| $4.02\times10^{5}$ | $3.17\times10^{5}$ | $6.82\times10^{6}$ | $1.75\times10^{6}$ | $2.74\times10^{5}$ | $1.31\times10^{5}$ | POF1B (C/CS) |
| $6.42\times10^{3}$ | $8.59\times10^{3}$ | $1.07\times10^{5}$ | $7.89\times10^{4}$ | $1.79\times10^{4}$ | $2.25\times10^{4}$ | PPA1 (C) |
| $1.81\times10^{5}$ | $4.05\times10^{5}$ | $7.80\times10^{5}$ | $2.70\times10^{6}$ | $4.77\times10^{5}$ | $9.50\times10^{5}$ | PPIA (C/NU/EC) |
| $5.94\times10^{3}$ | $7.19\times10^{3}$ | $1.38\times10^{5}$ | $1.99\times10^{5}$ | $3.98\times10^{3}$ | $4.23\times10^{3}$ | PPIB (ER) |
| $9.38\times10^{4}$ | $5.38\times10^{4}$ | $1.83\times10^{6}$ | $2.31\times10^{6}$ | $5.77\times10^{4}$ | $2.58\times10^{4}$ | PPL (C/CS/PM) |
| $7.86\times10^{4}$ | $2.17\times10^{4}$ | $6.24\times10^{4}$ | $1.85\times10^{5}$ | $1.02\times10^{5}$ | $5.73\times10^{4}$ | PPP2R1A (C/NU/PM) |
| $7.75\times10^{5}$ | $3.61\times10^{6}$ | $6.15\times10^{6}$ | $8.89\times10^{6}$ | $1.43\times10^{7}$ | $3.12\times10^{7}$ | PRDX1 (C) |
| $4.74\times10^{5}$ | $1.50\times10^{6}$ | $3.52\times10^{6}$ | $2.86\times10^{6}$ | $4.64\times10^{6}$ | $6.72\times10^{6}$ | PRDX2 (C) |
| $2.94\times10^{5}$ | $6.00\times10^{5}$ | $3.81\times10^{5}$ | $9.73\times10^{5}$ | $1.16\times10^{6}$ | $2.89\times10^{6}$ | PRDX3 (MT/C/EE) |
| $1.38\times10^{4}$ | $6.61\times10^{4}$ | $5.82\times10^{4}$ | $5.65\times10^{4}$ | $8.56\times10^{4}$ | $1.30\times10^{5}$ | PRDX4 (C/ER) |
| $5.08\times10^{4}$ | $4.98\times10^{4}$ | $5.23\times10^{5}$ | $8.20\times10^{5}$ | $7.72\times10^{5}$ | $2.40\times10^{6}$ | PRDX6 (C/LY) |
| $2.53\times10^{2}$ | $2.96\times10^{2}$ | $4.99\times10^{4}$ | $1.13\times10^{4}$ | 0 | $9.96\times10^{2}$ | PREP (C) |
| $2.44\times10^{3}$ | $5.08\times10^{3}$ | $1.14\times10^{4}$ | $3.37\times10^{4}$ | $3.14\times10^{5}$ | $1.17\times10^{4}$ | PRKD2 (C/NU/GA/PM) |
| $4.26\times10^{3}$ | $9.11\times10^{3}$ | $1.88\times10^{4}$ | $3.49\times10^{4}$ | $3.17\times10^{4}$ | $1.82\times10^{5}$ | PRMT5 (C/NU/GA) |
| $4.44\times10^{2}$ | $9.81\times10^{2}$ | $1.34\times10^{4}$ | $1.45\times10^{4}$ | $1.93\times10^{3}$ | $6.17\times10^{2}$ | PRXL2A (C/EC) |
| $4.28\times10^{3}$ | $1.11\times10^{4}$ | $1.27\times10^{5}$ | $2.29\times10^{4}$ | $2.80\times10^{4}$ | $8.95\times10^{3}$ | PSMA2 (C/NU) |
| $2.14\times10^{4}$ | $1.80\times10^{4}$ | $2.17\times10^{5}$ | $7.30\times10^{5}$ | $3.19\times10^{4}$ | $9.73\times10^{3}$ | PSMA3 (C/NU) |
| $6.66\times10^{4}$ | $6.84\times10^{4}$ | $6.07\times10^{5}$ | $2.28\times10^{5}$ | $2.62\times10^{5}$ | $3.38\times10^{4}$ | PSMA5 (C/NU) |
| $3.72\times10^{4}$ | $2.52\times10^{4}$ | $2.48\times10^{5}$ | $8.77\times10^{4}$ | $3.76\times10^{4}$ | $9.28\times10^{3}$ | PSMA6 (C/NU) |
| $7.14\times10^{4}$ | $3.79\times10^{4}$ | $4.55\times10^{5}$ | $1.06\times10^{5}$ | $8.19\times10^{4}$ | $2.29\times10^{4}$ | PSMA7 (C/NU) |
| $1.92\times10^{4}$ | $1.63\times10^{4}$ | $3.25\times10^{5}$ | $8.74\times10^{4}$ | $3.90\times10^{4}$ | $1.04\times10^{4}$ | PSMB4 (C/NU) |
| $2.95\times10^{4}$ | $1.96\times10^{4}$ | $7.74\times10^{4}$ | $3.00\times10^{4}$ | $1.48\times10^{4}$ | $7.07\times10^{3}$ | PSMB5 (C/NU) |
| $2.05\times10^{4}$ | $2.37\times10^{4}$ | $1.71\times10^{5}$ | $9.27\times10^{4}$ | $2.76\times10^{4}$ | $1.32\times10^{4}$ | PSMB6 (C/NU) |
| $3.57\times10^{3}$ | $8.09\times10^{2}$ | $3.15\times10^{4}$ | $1.20\times10^{4}$ | $2.21\times10^{3}$ | $1.78\times10^{3}$ | PSMB8 (C/NU) |
| 0 | $3.28\times10^{2}$ | $1.34\times10^{4}$ | $3.21\times10^{4}$ | 0 | $6.61\times10^{2}$ | PSMC3 (C/NU) |
| 0 | $1.36\times10^{3}$ | $5.67\times10^{3}$ | $2.39\times10^{4}$ | 0 | $2.38\times10^{3}$ | PSMC5 (C/NU) |
| 0 | $1.22\times10^{2}$ | $2.30\times10^{4}$ | $3.60\times10^{4}$ | 0 | $1.40\times10^{4}$ | PSMD14 (C/NU/EC) |
| $4.71\times10^{3}$ | $1.34\times10^{4}$ | $1.11\times10^{5}$ | $1.01\times10^{5}$ | $1.89\times10^{4}$ | $1.48\times10^{4}$ | PTBP1 (NU) |
| $2.65\times10^{3}$ | $6.13\times10^{3}$ | $7.46\times10^{2}$ | $1.99\times10^{4}$ | $3.63\times10^{3}$ | $3.35\times10^{2}$ | PTCD1 (MT) |
| $5.85\times10^{3}$ | $4.09\times10^{3}$ | $2.80\times10^{3}$ | $3.86\times10^{4}$ | $2.98\times10^{3}$ | $2.93\times10^{3}$ | PTMA (NU) |
| $2.59\times10^{3}$ | $3.91\times10^{2}$ | $5.22\times10^{2}$ | $4.06\times10^{4}$ | $2.90\times10^{3}$ | $3.32\times10^{2}$ | PYCARD (C/NU/MT/ER) |
| $1.31\times10^{4}$ | $1.48\times10^{4}$ | $1.07\times10^{5}$ | $1.13\times10^{5}$ | $1.33\times10^{4}$ | $7.44\times10^{3}$ | RAB10 (N/CS/ER/GA/ES |
| $3.84\times10^{3}$ | $6.40\times10^{3}$ | $1.42\times10^{5}$ | $2.77\times10^{5}$ | $6.62\times10^{3}$ | $2.14\times10^{3}$ | RAB11A (PM/ES) |
| $1.38\times10^{4}$ | $1.19\times10^{4}$ | $1.34\times10^{5}$ | $1.74\times10^{5}$ | $4.70\times10^{3}$ | $4.97\times10^{3}$ | RAB14 (GA/ES) |
| $6.81\times10^{3}$ | $3.64\times10^{4}$ | $1.09\times10^{5}$ | $1.19\times10^{5}$ | $1.90\times10^{4}$ | $1.55\times10^{4}$ | RAB1A (C/ES/GA/PM) |
| $5.29\times10^{3}$ | $3.33\times10^{3}$ | $6.56\times10^{4}$ | $5.03\times10^{4}$ | $1.14\times10^{3}$ | 0 | RAB2A (GA/ER) |
| $4.27\times10^{4}$ | $2.30\times10^{4}$ | $2.77\times10^{5}$ | $2.10\times10^{5}$ | $2.40\times10^{4}$ | $1.85\times10^{4}$ | RAB7A (LY/ES) |
| $5.72\times10^{4}$ | $1.06\times10^{5}$ | $3.62\times10^{5}$ | $4.06\times10^{5}$ | $6.09\times10^{4}$ | $1.03\times10^{5}$ | RACK1 (C/NU/PM) |
| $7.01\times10^{2}$ | $8.89\times10^{2}$ | $4.70\times10^{4}$ | $4.40\times10^{4}$ | $9.44\times10^{1}$ | $1.16\times10^{3}$ | RAD23B (C/NU) |
| $1.15\times10^{5}$ | $2.11\times10^{5}$ | $3.55\times10^{5}$ | $3.59\times10^{5}$ | $1.32\times10^{5}$ | $2.20\times10^{5}$ | RAN (C/NU) |
| $3.71\times10^{2}$ | $6.52\times10^{3}$ | $2.44\times10^{4}$ | $3.78\times10^{4}$ | $2.34\times10^{2}$ | $6.20\times10^{2}$ | RARS1 (C) |
| $5.00\times10^{3}$ | $1.72\times10^{3}$ | $2.14\times10^{5}$ | $2.94\times10^{5}$ | $3.27\times10^{3}$ | $3.58\times10^{3}$ | RNH1 (C) |

**Table S3**
**(page 7)**

**po-$H_2O_2$ / c-YAP1C**

raw abundance

$1\times10^{7}$
$8\times10^{6}$
$6\times10^{6}$
$4\times10^{6}$
$2\times10^{6}$
0

| | 0 | 2 | 5 | 15 | 30 | 60 |
|---|---|---|---|---|---|---|
| RO60 (C) | $5.98\times10^{3}$ | $3.89\times10^{3}$ | $2.00\times10^{4}$ | $1.44\times10^{4}$ | $4.00\times10^{3}$ | $3.02\times10^{3}$ |
| RPL10 (C/NU/ER) | $1.77\times10^{3}$ | $3.42\times10^{3}$ | $7.26\times10^{3}$ | $7.22\times10^{4}$ | $5.45\times10^{3}$ | $9.61\times10^{3}$ |
| RPL11 (C/NU) | $4.43\times10^{4}$ | $1.05\times10^{5}$ | $1.37\times10^{5}$ | $3.86\times10^{5}$ | $2.53\times10^{4}$ | $7.35\times10^{4}$ |
| RPL12 (C/EC) | $6.79\times10^{3}$ | $9.70\times10^{3}$ | $4.78\times10^{4}$ | $1.79\times10^{5}$ | $4.42\times10^{3}$ | $5.27\times10^{3}$ |
| RPL13 (C) | $6.23\times10^{3}$ | $2.96\times10^{4}$ | $1.09\times10^{4}$ | $2.13\times10^{5}$ | $1.41\times10^{4}$ | $2.94\times10^{4}$ |
| RPL13A (C) | $4.76\times10^{3}$ | $1.01\times10^{4}$ | $2.44\times10^{4}$ | $1.75\times10^{5}$ | $1.03\times10^{4}$ | $2.26\times10^{4}$ |
| RPL14 (C/EC) | $3.53\times10^{3}$ | $9.33\times10^{3}$ | $1.02\times10^{4}$ | $1.91\times10^{5}$ | $9.00\times10^{3}$ | $1.46\times10^{4}$ |
| RPL15 (PM) | $1.03\times10^{4}$ | $3.58\times10^{4}$ | $4.58\times10^{4}$ | $2.44\times10^{5}$ | $4.69\times10^{4}$ | $2.72\times10^{4}$ |
| RPL17-C18orf32 (C) | $1.84\times10^{3}$ | $1.30\times10^{4}$ | $2.92\times10^{3}$ | $9.17\times10^{4}$ | $8.22\times10^{3}$ | $1.54\times10^{4}$ |
| RPL18A (C/PM) | $1.09\times10^{4}$ | $2.37\times10^{4}$ | $7.64\times10^{4}$ | $4.85\times10^{5}$ | $6.12\times10^{4}$ | $5.47\times10^{4}$ |
| RPL19 (CC/NU) | $3.08\times10^{4}$ | $3.18\times10^{4}$ | $2.71\times10^{4}$ | $1.11\times10^{5}$ | $2.82\times10^{4}$ | $2.70\times10^{4}$ |
| RPL21 (C/ER) | $1.59\times10^{3}$ | $5.63\times10^{3}$ | $1.04\times10^{2}$ | $1.35\times10^{5}$ | $5.42\times10^{3}$ | $7.40\times10^{3}$ |
| RPL22 (C/NU/EC) | $1.51\times10^{4}$ | $6.67\times10^{4}$ | $2.22\times10^{5}$ | $2.72\times10^{5}$ | $7.93\times10^{3}$ | $4.63\times10^{4}$ |
| RPL23 (C/NU/EC) | $2.80\times10^{4}$ | $3.77\times10^{4}$ | $4.56\times10^{4}$ | $2.27\times10^{5}$ | $1.50\times10^{4}$ | $2.02\times10^{4}$ |
| RPL23A (C/NU/EC) | $1.54\times10^{5}$ | $1.94\times10^{5}$ | $2.48\times10^{5}$ | $6.03\times10^{5}$ | $1.93\times10^{5}$ | $1.73\times10^{5}$ |
| RPL27A (C/ER) | $4.85\times10^{3}$ | $1.55\times10^{4}$ | $3.16\times10^{3}$ | $1.73\times10^{5}$ | $1.06\times10^{4}$ | $1.48\times10^{4}$ |
| RPL28 (C/EC) | $2.89\times10^{3}$ | $1.09\times10^{4}$ | $3.96\times10^{3}$ | $1.29\times10^{5}$ | $3.04\times10^{3}$ | $1.37\times10^{4}$ |
| RPL29 (C) | 0 | $2.60\times10^{4}$ | $7.00\times10^{3}$ | $1.68\times10^{5}$ | $4.40\times10^{4}$ | $2.72\times10^{4}$ |
| RPL3 (C/NU) | $3.91\times10^{4}$ | $1.95\times10^{4}$ | $2.61\times10^{4}$ | $2.09\times10^{5}$ | $1.56\times10^{4}$ | $2.02\times10^{4}$ |
| RPL36A (C) | $1.20\times10^{2}$ | $5.97\times10^{2}$ | 0 | $5.77\times10^{3}$ | $3.94\times10^{1}$ | $7.01\times10^{2}$ |
| RPL4 (C/NU/ER/EC) | $2.35\times10^{4}$ | $3.16\times10^{4}$ | $3.92\times10^{4}$ | $4.00\times10^{5}$ | $3.17\times10^{4}$ | $4.42\times10^{4}$ |
| RPL5 (C/NU) | $8.38\times10^{3}$ | $1.04\times10^{4}$ | $5.79\times10^{3}$ | $1.44\times10^{5}$ | $5.25\times10^{3}$ | $1.69\times10^{4}$ |
| RPL6 (C/ER) | $3.94\times10^{4}$ | $6.24\times10^{4}$ | $2.19\times10^{5}$ | $7.23\times10^{5}$ | $8.42\times10^{4}$ | $8.29\times10^{4}$ |
| RPL7 (C/NU) | $2.16\times10^{4}$ | $3.73\times10^{4}$ | $4.50\times10^{4}$ | $3.93\times10^{5}$ | $3.00\times10^{4}$ | $6.33\times10^{4}$ |
| RPL7A (C/NU) | $2.18\times10^{4}$ | $9.65\times10^{4}$ | $1.29\times10^{5}$ | $7.59\times10^{5}$ | $1.58\times10^{5}$ | $9.09\times10^{4}$ |
| RPL8 (C) | $2.67\times10^{3}$ | $1.21\times10^{4}$ | $1.82\times10^{4}$ | $1.38\times10^{5}$ | $8.94\times10^{3}$ | $1.56\times10^{4}$ |
| RPLP0 (C/NU) | $9.01\times10^{2}$ | $1.68\times10^{4}$ | $4.95\times10^{4}$ | $5.17\times10^{5}$ | $4.22\times10^{4}$ | $3.22\times10^{4}$ |
| RPLP2 (C/EC) | $6.47\times10^{4}$ | $7.86\times10^{4}$ | $7.04\times10^{5}$ | $1.63\times10^{6}$ | $2.88\times10^{4}$ | $6.93\times10^{4}$ |
| RPN1 (ER) | $2.13\times10^{4}$ | $3.52\times10^{4}$ | $1.98\times10^{5}$ | $1.33\times10^{5}$ | $4.55\times10^{4}$ | $5.58\times10^{4}$ |
| RPN2 (ER) | $3.68\times10^{3}$ | $1.48\times10^{4}$ | $4.49\times10^{4}$ | $2.56\times10^{4}$ | $5.52\times10^{3}$ | $4.63\times10^{3}$ |
| RPS10 (C/NU) | $2.38\times10^{4}$ | $1.55\times10^{5}$ | $3.41\times10^{4}$ | $1.82\times10^{5}$ | $2.49\times10^{4}$ | $4.96\times10^{4}$ |
| RPS13 (C/NU/EC) | $1.26\times10^{4}$ | $2.63\times10^{4}$ | $4.48\times10^{4}$ | $2.21\times10^{5}$ | $1.47\times10^{4}$ | $2.88\times10^{4}$ |
| RPS14 (C/NU/EC) | $1.37\times10^{4}$ | $2.22\times10^{4}$ | $6.38\times10^{3}$ | $1.22\times10^{5}$ | $1.16\times10^{4}$ | $2.04\times10^{4}$ |
| RPS15A (C/NU/EC) | $4.94\times10^{4}$ | $4.71\times10^{4}$ | $1.65\times10^{5}$ | $2.58\times10^{5}$ | $3.91\times10^{4}$ | $2.96\times10^{4}$ |
| RPS16 (C/NU/EC) | $2.30\times10^{4}$ | $3.07\times10^{4}$ | $1.58\times10^{4}$ | $1.40\times10^{5}$ | $1.37\times10^{4}$ | $1.92\times10^{4}$ |
| RPS18 (C) | $3.87\times10^{4}$ | $8.35\times10^{4}$ | $3.16\times10^{4}$ | $2.11\times10^{5}$ | $2.08\times10^{4}$ | $4.42\times10^{4}$ |
| RPS19 (NU) | $4.41\times10^{4}$ | $1.75\times10^{5}$ | $2.03\times10^{4}$ | $3.26\times10^{5}$ | $5.75\times10^{4}$ | $9.02\times10^{4}$ |
| RPS21 (C/ER) | $5.27\times10^{3}$ | $5.59\times10^{3}$ | $5.53\times10^{4}$ | $1.01\times10^{5}$ | $1.57\times10^{4}$ | $1.10\times10^{4}$ |
| RPS23 (C/ER) | $3.87\times10^{3}$ | $1.23\times10^{4}$ | $2.40\times10^{3}$ | $9.20\times10^{4}$ | $2.24\times10^{3}$ | $1.05\times10^{4}$ |
| RPS25 (C/NU/EC) | $5.08\times10^{4}$ | $1.54\times10^{5}$ | $1.23\times10^{5}$ | $4.45\times10^{5}$ | $7.19\times10^{4}$ | $1.10\times10^{5}$ |
| RPS26 (C/ER) | $2.89\times10^{4}$ | $1.78\times10^{4}$ | $1.76\times10^{4}$ | $1.95\times10^{5}$ | $1.87\times10^{4}$ | $3.45\times10^{4}$ |
| RPS27A (C/NU) | $1.84\times10^{6}$ | $1.82\times10^{6}$ | $1.04\times10^{7}$ | $5.58\times10^{6}$ | $2.37\times10^{6}$ | $1.43\times10^{6}$ |
| RPS28 (C/ER) | $4.20\times10^{3}$ | $3.77\times10^{3}$ | $5.11\times10^{3}$ | $5.01\times10^{4}$ | $1.72\times10^{3}$ | $3.48\times10^{3}$ |
| RPS3 (C/NU/MT/CS) | $1.15\times10^{5}$ | $2.46\times10^{5}$ | $5.94\times10^{5}$ | $1.10\times10^{6}$ | $1.03\times10^{5}$ | $2.38\times10^{5}$ |
| RPS3A (C/NU) | $1.65\times10^{4}$ | $2.33\times10^{4}$ | $3.60\times10^{4}$ | $2.79\times10^{5}$ | $9.27\times10^{3}$ | $2.60\times10^{4}$ |
| RPS4X (C) | $1.48\times10^{4}$ | $3.56\times10^{4}$ | $1.69\times10^{4}$ | $3.39\times10^{5}$ | $1.77\times10^{4}$ | $4.07\times10^{4}$ |
| RPS5 (C/NU/EC) | $1.33\times10^{4}$ | $5.50\times10^{4}$ | $2.75\times10^{4}$ | $7.31\times10^{4}$ | $9.80\times10^{3}$ | $1.41\times10^{4}$ |
| RPS6 (C/NU/ER) | $9.45\times10^{3}$ | $2.20\times10^{4}$ | $1.63\times10^{4}$ | $1.86\times10^{5}$ | $1.17\times10^{4}$ | $2.43\times10^{4}$ |
| RPS7 (C) | $1.18\times10^{4}$ | $3.45\times10^{4}$ | $6.16\times10^{4}$ | $2.23\times10^{5}$ | $1.32\times10^{4}$ | $3.74\times10^{4}$ |
| RPS8 (C/PM) | $3.60\times10^{4}$ | $1.35\times10^{5}$ | $3.88\times10^{5}$ | $7.36\times10^{5}$ | $7.67\times10^{4}$ | $1.35\times10^{5}$ |

### Table S3
### (page 8)

po-$H_2O_2$ / c-YAP1C

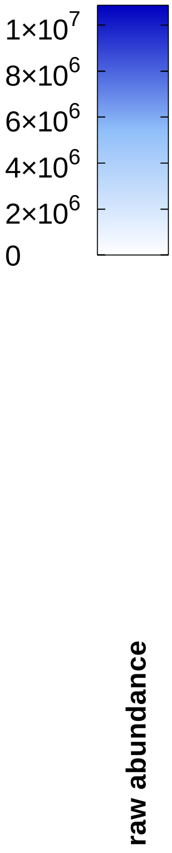


| | 0 | 2 | 5 | 15 | 30 | 60 |
|---|---|---|---|---|---|---|
| RPSA (NU/C/PM) | $6.20\times10^{3}$ | $1.93\times10^{4}$ | $4.37\times10^{5}$ | $6.73\times10^{5}$ | $4.07\times10^{4}$ | $9.49\times10^{4}$ |
| RTCB (C/NU) | $6.96\times10^{3}$ | $5.64\times10^{4}$ | $5.73\times10^{4}$ | $4.95\times10^{4}$ | $1.42\times10^{3}$ | $2.97\times10^{3}$ |
| RTN4 (PM/ER) | $4.43\times10^{2}$ | $2.26\times10^{2}$ | $3.03\times10^{4}$ | $9.60\times10^{4}$ | $5.38\times10^{2}$ | $2.73\times10^{1}$ |
| S100A14 (C) | $2.38\times10^{5}$ | $1.25\times10^{5}$ | $1.13\times10^{6}$ | $5.10\times10^{5}$ | $2.92\times10^{5}$ | $8.45\times10^{4}$ |
| S100A16 (C/NU) | $2.86\times10^{4}$ | $1.84\times10^{4}$ | $2.85\times10^{5}$ | $1.47\times10^{5}$ | $3.05\times10^{4}$ | $5.74\times10^{3}$ |
| S100A7 (C/EC) | $1.08\times10^{5}$ | $9.53\times10^{4}$ | $3.62\times10^{6}$ | $1.66\times10^{5}$ | $1.23\times10^{5}$ | $8.65\times10^{4}$ |
| S100A7A (C) | 0 | $3.03\times10^{1}$ | $4.06\times10^{5}$ | $1.51\times10^{4}$ | $9.79\times10^{1}$ | $5.82\times10^{1}$ |
| S100A8 (C/PM/EC/CS) | $4.86\times10^{5}$ | $4.21\times10^{5}$ | $5.11\times10^{7}$ | $3.09\times10^{6}$ | $1.01\times10^{6}$ | $3.22\times10^{5}$ |
| S100A9 (C/CS/PM/EC) | $4.70\times10^{5}$ | $6.53\times10^{5}$ | $9.09\times10^{7}$ | $3.69\times10^{6}$ | $9.22\times10^{5}$ | $5.49\times10^{5}$ |
| S100B (C/NU) | $6.69\times10^{4}$ | $6.67\times10^{4}$ | $1.16\times10^{6}$ | $9.71\times10^{5}$ | $2.92\times10^{4}$ | $2.28\times10^{4}$ |
| S100P (C/NU/PM) | $1.97\times10^{3}$ | $1.39\times10^{4}$ | $3.75\times10^{5}$ | $4.47\times10^{4}$ | $1.24\times10^{4}$ | $6.34\times10^{2}$ |
| SCEL (C/PM) | $2.85\times10^{2}$ | $8.29\times10^{2}$ | $5.37\times10^{1}$ | $4.40\times10^{4}$ | $5.38\times10^{2}$ | $1.10\times10^{3}$ |
| SDCBP2 (C/NU/PM) | $5.10\times10^{2}$ | 0 | $4.27\times10^{4}$ | $3.78\times10^{4}$ | 0 | $9.51\times10^{2}$ |
| SEPTIN7 (C/CS) | $2.16\times10^{2}$ | $2.48\times10^{2}$ | $2.29\times10^{3}$ | $1.50\times10^{4}$ | $4.33\times10^{1}$ | $8.79\times10^{2}$ |
| SERBP1 (C/NU) | $6.69\times10^{3}$ | $4.07\times10^{4}$ | $9.32\times10^{3}$ | $9.31\times10^{4}$ | $1.60\times10^{4}$ | $3.34\times10^{4}$ |
| SERPINA1 (ER/EC) | $3.05\times10^{4}$ | $3.20\times10^{4}$ | $3.21\times10^{5}$ | $6.83\times10^{5}$ | $9.63\times10^{4}$ | $1.78\times10^{4}$ |
| SERPINB1 (C/ES/LY/EC) | 0 | $8.41\times10^{-1}$ | $4.53\times10^{3}$ | $2.57\times10^{4}$ | $7.53\times10^{2}$ | $2.50\times10^{3}$ |
| SERPINB12 (C) | $1.84\times10^{6}$ | $8.47\times10^{5}$ | $7.61\times10^{6}$ | $6.18\times10^{5}$ | $2.04\times10^{6}$ | $4.19\times10^{5}$ |
| SERPINB13 (C) | $3.68\times10^{4}$ | $1.25\times10^{4}$ | $2.08\times10^{5}$ | $1.07\times10^{5}$ | $1.59\times10^{4}$ | $1.45\times10^{4}$ |
| SERPINB2 (C/EC) | $9.09\times10^{4}$ | $4.15\times10^{4}$ | $3.14\times10^{5}$ | $3.45\times10^{5}$ | $8.52\times10^{4}$ | $1.77\times10^{4}$ |
| SERPINB3 (C) | $6.56\times10^{4}$ | $8.42\times10^{4}$ | $1.09\times10^{7}$ | $2.54\times10^{6}$ | $2.93\times10^{5}$ | $5.44\times10^{4}$ |
| SERPINB4 (C) | $4.28\times10^{4}$ | $6.53\times10^{4}$ | $6.37\times10^{6}$ | $4.05\times10^{6}$ | $1.61\times10^{5}$ | $5.22\times10^{4}$ |
| SERPINB7 (C) | $2.29\times10^{4}$ | $1.66\times10^{4}$ | $2.24\times10^{5}$ | $3.87\times10^{4}$ | $2.63\times10^{4}$ | $2.21\times10^{3}$ |
| SERPINB8 (C) | $3.42\times10^{4}$ | $8.79\times10^{3}$ | $1.15\times10^{5}$ | $4.72\times10^{4}$ | $1.33\times10^{4}$ | $2.72\times10^{3}$ |
| SFN (C/NU/EC) | $1.43\times10^{5}$ | $9.79\times10^{4}$ | $6.25\times10^{6}$ | $1.17\times10^{7}$ | $1.05\times10^{5}$ | $8.99\times10^{4}$ |
| SFPQ (C/NU) | $6.00\times10^{3}$ | $1.15\times10^{4}$ | $5.58\times10^{4}$ | $1.72\times10^{5}$ | $1.41\times10^{4}$ | $3.20\times10^{4}$ |
| SHMT2(C/NU/MT) | $1.02\times10^{4}$ | $4.27\times10^{4}$ | $1.61\times10^{1}$ | $1.45\times10^{4}$ | $4.50\times10^{3}$ | $4.55\times10^{4}$ |
| SKP1 (C/NU) | $2.95\times10^{3}$ | $7.29\times10^{3}$ | $3.75\times10^{4}$ | $3.65\times10^{4}$ | $1.07\times10^{5}$ | $1.17\times10^{6}$ |
| SLC25A5 (MT) | $4.94\times10^{4}$ | $9.14\times10^{4}$ | $4.14\times10^{4}$ | $2.77\times10^{5}$ | $1.39\times10^{4}$ | $5.80\times10^{4}$ |
| SLC25A6 (MT) | $4.80\times10^{3}$ | $7.67\times10^{3}$ | $5.97\times10^{3}$ | $4.33\times10^{4}$ | $2.37\times10^{3}$ | $6.75\times10^{3}$ |
| SND1 (C/NU) | 0 | $4.14\times10^{2}$ | $1.60\times10^{4}$ | $5.09\times10^{4}$ | 0 | $4.44\times10^{3}$ |
| SNRPB (C/NU) | $2.64\times10^{3}$ | $5.98\times10^{3}$ | $1.58\times10^{4}$ | $1.64\times10^{4}$ | $2.18\times10^{3}$ | $1.42\times10^{3}$ |
| SNRPB2 (NU) | $9.49\times10^{1}$ | $1.53\times10^{3}$ | 0 | $5.21\times10^{3}$ | $1.48\times10^{2}$ | $5.75\times10^{2}$ |
| SNRPD3 (C/NU) | $1.13\times10^{4}$ | $2.59\times10^{4}$ | $6.03\times10^{4}$ | $5.14\times10^{4}$ | $1.21\times10^{4}$ | $1.19\times10^{4}$ |
| SNRPF (C/NU) | $1.60\times10^{3}$ | $9.59\times10^{3}$ | $1.42\times10^{4}$ | $2.16\times10^{4}$ | $3.56\times10^{3}$ | $9.24\times10^{3}$ |
| SOD2 (MT) | $1.30\times10^{3}$ | $2.52\times10^{3}$ | $5.07\times10^{4}$ | $6.39\times10^{4}$ | $1.19\times10^{3}$ | $5.20\times10^{2}$ |
| SPTAN1 (CS/PM) | $3.94\times10^{1}$ | $1.44\times10^{2}$ | $3.28\times10^{4}$ | $1.05\times10^{5}$ | $3.29\times10^{4}$ | $9.90\times10^{1}$ |
| SRSF2 (NU) | $2.22\times10^{4}$ | $6.38\times10^{4}$ | $2.74\times10^{4}$ | $1.55\times10^{5}$ | $4.77\times10^{4}$ | $3.72\times10^{4}$ |
| SRSF7 (C/NU) | $2.93\times10^{4}$ | $1.05\times10^{5}$ | $1.73\times10^{5}$ | $1.73\times10^{5}$ | $1.04\times10^{5}$ | $9.17\times10^{4}$ |
| SSB (NU) | 0 | $2.70\times10^{3}$ | $3.73\times10^{3}$ | $2.95\times10^{4}$ | $1.30\times10^{3}$ | $4.93\times10^{3}$ |
| SSR4 (ER) | $4.59\times10^{3}$ | $6.08\times10^{3}$ | $8.55\times10^{4}$ | $2.94\times10^{4}$ | $1.28\times10^{4}$ | $2.00\times10^{3}$ |
| ST13 (C) | $1.16\times10^{3}$ | $1.51\times10^{3}$ | $1.51\times10^{3}$ | $8.01\times10^{4}$ | $1.14\times10^{3}$ | $8.71\times10^{3}$ |
| SULT2B1 (C/NU/ER) | $2.73\times10^{4}$ | $9.76\times10^{3}$ | $1.70\times10^{5}$ | $4.15\times10^{5}$ | $4.49\times10^{3}$ | $6.64\times10^{3}$ |
| SYNCRIP (C/NU/ER) | $1.48\times10^{3}$ | $7.12\times10^{3}$ | $8.18\times10^{3}$ | $6.28\times10^{4}$ | $2.84\times10^{3}$ | $1.60\times10^{4}$ |
| TACSTD2 (PM) | 0 | 0 | $3.47\times10^{3}$ | $6.59\times10^{4}$ | 0 | $4.02\times10^{2}$ |
| TAF15 (C/NU) | $4.77\times10^{3}$ | $2.70\times10^{4}$ | $3.93\times10^{4}$ | $1.18\times10^{5}$ | $2.77\times10^{5}$ | $3.37\times10^{3}$ |
| TAGLN2 (C/EC) | $1.96\times10^{4}$ | $7.78\times10^{3}$ | $5.66\times10^{4}$ | $6.14\times10^{5}$ | $2.74\times10^{3}$ | $8.57\times10^{3}$ |
| TALDO1 (C) | $1.64\times10^{4}$ | $2.12\times10^{4}$ | $3.24\times10^{5}$ | $2.09\times10^{5}$ | $3.43\times10^{4}$ | $3.40\times10^{4}$ |
| TCP1 (C/CS) | $3.42\times10^{2}$ | $6.82\times10^{2}$ | $5.48\times10^{3}$ | $5.20\times10^{4}$ | $5.51\times10^{2}$ | $1.70\times10^{3}$ |
| TGM1 (C/PM) | $5.21\times10^{5}$ | $4.34\times10^{5}$ | $4.00\times10^{6}$ | $8.58\times10^{5}$ | $8.10\times10^{5}$ | $4.16\times10^{5}$ |

### Table S3
### (page 9)

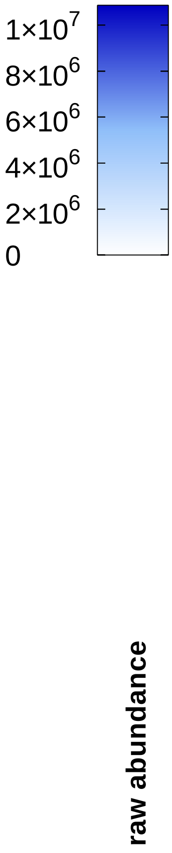


**po-$H_2O_2$ / c-YAP1C**

| 0 | 2 | 5 | 15 | 30 | 60 | |
|---|---|---|---|---|---|---|
| $9.92\times10^{5}$ | $1.02\times10^{6}$ | $7.20\times10^{6}$ | $1.45\times10^{6}$ | $1.25\times10^{6}$ | $3.90\times10^{5}$ | TGM3 (C) |
| $6.80\times10^{3}$ | $4.69\times10^{3}$ | $2.21\times10^{5}$ | $1.83\times10^{4}$ | $1.19\times10^{4}$ | $7.22\times10^{2}$ | TGM5 (C) |
| $1.40\times10^{5}$ | $9.93\times10^{4}$ | $9.16\times10^{5}$ | $3.95\times10^{5}$ | $7.09\times10^{4}$ | $6.02\times10^{4}$ | TKT (C/NU/ER/PO/EC) |
| $1.10\times10^{4}$ | $1.24\times10^{4}$ | $9.88\times10^{4}$ | $9.20\times10^{4}$ | $1.35\times10^{4}$ | $8.88\times10^{3}$ | TMED10 (PM/ER/GA) |
| 0 | 0 | $4.83\times10^{4}$ | 0 | 0 | 0 | TMEM40 (PM) |
| $1.38\times10^{4}$ | $1.93\times10^{4}$ | $1.02\times10^{5}$ | $9.20\times10^{4}$ | $1.54\times10^{4}$ | $6.88\times10^{3}$ | TOLLIP (C) |
| $3.84\times10^{5}$ | $3.21\times10^{5}$ | $2.83\times10^{6}$ | $1.10\times10^{6}$ | $2.70\times10^{5}$ | $1.34\times10^{5}$ | TPI1 (C) |
| $8.05\times10^{4}$ | $1.04\times10^{5}$ | $6.74\times10^{5}$ | $6.80\times10^{5}$ | $3.31\times10^{5}$ | $1.14\times10^{6}$ | TPM3 (CS) |
| $3.39\times10^{3}$ | $4.86\times10^{3}$ | $1.26\times10^{4}$ | $5.18\times10^{4}$ | $1.49\times10^{4}$ | $6.39\times10^{4}$ | TPM4 (CS) |
| $2.98\times10^{3}$ | 0 | $9.83\times10^{3}$ | $1.30\times10^{5}$ | 0 | $7.30\times10^{2}$ | TPPP3 (C/CS) |
| $5.99\times10^{3}$ | $7.96\times10^{3}$ | $2.27\times10^{5}$ | $2.50\times10^{5}$ | $3.76\times10^{3}$ | $1.14\times10^{4}$ | TPT1 (C) |
| $5.35\times10^{4}$ | $1.24\times10^{5}$ | $3.73\times10^{6}$ | $1.14\times10^{6}$ | $3.93\times10^{5}$ | $1.02\times10^{5}$ | TRIM29 (C/LY) |
| $3.05\times10^{4}$ | $5.67\times10^{4}$ | $4.33\times10^{5}$ | $1.14\times10^{6}$ | $5.27\times10^{4}$ | $1.06\times10^{5}$ | TUBA1C (CS) |
| $4.97\times10^{3}$ | $2.14\times10^{3}$ | $2.50\times10^{5}$ | $5.09\times10^{5}$ | $3.82\times10^{3}$ | $2.29\times10^{3}$ | TUBA4A (CS) |
| $2.05\times10^{4}$ | $1.72\times10^{4}$ | $1.27\times10^{5}$ | $4.48\times10^{5}$ | $1.70\times10^{4}$ | $4.62\times10^{4}$ | TUBB (CS) |
| $4.82\times10^{3}$ | $4.71\times10^{3}$ | $3.70\times10^{5}$ | $1.67\times10^{6}$ | $8.33\times10^{3}$ | $1.96\times10^{4}$ | TUBB4B (CS) |
| 0 | 0 | $1.38\times10^{4}$ | $2.41\times10^{4}$ | 0 | 0 | TUBB6 (CS) |
| $2.67\times10^{3}$ | $1.07\times10^{5}$ | $2.37\times10^{4}$ | $8.94\times10^{4}$ | $1.43\times10^{4}$ | $3.96\times10^{4}$ | TUFM (MT) |
| $1.20\times10^{6}$ | $1.12\times10^{6}$ | $3.46\times10^{6}$ | $1.89\times10^{6}$ | $1.81\times10^{6}$ | $4.07\times10^{6}$ | TXN (C/NU/EC) |
| $1.00\times10^{4}$ | $1.46\times10^{4}$ | $1.38\times10^{6}$ | $2.78\times10^{5}$ | $1.20\times10^{4}$ | $2.66\times10^{4}$ | TYMP (C) |
| $1.04\times10^{4}$ | $1.25\times10^{4}$ | $1.51\times10^{5}$ | $3.96\times10^{5}$ | $3.24\times10^{4}$ | $1.15\times10^{5}$ | UBA1 (C/NU/MT) |
| $1.03\times10^{3}$ | $2.36\times10^{3}$ | $4.18\times10^{4}$ | $9.10\times10^{4}$ | $2.43\times10^{2}$ | $4.97\times10^{3}$ | UBE2N (C/NU) |
| $4.08\times10^{3}$ | $1.30\times10^{4}$ | $1.75\times10^{4}$ | $5.86\times10^{4}$ | $1.26\times10^{4}$ | $9.74\times10^{4}$ | UCHL1 (C/ER) |
| $3.24\times10^{3}$ | $2.69\times10^{3}$ | $2.91\times10^{4}$ | $4.60\times10^{4}$ | $1.48\times10^{4}$ | $6.15\times10^{4}$ | UCHL3 (C) |
| $1.24\times10^{4}$ | $4.98\times10^{3}$ | $5.23\times10^{3}$ | $2.87\times10^{4}$ | $4.00\times10^{3}$ | $9.31\times10^{3}$ | UFM1 (C/NU) |
| $1.48\times10^{3}$ | $4.15\times10^{2}$ | $2.69\times10^{3}$ | $9.83\times10^{3}$ | 0 | $9.22\times10^{2}$ | UGP2 (C) |
| $2.84\times10^{3}$ | $2.98\times10^{3}$ | $6.55\times10^{4}$ | $8.28\times10^{4}$ | $2.18\times10^{3}$ | $9.87\times10^{3}$ | USP5 (C/NU/LY) |
| $1.79\times10^{3}$ | $1.71\times10^{3}$ | $4.05\times10^{4}$ | $1.70\times10^{5}$ | $2.65\times10^{3}$ | $2.63\times10^{2}$ | VAT1 (C/MT) |
| $6.64\times10^{4}$ | $4.08\times10^{4}$ | $5.28\times10^{5}$ | $2.03\times10^{5}$ | $6.69\times10^{4}$ | $2.73\times10^{4}$ | VCL (PM/CS) |
| $5.32\times10^{4}$ | $5.00\times10^{4}$ | $1.50\times10^{6}$ | $1.16\times10^{6}$ | $4.96\times10^{4}$ | $6.10\times10^{4}$ | VCP (C/NU/ER) |
| $4.11\times10^{4}$ | $2.12\times10^{4}$ | $6.34\times10^{5}$ | $4.38\times10^{5}$ | $2.34\times10^{4}$ | $1.08\times10^{4}$ | VDAC1 (MT/PM) |
| $2.25\times10^{4}$ | $1.53\times10^{4}$ | $1.09\times10^{5}$ | $1.14\times10^{5}$ | $7.94\times10^{3}$ | $8.73\times10^{3}$ | VDAC2 (MT) |
| $6.29\times10^{5}$ | $1.71\times10^{6}$ | $2.74\times10^{5}$ | $2.34\times10^{6}$ | $6.86\times10^{5}$ | $4.66\times10^{5}$ | VIM (C/NU/PM) |
| $4.97\times10^{3}$ | $1.28\times10^{4}$ | $4.60\times10^{4}$ | $8.37\times10^{4}$ | $1.42\times10^{4}$ | $1.16\times10^{4}$ | WDR1 (C/CS) |
| $1.17\times10^{4}$ | $8.76\times10^{3}$ | $2.44\times10^{4}$ | $1.76\times10^{4}$ | $2.59\times10^{4}$ | $1.16\times10^{5}$ | WDR77 (C/NU) |
| $1.97\times10^{3}$ | $1.43\times10^{3}$ | $2.87\times10^{4}$ | $6.38\times10^{4}$ | $2.99\times10^{3}$ | $6.70\times10^{3}$ | XRCC6 (NU) |
| $2.03\times10^{4}$ | $1.34\times10^{4}$ | $1.83\times10^{5}$ | $4.37\times10^{5}$ | $1.73\times10^{4}$ | $4.52\times10^{4}$ | YWHAB (C) |
| $1.48\times10^{5}$ | $1.32\times10^{5}$ | $4.39\times10^{5}$ | $6.12\times10^{5}$ | $1.13\times10^{5}$ | $2.29\times10^{5}$ | YWHAE (C/NU) |
| 0 | $2.12\times10^{2}$ | $1.24\times10^{4}$ | $9.07\times10^{3}$ | $1.68\times10^{2}$ | $1.57\times10^{3}$ | YWHAG (C) |
| $1.54\times10^{3}$ | $6.86\times10^{3}$ | $7.51\times10^{4}$ | $1.64\times10^{5}$ | $1.36\times10^{4}$ | $8.01\times10^{4}$ | YWHAQ (C) |
| $7.31\times10^{4}$ | $9.86\times10^{4}$ | $2.24\times10^{6}$ | $3.50\times10^{6}$ | $1.13\times10^{5}$ | $2.52\times10^{5}$ | YWHAZ (C) |
| $1.19\times10^{3}$ | $1.31\times10^{3}$ | $3.73\times10^{4}$ | $1.16\times10^{5}$ | $1.12\times10^{3}$ | $1.47\times10^{3}$ | ZNF185 (CS) |

#### Table S4
#### (page 1)

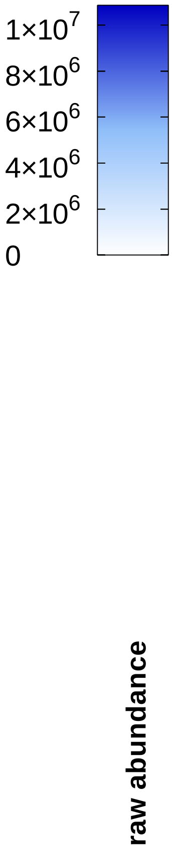


**po-$H_2O_2$ / po-YAP1C**

| | 0 | 2 | 5 | 15 | 30 | 60 |
|---|---|---|---|---|---|---|
| ACTG1 (CS) | $1.41\times10^{6}$ | $9.73\times10^{5}$ | $1.79\times10^{6}$ | $1.46\times10^{6}$ | $8.84\times10^{5}$ | $3.21\times10^{6}$ |
| ALDOA (C/NU) | $9.32\times10^{4}$ | $8.38\times10^{4}$ | $7.67\times10^{4}$ | $1.95\times10^{5}$ | $2.37\times10^{5}$ | $1.04\times10^{5}$ |
| ALOX12B (C) | $6.73\times10^{4}$ | $1.76\times10^{5}$ | $1.68\times10^{5}$ | $3.55\times10^{4}$ | $4.19\times10^{5}$ | $1.14\times10^{5}$ |
| ALOXE3 (C) | $3.01\times10^{3}$ | $3.91\times10^{4}$ | $2.23\times10^{4}$ | $3.03\times10^{3}$ | $9.54\times10^{4}$ | $1.28\times10^{4}$ |
| ANXA1 (C/NU/PM/EC) | $1.05\times10^{4}$ | $9.41\times10^{4}$ | $6.16\times10^{3}$ | $2.13\times10^{4}$ | $1.81\times10^{4}$ | $6.72\times10^{3}$ |
| ANXA2 (PM/EC) | $6.03\times10^{5}$ | $8.87\times10^{5}$ | $8.98\times10^{5}$ | $5.98\times10^{5}$ | $2.71\times10^{6}$ | $2.09\times10^{6}$ |
| APOA1 (C/PM/EC) | $2.60\times10^{5}$ | $1.84\times10^{5}$ | $1.98\times10^{5}$ | $1.76\times10^{5}$ | $7.33\times10^{6}$ | $1.84\times10^{5}$ |
| ARF5 (NU/GA) | $8.36\times10^{3}$ | $9.99\times10^{3}$ | $1.87\times10^{4}$ | $1.97\times10^{4}$ | $1.03\times10^{4}$ | $2.56\times10^{4}$ |
| ARG1 (C) | $4.40\times10^{5}$ | $5.86\times10^{5}$ | $5.79\times10^{5}$ | $2.37\times10^{5}$ | $3.05\times10^{6}$ | $4.36\times10^{5}$ |
| ASAH1 (C/LY/EC) | $3.36\times10^{3}$ | $3.02\times10^{4}$ | $2.05\times10^{4}$ | $5.39\times10^{3}$ | $3.37\times10^{4}$ | $7.84\times10^{3}$ |
| ASPRV1 (C/NU) | $7.54\times10^{3}$ | $1.75\times10^{4}$ | $4.60\times10^{4}$ | $1.75\times10^{4}$ | $8.24\times10^{4}$ | $4.60\times10^{4}$ |
| BLMH (C) | $1.53\times10^{5}$ | $7.33\times10^{5}$ | $3.99\times10^{5}$ | $2.06\times10^{5}$ | $2.34\times10^{6}$ | $2.38\times10^{5}$ |
| CAPN1 (C/PM) | $1.27\times10^{4}$ | $4.37\times10^{4}$ | $2.61\times10^{4}$ | $1.08\times10^{4}$ | $8.57\times10^{4}$ | $2.96\times10^{4}$ |
| CASP14 (C/NU) | $4.47\times10^{5}$ | $1.02\times10^{6}$ | $7.74\times10^{5}$ | $2.78\times10^{5}$ | $2.27\times10^{6}$ | $4.40\times10^{5}$ |
| CAT (PO) | $4.44\times10^{5}$ | $4.93\times10^{5}$ | $7.51\times10^{5}$ | $2.84\times10^{5}$ | $2.03\times10^{6}$ | $8.04\times10^{5}$ |
| CDK4 (C/NU) | $2.57\times10^{3}$ | $4.51\times10^{3}$ | $3.16\times10^{3}$ | $5.05\times10^{3}$ | $6.50\times10^{4}$ | $8.27\times10^{4}$ |
| CDKN2A (C/NU) | $2.38\times10^{3}$ | $2.97\times10^{3}$ | $2.49\times10^{3}$ | $1.87\times10^{4}$ | $3.72\times10^{4}$ | $1.62\times10^{5}$ |
| CLU (C) | $9.00\times10^{4}$ | $8.96\times10^{4}$ | $5.30\times10^{6}$ | $1.52\times10^{5}$ | $7.68\times10^{5}$ | $4.84\times10^{4}$ |
| CRKL (C/NU/PM) | $3.18\times10^{1}$ | 0 | $7.92\times10^{3}$ | $2.51\times10^{2}$ | $2.76\times10^{3}$ | $1.62\times10^{4}$ |
| CSTA (C) | $3.44\times10^{5}$ | $6.45\times10^{5}$ | $6.43\times10^{5}$ | $2.59\times10^{5}$ | $8.01\times10^{5}$ | $3.50\times10^{5}$ |
| CSTB (C/NU) | $5.21\times10^{3}$ | $3.95\times10^{4}$ | $2.01\times10^{4}$ | $7.00\times10^{4}$ | $9.24\times10^{4}$ | $5.23\times10^{5}$ |
| CTSA (LY) | $7.28\times10^{3}$ | $1.40\times10^{4}$ | $1.92\times10^{4}$ | $7.16\times10^{3}$ | $8.23\times10^{4}$ | $1.56\times10^{4}$ |
| CTSD (LY/EC) | $1.91\times10^{5}$ | $2.31\times10^{5}$ | $5.32\times10^{5}$ | $1.98\times10^{5}$ | $9.86\times10^{5}$ | $2.36\times10^{5}$ |
| DSG1 (PM) | $4.15\times10^{6}$ | $4.90\times10^{6}$ | $7.95\times10^{6}$ | $3.08\times10^{6}$ | $1.87\times10^{7}$ | $6.87\times10^{6}$ |
| DSP (CS/PM) | $7.17\times10^{6}$ | $7.33\times10^{6}$ | $1.22\times10^{7}$ | $4.76\times10^{6}$ | $1.20\times10^{7}$ | $1.72\times10^{7}$ |
| EEF1A1 (C/NU/PM) | $6.33\times10^{4}$ | $1.28\times10^{5}$ | $1.26\times10^{5}$ | $9.69\times10^{4}$ | $6.24\times10^{4}$ | $2.86\times10^{5}$ |
| ENO1 (C/NU) | $4.59\times10^{5}$ | $2.91\times10^{5}$ | $8.11\times10^{5}$ | $5.06\times10^{5}$ | $3.96\times10^{5}$ | $1.15\times10^{6}$ |
| FABP5 (C/NU/EC) | $2.34\times10^{5}$ | $3.28\times10^{5}$ | $9.70\times10^{5}$ | $4.76\times10^{5}$ | $5.28\times10^{5}$ | $3.01\times10^{6}$ |
| FLG (C/PM) | $3.09\times10^{5}$ | $2.41\times10^{5}$ | $8.92\times10^{5}$ | $2.09\times10^{5}$ | $1.63\times10^{6}$ | $4.76\times10^{5}$ |
| FLG2 (C) | $1.59\times10^{6}$ | $2.49\times10^{6}$ | $3.01\times10^{6}$ | $1.18\times10^{6}$ | $8.97\times10^{6}$ | $2.23\times10^{6}$ |
| GAPDH (C/NU/CS) | $9.01\times10^{5}$ | $1.19\times10^{6}$ | $1.12\times10^{6}$ | $9.99\times10^{5}$ | $1.89\times10^{6}$ | $1.49\times10^{6}$ |
| GDA (C) | $2.07\times10^{3}$ | $1.00\times10^{3}$ | $1.97\times10^{3}$ | $1.39\times10^{3}$ | $1.27\times10^{4}$ | $2.46\times10^{3}$ |
| GGCT(C/EC) | $1.46\times10^{5}$ | $2.01\times10^{5}$ | $3.21\times10^{5}$ | $9.20\times10^{4}$ | $7.83\times10^{5}$ | $1.84\times10^{5}$ |
| GSDMA (C/PM) | $1.19\times10^{5}$ | $3.13\times10^{5}$ | $3.23\times10^{5}$ | $1.44\times10^{5}$ | $9.65\times10^{5}$ | $2.24\times10^{5}$ |
| GSR (C/MT) | $6.82\times10^{3}$ | $5.84\times10^{3}$ | $6.45\times10^{3}$ | $3.00\times10^{4}$ | $5.86\times10^{4}$ | $2.32\times10^{5}$ |
| GSTP1 (C/NU/MT) | $5.09\times10^{3}$ | $1.91\times10^{4}$ | $3.60\times10^{4}$ | $3.36\times10^{4}$ | $2.34\times10^{4}$ | $5.85\times10^{4}$ |
| HAL (C) | $6.57\times10^{4}$ | $2.36\times10^{5}$ | $1.07\times10^{5}$ | $8.40\times10^{4}$ | $8.38\times10^{5}$ | $1.16\times10^{5}$ |
| HPRT1 (C) | $2.38\times10^{3}$ | $1.31\times10^{3}$ | $6.44\times10^{3}$ | $2.26\times10^{4}$ | $5.75\times10^{4}$ | $2.03\times10^{5}$ |
| HSD17B4 (PO) | $1.96\times10^{2}$ | $6.13\times10^{3}$ | $4.00\times10^{3}$ | $8.80\times10^{2}$ | $3.38\times10^{3}$ | $1.06\times10^{4}$ |
| HSP90AA1 (C/NU/MT/PM) | $1.23\times10^{4}$ | $4.52\times10^{3}$ | $7.91\times10^{3}$ | $1.55\times10^{4}$ | $1.56\times10^{4}$ | $3.17\times10^{4}$ |
| HSPA1A (C/NU/EC) | $2.47\times10^{5}$ | $3.04\times10^{5}$ | $4.67\times10^{5}$ | $3.67\times10^{5}$ | $2.11\times10^{5}$ | $5.98\times10^{5}$ |
| HSPA9 (MT) | $4.22\times10^{3}$ | $6.16\times10^{2}$ | $6.60\times10^{3}$ | $1.69\times10^{3}$ | $3.12\times10^{3}$ | $3.38\times10^{4}$ |
| HSPB1 (C/NU/CS) | $1.36\times10^{5}$ | $2.71\times10^{5}$ | $1.23\times10^{5}$ | $1.59\times10^{5}$ | $1.45\times10^{5}$ | $8.70\times10^{5}$ |
| HSPD1 (MT) | $2.53\times10^{4}$ | $1.29\times10^{4}$ | $3.67\times10^{4}$ | $2.39\times10^{4}$ | $1.33\times10^{4}$ | $8.05\times10^{4}$ |
| LAMP1 (LY) | $1.64\times10^{4}$ | $2.75\times10^{4}$ | $5.84\times10^{4}$ | $2.03\times10^{4}$ | $1.01\times10^{5}$ | $4.42\times10^{4}$ |
| LDHA (C) | $7.30\times10^{4}$ | $5.35\times10^{4}$ | $8.74\times10^{4}$ | $9.84\times10^{4}$ | $6.96\times10^{4}$ | $2.03\times10^{5}$ |
| LDHB (C/MT) | $3.75\times10^{4}$ | $1.91\times10^{4}$ | $4.03\times10^{4}$ | $5.23\times10^{4}$ | $2.67\times10^{4}$ | $8.24\times10^{4}$ |
| LGALS7 (C/NU/EC) | $1.56\times10^{5}$ | $2.52\times10^{5}$ | $2.25\times10^{5}$ | $2.83\times10^{5}$ | $2.27\times10^{5}$ | $1.69\times10^{6}$ |
| LTF (C/EC) | $5.75\times10^{4}$ | $3.45\times10^{4}$ | $2.26\times10^{4}$ | $1.94\times10^{5}$ | $1.15\times10^{6}$ | $5.71\times10^{4}$ |
| NCCRP1 (C/NU/MT) | $7.92\times10^{4}$ | $1.75\times10^{5}$ | $1.94\times10^{5}$ | $8.07\times10^{4}$ | $4.11\times10^{5}$ | $2.14\times10^{5}$ |

### Table S4
### (page 2)

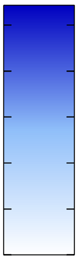


**po-$H_2O_2$ / po-YAP1C**

| Protein | 0 | 2 | 5 | 15 | 30 | 60 |
|---|---|---|---|---|---|---|
| NPM1 (NU) | $2.30\times10^{4}$ | $2.95\times10^{4}$ | $3.92\times10^{4}$ | $4.41\times10^{4}$ | $9.01\times10^{4}$ | $2.70\times10^{5}$ |
| NUDC (CS/NU) | $1.18\times10^{4}$ | $4.53\times10^{3}$ | $9.61\times10^{3}$ | $1.32\times10^{4}$ | $3.22\times10^{4}$ | $9.17\times10^{4}$ |
| PCCA (MT) | $3.27\times10^{5}$ | $2.62\times10^{5}$ | $3.39\times10^{5}$ | $3.70\times10^{5}$ | $2.33\times10^{5}$ | $7.32\times10^{5}$ |
| PCCB (MT) | $1.11\times10^{5}$ | $3.63\times10^{4}$ | $2.76\times10^{5}$ | $5.70\times10^{4}$ | $3.98\times10^{4}$ | $1.24\times10^{6}$ |
| PCMT1 (C) | $1.81\times10^{3}$ | $1.24\times10^{3}$ | $2.69\times10^{3}$ | $5.67\times10^{2}$ | $1.37\times10^{4}$ | $2.09\times10^{4}$ |
| PEBP1 (C) | $1.29\times10^{4}$ | $5.27\times10^{3}$ | $1.95\times10^{4}$ | $1.26\times10^{4}$ | $4.73\times10^{4}$ | $3.79\times10^{4}$ |
| PKM (C/NU) | $4.26\times10^{4}$ | $4.45\times10^{4}$ | $1.10\times10^{5}$ | $7.49\times10^{4}$ | $1.81\times10^{5}$ | $2.24\times10^{5}$ |
| PLEC (CS) | $1.07\times10^{4}$ | $3.83\times10^{4}$ | $2.76\times10^{4}$ | $1.00\times10^{4}$ | $7.26\times10^{4}$ | $1.87\times10^{4}$ |
| PLS3 (C) | $3.33\times10^{3}$ | $5.51\times10^{3}$ | $8.99\times10^{2}$ | $7.61\times10^{3}$ | $1.42\times10^{4}$ | $6.66\times10^{4}$ |
| PNP (C) | $8.87\times10^{3}$ | $1.66\times10^{4}$ | $1.71\times10^{4}$ | $6.89\times10^{3}$ | $3.44\times10^{4}$ | $1.51\times10^{4}$ |
| POF1B (C/CS) | $8.57\times10^{4}$ | $1.20\times10^{5}$ | $1.81\times10^{5}$ | $5.08\times10^{4}$ | $1.49\times10^{5}$ | $2.16\times10^{5}$ |
| PPP2R2B (C/MT/CS) | $7.63\times10^{3}$ | $5.71\times10^{3}$ | $1.78\times10^{4}$ | $1.97\times10^{4}$ | $6.71\times10^{4}$ | $2.33\times10^{5}$ |
| PRDX1 (C) | $3.81\times10^{5}$ | $4.34\times10^{5}$ | $1.22\times10^{6}$ | $2.40\times10^{6}$ | $4.81\times10^{6}$ | $1.24\times10^{7}$ |
| PRDX2 (C) | $3.72\times10^{5}$ | $2.47\times10^{5}$ | $5.47\times10^{5}$ | $7.20\times10^{5}$ | $1.68\times10^{6}$ | $2.01\times10^{6}$ |
| PRDX3 (MT/C/EE) | $1.17\times10^{4}$ | $1.22\times10^{4}$ | $1.75\times10^{4}$ | $4.38\times10^{4}$ | $4.43\times10^{4}$ | $1.48\times10^{5}$ |
| PRDX6 (C/LY) | $4.42\times10^{4}$ | $3.69\times10^{4}$ | $2.16\times10^{4}$ | $5.31\times10^{4}$ | $1.65\times10^{5}$ | $6.80\times10^{5}$ |
| PRMT5 (C/NU/GA) | $2.04\times10^{3}$ | $2.84\times10^{3}$ | $4.74\times10^{3}$ | $2.19\times10^{3}$ | $8.83\times10^{3}$ | $4.73\times10^{4}$ |
| PSMA7 (C/NU) | $8.78\times10^{3}$ | $1.16\times10^{4}$ | $1.25\times10^{4}$ | $4.63\times10^{3}$ | $3.08\times10^{4}$ | $1.17\times10^{4}$ |
| PSMB5 (C/NU) | $5.78\times10^{3}$ | $7.57\times10^{3}$ | $1.26\times10^{4}$ | $4.21\times10^{3}$ | $3.24\times10^{4}$ | $1.10\times10^{4}$ |
| RPS27A (C/NU) | $1.61\times10^{5}$ | $1.73\times10^{5}$ | $2.84\times10^{5}$ | $1.35\times10^{5}$ | $4.33\times10^{5}$ | $3.21\times10^{5}$ |
| S100A14 (C) | $9.09\times10^{4}$ | $1.56\times10^{5}$ | $3.32\times10^{5}$ | $8.51\times10^{4}$ | $3.57\times10^{5}$ | $2.23\times10^{5}$ |
| S100A16 (C/NU) | $1.51\times10^{4}$ | $4.99\times10^{4}$ | $5.03\times10^{4}$ | $1.78\times10^{4}$ | $1.19\times10^{5}$ | $3.41\times10^{4}$ |
| SERPINB12 (C) | $8.21\times10^{5}$ | $1.11\times10^{6}$ | $1.22\times10^{6}$ | $5.97\times10^{5}$ | $1.74\times10^{6}$ | $9.40\times10^{5}$ |
| SERPINB7 (C) | $8.38\times10^{3}$ | $1.74\times10^{4}$ | $3.26\times10^{4}$ | $8.91\times10^{3}$ | $4.32\times10^{4}$ | $1.90\times10^{4}$ |
| SFN (C/NU/EC) | $1.98\times10^{4}$ | $1.41\times10^{4}$ | $1.24\times10^{4}$ | $7.69\times10^{4}$ | $5.64\times10^{4}$ | $8.29\times10^{4}$ |
| SKP1 (C/NU) | $8.23\times10^{4}$ | $1.96\times10^{4}$ | $2.36\times10^{4}$ | $1.54\times10^{5}$ | $1.23\times10^{6}$ | $5.89\times10^{6}$ |
| TGM1 (C/PM) | $2.73\times10^{5}$ | $5.38\times10^{5}$ | $5.60\times10^{5}$ | $1.95\times10^{5}$ | $1.18\times10^{6}$ | $4.23\times10^{5}$ |
| TPI1 (C) | $4.36\times10^{4}$ | $4.86\times10^{4}$ | $1.04\times10^{5}$ | $4.65\times10^{4}$ | $8.97\times10^{4}$ | $1.25\times10^{5}$ |
| TRAP1 (MT) | $2.35\times10^{4}$ | $1.36\times10^{4}$ | $1.54\times10^{4}$ | $4.04\times10^{4}$ | $1.64\times10^{4}$ | $6.98\times10^{4}$ |
| TXN (C/NU/EC) | $3.63\times10^{5}$ | $4.73\times10^{5}$ | $6.09\times10^{5}$ | $5.13\times10^{5}$ | $1.53\times10^{6}$ | $2.27\times10^{6}$ |
| YWHAB (C) | $1.68\times10^{3}$ | $1.18\times10^{3}$ | $1.59\times10^{2}$ | $8.15\times10^{3}$ | $4.98\times10^{3}$ | $4.16\times10^{4}$ |
| YWHAG (C) | $2.82\times10^{2}$ | $1.99\times10^{0}$ | $2.05\times10^{2}$ | $1.23\times10^{3}$ | 0 | $7.10\times10^{3}$ |
| YWHAQ (C) | $3.16\times10^{3}$ | $9.14\times10^{2}$ | $5.14\times10^{2}$ | $9.87\times10^{3}$ | $1.12\times10^{4}$ | $6.63\times10^{4}$ |
| YWHAZ (C) | $8.69\times10^{3}$ | $6.15\times10^{3}$ | $6.54\times10^{3}$ | $3.85\times10^{4}$ | $1.78\times10^{4}$ | $1.52\times10^{5}$ |

### Table S5

po-$H_2O_2$ / mt-YAP1C

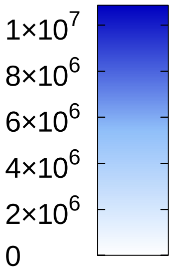


| | 0 | 2 | 5 | 15 | 30 | 60 |
|---|---|---|---|---|---|---|
| ACTG1 (CS) | $1.52\times10^{6}$ | $2.31\times10^{6}$ | $3.31\times10^{6}$ | $2.39\times10^{6}$ | $4.07\times10^{6}$ | $3.30\times10^{6}$ |
| ANXA1 (C/NU/PM/EC) | $1.15\times10^{4}$ | $2.16\times10^{4}$ | $1.89\times10^{4}$ | $1.52\times10^{5}$ | $2.81\times10^{5}$ | $1.49\times10^{4}$ |
| ANXA2 (PM/EC) | $4.78\times10^{5}$ | $7.78\times10^{5}$ | $1.39\times10^{6}$ | $9.19\times10^{5}$ | $1.49\times10^{6}$ | $1.40\times10^{6}$ |
| ARHGDIA (C) | $8.24\times10^{3}$ | $1.08\times10^{4}$ | $2.74\times10^{4}$ | $3.53\times10^{3}$ | $2.42\times10^{4}$ | $2.60\times10^{5}$ |
| ATP5F1A (MT) | $1.64\times10^{4}$ | $5.60\times10^{4}$ | $4.34\times10^{4}$ | $3.06\times10^{4}$ | $1.14\times10^{5}$ | $5.31\times10^{4}$ |
| BLMH (C) | $1.20\times10^{5}$ | $6.50\times10^{4}$ | $1.81\times10^{5}$ | $1.92\times10^{5}$ | $2.45\times10^{5}$ | $3.79\times10^{5}$ |
| CDK4 (C/NU) | $1.86\times10^{2}$ | $5.82\times10^{3}$ | $3.02\times10^{3}$ | $5.70\times10^{3}$ | $8.55\times10^{4}$ | $1.41\times10^{5}$ |
| CKB (C) | $6.99\times10^{4}$ | $1.07\times10^{5}$ | $9.52\times10^{4}$ | $1.09\times10^{5}$ | $2.61\times10^{5}$ | $1.77\times10^{5}$ |
| CSTA (C) | $2.24\times10^{5}$ | $4.64\times10^{5}$ | $4.20\times10^{5}$ | $6.56\times10^{5}$ | $4.21\times10^{5}$ | $2.05\times10^{5}$ |
| CSTB (C/NU) | $8.50\times10^{2}$ | $1.87\times10^{4}$ | $1.72\times10^{4}$ | $1.07\times10^{5}$ | $2.66\times10^{5}$ | $7.06\times10^{4}$ |
| CTSA (LY) | $4.49\times10^{3}$ | $6.95\times10^{3}$ | $1.53\times10^{4}$ | $5.52\times10^{3}$ | $6.28\times10^{3}$ | $3.56\times10^{3}$ |
| CTSD (LY/EC) | $6.49\times10^{4}$ | $1.69\times10^{5}$ | $2.92\times10^{5}$ | $1.48\times10^{5}$ | $1.89\times10^{5}$ | $2.70\times10^{5}$ |
| DSG1 (PM) | $2.92\times10^{6}$ | $5.68\times10^{6}$ | $8.28\times10^{6}$ | $4.95\times10^{6}$ | $5.98\times10^{6}$ | $7.33\times10^{6}$ |
| DSP (CS/PM) | $3.67\times10^{6}$ | $7.58\times10^{6}$ | $9.27\times10^{6}$ | $7.38\times10^{6}$ | $6.57\times10^{6}$ | $6.43\times10^{6}$ |
| EEF1A1 (C/NU/PM) | $1.08\times10^{5}$ | $2.12\times10^{5}$ | $4.12\times10^{5}$ | $1.59\times10^{5}$ | $2.03\times10^{5}$ | $7.41\times10^{5}$ |
| ENO1 (C/NU) | $8.38\times10^{5}$ | $1.41\times10^{6}$ | $1.18\times10^{6}$ | $1.37\times10^{6}$ | $1.81\times10^{6}$ | $1.64\times10^{6}$ |
| FABP5 (C/NU/EC) | $6.08\times10^{4}$ | $2.33\times10^{5}$ | $5.06\times10^{5}$ | $5.52\times10^{5}$ | $2.19\times10^{5}$ | $1.59\times10^{5}$ |
| GGCT(C/EC) | $7.44\times10^{4}$ | $1.12\times10^{5}$ | $1.79\times10^{5}$ | $1.54\times10^{5}$ | $1.74\times10^{5}$ | $1.95\times10^{5}$ |
| GSDMA (C/PM) | $2.22\times10^{5}$ | $2.59\times10^{5}$ | $4.03\times10^{5}$ | $3.53\times10^{5}$ | $4.86\times10^{5}$ | $4.49\times10^{5}$ |
| GSR (C/MT) | $3.26\times10^{4}$ | $6.97\times10^{4}$ | $1.04\times10^{5}$ | $2.76\times10^{5}$ | $7.81\times10^{5}$ | $1.10\times10^{6}$ |
| HNRNPA1 (NU/C) | $3.97\times10^{2}$ | $1.23\times10^{4}$ | $1.01\times10^{4}$ | $2.50\times10^{3}$ | $7.01\times10^{3}$ | $9.92\times10^{3}$ |
| HPRT1 (C) | $6.85\times10^{2}$ | $1.53\times10^{4}$ | $2.53\times10^{4}$ | $5.31\times10^{4}$ | $1.64\times10^{5}$ | $2.49\times10^{5}$ |
| HSD17B10 (MT) | $3.11\times10^{4}$ | $6.63\times10^{4}$ | $1.33\times10^{5}$ | $3.48\times10^{5}$ | $8.72\times10^{5}$ | $2.00\times10^{6}$ |
| HSP90AA1 (C/NU/MT/PM) | $1.79\times10^{4}$ | $1.63\times10^{4}$ | $1.16\times10^{4}$ | $9.09\times10^{3}$ | $4.23\times10^{4}$ | $1.33\times10^{5}$ |
| HSP90AB1 (C/NU/PM/EC) | $3.79\times10^{3}$ | $5.24\times10^{3}$ | $9.77\times10^{3}$ | $3.84\times10^{3}$ | $3.13\times10^{4}$ | $8.71\times10^{4}$ |
| HSPA8 (C/PM/NU) | $1.64\times10^{5}$ | $3.24\times10^{5}$ | $3.31\times10^{5}$ | $2.03\times10^{5}$ | $3.54\times10^{5}$ | $2.13\times10^{5}$ |
| HSPA9 (MT) | $8.56\times10^{5}$ | $9.03\times10^{5}$ | $1.32\times10^{6}$ | $1.03\times10^{6}$ | $1.52\times10^{6}$ | $2.05\times10^{6}$ |
| HSPB1 (C/NU/CS) | $1.89\times10^{4}$ | $1.19\times10^{5}$ | $5.26\times10^{5}$ | $1.92\times10^{5}$ | $2.11\times10^{5}$ | $1.62\times10^{5}$ |
| HSPD1 (MT) | $2.21\times10^{5}$ | $2.04\times10^{5}$ | $2.58\times10^{5}$ | $2.22\times10^{5}$ | $3.86\times10^{5}$ | $6.23\times10^{5}$ |
| JUP (CS/PM) | $1.58\times10^{6}$ | $4.01\times10^{6}$ | $4.04\times10^{6}$ | $3.45\times10^{6}$ | $2.94\times10^{6}$ | $2.91\times10^{6}$ |
| LGALS7 (C/NU/EC) | $1.26\times10^{5}$ | $2.73\times10^{5}$ | $3.36\times10^{6}$ | $3.24\times10^{5}$ | $3.04\times10^{5}$ | $2.16\times10^{5}$ |
| PC (MT) | $3.51\times10^{3}$ | $6.27\times10^{2}$ | $3.17\times10^{3}$ | $6.92\times10^{2}$ | $3.97\times10^{3}$ | $4.73\times10^{5}$ |
| PCCA (MT) | $3.21\times10^{5}$ | $4.46\times10^{5}$ | $4.24\times10^{5}$ | $4.48\times10^{5}$ | $4.65\times10^{5}$ | $6.76\times10^{5}$ |
| PCCB (MT) | $1.58\times10^{5}$ | $3.49\times10^{5}$ | $3.13\times10^{5}$ | $3.82\times10^{5}$ | $7.76\times10^{5}$ | $1.96\times10^{6}$ |
| PCMT1 (C) | $2.70\times10^{4}$ | $5.34\times10^{4}$ | $5.86\times10^{4}$ | $3.61\times10^{4}$ | $8.84\times10^{4}$ | $1.07\times10^{5}$ |
| PKP1 (NU) | $2.44\times10^{5}$ | $5.17\times10^{5}$ | $8.26\times10^{5}$ | $5.13\times10^{5}$ | $3.19\times10^{5}$ | $2.18\times10^{5}$ |
| POF1B (C/CS) | $8.02\times10^{4}$ | $1.13\times10^{5}$ | $3.14\times10^{5}$ | $1.28\times10^{5}$ | $1.19\times10^{5}$ | $8.58\times10^{4}$ |
| PPIA (C/NU/EC) | $3.92\times10^{4}$ | $1.76\times10^{5}$ | $2.11\times10^{5}$ | $9.91\times10^{4}$ | $3.56\times10^{5}$ | $6.38\times10^{5}$ |
| PPP2R2B (C/MT/CS) | $4.38\times10^{2}$ | $1.03\times10^{4}$ | $1.97\times10^{4}$ | $4.93\times10^{4}$ | $1.04\times10^{5}$ | $1.57\times10^{5}$ |
| PRDX1 (C) | $9.01\times10^{5}$ | $1.79\times10^{6}$ | $5.03\times10^{6}$ | $7.35\times10^{6}$ | $1.47\times10^{7}$ | $2.13\times10^{7}$ |
| PRDX2 (C) | $2.49\times10^{5}$ | $4.35\times10^{5}$ | $9.87\times10^{5}$ | $1.12\times10^{6}$ | $1.73\times10^{6}$ | $1.77\times10^{6}$ |
| PRDX3 (MT/C/EE) | $4.73\times10^{5}$ | $3.00\times10^{6}$ | $1.03\times10^{7}$ | $9.98\times10^{6}$ | $1.64\times10^{7}$ | $2.33\times10^{7}$ |
| PRDX6 (C/LY) | $3.44\times10^{3}$ | $2.45\times10^{4}$ | $4.69\times10^{4}$ | $1.01\times10^{5}$ | $5.00\times10^{5}$ | $6.38\times10^{5}$ |
| PRMT5 (C/NU/GA) | $3.43\times10^{3}$ | $3.72\times10^{3}$ | $9.76\times10^{3}$ | $1.80\times10^{4}$ | $4.61\times10^{4}$ | $8.25\times10^{4}$ |
| RPSA (NU/C/PM) | $1.93\times10^{4}$ | $5.35\times10^{4}$ | $5.40\times10^{4}$ | $3.19\times10^{4}$ | $6.65\times10^{4}$ | $4.04\times10^{4}$ |
| S100A14 (C) | $8.61\times10^{4}$ | $1.10\times10^{5}$ | $2.39\times10^{5}$ | $1.32\times10^{5}$ | $1.24\times10^{5}$ | $9.16\times10^{4}$ |
| S100A7 (C/EC) | $4.19\times10^{3}$ | $1.74\times10^{5}$ | $2.67\times10^{5}$ | $7.93\times10^{4}$ | $1.63\times10^{5}$ | $1.45\times10^{5}$ |
| S100A8 (C/PM/EC/CS) | $5.13\times10^{5}$ | $5.69\times10^{5}$ | $1.03\times10^{6}$ | $1.10\times10^{6}$ | $1.09\times10^{6}$ | $5.77\times10^{5}$ |
| SERPINB3 (C) | $1.20\times10^{5}$ | $2.54\times10^{5}$ | $5.20\times10^{5}$ | $3.40\times10^{5}$ | $5.10\times10^{5}$ | $4.70\times10^{5}$ |
| SKP1 (C/NU) | $7.88\times10^{2}$ | $3.54\times10^{3}$ | $1.53\times10^{4}$ | $5.25\times10^{4}$ | $4.43\times10^{5}$ | $1.16\times10^{6}$ |
| TUBA1C (CS) | $1.02\times10^{3}$ | $1.09\times10^{5}$ | $4.26\times10^{4}$ | $9.09\times10^{3}$ | $1.68\times10^{5}$ | $8.09\times10^{4}$ |
| TXN (C/NU/EC) | $3.19\times10^{5}$ | $5.27\times10^{5}$ | $7.39\times10^{5}$ | $8.25\times10^{5}$ | $1.96\times10^{6}$ | $2.43\times10^{6}$ |
| UCHL1 (C/ER) | $1.47\times10^{4}$ | $7.21\times10^{3}$ | $9.42\times10^{3}$ | $7.20\times10^{3}$ | $1.99\times10^{4}$ | $3.95\times10^{4}$ |
| VDAC1 (MT/PM) | 0 | $2.64\times10^{3}$ | $1.36\times10^{4}$ | 0 | $1.32\times10^{4}$ | 0 |
| YWHAQ (C) | $8.01\times10^{3}$ | $6.92\times10^{3}$ | $7.82\times10^{3}$ | $1.59\times10^{4}$ | $4.80\times10^{4}$ | $8.31\times10^{4}$ |
| YWHAZ (C) | $3.30\times10^{4}$ | $2.39\times10^{4}$ | $4.68\times10^{4}$ | $5.49\times10^{4}$ | $1.07\times10^{5}$ | $1.39\times10^{5}$ |

### Table S6 (page 1)

#### external-$H_2O_2$ / c-YAP1C / 10 min

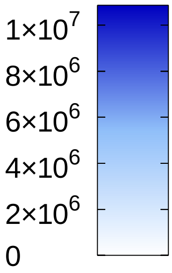


raw abundance

| 0 | 10 | 30 | 100 | 300 | 1000 | |
|---|---|---|---|---|---|---|
| $5.66\times10^{4}$ | $4.59\times10^{4}$ | $5.82\times10^{4}$ | $1.31\times10^{5}$ | $1.44\times10^{5}$ | $2.05\times10^{5}$ | AAMDC (C) |
| $9.52\times10^{4}$ | $1.56\times10^{5}$ | $6.25\times10^{5}$ | $1.11\times10^{6}$ | $9.53\times10^{5}$ | $8.51\times10^{5}$ | AASDHPPT (C) |
| $4.63\times10^{4}$ | $5.61\times10^{4}$ | $6.09\times10^{4}$ | $9.36\times10^{4}$ | $1.43\times10^{5}$ | $2.61\times10^{5}$ | ACP1 (C) |
| 0 | 0 | $1.29\times10^{3}$ | $3.62\times10^{3}$ | $1.96\times10^{4}$ | $6.24\times10^{4}$ | ACP6 (MT) |
| $3.49\times10^{3}$ | $1.00\times10^{4}$ | $2.78\times10^{4}$ | $2.85\times10^{4}$ | $7.03\times10^{4}$ | $5.80\times10^{4}$ | ACT (???) |
| $7.00\times10^{4}$ | $9.97\times10^{4}$ | $2.26\times10^{5}$ | $3.69\times10^{5}$ | $3.09\times10^{5}$ | $3.47\times10^{5}$ | ACTN1 (C/PM) |
| $4.60\times10^{1}$ | $3.57\times10^{2}$ | $4.31\times10^{3}$ | $1.09\times10^{3}$ | $7.28\times10^{2}$ | $2.44\times10^{3}$ | ACYP2 (?) |
| $2.16\times10^{6}$ | $2.37\times10^{6}$ | $3.31\times10^{6}$ | $8.10\times10^{6}$ | $7.37\times10^{6}$ | $7.50\times10^{6}$ | AHCY (C) |
| $1.32\times10^{3}$ | $2.41\times10^{3}$ | $2.88\times10^{4}$ | $5.13\times10^{4}$ | $4.03\times10^{4}$ | $5.05\times10^{4}$ | AIF1L (CS/PM) |
| $3.71\times10^{5}$ | $4.99\times10^{5}$ | $8.70\times10^{5}$ | $1.68\times10^{6}$ | $1.66\times10^{6}$ | $2.71\times10^{6}$ | AK2 (MT) |
| $5.20\times10^{5}$ | $6.22\times10^{5}$ | $8.18\times10^{5}$ | $1.63\times10^{6}$ | $1.32\times10^{6}$ | $1.08\times10^{6}$ | AKAP12 |
| $1.81\times10^{4}$ | $1.71\times10^{4}$ | $1.35\times10^{4}$ | $3.35\times10^{4}$ | $2.08\times10^{4}$ | $1.41\times10^{5}$ | ANXA1 (C/NU/PM/EC) |
| $5.16\times10^{6}$ | $7.30\times10^{6}$ | $1.90\times10^{7}$ | $2.61\times10^{7}$ | $2.23\times10^{7}$ | $2.15\times10^{7}$ | ANXA2 (PM/EC) |
| $3.26\times10^{3}$ | $7.29\times10^{3}$ | $6.34\times10^{3}$ | $3.95\times10^{3}$ | $8.53\times10^{3}$ | $1.05\times10^{4}$ | ANXA6 (C) |
| $1.08\times10^{4}$ | $2.56\times10^{4}$ | $3.93\times10^{4}$ | $6.00\times10^{4}$ | $6.72\times10^{4}$ | $6.08\times10^{4}$ | APIP (C) |
| $1.86\times10^{5}$ | $1.28\times10^{5}$ | $1.16\times10^{5}$ | $1.94\times10^{5}$ | $1.41\times10^{5}$ | $2.20\times10^{6}$ | APOA1 (C/PM/EC) |
| $4.85\times10^{4}$ | $7.57\times10^{4}$ | $6.18\times10^{4}$ | $8.47\times10^{4}$ | $9.74\times10^{4}$ | $7.00\times10^{4}$ | ARF4 (GA/PM) |
| 0 | 0 | $2.88\times10^{4}$ | $5.74\times10^{4}$ | $5.37\times10^{4}$ | $8.61\times10^{4}$ | ARL6IP4 (NU) |
| $1.00\times10^{4}$ | $6.18\times10^{4}$ | $2.96\times10^{5}$ | $4.60\times10^{5}$ | $4.81\times10^{5}$ | $4.77\times10^{5}$ | ASF1A (NU) |
| $1.28\times10^{5}$ | $2.43\times10^{5}$ | $7.00\times10^{5}$ | $1.15\times10^{6}$ | $9.31\times10^{5}$ | $7.64\times10^{5}$ | ASF1B (NU) |
| $2.66\times10^{4}$ | $2.90\times10^{4}$ | $2.44\times10^{4}$ | $1.85\times10^{5}$ | $5.12\times10^{5}$ | $9.80\times10^{5}$ | ASNS (C) |
| 0 | $1.21\times10^{4}$ | $4.27\times10^{4}$ | $1.18\times10^{5}$ | $1.08\times10^{5}$ | $1.06\times10^{5}$ | ASRGL1 (C) |
| $4.09\times10^{4}$ | $4.74\times10^{4}$ | $5.90\times10^{4}$ | $1.46\times10^{5}$ | $8.48\times10^{4}$ | $1.07\times10^{5}$ | ATG3 (C) |
| $6.31\times10^{3}$ | $9.52\times10^{3}$ | $2.31\times10^{4}$ | $4.08\times10^{4}$ | $4.75\times10^{4}$ | $1.24\times10^{5}$ | ATOX1 (C) |
| $2.32\times10^{2}$ | $1.57\times10^{3}$ | $1.11\times10^{3}$ | $1.86\times10^{3}$ | $1.15\times10^{3}$ | 0 | ATP5F1B (MT) |
| $1.67\times10^{5}$ | $1.43\times10^{5}$ | $2.18\times10^{5}$ | $3.55\times10^{5}$ | $3.40\times10^{5}$ | $3.91\times10^{5}$ | ATP6V1A (C) |
| $3.05\times10^{4}$ | $4.74\times10^{4}$ | $2.94\times10^{4}$ | $7.57\times10^{4}$ | $5.44\times10^{4}$ | $3.59\times10^{4}$ | ATPAF1 (MT) |
| $5.50\times10^{4}$ | $8.88\times10^{4}$ | $3.90\times10^{5}$ | $8.45\times10^{5}$ | $6.16\times10^{5}$ | $6.73\times10^{5}$ | BAG3 (C/NU) |
| $1.65\times10^{4}$ | $2.90\times10^{4}$ | $5.45\times10^{4}$ | $1.40\times10^{5}$ | $1.31\times10^{5}$ | $1.03\times10^{5}$ | BAG4 (C) |
| $7.34\times10^{4}$ | $1.98\times10^{5}$ | $1.08\times10^{6}$ | $1.58\times10^{6}$ | $1.41\times10^{6}$ | $1.34\times10^{6}$ | BAG5 (C/NU/MT) |
| $1.92\times10^{5}$ | $2.32\times10^{5}$ | $2.37\times10^{5}$ | $4.53\times10^{5}$ | $3.58\times10^{5}$ | $2.48\times10^{5}$ | BCAP31 (ER) |
| $6.56\times10^{2}$ | $2.48\times10^{4}$ | $4.29\times10^{4}$ | $7.68\times10^{4}$ | $6.52\times10^{4}$ | $8.22\times10^{4}$ | BICD2 (C/NU/CS/GA) |
| $5.83\times10^{4}$ | $2.38\times10^{5}$ | $5.10\times10^{5}$ | $8.75\times10^{5}$ | $7.80\times10^{5}$ | $8.82\times10^{5}$ | BID (C/MT) |
| $7.54\times10^{5}$ | $8.38\times10^{5}$ | $7.28\times10^{5}$ | $1.86\times10^{6}$ | $1.26\times10^{6}$ | $4.43\times10^{5}$ | BLMH (C) |
| $6.09\times10^{4}$ | $7.13\times10^{4}$ | $2.56\times10^{5}$ | $2.94\times10^{5}$ | $2.30\times10^{5}$ | $2.16\times10^{5}$ | BOLA1 (MT) |
| $1.49\times10^{6}$ | $1.58\times10^{6}$ | $6.39\times10^{6}$ | $5.92\times10^{6}$ | $4.42\times10^{6}$ | $6.05\times10^{6}$ | BOLA2B (C/NU) |
| $2.41\times10^{4}$ | $4.46\times10^{4}$ | $2.15\times10^{5}$ | $2.83\times10^{5}$ | $2.66\times10^{5}$ | $2.44\times10^{5}$ | C11orf54 (NU) |
| $6.56\times10^{4}$ | $1.61\times10^{5}$ | $1.60\times10^{6}$ | $2.56\times10^{6}$ | $1.46\times10^{6}$ | $2.04\times10^{6}$ | C11orf58 (???) |
| $2.26\times10^{3}$ | $5.12\times10^{3}$ | $9.30\times10^{4}$ | $2.11\times10^{5}$ | $1.70\times10^{5}$ | $1.30\times10^{5}$ | C12orf29 (???) |
| 0 | 0 | 0 | 0 | $4.62\times10^{2}$ | $1.68\times10^{3}$ | C1orf174 (NU) |
| $1.40\times10^{4}$ | $1.07\times10^{4}$ | $1.77\times10^{4}$ | $2.85\times10^{4}$ | $3.25\times10^{4}$ | $2.29\times10^{4}$ | C1orf50 (???) |
| $8.12\times10^{3}$ | $1.10\times10^{4}$ | $1.57\times10^{5}$ | $3.50\times10^{5}$ | $3.08\times10^{5}$ | $4.12\times10^{5}$ | C1orf52 (NU) |
| $1.10\times10^{6}$ | $1.65\times10^{6}$ | $4.12\times10^{6}$ | $7.07\times10^{6}$ | $6.00\times10^{6}$ | $5.70\times10^{6}$ | CACYBP (C/NU) |
| $1.20\times10^{5}$ | $1.80\times10^{5}$ | $3.83\times10^{5}$ | $9.92\times10^{5}$ | $7.25\times10^{5}$ | $4.65\times10^{5}$ | CALD1 (CS) |
| $1.82\times10^{4}$ | $2.87\times10^{4}$ | $2.92\times10^{4}$ | $4.41\times10^{4}$ | $3.57\times10^{4}$ | $3.70\times10^{4}$ | CAPNS1 (C/PM) |
| $5.97\times10^{4}$ | $1.23\times10^{5}$ | $1.33\times10^{5}$ | $2.29\times10^{5}$ | $2.46\times10^{5}$ | $1.83\times10^{5}$ | CAPZA2 (C/CS/EC) |
| $2.23\times10^{6}$ | $2.63\times10^{6}$ | $2.91\times10^{6}$ | $5.09\times10^{6}$ | $3.92\times10^{6}$ | $2.86\times10^{6}$ | CAPZB (CS) |
| $7.13\times10^{5}$ | $7.93\times10^{5}$ | $1.12\times10^{6}$ | $1.94\times10^{6}$ | $1.54\times10^{6}$ | $1.71\times10^{6}$ | CARHSP1 (C) |
| $2.25\times10^{5}$ | $3.27\times10^{5}$ | $5.20\times10^{5}$ | $7.18\times10^{5}$ | $7.13\times10^{5}$ | $6.64\times10^{5}$ | CASP3 (C) |
| $4.57\times10^{5}$ | $1.15\times10^{6}$ | $7.44\times10^{6}$ | $1.15\times10^{7}$ | $9.50\times10^{6}$ | $8.93\times10^{6}$ | CAST (C/ER) |

**Table S6**
**(page 2)**

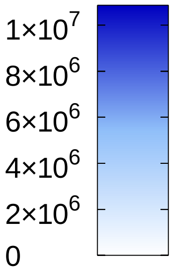


raw abundance

**external-$H_2O_2$ / c-YAP1C / 10 min**

| | 0 | 10 | 30 | 100 | 300 | 1000 |
|---|---|---|---|---|---|---|
| CCDC50 (C) | $2.23\times10^{4}$ | $1.07\times10^{5}$ | $3.57\times10^{5}$ | $5.79\times10^{5}$ | $4.42\times10^{5}$ | $4.19\times10^{5}$ |
| CCDC6 (C/CS) | $3.23\times10^{5}$ | $4.35\times10^{5}$ | $1.56\times10^{6}$ | $3.23\times10^{6}$ | $2.59\times10^{6}$ | $2.33\times10^{6}$ |
| CCS (C) | $2.11\times10^{5}$ | $2.46\times10^{5}$ | $5.50\times10^{5}$ | $1.16\times10^{6}$ | $1.22\times10^{6}$ | $1.32\times10^{6}$ |
| CDK4 (C/NU) | $8.32\times10^{5}$ | $2.45\times10^{6}$ | $4.23\times10^{6}$ | $6.51\times10^{6}$ | $5.60\times10^{6}$ | $4.79\times10^{6}$ |
| CDK6 (C/NU/CS) | $2.17\times10^{4}$ | $2.92\times10^{4}$ | $1.21\times10^{5}$ | $2.29\times10^{5}$ | $1.69\times10^{5}$ | $1.62\times10^{5}$ |
| CDKN2A (C/NU) | $3.64\times10^{5}$ | $7.59\times10^{5}$ | $1.42\times10^{6}$ | $1.86\times10^{6}$ | $1.80\times10^{6}$ | $1.48\times10^{6}$ |
| CHAC2 (C) | $1.57\times10^{4}$ | $4.75\times10^{4}$ | $9.70\times10^{4}$ | $1.75\times10^{5}$ | $1.43\times10^{5}$ | $1.68\times10^{5}$ |
| CHCHD4 (MT) | $4.34\times10^{5}$ | $4.95\times10^{5}$ | $9.87\times10^{5}$ | $1.15\times10^{6}$ | $8.28\times10^{5}$ | $9.70\times10^{5}$ |
| CHMP2B (C/ES) | $2.08\times10^{4}$ | $4.21\times10^{4}$ | $2.97\times10^{4}$ | $5.26\times10^{4}$ | $4.40\times10^{4}$ | $3.85\times10^{4}$ |
| CHMP5 (C/ES) | $5.56\times10^{4}$ | $1.08\times10^{5}$ | $3.24\times10^{5}$ | $9.57\times10^{5}$ | $8.02\times10^{5}$ | $5.96\times10^{5}$ |
| CHORDC1 (???) | $2.45\times10^{6}$ | $3.12\times10^{6}$ | $3.48\times10^{6}$ | $5.17\times10^{6}$ | $2.85\times10^{6}$ | $1.67\times10^{6}$ |
| CIAO2A (C/NU) | $3.49\times10^{4}$ | $3.93\times10^{4}$ | $5.66\times10^{4}$ | $8.79\times10^{4}$ | $9.06\times10^{4}$ | $6.60\times10^{4}$ |
| CIAPIN1 (C/NU/MT) | $1.22\times10^{6}$ | $3.55\times10^{6}$ | $1.69\times10^{7}$ | $2.68\times10^{7}$ | $2.45\times10^{7}$ | $2.61\times10^{7}$ |
| CKMT1A (MT) | $3.28\times10^{3}$ | $2.14\times10^{3}$ | $4.71\times10^{3}$ | $1.61\times10^{4}$ | $3.21\times10^{4}$ | $7.01\times10^{4}$ |
| CLIC1 (C/NU/PM) | $6.07\times10^{4}$ | $6.67\times10^{4}$ | $6.96\times10^{4}$ | $1.27\times10^{5}$ | $1.50\times10^{5}$ | $1.35\times10^{5}$ |
| CLU (C) | $1.18\times10^{5}$ | $1.37\times10^{5}$ | $1.17\times10^{5}$ | $1.48\times10^{6}$ | $3.79\times10^{5}$ | $6.04\times10^{5}$ |
| CNN3 (C/CS) | $1.09\times10^{6}$ | $1.75\times10^{6}$ | $1.63\times10^{6}$ | $3.01\times10^{6}$ | $2.63\times10^{6}$ | $2.10\times10^{6}$ |
| COX17 (C/MT) | $1.14\times10^{5}$ | $4.71\times10^{4}$ | $4.09\times10^{5}$ | $1.95\times10^{5}$ | $9.25\times10^{4}$ | $2.62\times10^{5}$ |
| CPOX (MT) | $5.09\times10^{5}$ | $1.01\times10^{6}$ | $2.05\times10^{6}$ | $3.52\times10^{6}$ | $3.98\times10^{6}$ | $5.88\times10^{6}$ |
| CPPED1 (C) | $1.57\times10^{3}$ | $2.62\times10^{4}$ | $8.86\times10^{4}$ | $1.22\times10^{5}$ | $1.32\times10^{5}$ | $1.79\times10^{5}$ |
| CRKL (C/NU/PM) | $1.37\times10^{6}$ | $1.94\times10^{6}$ | $1.08\times10^{7}$ | $1.81\times10^{7}$ | $1.67\times10^{7}$ | $1.60\times10^{7}$ |
| CRMP1 (C/CS) | $1.78\times10^{4}$ | $8.33\times10^{4}$ | $2.12\times10^{5}$ | $3.70\times10^{5}$ | $3.76\times10^{5}$ | $2.92\times10^{5}$ |
| CRYZL1 (C) | $1.77\times10^{4}$ | $7.77\times10^{4}$ | $1.46\times10^{5}$ | $2.00\times10^{5}$ | $1.94\times10^{5}$ | $1.88\times10^{5}$ |
| CSTB (C/NU) | $7.85\times10^{5}$ | $1.71\times10^{6}$ | $1.33\times10^{7}$ | $1.18\times10^{7}$ | $8.26\times10^{6}$ | $1.57\times10^{7}$ |
| CTH (C) | $3.39\times10^{4}$ | $4.89\times10^{4}$ | $2.35\times10^{5}$ | $5.99\times10^{5}$ | $6.67\times10^{5}$ | $8.13\times10^{5}$ |
| CTSB (LY/EC) | $4.07\times10^{6}$ | $4.80\times10^{6}$ | $1.09\times10^{7}$ | $8.74\times10^{6}$ | $5.96\times10^{6}$ | $6.65\times10^{6}$ |
| CWC15 (NU) | $2.00\times10^{0}$ | $1.07\times10^{3}$ | $1.76\times10^{4}$ | $5.18\times10^{4}$ | $3.84\times10^{4}$ | $5.18\times10^{4}$ |
| CYB5R3 (C/MT/ER) | $1.01\times10^{5}$ | $1.06\times10^{5}$ | $1.15\times10^{5}$ | $2.33\times10^{5}$ | $2.09\times10^{5}$ | $1.52\times10^{5}$ |
| DBN1 (C) | $3.39\times10^{5}$ | $3.34\times10^{5}$ | $3.34\times10^{5}$ | $5.63\times10^{5}$ | $6.37\times10^{5}$ | $7.13\times10^{5}$ |
| DBNL (C/GA/PM/EE/CS) | $2.49\times10^{3}$ | $5.37\times10^{3}$ | $7.98\times10^{3}$ | $1.44\times10^{4}$ | $1.18\times10^{4}$ | $9.20\times10^{3}$ |
| DCPS (C/NU) | $3.24\times10^{4}$ | $2.91\times10^{4}$ | $1.83\times10^{5}$ | $2.88\times10^{5}$ | $2.64\times10^{5}$ | $2.83\times10^{5}$ |
| DCTPP1 (C/NU/MT) | $3.38\times10^{5}$ | $8.84\times10^{5}$ | $4.17\times10^{6}$ | $6.63\times10^{6}$ | $6.44\times10^{6}$ | $5.53\times10^{6}$ |
| DENR (???) | $2.28\times10^{4}$ | $5.83\times10^{4}$ | $1.01\times10^{5}$ | $3.03\times10^{5}$ | $2.89\times10^{5}$ | $3.12\times10^{5}$ |
| DFFA (C) | $2.25\times10^{5}$ | $3.95\times10^{5}$ | $5.05\times10^{5}$ | $9.70\times10^{5}$ | $8.98\times10^{5}$ | $6.40\times10^{5}$ |
| DHPS (C) | $4.01\times10^{4}$ | $6.66\times10^{4}$ | $3.10\times10^{5}$ | $4.32\times10^{5}$ | $3.72\times10^{5}$ | $4.55\times10^{5}$ |
| DLAT (MT) | $3.12\times10^{3}$ | $6.05\times10^{3}$ | $6.11\times10^{3}$ | $2.91\times10^{4}$ | $1.81\times10^{4}$ | $1.88\times10^{4}$ |
| DLST (NU/MT) | $1.27\times10^{5}$ | $1.47\times10^{5}$ | $1.92\times10^{5}$ | $2.91\times10^{5}$ | $2.61\times10^{5}$ | $2.10\times10^{5}$ |
| DLST (NU/MT) | 0 | 0 | 0 | 0 | 0 | $8.20\times10^{1}$ |
| DNAJB11 (ER) | $5.91\times10^{6}$ | $1.15\times10^{7}$ | $5.82\times10^{6}$ | $1.25\times10^{7}$ | $1.11\times10^{7}$ | $6.94\times10^{6}$ |
| DNAJC9 (C/NU/PM) | $1.92\times10^{4}$ | $2.45\times10^{4}$ | $2.37\times10^{4}$ | $4.48\times10^{4}$ | $4.62\times10^{4}$ | $2.46\times10^{4}$ |
| DNPEP (C) | $1.39\times10^{5}$ | $5.96\times10^{5}$ | $2.69\times10^{6}$ | $5.39\times10^{6}$ | $6.09\times10^{6}$ | $7.63\times10^{6}$ |
| DPYSL2 (C/CS/PM) | $5.64\times10^{5}$ | $1.06\times10^{6}$ | $2.75\times10^{6}$ | $4.98\times10^{6}$ | $4.30\times10^{6}$ | $3.37\times10^{6}$ |
| DPYSL3 (C) | $7.76\times10^{5}$ | $2.13\times10^{6}$ | $5.57\times10^{6}$ | $1.06\times10^{7}$ | $9.16\times10^{6}$ | $7.32\times10^{6}$ |
| DPYSL5 (C) | $7.38\times10^{4}$ | $9.79\times10^{4}$ | $2.04\times10^{5}$ | $3.84\times10^{5}$ | $3.84\times10^{5}$ | $2.33\times10^{5}$ |
| DRG1 (C/NU) | $1.21\times10^{3}$ | $3.22\times10^{2}$ | $4.01\times10^{2}$ | $2.88\times10^{3}$ | $3.46\times10^{3}$ | $1.99\times10^{3}$ |
| DSTN (C/CS/EC) | $2.07\times10^{5}$ | $2.22\times10^{5}$ | $3.39\times10^{5}$ | $5.09\times10^{5}$ | $4.40\times10^{5}$ | $4.03\times10^{5}$ |
| DUSP3 (NU) | $2.90\times10^{4}$ | $4.77\times10^{4}$ | $4.23\times10^{4}$ | $1.33\times10^{5}$ | $2.02\times10^{5}$ | $2.73\times10^{5}$ |
| DYNLL1 (NU/CS/MT) | $1.40\times10^{5}$ | $2.50\times10^{5}$ | $7.81\times10^{5}$ | $1.16\times10^{6}$ | $7.20\times10^{5}$ | $7.76\times10^{5}$ |
| EFHD2 (PM) | $1.26\times10^{5}$ | $3.38\times10^{5}$ | $8.54\times10^{5}$ | $1.38\times10^{6}$ | $1.20\times10^{6}$ | $1.35\times10^{6}$ |
| EIF1AD (NU) | $1.11\times10^{4}$ | $1.78\times10^{4}$ | $1.06\times10^{5}$ | $1.66\times10^{5}$ | $1.76\times10^{5}$ | $1.97\times10^{5}$ |

### Table S6 (page 3)

#### external-$H_2O_2$ / c-YAP1C / 10 min

raw abundance: $1\times10^{7}$, $8\times10^{6}$, $6\times10^{6}$, $4\times10^{6}$, $2\times10^{6}$, 0

| 0 | 10 | 30 | 100 | 300 | 1000 | |
|---|---|---|---|---|---|---|
| $1.24\times10^{3}$ | $2.88\times10^{3}$ | $3.16\times10^{3}$ | $7.98\times10^{3}$ | $1.17\times10^{4}$ | $4.71\times10^{3}$ | EIF3B (C) |
| $2.76\times10^{5}$ | $6.63\times10^{5}$ | $3.61\times10^{5}$ | $7.32\times10^{5}$ | $7.21\times10^{5}$ | $5.70\times10^{5}$ | EIF3G (C/NU) |
| $5.30\times10^{4}$ | $8.59\times10^{4}$ | $7.04\times10^{4}$ | $1.14\times10^{5}$ | $1.33\times10^{5}$ | $1.09\times10^{5}$ | EIF3I (C) |
| $3.20\times10^{1}$ | $7.20\times10^{1}$ | $7.40\times10^{1}$ | $5.00\times10^{0}$ | $8.60\times10^{1}$ | $2.40\times10^{1}$ | EIF4A2 (C) |
| $2.23\times10^{4}$ | $5.02\times10^{4}$ | $2.43\times10^{5}$ | $4.85\times10^{5}$ | $4.22\times10^{5}$ | $5.76\times10^{5}$ | EIF4B (C) |
| $8.03\times10^{3}$ | $4.79\times10^{4}$ | $1.86\times10^{5}$ | $1.22\times10^{5}$ | $6.46\times10^{4}$ | $6.74\times10^{4}$ | EIF4EBP1 (C/NU) |
| $1.82\times10^{6}$ | $1.83\times10^{6}$ | $1.90\times10^{6}$ | $4.98\times10^{6}$ | $4.01\times10^{6}$ | $3.38\times10^{6}$ | EIF5A (C/NU/ER) |
| $4.95\times10^{5}$ | $4.49\times10^{5}$ | $9.37\times10^{5}$ | $1.19\times10^{6}$ | $9.23\times10^{5}$ | $9.24\times10^{5}$ | EIF6 (C/NU) |
| $4.55\times10^{5}$ | $5.61\times10^{5}$ | $5.67\times10^{5}$ | $9.87\times10^{5}$ | $8.38\times10^{5}$ | $5.58\times10^{5}$ | EPDR1 (LY/EC) |
| $4.41\times10^{4}$ | $8.83\times10^{4}$ | $8.57\times10^{4}$ | $1.82\times10^{5}$ | $1.52\times10^{5}$ | $1.55\times10^{5}$ | EPS15L1 (NU/PM) |
| $8.55\times10^{4}$ | $2.58\times10^{4}$ | $4.77\times10^{5}$ | $6.99\times10^{5}$ | $6.46\times10^{5}$ | $7.32\times10^{5}$ | ERICH5 (???) |
| $2.04\times10^{4}$ | $2.87\times10^{4}$ | $5.55\times10^{4}$ | $7.58\times10^{4}$ | $6.33\times10^{4}$ | $8.17\times10^{4}$ | EVPL (CS) |
| $9.03\times10^{4}$ | $3.01\times10^{5}$ | $6.04\times10^{5}$ | $9.12\times10^{5}$ | $6.05\times10^{5}$ | $5.05\times10^{5}$ | EWSR1 (C/NU/PM) |
| $3.17\times10^{3}$ | $5.29\times10^{3}$ | $6.64\times10^{3}$ | $5.56\times10^{3}$ | $3.09\times10^{3}$ | $3.55\times10^{3}$ | EXOSC6 (C/NU) |
| $1.90\times10^{3}$ | $7.17\times10^{3}$ | $6.34\times10^{4}$ | $8.50\times10^{4}$ | $7.73\times10^{4}$ | $1.09\times10^{5}$ | FAH (C/EC) |
| $8.94\times10^{3}$ | $1.55\times10^{4}$ | $1.15\times10^{4}$ | $1.87\times10^{4}$ | $1.57\times10^{4}$ | $1.33\times10^{4}$ | FAM234A (PM) |
| $2.50\times10^{4}$ | $5.20\times10^{4}$ | $1.50\times10^{4}$ | $3.19\times10^{4}$ | $7.36\times10^{4}$ | $3.35\times10^{4}$ | FKBP3 (NU) |
| $8.02\times10^{5}$ | $1.39\times10^{6}$ | $7.36\times10^{5}$ | $1.05\times10^{6}$ | $1.81\times10^{6}$ | $3.06\times10^{5}$ | FLG (C/PM) |
| $1.12\times10^{4}$ | $4.33\times10^{4}$ | $5.18\times10^{4}$ | $7.47\times10^{4}$ | $6.59\times10^{4}$ | $7.99\times10^{4}$ | FLYWCH2 (???) |
| $2.15\times10^{5}$ | $5.79\times10^{5}$ | $1.29\times10^{6}$ | $2.09\times10^{6}$ | $1.68\times10^{6}$ | $1.69\times10^{6}$ | FN3KRP (C) |
| $2.00\times10^{4}$ | $5.40\times10^{4}$ | $2.15\times10^{5}$ | $3.62\times10^{5}$ | $2.65\times10^{5}$ | $2.56\times10^{5}$ | FNTA (C/PM/CS) |
| $1.58\times10^{4}$ | $3.48\times10^{4}$ | $1.52\times10^{5}$ | $2.26\times10^{5}$ | $1.95\times10^{5}$ | $2.23\times10^{5}$ | FNTB (C/CS) |
| $1.59\times10^{6}$ | $2.10\times10^{6}$ | $2.72\times10^{6}$ | $4.37\times10^{6}$ | $3.64\times10^{6}$ | $2.44\times10^{6}$ | FSCN1 (C/CS) |
| $2.66\times10^{4}$ | $6.97\times10^{4}$ | $2.52\times10^{5}$ | $3.39\times10^{5}$ | $2.94\times10^{5}$ | $3.41\times10^{5}$ | FTH1 (C/NU/LY/EC) |
| $2.66\times10^{5}$ | $3.05\times10^{5}$ | $3.83\times10^{5}$ | $6.80\times10^{5}$ | $5.58\times10^{5}$ | $5.29\times10^{5}$ | GARS1 (C/EC) |
| $5.26\times10^{4}$ | $1.76\times10^{5}$ | $3.07\times10^{5}$ | $5.25\times10^{5}$ | $4.56\times10^{5}$ | $4.46\times10^{5}$ | GCLC (C) |
| $8.30\times10^{4}$ | $3.20\times10^{5}$ | $9.84\times10^{5}$ | $1.62\times10^{6}$ | $1.20\times10^{6}$ | $1.08\times10^{6}$ | GCLM (C) |
| $1.40\times10^{5}$ | $2.15\times10^{5}$ | $2.37\times10^{5}$ | $5.03\times10^{5}$ | $4.11\times10^{5}$ | $3.73\times10^{5}$ | GEMIN6 (C/NU) |
| $7.74\times10^{4}$ | $1.73\times10^{5}$ | $3.90\times10^{5}$ | $7.52\times10^{5}$ | $6.56\times10^{5}$ | $9.19\times10^{5}$ | GFER (C/MT/EC) |
| $4.39\times10^{3}$ | $9.36\times10^{3}$ | $2.19\times10^{4}$ | $6.31\times10^{4}$ | $6.36\times10^{4}$ | $6.21\times10^{4}$ | GLDC (MT) |
| $1.23\times10^{5}$ | $4.26\times10^{5}$ | $1.26\times10^{6}$ | $1.72\times10^{6}$ | $1.49\times10^{6}$ | $1.69\times10^{6}$ | GLOD4 (MT) |
| $2.83\times10^{4}$ | $4.08\times10^{4}$ | $9.66\times10^{4}$ | $1.72\times10^{5}$ | $1.65\times10^{5}$ | $1.64\times10^{5}$ | GLRX3 (C) |
| $1.85\times10^{3}$ | $1.03\times10^{3}$ | $1.24\times10^{3}$ | $6.75\times10^{3}$ | $3.84\times10^{3}$ | $1.39\times10^{4}$ | GPC4 (PM/EC) |
| $4.78\times10^{5}$ | $1.83\times10^{6}$ | $6.00\times10^{6}$ | $9.97\times10^{6}$ | $8.63\times10^{6}$ | $7.96\times10^{6}$ | GPHN (C/PM) |
| $4.78\times10^{4}$ | $7.60\times10^{4}$ | $1.43\times10^{5}$ | $2.74\times10^{5}$ | $2.43\times10^{5}$ | $2.65\times10^{5}$ | GPX1 (C) |
| $4.60\times10^{5}$ | $8.03\times10^{5}$ | $1.71\times10^{6}$ | $2.51\times10^{6}$ | $2.27\times10^{6}$ | $2.23\times10^{6}$ | GPX4 (C/MT) |
| $1.22\times10^{6}$ | $3.72\times10^{5}$ | $1.83\times10^{6}$ | $2.76\times10^{6}$ | $2.43\times10^{6}$ | $2.50\times10^{6}$ | GRB2 (C/MT/EE/GA) |
| $5.70\times10^{4}$ | $8.14\times10^{4}$ | $2.35\times10^{5}$ | $5.27\times10^{5}$ | $4.50\times10^{5}$ | $3.27\times10^{5}$ | GRIPAP1 (ES) |
| $4.16\times10^{3}$ | $4.15\times10^{4}$ | $5.49\times10^{3}$ | $1.28\times10^{4}$ | $1.32\times10^{4}$ | $1.34\times10^{4}$ | GRN (LY/EC) |
| $3.44\times10^{5}$ | $3.91\times10^{5}$ | $3.78\times10^{5}$ | $7.10\times10^{5}$ | $5.09\times10^{5}$ | $5.67\times10^{5}$ | GRPEL1 (MT) |
| $1.78\times10^{6}$ | $6.10\times10^{6}$ | $3.36\times10^{7}$ | $4.93\times10^{7}$ | $4.52\times10^{7}$ | $4.07\times10^{7}$ | GSR (C/MT) |
| $5.80\times10^{1}$ | $4.04\times10^{2}$ | $3.66\times10^{3}$ | $4.01\times10^{3}$ | $5.76\times10^{3}$ | $6.89\times10^{3}$ | H2AC21 (NU) |
| $3.23\times10^{4}$ | $7.83\times10^{4}$ | $8.91\times10^{4}$ | $1.42\times10^{5}$ | $1.47\times10^{5}$ | $1.27\times10^{5}$ | H2BC11 (NU) |
| $7.09\times10^{3}$ | $8.46\times10^{3}$ | $1.04\times10^{4}$ | $1.61\times10^{4}$ | $1.61\times10^{4}$ | $1.55\times10^{4}$ | HAGH (C/MT) |
| $6.95\times10^{4}$ | $8.05\times10^{4}$ | $1.96\times10^{5}$ | $2.87\times10^{5}$ | $2.64\times10^{5}$ | $2.30\times10^{5}$ | HAT1 (C/NU/MT) |
| $1.69\times10^{5}$ | $1.43\times10^{4}$ | $1.01\times10^{4}$ | $2.19\times10^{4}$ | $4.90\times10^{4}$ | $3.91\times10^{6}$ | HBA2 (C/EC) |
| $1.80\times10^{5}$ | $3.03\times10^{4}$ | $2.85\times10^{4}$ | $5.28\times10^{4}$ | $8.15\times10^{4}$ | $7.85\times10^{5}$ | HBB (C/EC) |
| $3.97\times10^{4}$ | $9.84\times10^{4}$ | $1.86\times10^{5}$ | $3.89\times10^{5}$ | $2.50\times10^{5}$ | $2.37\times10^{5}$ | HEXIM1 (C/NU) |
| $1.16\times10^{4}$ | $9.45\times10^{3}$ | $3.09\times10^{4}$ | $6.46\times10^{4}$ | $4.74\times10^{4}$ | $4.05\times10^{4}$ | HIRIP3 (NU) |
| $8.54\times10^{3}$ | $1.58\times10^{4}$ | $1.60\times10^{4}$ | $3.12\times10^{4}$ | $2.17\times10^{4}$ | $2.01\times10^{4}$ | HMGA1 (NU) |

### Table S6 (page 4)

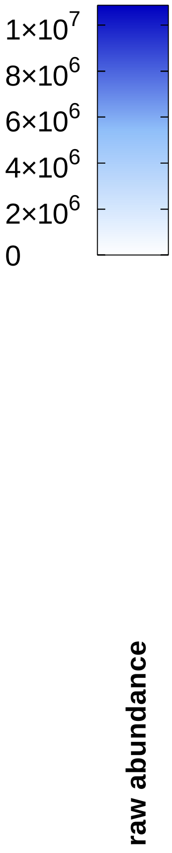


**external-$H_2O_2$ / c-YAP1C / 10 min**

| | 0 | 10 | 30 | 100 | 300 | 1000 |
|---|---|---|---|---|---|---|
| HMGCS1 (C) | $2.21\times10^{5}$ | $2.78\times10^{5}$ | $3.85\times10^{5}$ | $6.05\times10^{5}$ | $4.58\times10^{5}$ | $3.26\times10^{5}$ |
| HPRT1 (C) | $1.08\times10^{6}$ | $4.49\times10^{6}$ | $1.50\times10^{7}$ | $2.24\times10^{7}$ | $2.36\times10^{7}$ | $2.78\times10^{7}$ |
| HSD17B10 (MT) | $1.73\times10^{6}$ | $2.87\times10^{6}$ | $1.07\times10^{7}$ | $2.33\times10^{7}$ | $2.24\times10^{7}$ | $2.39\times10^{7}$ |
| HSD17B8 (MT) | $2.17\times10^{3}$ | $1.04\times10^{4}$ | $5.63\times10^{4}$ | $9.99\times10^{4}$ | $1.10\times10^{5}$ | $1.12\times10^{5}$ |
| HSPA4L (C/NU) | $1.74\times10^{5}$ | $1.54\times10^{5}$ | $2.65\times10^{5}$ | $4.47\times10^{5}$ | $3.95\times10^{5}$ | $3.92\times10^{5}$ |
| HSPH1 (C) | $5.58\times10^{5}$ | $6.59\times10^{5}$ | $7.86\times10^{5}$ | $1.24\times10^{6}$ | $1.19\times10^{6}$ | $1.09\times10^{6}$ |
| IAH1 (???) | $9.71\times10^{4}$ | $1.38\times10^{5}$ | $2.73\times10^{5}$ | $4.28\times10^{5}$ | $3.93\times10^{5}$ | $5.47\times10^{5}$ |
| IBA57 (MT) | 0 | $2.34\times10^{4}$ | $5.96\times10^{4}$ | $1.28\times10^{5}$ | $1.27\times10^{5}$ | $1.34\times10^{5}$ |
| IDI1 (C/PO) | $2.97\times10^{4}$ | $3.95\times10^{4}$ | $2.88\times10^{4}$ | $5.69\times10^{4}$ | $7.37\times10^{4}$ | $4.32\times10^{4}$ |
| IKBKG (C/NU) | $2.05\times10^{4}$ | $2.16\times10^{4}$ | $3.58\times10^{4}$ | $7.24\times10^{4}$ | $4.94\times10^{4}$ | $4.24\times10^{4}$ |
| IMPDH1 (C/NU) | 0 | $1.65\times10^{4}$ | $4.29\times10^{4}$ | $1.13\times10^{5}$ | $8.91\times10^{4}$ | $1.34\times10^{5}$ |
| IMPDH2 (C/NU) | $1.51\times10^{5}$ | $2.98\times10^{5}$ | $5.93\times10^{5}$ | $1.22\times10^{6}$ | $1.08\times10^{6}$ | $1.21\times10^{6}$ |
| ISCA2 (MT) | $1.13\times10^{5}$ | $2.23\times10^{5}$ | $4.01\times10^{5}$ | $9.51\times10^{5}$ | $9.23\times10^{5}$ | $6.56\times10^{5}$ |
| ISCU (C/NU/MT) | $6.77\times10^{4}$ | $1.20\times10^{5}$ | $2.87\times10^{5}$ | $7.78\times10^{5}$ | $6.41\times10^{5}$ | $9.01\times10^{5}$ |
| ISYNA1 (C) | $3.83\times10^{6}$ | $5.59\times10^{6}$ | $2.00\times10^{7}$ | $2.75\times10^{7}$ | $2.69\times10^{7}$ | $3.04\times10^{7}$ |
| JMJD6 (C/NU) | $4.02\times10^{4}$ | $5.24\times10^{4}$ | $1.64\times10^{5}$ | $2.41\times10^{5}$ | $1.78\times10^{5}$ | $1.71\times10^{5}$ |
| JPT2 (C/NU) | $2.72\times10^{5}$ | $1.52\times10^{6}$ | $8.25\times10^{6}$ | $8.58\times10^{6}$ | $4.95\times10^{6}$ | $8.15\times10^{6}$ |
| KCTD5 (C/NU) | $1.06\times10^{3}$ | $1.05\times10^{4}$ | $3.93\times10^{4}$ | $6.88\times10^{4}$ | $5.08\times10^{4}$ | $5.02\times10^{4}$ |
| KHDRBS1 (C/NU/PM) | $2.47\times10^{4}$ | $1.96\times10^{4}$ | $4.04\times10^{4}$ | $6.75\times10^{4}$ | $4.66\times10^{4}$ | $1.81\times10^{4}$ |
| KHSRP (C/NU) | $2.49\times10^{4}$ | $7.49\times10^{4}$ | $1.29\times10^{5}$ | $2.66\times10^{5}$ | $2.23\times10^{5}$ | $2.42\times10^{5}$ |
| KIF3A (CS) | $4.26\times10^{4}$ | $7.01\times10^{4}$ | $1.04\times10^{5}$ | $1.39\times10^{5}$ | $1.41\times10^{5}$ | $1.22\times10^{5}$ |
| LARP7 (NU) | $6.35\times10^{3}$ | $9.50\times10^{3}$ | $5.05\times10^{4}$ | $9.99\times10^{4}$ | $9.43\times10^{4}$ | $1.06\times10^{5}$ |
| LBH (C/NU) | $9.69\times10^{3}$ | $3.38\times10^{4}$ | $2.01\times10^{5}$ | $2.19\times10^{5}$ | $1.70\times10^{5}$ | $1.15\times10^{5}$ |
| LCP1 (PM/CS) | $8.04\times10^{5}$ | $7.36\times10^{5}$ | $2.99\times10^{6}$ | $4.78\times10^{6}$ | $4.09\times10^{6}$ | $4.12\times10^{6}$ |
| LRRFIP1 (C/NU) | $2.50\times10^{4}$ | $6.44\times10^{4}$ | $1.58\times10^{5}$ | $2.93\times10^{5}$ | $3.08\times10^{5}$ | $3.11\times10^{5}$ |
| LTF (C/EC) | $2.70\times10^{3}$ | $1.04\times10^{4}$ | $3.02\times10^{3}$ | $5.30\times10^{3}$ | $1.54\times10^{4}$ | $1.04\times10^{4}$ |
| LUC7L2(NU) | $7.79\times10^{4}$ | $1.62\times10^{5}$ | $1.51\times10^{5}$ | $2.29\times10^{5}$ | $1.99\times10^{5}$ | $1.98\times10^{5}$ |
| LYRM7 (MT) | $3.20\times10^{4}$ | $1.27\times10^{5}$ | $1.87\times10^{5}$ | $4.65\times10^{5}$ | $2.96\times10^{5}$ | $3.51\times10^{5}$ |
| MAD2L1(C/CS/NU) | $1.33\times10^{5}$ | $2.25\times10^{5}$ | $2.29\times10^{5}$ | $3.07\times10^{5}$ | $2.79\times10^{5}$ | $3.27\times10^{5}$ |
| MALSU1 (MT) | $3.23\times10^{5}$ | $3.94\times10^{5}$ | $4.67\times10^{5}$ | $6.46\times10^{5}$ | $6.58\times10^{5}$ | $6.31\times10^{5}$ |
| MAP4 (CS) | $3.07\times10^{5}$ | $3.67\times10^{5}$ | $1.25\times10^{6}$ | $2.21\times10^{6}$ | $1.89\times10^{6}$ | $1.84\times10^{6}$ |
| MAPK1 (C/CS/NU) | $2.30\times10^{3}$ | $7.30\times10^{3}$ | $1.61\times10^{4}$ | $3.41\times10^{4}$ | $3.59\times10^{4}$ | $2.88\times10^{4}$ |
| MAPK14 (C/NU) | $2.00\times10^{4}$ | $1.88\times10^{4}$ | $1.07\times10^{5}$ | $2.27\times10^{5}$ | $1.60\times10^{5}$ | $1.40\times10^{5}$ |
| MARCKS (CS/PM) | $5.64\times10^{5}$ | $1.53\times10^{6}$ | $2.78\times10^{6}$ | $6.35\times10^{6}$ | $5.02\times10^{6}$ | $4.21\times10^{6}$ |
| MARCKSL1 (CS/PM) | $6.71\times10^{4}$ | $3.29\times10^{5}$ | $1.39\times10^{6}$ | $3.46\times10^{6}$ | $2.64\times10^{6}$ | $2.84\times10^{6}$ |
| MCMBP (NU) | $3.65\times10^{3}$ | $1.72\times10^{4}$ | $4.51\times10^{4}$ | $9.06\times10^{4}$ | $7.36\times10^{4}$ | $1.54\times10^{5}$ |
| MDH1 (C) | $2.13\times10^{5}$ | $2.22\times10^{5}$ | $7.48\times10^{5}$ | $1.16\times10^{6}$ | $9.44\times10^{5}$ | $9.73\times10^{5}$ |
| METTL26 (???) | $4.24\times10^{3}$ | $2.91\times10^{4}$ | $4.61\times10^{4}$ | $6.86\times10^{4}$ | $5.47\times10^{4}$ | $4.03\times10^{4}$ |
| MINDY3 (NU) | $1.11\times10^{5}$ | $9.48\times10^{4}$ | $2.77\times10^{5}$ | $4.53\times10^{5}$ | $5.25\times10^{5}$ | $4.17\times10^{5}$ |
| MMAB (MT) | $2.41\times10^{4}$ | $3.23\times10^{4}$ | $6.75\times10^{4}$ | $1.29\times10^{5}$ | $1.41\times10^{5}$ | $1.58\times10^{5}$ |
| MOCS2 (C) | $3.50\times10^{4}$ | $7.29\times10^{4}$ | $2.24\times10^{5}$ | $3.96\times10^{5}$ | $3.38\times10^{5}$ | $3.18\times10^{5}$ |
| MPP1 (PM) | $1.57\times10^{4}$ | $5.81\times10^{4}$ | $1.31\times10^{5}$ | $2.83\times10^{5}$ | $2.38\times10^{5}$ | $2.26\times10^{5}$ |
| MRPL1 (MT) | $3.06\times10^{3}$ | $4.19\times10^{3}$ | $3.13\times10^{3}$ | $6.19\times10^{3}$ | $2.84\times10^{3}$ | $3.64\times10^{3}$ |
| MRPL2 (MT) | 0 | 0 | $6.94\times10^{2}$ | $7.72\times10^{2}$ | $2.55\times10^{3}$ | $3.67\times10^{2}$ |
| MSRB3 (MT/ER) | $4.93\times10^{3}$ | $3.93\times10^{4}$ | $9.31\times10^{4}$ | $1.53\times10^{5}$ | $1.37\times10^{5}$ | $1.36\times10^{5}$ |
| MTAP (C/NU) | $6.72\times10^{4}$ | $1.22\times10^{5}$ | $3.62\times10^{5}$ | $6.06\times10^{5}$ | $6.36\times10^{5}$ | $5.07\times10^{5}$ |
| MTHFD2 (MT) | $4.26\times10^{4}$ | $4.22\times10^{4}$ | $4.51\times10^{4}$ | $9.23\times10^{4}$ | $5.60\times10^{4}$ | $5.05\times10^{4}$ |
| MYL6 (C/CS/EC) | $2.70\times10^{5}$ | $4.00\times10^{5}$ | $2.88\times10^{5}$ | $6.30\times10^{5}$ | $6.31\times10^{5}$ | $5.48\times10^{5}$ |
| NAA10 (C/NU) | $9.19\times10^{4}$ | $1.57\times10^{5}$ | $1.83\times10^{5}$ | $4.18\times10^{5}$ | $2.94\times10^{5}$ | $2.88\times10^{5}$ |
| NAP1L4 (C/NU) | $1.93\times10^{4}$ | $3.60\times10^{4}$ | $2.62\times10^{4}$ | $4.10\times10^{4}$ | $4.47\times10^{4}$ | $3.93\times10^{4}$ |

#### Table S6
#### (page 5)

##### external-$H_2O_2$ / c-YAP1C / 10 min

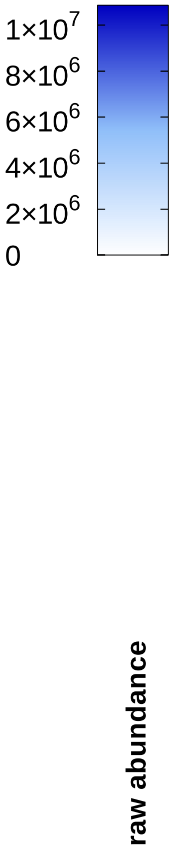


| | 0 | 10 | 30 | 100 | 300 | 1000 |
|---|---|---|---|---|---|---|
| NASP (C/NU) | $3.01\times10^{6}$ | $3.64\times10^{6}$ | $4.82\times10^{6}$ | $7.34\times10^{6}$ | $6.77\times10^{6}$ | $6.24\times10^{6}$ |
| NDUFAB1 (MT) | $1.93\times10^{5}$ | $2.14\times10^{5}$ | $6.51\times10^{5}$ | $1.05\times10^{6}$ | $8.34\times10^{5}$ | $6.78\times10^{5}$ |
| NDUFS6 (MT) | $3.64\times10^{5}$ | $4.30\times10^{5}$ | $7.67\times10^{5}$ | $9.89\times10^{5}$ | $7.64\times10^{5}$ | $9.48\times10^{5}$ |
| NDUFV3 (MT) | $5.62\times10^{4}$ | $7.05\times10^{4}$ | $9.53\times10^{4}$ | $2.35\times10^{5}$ | $1.15\times10^{5}$ | $1.11\times10^{5}$ |
| NECAP2 (PM) | $2.78\times10^{5}$ | $5.13\times10^{5}$ | $2.25\times10^{5}$ | $5.57\times10^{5}$ | $5.70\times10^{5}$ | $3.02\times10^{5}$ |
| NFU1 (C/MT) | $4.95\times10^{5}$ | $7.11\times10^{5}$ | $3.63\times10^{6}$ | $9.03\times10^{6}$ | $7.79\times10^{6}$ | $7.47\times10^{6}$ |
| NOMO1 (ER) | $8.12\times10^{4}$ | $1.20\times10^{5}$ | $1.00\times10^{5}$ | $1.63\times10^{5}$ | $1.43\times10^{5}$ | $1.16\times10^{5}$ |
| NPEPPS (C/NU) | $2.30\times10^{6}$ | $2.22\times10^{6}$ | $6.09\times10^{6}$ | $1.00\times10^{7}$ | $8.79\times10^{6}$ | $7.85\times10^{6}$ |
| NRDC (C/MT) | $1.18\times10^{5}$ | $1.56\times10^{5}$ | $1.83\times10^{5}$ | $2.94\times10^{5}$ | $2.98\times10^{5}$ | $2.85\times10^{5}$ |
| NT5DC2 (???) | $1.38\times10^{4}$ | $3.17\times10^{4}$ | $1.41\times10^{5}$ | $4.56\times10^{5}$ | $3.68\times10^{5}$ | $3.54\times10^{5}$ |
| NUDC (CS/NU) | $8.35\times10^{5}$ | $4.71\times10^{6}$ | $1.67\times10^{7}$ | $2.75\times10^{7}$ | $2.37\times10^{7}$ | $2.54\times10^{7}$ |
| NUDT1 (C/NU/MT) | $8.99\times10^{3}$ | $3.04\times10^{4}$ | $2.54\times10^{4}$ | $3.40\times10^{4}$ | $3.57\times10^{4}$ | $4.73\times10^{4}$ |
| OGFR (C/NU) | $2.22\times10^{4}$ | $8.22\times10^{4}$ | $8.94\times10^{5}$ | $1.37\times10^{6}$ | $1.20\times10^{6}$ | $1.25\times10^{6}$ |
| OPTN (C/GA/RE) | $9.16\times10^{3}$ | $6.42\times10^{4}$ | $9.36\times10^{4}$ | $1.68\times10^{5}$ | $1.49\times10^{5}$ | $1.36\times10^{5}$ |
| OTUB1 (C) | $2.63\times10^{4}$ | $3.43\times10^{4}$ | $2.94\times10^{4}$ | $1.90\times10^{5}$ | $4.49\times10^{5}$ | $1.22\times10^{6}$ |
| P3H1 (ER/EC) | $8.32\times10^{4}$ | $1.19\times10^{5}$ | $1.03\times10^{5}$ | $1.85\times10^{5}$ | $1.41\times10^{5}$ | $1.45\times10^{5}$ |
| PA2G4 (C/NU) | $2.76\times10^{5}$ | $2.98\times10^{5}$ | $3.36\times10^{5}$ | $5.91\times10^{5}$ | $4.99\times10^{5}$ | $5.26\times10^{5}$ |
| PABIR2 (???) | $9.81\times10^{4}$ | $1.14\times10^{5}$ | $1.51\times10^{5}$ | $2.60\times10^{5}$ | $2.27\times10^{5}$ | $1.89\times10^{5}$ |
| PABPC1 (C/NU) | $7.78\times10^{4}$ | $1.12\times10^{5}$ | $1.97\times10^{5}$ | $3.44\times10^{5}$ | $3.35\times10^{5}$ | $4.05\times10^{5}$ |
| PABPC4 (C) | $3.89\times10^{3}$ | $4.10\times10^{4}$ | $9.59\times10^{4}$ | $1.94\times10^{5}$ | $2.00\times10^{5}$ | $2.10\times10^{5}$ |
| PAICS (C/EC) | $6.90\times10^{5}$ | $1.50\times10^{6}$ | $2.90\times10^{6}$ | $6.35\times10^{6}$ | $5.51\times10^{6}$ | $3.34\times10^{6}$ |
| PAWR (C/NU) | $6.29\times10^{4}$ | $1.36\times10^{5}$ | $3.43\times10^{5}$ | $7.97\times10^{5}$ | $7.31\times10^{5}$ | $6.45\times10^{5}$ |
| PBK (NU) | $1.71\times10^{6}$ | $1.62\times10^{6}$ | $3.52\times10^{6}$ | $5.71\times10^{6}$ | $5.33\times10^{6}$ | $4.81\times10^{6}$ |
| PCBP1 (C/NU) | $9.18\times10^{4}$ | $1.67\times10^{5}$ | $6.29\times10^{5}$ | $1.45\times10^{6}$ | $1.23\times10^{6}$ | $9.36\times10^{5}$ |
| PCMT1 (C) | $3.01\times10^{6}$ | $5.48\times10^{6}$ | $1.31\times10^{7}$ | $2.01\times10^{7}$ | $1.95\times10^{7}$ | $1.79\times10^{7}$ |
| PDCD6IP (C/CS/EC) | $8.67\times10^{3}$ | $1.55\times10^{4}$ | $3.55\times10^{4}$ | $7.53\times10^{4}$ | $5.32\times10^{4}$ | $4.04\times10^{4}$ |
| PDHB (MT) | $2.48\times10^{5}$ | $1.89\times10^{5}$ | $2.74\times10^{5}$ | $7.02\times10^{5}$ | $9.47\times10^{5}$ | $1.14\times10^{6}$ |
| PDIA3 (ER) | $6.54\times10^{6}$ | $1.17\times10^{7}$ | $1.07\times10^{7}$ | $1.76\times10^{7}$ | $1.50\times10^{7}$ | $1.11\times10^{7}$ |
| PDIA6 (ER/PM) | $1.73\times10^{6}$ | $2.80\times10^{6}$ | $2.16\times10^{6}$ | $3.49\times10^{6}$ | $3.25\times10^{6}$ | $2.58\times10^{6}$ |
| PDLIM1 (C/CS) | $7.37\times10^{5}$ | $1.08\times10^{6}$ | $7.46\times10^{5}$ | $2.00\times10^{6}$ | $1.50\times10^{6}$ | $1.12\times10^{6}$ |
| PFDN1 (C) | $3.29\times10^{4}$ | $2.88\times10^{4}$ | $5.75\times10^{4}$ | $8.80\times10^{4}$ | $6.43\times10^{4}$ | $8.12\times10^{4}$ |
| PFDN2 (C/NU/MT) | $1.51\times10^{5}$ | $2.97\times10^{5}$ | $1.23\times10^{6}$ | $1.70\times10^{6}$ | $1.47\times10^{6}$ | $1.89\times10^{6}$ |
| PFDN4 (C/NU/MT) | $6.33\times10^{5}$ | $1.26\times10^{6}$ | $3.54\times10^{6}$ | $5.06\times10^{6}$ | $5.05\times10^{6}$ | $4.73\times10^{6}$ |
| PFDN5 (C/NU) | $8.60\times10^{4}$ | $1.58\times10^{5}$ | $7.46\times10^{5}$ | $1.02\times10^{6}$ | $8.04\times10^{5}$ | $1.10\times10^{6}$ |
| PFN2 (CS) | $5.22\times10^{4}$ | $2.33\times10^{5}$ | $9.19\times10^{5}$ | $9.45\times10^{5}$ | $1.10\times10^{6}$ | $1.11\times10^{6}$ |
| PGLS (C) | $2.21\times10^{5}$ | $2.56\times10^{5}$ | $5.83\times10^{5}$ | $7.14\times10^{5}$ | $7.33\times10^{5}$ | $6.08\times10^{5}$ |
| PHAX (C/NU) | $2.41\times10^{4}$ | $6.80\times10^{4}$ | $5.20\times10^{5}$ | $1.17\times10^{6}$ | $1.24\times10^{6}$ | $1.18\times10^{6}$ |
| PIN1 (C/NU) | $1.51\times10^{4}$ | $8.05\times10^{3}$ | $1.36\times10^{4}$ | $5.31\times10^{4}$ | $1.17\times10^{5}$ | $2.09\times10^{5}$ |
| PITHD1 (C) | $2.44\times10^{5}$ | $1.10\times10^{6}$ | $1.02\times10^{7}$ | $1.53\times10^{7}$ | $1.28\times10^{7}$ | $1.10\times10^{7}$ |
| PITRM1 (MT) | $8.84\times10^{3}$ | $1.26\times10^{4}$ | $3.16\times10^{4}$ | $6.85\times10^{4}$ | $7.69\times10^{4}$ | $1.04\times10^{5}$ |
| PLS3 (C) | $1.29\times10^{6}$ | $1.88\times10^{6}$ | $5.96\times10^{6}$ | $9.55\times10^{6}$ | $8.07\times10^{6}$ | $8.51\times10^{6}$ |
| PM20D2 (NU) | $7.12\times10^{5}$ | $9.36\times10^{5}$ | $7.04\times10^{5}$ | $1.51\times10^{6}$ | $1.33\times10^{6}$ | $1.16\times10^{6}$ |
| PMPCB (MT) | $3.40\times10^{3}$ | $3.56\times10^{4}$ | $2.83\times10^{4}$ | $8.58\times10^{4}$ | $4.15\times10^{4}$ | $4.23\times10^{4}$ |
| PNPO (C) | $2.44\times10^{5}$ | $5.38\times10^{5}$ | $2.04\times10^{6}$ | $2.87\times10^{6}$ | $2.52\times10^{6}$ | $2.78\times10^{6}$ |
| POLDIP2 (NU/MT) | $2.84\times10^{4}$ | $4.65\times10^{4}$ | $6.87\times10^{4}$ | $1.64\times10^{5}$ | $1.24\times10^{5}$ | $1.43\times10^{5}$ |
| POMP (C/NU/ER) | $7.52\times10^{4}$ | $1.55\times10^{5}$ | $6.54\times10^{4}$ | $1.23\times10^{5}$ | $1.58\times10^{5}$ | $9.56\times10^{4}$ |
| PON2 (PM) | $4.54\times10^{4}$ | $8.50\times10^{4}$ | $5.67\times10^{4}$ | $1.08\times10^{5}$ | $8.22\times10^{4}$ | $6.88\times10^{4}$ |
| PPAT (C) | $1.21\times10^{4}$ | $9.33\times10^{4}$ | $3.25\times10^{5}$ | $5.00\times10^{5}$ | $4.07\times10^{5}$ | $7.18\times10^{5}$ |
| PPCDC (C) | $2.48\times10^{4}$ | $1.38\times10^{5}$ | $6.25\times10^{5}$ | $8.42\times10^{5}$ | $7.51\times10^{5}$ | $8.02\times10^{5}$ |
| PPL (C/CS/PM) | $2.15\times10^{2}$ | 0 | $4.40\times10^{1}$ | 0 | 0 | $9.10\times10^{2}$ |

### Table S6 (page 6)

**external-$H_2O_2$ / c-YAP1C / 10 min**

raw abundance (colour scale): $1\times10^{7}$, $8\times10^{6}$, $6\times10^{6}$, $4\times10^{6}$, $2\times10^{6}$, 0

| | 0 | 10 | 30 | 100 | 300 | 1000 |
|---|---|---|---|---|---|---|
| PPP1CC (C/NU/MT) | $1.45\times10^{3}$ | $1.08\times10^{3}$ | $2.19\times10^{3}$ | $6.84\times10^{3}$ | $4.02\times10^{3}$ | $4.61\times10^{3}$ |
| PPP1R11 (C/NU) | $3.99\times10^{5}$ | $1.17\times10^{6}$ | $6.23\times10^{6}$ | $7.75\times10^{6}$ | $6.00\times10^{6}$ | $6.75\times10^{6}$ |
| PPP1R2 (???) | 0 | 0 | $4.68\times10^{4}$ | $9.10\times10^{4}$ | $4.77\times10^{4}$ | $1.08\times10^{5}$ |
| PPP1R7 (NU) | $3.08\times10^{4}$ | $4.52\times10^{4}$ | $1.55\times10^{5}$ | $2.06\times10^{5}$ | $2.03\times10^{5}$ | $1.83\times10^{5}$ |
| PPP2R2D (C) | $2.43\times10^{3}$ | $2.82\times10^{4}$ | $1.72\times10^{5}$ | $2.58\times10^{5}$ | $2.35\times10^{5}$ | $2.30\times10^{5}$ |
| PRDX1 (C) | $5.42\times10^{7}$ | $2.36\times10^{8}$ | $7.01\times10^{8}$ | $8.78\times10^{8}$ | $7.59\times10^{8}$ | $6.49\times10^{8}$ |
| PRDX2 (C) | $1.33\times10^{7}$ | $3.88\times10^{7}$ | $9.88\times10^{7}$ | $1.16\times10^{8}$ | $9.88\times10^{7}$ | $6.24\times10^{7}$ |
| PRDX3 (MT/C/EE) | $1.08\times10^{7}$ | $2.45\times10^{7}$ | $5.44\times10^{7}$ | $9.06\times10^{7}$ | $6.27\times10^{7}$ | $3.53\times10^{7}$ |
| PRDX4 (C/ER) | $6.50\times10^{6}$ | $1.06\times10^{7}$ | $1.74\times10^{7}$ | $2.82\times10^{7}$ | $2.42\times10^{7}$ | $2.08\times10^{7}$ |
| PRDX5 (C/NU/MT/PO) | $1.32\times10^{6}$ | $3.00\times10^{6}$ | $7.06\times10^{6}$ | $1.43\times10^{7}$ | $1.41\times10^{7}$ | $1.14\times10^{7}$ |
| PRDX6 (C/LY) | $3.09\times10^{6}$ | $8.12\times10^{6}$ | $1.57\times10^{7}$ | $3.43\times10^{7}$ | $3.24\times10^{7}$ | $2.66\times10^{7}$ |
| PREP (C) | $5.98\times10^{4}$ | $1.26\times10^{5}$ | $3.03\times10^{5}$ | $4.40\times10^{5}$ | $5.06\times10^{5}$ | $5.35\times10^{5}$ |
| PRMT1 (C/NU) | $2.87\times10^{4}$ | $3.44\times10^{4}$ | $4.39\times10^{4}$ | $7.10\times10^{4}$ | $7.36\times10^{4}$ | $5.69\times10^{4}$ |
| PRMT5 (C/NU/GA) | $7.69\times10^{5}$ | $2.34\times10^{6}$ | $5.15\times10^{6}$ | $8.08\times10^{6}$ | $7.43\times10^{6}$ | $8.12\times10^{6}$ |
| PSMD9 (C/NU) | $7.59\times10^{5}$ | $3.87\times10^{5}$ | $1.75\times10^{6}$ | $2.52\times10^{6}$ | $2.02\times10^{6}$ | $2.11\times10^{6}$ |
| PSME3 (C/NU) | $8.71\times10^{5}$ | $1.18\times10^{6}$ | $3.65\times10^{6}$ | $5.49\times10^{6}$ | $5.06\times10^{6}$ | $4.23\times10^{6}$ |
| PSME3IP1 (NU) | $1.04\times10^{4}$ | $6.48\times10^{4}$ | $2.53\times10^{5}$ | $4.04\times10^{5}$ | $3.57\times10^{5}$ | $3.31\times10^{5}$ |
| PSMG4 (C) | $5.51\times10^{4}$ | $1.13\times10^{5}$ | $5.05\times10^{4}$ | $1.19\times10^{5}$ | $1.55\times10^{5}$ | $1.03\times10^{5}$ |
| PXN (C/CS) | $1.97\times10^{5}$ | $3.14\times10^{5}$ | $2.30\times10^{5}$ | $4.11\times10^{5}$ | $3.07\times10^{5}$ | $2.10\times10^{5}$ |
| RAB21 (GA/ES/ER) | $2.38\times10^{4}$ | $2.46\times10^{4}$ | $2.94\times10^{4}$ | $5.08\times10^{4}$ | $3.18\times10^{4}$ | $2.59\times10^{4}$ |
| RAB8A (C/PM/ES/CS/GA) | $1.35\times10^{3}$ | $1.45\times10^{3}$ | $1.12\times10^{3}$ | $4.30\times10^{3}$ | $3.42\times10^{3}$ | $3.12\times10^{3}$ |
| RABEP2 (C/CS/ES) | $1.43\times10^{5}$ | $3.26\times10^{5}$ | $5.51\times10^{5}$ | $8.69\times10^{5}$ | $8.00\times10^{5}$ | $6.98\times10^{5}$ |
| RANBP3 (C/NU) | $6.60\times10^{3}$ | $1.14\times10^{4}$ | $1.51\times10^{5}$ | $2.67\times10^{5}$ | $2.17\times10^{5}$ | $2.41\times10^{5}$ |
| RBBP7 (NU) | $6.02\times10^{5}$ | $1.33\times10^{6}$ | $3.86\times10^{6}$ | $5.30\times10^{6}$ | $4.43\times10^{6}$ | $4.47\times10^{6}$ |
| RBM8A (C/NU) | $1.14\times10^{5}$ | $1.43\times10^{5}$ | $3.44\times10^{5}$ | $5.51\times10^{5}$ | $5.06\times10^{5}$ | $5.40\times10^{5}$ |
| RBMX (NU) | $1.07\times10^{5}$ | $1.39\times10^{5}$ | $1.95\times10^{5}$ | $2.73\times10^{5}$ | $2.14\times10^{5}$ | $1.96\times10^{5}$ |
| RCC2 (NU/PM/CS) | $1.20\times10^{5}$ | $2.19\times10^{5}$ | $8.51\times10^{5}$ | $1.42\times10^{6}$ | $1.38\times10^{6}$ | $1.59\times10^{6}$ |
| RILPL1 (C/CS) | $1.64\times10^{3}$ | $2.29\times10^{4}$ | $6.19\times10^{4}$ | $9.33\times10^{4}$ | $8.36\times10^{4}$ | $1.32\times10^{5}$ |
| RNASEH2C (NU) | $1.77\times10^{5}$ | $1.99\times10^{5}$ | $1.87\times10^{5}$ | $3.59\times10^{5}$ | $2.96\times10^{5}$ | $2.32\times10^{5}$ |
| RNPS1 (C/NU) | $3.06\times10^{4}$ | $3.36\times10^{4}$ | $4.81\times10^{4}$ | $7.86\times10^{4}$ | $9.04\times10^{4}$ | $5.81\times10^{4}$ |
| RPIA (C) | $1.70\times10^{4}$ | $3.84\times10^{4}$ | $1.58\times10^{5}$ | $2.61\times10^{5}$ | $2.59\times10^{5}$ | $2.67\times10^{5}$ |
| RPL15 (PM) | $9.08\times10^{4}$ | $1.57\times10^{5}$ | $1.16\times10^{5}$ | $1.73\times10^{5}$ | $1.47\times10^{5}$ | $2.25\times10^{5}$ |
| RPL27 (C/ER) | $2.60\times10^{4}$ | $3.76\times10^{4}$ | $6.81\times10^{4}$ | $1.56\times10^{5}$ | $8.26\times10^{4}$ | $7.59\times10^{4}$ |
| RPL35A (C/EC) | $8.49\times10^{3}$ | $5.97\times10^{3}$ | $1.09\times10^{4}$ | $2.21\times10^{4}$ | $1.60\times10^{4}$ | $1.50\times10^{4}$ |
| RPL36A (C) | $5.95\times10^{3}$ | $1.23\times10^{4}$ | $9.80\times10^{3}$ | $1.69\times10^{4}$ | $1.65\times10^{4}$ | $8.63\times10^{3}$ |
| RPL6 (C/ER) | $6.84\times10^{4}$ | $1.14\times10^{5}$ | $7.55\times10^{4}$ | $7.31\times10^{4}$ | $1.41\times10^{5}$ | $5.28\times10^{4}$ |
| RPS27 (C/NU) | $4.22\times10^{4}$ | $3.85\times10^{4}$ | $2.14\times10^{4}$ | $5.14\times10^{4}$ | $3.41\times10^{4}$ | $9.14\times10^{4}$ |
| RPS27A (C/NU) | $9.70\times10^{6}$ | $1.14\times10^{7}$ | $1.46\times10^{7}$ | $2.38\times10^{7}$ | $2.35\times10^{7}$ | $2.23\times10^{7}$ |
| RRM2 (C) | $3.23\times10^{5}$ | $4.46\times10^{5}$ | $1.08\times10^{6}$ | $3.34\times10^{6}$ | $3.57\times10^{6}$ | $4.12\times10^{6}$ |
| RRM2B (C/NU) | $8.38\times10^{4}$ | $2.06\times10^{5}$ | $2.53\times10^{5}$ | $5.80\times10^{5}$ | $4.57\times10^{5}$ | $4.91\times10^{5}$ |
| S100A9 (C/CS/PM/EC) | $1.55\times10^{5}$ | $1.69\times10^{5}$ | $3.84\times10^{5}$ | $1.78\times10^{5}$ | $2.41\times10^{5}$ | $5.30\times10^{5}$ |
| SAP30BP (NU) | $6.59\times10^{4}$ | $9.11\times10^{4}$ | $5.49\times10^{4}$ | $1.41\times10^{5}$ | $1.13\times10^{5}$ | $7.29\times10^{4}$ |
| SCO2 (MT) | $1.52\times10^{5}$ | $2.48\times10^{5}$ | $2.19\times10^{5}$ | $3.28\times10^{5}$ | $3.08\times10^{5}$ | $2.86\times10^{5}$ |
| SDE2 (NU) | $9.01\times10^{3}$ | $2.79\times10^{4}$ | $6.41\times10^{4}$ | $1.95\times10^{5}$ | $1.69\times10^{5}$ | $1.20\times10^{5}$ |
| SEC22B (GA/ER) | $3.29\times10^{4}$ | $4.74\times10^{4}$ | $3.91\times10^{4}$ | $1.03\times10^{5}$ | $6.17\times10^{4}$ | $5.18\times10^{4}$ |
| SELENBP1 (C/NU) | $1.51\times10^{5}$ | $1.96\times10^{5}$ | $8.68\times10^{5}$ | $1.36\times10^{6}$ | $1.24\times10^{6}$ | $1.30\times10^{6}$ |
| SERPINA1 (ER/EC) | $3.03\times10^{4}$ | $3.20\times10^{4}$ | $4.41\times10^{4}$ | $7.16\times10^{4}$ | $6.03\times10^{4}$ | $1.32\times10^{6}$ |
| SERPINB1 (C/LY/ES/EC) | $2.14\times10^{5}$ | $4.49\times10^{5}$ | $2.25\times10^{6}$ | $3.58\times10^{6}$ | $3.22\times10^{6}$ | $3.22\times10^{6}$ |
| SERPINB6 (C) | $1.50\times10^{5}$ | $1.49\times10^{5}$ | $2.16\times10^{5}$ | $3.50\times10^{5}$ | $2.69\times10^{5}$ | $2.51\times10^{5}$ |
| SFN (C/NU/EC) | $1.26\times10^{4}$ | $1.10\times10^{4}$ | $3.96\times10^{4}$ | $6.15\times10^{4}$ | $4.48\times10^{4}$ | $4.30\times10^{4}$ |

### Table S6 (page 7)

external-$H_2O_2$ / c-YAP1C / 10 min

raw abundance (color scale: 0, $2\times10^6$, $4\times10^6$, $6\times10^6$, $8\times10^6$, $1\times10^7$)

| | 0 | 10 | 30 | 100 | 300 | 1000 |
|---|---|---|---|---|---|---|
| SH3BP5 (MT/VE) | 0 | $3.49\times10^2$ | $9.67\times10^3$ | $1.39\times10^4$ | $1.28\times10^4$ | $1.38\times10^4$ |
| SHTN1 (CS) | $2.31\times10^5$ | $4.27\times10^5$ | $9.99\times10^5$ | $2.58\times10^6$ | $2.52\times10^6$ | $2.52\times10^6$ |
| SKP1 (C/NU) | $1.74\times10^6$ | $5.86\times10^6$ | $3.86\times10^7$ | $5.45\times10^7$ | $4.38\times10^7$ | $3.14\times10^7$ |
| SLC9A3R (NU/PM) | $1.44\times10^3$ | $4.27\times10^3$ | $2.75\times10^4$ | $8.55\times10^4$ | $6.62\times10^4$ | $6.11\times10^4$ |
| SMAP1 (PM) | $1.55\times10^4$ | $9.95\times10^4$ | $2.57\times10^5$ | $8.67\times10^5$ | $7.04\times10^5$ | $5.30\times10^5$ |
| SNAP23 (PM) | $2.84\times10^5$ | $5.72\times10^5$ | $6.21\times10^5$ | $1.33\times10^6$ | $1.07\times10^6$ | $9.36\times10^5$ |
| SNAPIN (C/LY/GA/ES) | $5.83\times10^2$ | $6.22\times10^3$ | $1.23\times10^4$ | $2.82\times10^4$ | $2.23\times10^4$ | $2.51\times10^4$ |
| SNU13 (NU) | $1.88\times10^4$ | $3.02\times10^4$ | $6.35\times10^4$ | $1.31\times10^5$ | $1.20\times10^5$ | $1.39\times10^5$ |
| SNUPN (C/NU) | $5.27\times10^4$ | $1.38\times10^5$ | $7.47\times10^4$ | $1.45\times10^5$ | $1.29\times10^5$ | $1.07\times10^5$ |
| SOD1 (C/NU/MT) | $1.36\times10^6$ | $1.16\times10^6$ | $1.14\times10^6$ | $2.08\times10^6$ | $2.37\times10^6$ | $3.21\times10^6$ |
| SOD2 (MT) | $7.95\times10^3$ | $5.16\times10^3$ | $3.23\times10^4$ | $5.16\times10^4$ | $4.95\times10^4$ | $5.77\times10^4$ |
| SORBS3 (NU/CS) | $3.67\times10^4$ | $4.57\times10^4$ | $1.24\times10^5$ | $2.11\times10^5$ | $1.73\times10^5$ | $2.01\times10^5$ |
| SPTAN1 (CS/PM) | $1.26\times10^5$ | $1.46\times10^5$ | $1.64\times10^5$ | $3.84\times10^5$ | $3.61\times10^5$ | $3.68\times10^5$ |
| SRSF10 (C/NU) | $1.26\times10^4$ | $2.06\times10^4$ | $4.05\times10^4$ | $5.89\times10^4$ | $7.83\times10^4$ | $9.29\times10^4$ |
| SRXN1 (C) | $7.04\times10^4$ | $1.46\times10^5$ | $3.11\times10^5$ | $5.58\times10^5$ | $5.08\times10^5$ | $4.47\times10^5$ |
| SSB (NU) | $3.15\times10^5$ | $4.08\times10^5$ | $4.21\times10^5$ | $6.97\times10^5$ | $6.99\times10^5$ | $6.92\times10^5$ |
| ST13 (C) | $3.82\times10^5$ | $6.09\times10^5$ | $4.65\times10^5$ | $8.81\times10^5$ | $9.78\times10^5$ | $8.01\times10^5$ |
| STIP1 (C/NU) | $1.13\times10^6$ | $1.40\times10^6$ | $1.11\times10^6$ | $2.29\times10^6$ | $1.86\times10^6$ | $1.66\times10^6$ |
| STRAP (C/NU) | $1.29\times10^6$ | $1.83\times10^6$ | $4.69\times10^6$ | $7.15\times10^6$ | $6.00\times10^6$ | $5.60\times10^6$ |
| SUMF1 (ER) | $1.21\times10^5$ | $2.34\times10^5$ | $1.93\times10^5$ | $3.06\times10^5$ | $2.34\times10^5$ | $2.93\times10^5$ |
| SUMF2 (ER) | $2.41\times10^2$ | $8.37\times10^2$ | $2.37\times10^3$ | $5.77\times10^3$ | $2.89\times10^3$ | $9.12\times10^2$ |
| SUPT5H (NU) | $3.04\times10^5$ | $4.40\times10^5$ | $3.40\times10^5$ | $6.07\times10^5$ | $5.27\times10^5$ | $4.04\times10^5$ |
| TACC3 (C/CS) | $8.35\times10^4$ | $1.34\times10^5$ | $4.54\times10^5$ | $1.11\times10^6$ | $9.02\times10^5$ | $6.95\times10^5$ |
| TAF15 (C/NU) | 0 | $4.51\times10^2$ | $3.97\times10^2$ | 0 | $7.71\times10^2$ | $1.16\times10^4$ |
| TBCA (CS) | $3.37\times10^4$ | $2.49\times10^4$ | $1.25\times10^5$ | $8.57\times10^4$ | $6.73\times10^4$ | $8.29\times10^4$ |
| TCEAL4 (NU) | $5.38\times10^3$ | $2.04\times10^4$ | $2.97\times10^4$ | $8.55\times10^4$ | $7.45\times10^4$ | $5.62\times10^4$ |
| THOP1 (C) | $2.10\times10^5$ | $3.71\times10^5$ | $1.02\times10^6$ | $1.67\times10^6$ | $1.71\times10^6$ | $1.94\times10^6$ |
| TIMM13 (MT) | $1.83\times10^4$ | $2.07\times10^4$ | $9.20\times10^3$ | $3.90\times10^4$ | $2.70\times10^4$ | $1.91\times10^4$ |
| TIPRL (C) | $7.45\times10^5$ | $8.97\times10^5$ | $1.48\times10^6$ | $2.25\times10^6$ | $1.75\times10^6$ | $1.75\times10^6$ |
| TK1 (C) | $2.82\times10^4$ | $3.89\times10^3$ | $8.72\times10^4$ | $2.87\times10^5$ | $2.04\times10^5$ | $1.44\times10^5$ |
| TMA7 (???) | $2.17\times10^2$ | $1.58\times10^3$ | $4.82\times10^2$ | $2.39\times10^3$ | $1.33\times10^3$ | $1.47\times10^3$ |
| TPM1 (CS) | $2.94\times10^4$ | $8.74\times10^4$ | $2.05\times10^5$ | $2.48\times10^5$ | $2.39\times10^5$ | $2.70\times10^5$ |
| TPM3 (CS) | $1.02\times10^6$ | $6.62\times10^6$ | $2.07\times10^7$ | $2.60\times10^7$ | $2.06\times10^7$ | $1.88\times10^7$ |
| TPM4 (CS) | $1.83\times10^5$ | $9.34\times10^5$ | $3.23\times10^6$ | $4.63\times10^6$ | $5.02\times10^6$ | $4.40\times10^6$ |
| TPRKB (C/NU) | $5.91\times10^4$ | $9.23\times10^4$ | $8.46\times10^4$ | $1.38\times10^5$ | $1.34\times10^5$ | $1.14\times10^5$ |
| TSN (C/NU) | $1.42\times10^5$ | $2.60\times10^5$ | $8.72\times10^5$ | $1.30\times10^6$ | $1.16\times10^6$ | $1.15\times10^6$ |
| TSNAX (NU/GA/C) | $5.68\times10^4$ | $7.58\times10^4$ | $1.35\times10^5$ | $2.48\times10^5$ | $2.31\times10^5$ | $1.96\times10^5$ |
| TSR2 (???) | $4.22\times10^2$ | $2.48\times10^3$ | $2.86\times10^4$ | $1.99\times10^4$ | $4.24\times10^4$ | $3.00\times10^4$ |
| TTC1 (C/PO) | $6.10\times10^3$ | $1.68\times10^4$ | $8.03\times10^4$ | $1.66\times10^5$ | $1.44\times10^5$ | $1.27\times10^5$ |
| TTR (C/EC) | $1.01\times10^3$ | $7.33\times10^3$ | $4.51\times10^3$ | $3.23\times10^3$ | $7.92\times10^3$ | $9.62\times10^4$ |
| TUBA4A (CS) | $1.96\times10^4$ | $4.70\times10^3$ | $1.43\times10^5$ | $9.33\times10^3$ | $1.56\times10^4$ | $1.14\times10^4$ |
| TWF1 (C/CS) | $8.45\times10^4$ | $1.69\times10^5$ | $4.31\times10^5$ | $7.56\times10^5$ | $6.10\times10^5$ | $5.75\times10^5$ |
| TWF2 (C/CS) | $1.41\times10^5$ | $2.12\times10^5$ | $3.87\times10^5$ | $7.07\times10^5$ | $6.33\times10^5$ | $5.10\times10^5$ |
| TXLNA (C/EC) | 0 | 0 | 0 | $1.31\times10^3$ | $1.52\times10^3$ | $1.35\times10^3$ |
| TXN (C/NU/EC) | $4.97\times10^6$ | $3.36\times10^7$ | $1.39\times10^8$ | $1.68\times10^8$ | $1.59\times10^8$ | $1.38\times10^8$ |
| TXNDC17 (C) | $8.04\times10^4$ | $7.19\times10^5$ | $1.69\times10^6$ | $2.98\times10^6$ | $2.31\times10^6$ | $1.39\times10^6$ |
| TXNDC5 (ER) | $6.57\times10^5$ | $1.12\times10^6$ | $1.10\times10^6$ | $1.76\times10^6$ | $1.57\times10^6$ | $1.66\times10^6$ |
| TXNL1 (C/NU) | $4.91\times10^4$ | $3.88\times10^4$ | $8.70\times10^4$ | $1.32\times10^5$ | $1.38\times10^5$ | $1.29\times10^5$ |
| UBAP2 (C/NU) | $3.66\times10^2$ | $3.49\times10^3$ | $1.02\times10^3$ | $4.48\times10^3$ | $4.77\times10^3$ | $3.27\times10^3$ |
| UBE2A (C/NU) | $1.46\times10^5$ | $2.68\times10^5$ | $2.14\times10^6$ | $3.92\times10^6$ | $3.04\times10^6$ | $4.32\times10^6$ |

#### Table S6 (page 8)

##### external-$H_2O_2$ / c-YAP1C / 10 min

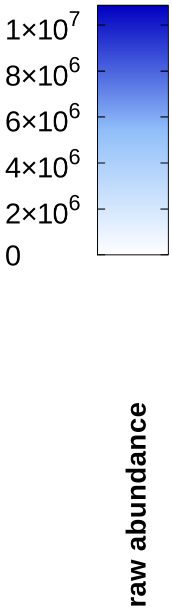


| | 0 | 10 | 30 | 100 | 300 | 1000 |
|---|---|---|---|---|---|---|
| UBE2B (NU/PM) | $1.18\times10^{5}$ | $1.90\times10^{5}$ | $1.09\times10^{6}$ | $2.12\times10^{6}$ | $2.34\times10^{6}$ | $2.31\times10^{6}$ |
| UBE2G1 (C/EC) | $7.85\times10^{4}$ | $6.89\times10^{4}$ | $6.63\times10^{4}$ | $1.11\times10^{5}$ | $1.63\times10^{5}$ | $1.96\times10^{5}$ |
| UBE2H (C/NU) | $2.27\times10^{4}$ | $1.54\times10^{4}$ | $2.38\times10^{4}$ | $5.81\times10^{4}$ | $9.52\times10^{4}$ | $1.70\times10^{5}$ |
| UBE2I (C/NU) | $1.42\times10^{5}$ | $1.59\times10^{5}$ | $2.27\times10^{5}$ | $8.17\times10^{5}$ | $1.27\times10^{6}$ | $3.21\times10^{6}$ |
| UBE2K (C) | $5.65\times10^{5}$ | $7.10\times10^{5}$ | $1.04\times10^{6}$ | $2.09\times10^{6}$ | $2.37\times10^{6}$ | $2.86\times10^{6}$ |
| UBE2L3 (C/NU) | $2.06\times10^{6}$ | $1.50\times10^{6}$ | $1.56\times10^{7}$ | $3.24\times10^{7}$ | $3.09\times10^{7}$ | $3.58\times10^{7}$ |
| UBE2N (C/NU) | $9.32\times10^{5}$ | $1.09\times10^{6}$ | $1.54\times10^{6}$ | $3.25\times10^{6}$ | $4.86\times10^{6}$ | $7.68\times10^{6}$ |
| UBFD1 (???) | $6.39\times10^{4}$ | $6.67\times10^{4}$ | $2.12\times10^{5}$ | $5.02\times10^{5}$ | $3.82\times10^{5}$ | $4.17\times10^{5}$ |
| UQCRC2 (MT) | $3.35\times10^{4}$ | $3.08\times10^{4}$ | $2.92\times10^{4}$ | $6.99\times10^{4}$ | $5.02\times10^{4}$ | $5.67\times10^{4}$ |
| UQCRH (MT) | $9.07\times10^{4}$ | $1.14\times10^{5}$ | $2.33\times10^{5}$ | $1.73\times10^{5}$ | $1.06\times10^{5}$ | $1.77\times10^{5}$ |
| USP5 (C/NU/LY) | $2.36\times10^{4}$ | $1.66\times10^{4}$ | $2.54\times10^{4}$ | $4.14\times10^{4}$ | $5.68\times10^{4}$ | $4.86\times10^{4}$ |
| VBP1 (C/NU) | $4.24\times10^{5}$ | $6.71\times10^{5}$ | $2.34\times10^{6}$ | $3.72\times10^{6}$ | $2.99\times10^{6}$ | $3.54\times10^{6}$ |
| VPS26B (C/ES) | $1.82\times10^{3}$ | $2.06\times10^{3}$ | $9.29\times10^{3}$ | $1.26\times10^{4}$ | $1.22\times10^{4}$ | $7.15\times10^{3}$ |
| WBP2 (C/NU) | $2.16\times10^{5}$ | $6.77\times10^{5}$ | $3.05\times10^{6}$ | $5.50\times10^{6}$ | $4.17\times10^{6}$ | $3.64\times10^{6}$ |
| WDR44 (C/GA/ES) | $1.65\times10^{5}$ | $1.17\times10^{5}$ | $3.58\times10^{5}$ | $5.70\times10^{5}$ | $5.40\times10^{5}$ | $6.77\times10^{5}$ |
| WDR5 (NU) | $3.67\times10^{5}$ | $4.54\times10^{5}$ | $8.92\times10^{5}$ | $1.42\times10^{6}$ | $1.36\times10^{6}$ | $1.04\times10^{6}$ |
| WDR77 (C/NU) | $3.94\times10^{5}$ | $9.41\times10^{5}$ | $2.34\times10^{6}$ | $3.78\times10^{6}$ | $3.25\times10^{6}$ | $3.52\times10^{6}$ |
| XRCC4 (NU) | $1.37\times10^{4}$ | $2.00\times10^{4}$ | $9.79\times10^{4}$ | $1.79\times10^{5}$ | $1.64\times10^{5}$ | $1.86\times10^{5}$ |
| YWHAE (C/NU) | $7.37\times10^{6}$ | $8.54\times10^{6}$ | $1.49\times10^{7}$ | $1.92\times10^{7}$ | $1.78\times10^{7}$ | $1.76\times10^{7}$ |
| YWHAG (C) | $6.56\times10^{5}$ | $9.69\times10^{5}$ | $1.31\times10^{6}$ | $1.73\times10^{6}$ | $1.52\times10^{6}$ | $1.65\times10^{6}$ |
| YWHAH (C/PM/MT/EC) | $1.95\times10^{5}$ | $1.96\times10^{5}$ | $2.87\times10^{5}$ | $4.20\times10^{5}$ | $2.97\times10^{5}$ | $2.83\times10^{5}$ |
| YWHAQ (C) | $1.95\times10^{6}$ | $3.48\times10^{6}$ | $9.87\times10^{6}$ | $1.26\times10^{7}$ | $1.12\times10^{7}$ | $1.18\times10^{7}$ |
| YWHAZ (C) | $3.70\times10^{6}$ | $5.20\times10^{6}$ | $7.74\times10^{6}$ | $9.93\times10^{6}$ | $9.20\times10^{6}$ | $8.96\times10^{6}$ |
| ZFAND5 (C) | 0 | 0 | 0 | $6.94\times10^{2}$ | 0 | 0 |
| ZFYVE16 (C/ES) | 0 | $3.07\times10^{2}$ | $3.45\times10^{3}$ | $1.02\times10^{4}$ | $8.47\times10^{3}$ | $5.35\times10^{3}$ |

#### Table S7
#### (page 1)

##### c-DD-DAO (10 min): no chase (NC) vs. chase (C)

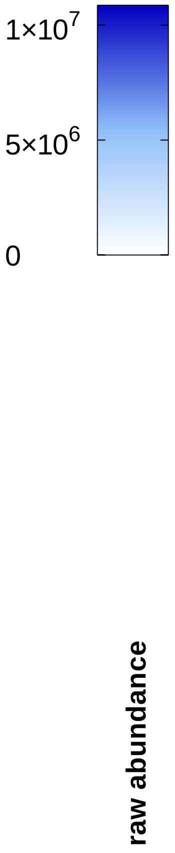


| NC (L-Ala) | NC (D-Ala) | C (L-Ala) | C (D-Ala) | |
|---|---|---|---|---|
| $1.59\times10^{5}$ | $4.55\times10^{5}$ | $2.34\times10^{5}$ | $2.08\times10^{5}$ | ACTC1 (CS) |
| $1.21\times10^{6}$ | $3.59\times10^{6}$ | $1.42\times10^{6}$ | $1.27\times10^{6}$ | ACTG1 (CS) |
| $1.47\times10^{4}$ | $1.11\times10^{5}$ | $2.60\times10^{4}$ | $3.69\times10^{4}$ | ACTN4 (C/NU/CS) |
| $1.55\times10^{5}$ | $5.83\times10^{5}$ | $2.16\times10^{5}$ | $2.23\times10^{5}$ | AHCY (C) |
| $4.46\times10^{5}$ | $1.09\times10^{6}$ | $5.14\times10^{5}$ | $6.68\times10^{5}$ | AHSA1 (C/ER) |
| $6.79\times10^{5}$ | $3.06\times10^{6}$ | $6.53\times10^{5}$ | $1.33\times10^{6}$ | AIFM1 (C/NU/MT) |
| $1.60\times10^{4}$ | $3.23\times10^{4}$ | $1.68\times10^{4}$ | $1.50\times10^{4}$ | AKR1B1 (C) |
| $6.93\times10^{3}$ | $2.25\times10^{4}$ | $6.63\times10^{3}$ | $8.22\times10^{3}$ | ALDH9A1 (C) |
| $2.71\times10^{5}$ | $5.90\times10^{5}$ | $2.35\times10^{5}$ | $2.01\times10^{5}$ | ALDOA (C/NU) |
| $2.36\times10^{4}$ | $7.38\times10^{4}$ | $3.07\times10^{4}$ | $2.15\times10^{4}$ | ALDOC (C/CS/EC) |
| $6.73\times10^{5}$ | $2.73\times10^{6}$ | $4.31\times10^{5}$ | $5.69\times10^{5}$ | ANXA2 (PM/EC) |
| $4.75\times10^{5}$ | $1.37\times10^{6}$ | $6.27\times10^{5}$ | $6.42\times10^{5}$ | ANXA5 (C/EC) |
| $8.39\times10^{4}$ | $8.08\times10^{4}$ | $1.60\times10^{4}$ | $5.47\times10^{4}$ | ANXA6 (C) |
| $2.24\times10^{5}$ | $5.79\times10^{5}$ | $2.53\times10^{5}$ | $2.41\times10^{5}$ | ARF3 (GA/C) |
| $4.83\times10^{3}$ | $1.95\times10^{4}$ | $8.40\times10^{3}$ | $7.08\times10^{3}$ | ARF4 (GA/PM) |
| $1.76\times10^{6}$ | $5.35\times10^{6}$ | $2.33\times10^{6}$ | $2.37\times10^{6}$ | ARHGDIA (C) |
| $2.12\times10^{2}$ | $2.36\times10^{4}$ | 0 | 0 | ASF1A (NU) |
| $3.72\times10^{3}$ | $2.22\times10^{4}$ | $3.83\times10^{3}$ | $8.62\times10^{3}$ | ASPRV1 (C/NU) |
| $1.03\times10^{5}$ | $3.28\times10^{5}$ | $1.46\times10^{5}$ | $1.25\times10^{5}$ | ATP1A1 (PM) |
| $1.28\times10^{6}$ | $2.66\times10^{6}$ | $9.70\times10^{5}$ | $9.25\times10^{5}$ | ATP5F1A (MT) |
| $5.88\times10^{6}$ | $1.34\times10^{7}$ | $4.92\times10^{6}$ | $5.16\times10^{6}$ | ATP5F1B (MT) |
| $2.25\times10^{4}$ | $7.35\times10^{4}$ | $2.41\times10^{4}$ | $2.35\times10^{4}$ | ATP5F1C (MT) |
| $3.34\times10^{5}$ | $8.78\times10^{5}$ | $2.89\times10^{5}$ | $3.46\times10^{5}$ | ATP5F1D (MT) |
| $2.36\times10^{4}$ | $6.38\times10^{4}$ | $2.16\times10^{4}$ | $2.17\times10^{4}$ | ATP5ME (MT) |
| $1.33\times10^{3}$ | $1.19\times10^{4}$ | $2.07\times10^{3}$ | $1.92\times10^{3}$ | BCAT1 (C) |
| $1.24\times10^{4}$ | $4.52\times10^{4}$ | $1.56\times10^{4}$ | $1.45\times10^{4}$ | BLVRB (C) |
| $3.97\times10^{4}$ | $5.51\times10^{5}$ | $3.48\times10^{4}$ | $4.01\times10^{4}$ | BOLA2B (C/NU) |
| $3.61\times10^{4}$ | $8.42\times10^{4}$ | $3.95\times10^{4}$ | $3.33\times10^{4}$ | BSG (PM/ER/ES) |
| 0 | $4.96\times10^{4}$ | $3.42\times10^{2}$ | $2.26\times10^{2}$ | C11orf58 (???) |
| $3.68\times10^{4}$ | $1.52\times10^{5}$ | $3.34\times10^{4}$ | $3.83\times10^{4}$ | C1QBP (C/NU/MT/PM/EC) |
| $5.62\times10^{2}$ | $6.14\times10^{3}$ | $9.81\times10^{2}$ | $1.75\times10^{3}$ | CACYBP (C/NU) |
| $2.17\times10^{6}$ | $4.77\times10^{6}$ | $2.25\times10^{6}$ | $3.03\times10^{6}$ | CALU (ER/GA/EC) |
| $6.62\times10^{5}$ | $1.39\times10^{6}$ | $6.28\times10^{5}$ | $6.36\times10^{5}$ | CANX (ER) |
| $7.33\times10^{4}$ | $1.91\times10^{5}$ | $6.86\times10^{4}$ | $5.33\times10^{4}$ | CAPZA1 (CS) |
| $1.24\times10^{5}$ | $3.95\times10^{5}$ | $1.91\times10^{5}$ | $1.12\times10^{5}$ | CAPZB (CS) |
| $6.43\times10^{3}$ | $7.07\times10^{4}$ | $1.42\times10^{4}$ | $2.06\times10^{4}$ | CARHSP1 (C) |
| $3.41\times10^{3}$ | $2.47\times10^{4}$ | $6.07\times10^{3}$ | $4.40\times10^{3}$ | CASP3 (C) |
| $8.19\times10^{3}$ | $2.21\times10^{5}$ | $3.69\times10^{3}$ | $4.55\times10^{3}$ | CAST(C/ER) |
| $2.39\times10^{4}$ | $7.34\times10^{4}$ | $3.19\times10^{4}$ | $4.02\times10^{4}$ | CBX1 (NU) |
| $3.71\times10^{5}$ | $1.25\times10^{6}$ | $4.29\times10^{5}$ | $6.22\times10^{5}$ | CBX3 (NU) |
| $2.15\times10^{3}$ | $1.67\times10^{4}$ | $4.57\times10^{3}$ | $8.00\times10^{3}$ | CBX5 (NU) |
| $2.84\times10^{3}$ | $2.42\times10^{5}$ | $1.74\times10^{3}$ | $6.44\times10^{3}$ | CDK4 (C/NU) |
| $1.26\times10^{4}$ | $2.14\times10^{5}$ | $1.37\times10^{4}$ | $1.61\times10^{4}$ | CDKN2A (C/NU) |
| $1.85\times10^{6}$ | $5.79\times10^{6}$ | $1.90\times10^{6}$ | $1.96\times10^{6}$ | CFL1 (C/NU/CS) |
| $1.23\times10^{4}$ | $4.36\times10^{4}$ | $2.08\times10^{4}$ | $1.44\times10^{4}$ | CFLAR (C/PM) |
| $6.44\times10^{3}$ | $2.16\times10^{4}$ | $1.25\times10^{4}$ | $1.42\times10^{4}$ | CHORDC1 (???) |
| $2.45\times10^{3}$ | $2.67\times10^{5}$ | $2.57\times10^{3}$ | $4.59\times10^{3}$ | CIAPIN1 (C/NU/MT) |
| $2.20\times10^{6}$ | $6.41\times10^{6}$ | $2.84\times10^{6}$ | $2.54\times10^{6}$ | CKB (C) |
| $4.92\times10^{4}$ | $1.23\times10^{5}$ | $5.39\times10^{4}$ | $5.78\times10^{4}$ | CNN3 (C/CS) |
| $7.58\times10^{4}$ | $1.99\times10^{5}$ | $1.19\times10^{4}$ | $2.70\times10^{3}$ | COX17 (C/MT) |

### Table S7
### (page 2)

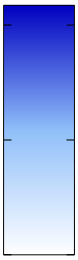


**c-DD-DAO (10 min): no chase (NC) vs. chase (C)**

| NC (L-Ala) | NC (D-Ala) | C (L-Ala) | C (D-Ala) | |
|---|---|---|---|---|
| $1.58\times10^{5}$ | $8.24\times10^{5}$ | $1.57\times10^{5}$ | $2.86\times10^{5}$ | COX5B (MT) |
| $1.95\times10^{4}$ | $4.80\times10^{4}$ | $2.18\times10^{4}$ | $2.00\times10^{4}$ | CPD (PM) |
| $1.43\times10^{4}$ | $1.59\times10^{5}$ | $1.71\times10^{4}$ | $1.72\times10^{4}$ | CPOX (MT) |
| $1.57\times10^{4}$ | $3.97\times10^{5}$ | $2.30\times10^{3}$ | $6.18\times10^{3}$ | CRKL (C/NU/PM) |
| $6.91\times10^{4}$ | $1.16\times10^{6}$ | $2.95\times10^{4}$ | $2.27\times10^{4}$ | CTSB (LY/EC) |
| $1.83\times10^{5}$ | $4.56\times10^{5}$ | $1.55\times10^{5}$ | $2.66\times10^{5}$ | CTSD (LY/EC) |
| $7.96\times10^{3}$ | $3.84\times10^{4}$ | $1.48\times10^{4}$ | $1.38\times10^{4}$ | CYB5R3 (C/MT/ER) |
| $3.17\times10^{2}$ | $6.68\times10^{4}$ | $1.04\times10^{2}$ | $2.30\times10^{2}$ | DCTPP1 (C/NU/MT) |
| $9.03\times10^{3}$ | $4.44\times10^{4}$ | $1.99\times10^{4}$ | $1.32\times10^{4}$ | DDB1 (C/NU) |
| $8.50\times10^{4}$ | $2.64\times10^{5}$ | $7.33\times10^{4}$ | $6.90\times10^{4}$ | DDOST (ER) |
| $1.43\times10^{5}$ | $4.18\times10^{5}$ | $1.89\times10^{5}$ | $1.66\times10^{5}$ | DDX39B (C/NU) |
| $2.66\times10^{5}$ | $1.41\times10^{6}$ | $3.01\times10^{5}$ | $5.74\times10^{5}$ | DNAJB11 (ER) |
| $1.02\times10^{5}$ | $5.21\times10^{5}$ | $1.04\times10^{5}$ | $1.55\times10^{5}$ | DPYSL2 (C/CS/PM) |
| $3.26\times10^{3}$ | $7.96\times10^{4}$ | $1.52\times10^{3}$ | $8.24\times10^{2}$ | DPYSL3 (C) |
| $6.31\times10^{4}$ | $1.36\times10^{5}$ | $7.79\times10^{4}$ | $1.09\times10^{5}$ | DSC3 (PM) |
| $4.43\times10^{6}$ | $9.58\times10^{6}$ | $2.43\times10^{6}$ | $4.42\times10^{6}$ | DSP (CS/PM) |
| $4.57\times10^{4}$ | $1.68\times10^{5}$ | $4.30\times10^{4}$ | $3.98\times10^{4}$ | DSTN (C/CS/EC) |
| $8.03\times10^{4}$ | $3.43\times10^{5}$ | $7.21\times10^{4}$ | $1.06\times10^{5}$ | DUT (NU/MT) |
| 0 | $1.36\times10^{4}$ | $1.39\times10^{3}$ | $3.20\times10^{2}$ | DYNLL1 (NU/MT/CS) |
| $3.36\times10^{4}$ | $7.64\times10^{4}$ | $1.37\times10^{4}$ | $2.44\times10^{4}$ | DYNLRB1 (CS) |
| $6.38\times10^{5}$ | $1.84\times10^{6}$ | $8.61\times10^{5}$ | $8.28\times10^{5}$ | EEF1A1 (C/NU/PM) |
| $1.30\times10^{5}$ | $4.29\times10^{5}$ | $1.04\times10^{5}$ | $1.56\times10^{5}$ | EEF1B2 (C) |
| $8.46\times10^{3}$ | $4.06\times10^{4}$ | $1.46\times10^{4}$ | $1.23\times10^{4}$ | EEF1D (NU) |
| $8.82\times10^{3}$ | $4.35\times10^{4}$ | $1.60\times10^{4}$ | $1.10\times10^{4}$ | EEF1G (C/NU/EC) |
| $4.36\times10^{5}$ | $1.15\times10^{6}$ | $6.29\times10^{5}$ | $6.04\times10^{5}$ | EEF2 (C/NU) |
| $5.13\times10^{3}$ | $1.24\times10^{4}$ | $4.94\times10^{3}$ | $3.44\times10^{3}$ | EIF3H (C) |
| $6.63\times10^{3}$ | $4.04\times10^{4}$ | $1.39\times10^{4}$ | $1.28\times10^{4}$ | EIF4A2 (C) |
| $9.60\times10^{4}$ | $2.69\times10^{5}$ | $6.33\times10^{4}$ | $6.80\times10^{4}$ | EIF5A (C/NU/ER) |
| $8.04\times10^{3}$ | $1.15\times10^{5}$ | $2.32\times10^{4}$ | $2.87\times10^{4}$ | EIF6 (C/NU) |
| $2.47\times10^{5}$ | $5.57\times10^{5}$ | $2.42\times10^{5}$ | $2.68\times10^{5}$ | ELOB (NU) |
| $2.68\times10^{5}$ | $8.61\times10^{5}$ | $2.86\times10^{5}$ | $2.68\times10^{5}$ | ELOC (NU) |
| $1.14\times10^{6}$ | $2.96\times10^{6}$ | $1.21\times10^{6}$ | $1.20\times10^{6}$ | ENO1 (C/NU) |
| $5.04\times10^{2}$ | $4.83\times10^{3}$ | $2.64\times10^{2}$ | $1.48\times10^{3}$ | ENO3 (C) |
| $1.40\times10^{3}$ | $1.64\times10^{4}$ | $4.23\times10^{3}$ | $3.52\times10^{3}$ | ENSG00000276612 (MT) |
| $3.64\times10^{5}$ | $7.50\times10^{5}$ | $2.94\times10^{5}$ | $3.81\times10^{5}$ | ENSG00000286022 (???) |
| $1.61\times10^{4}$ | $2.59\times10^{4}$ | $5.57\times10^{3}$ | $1.18\times10^{4}$ | FDX1 (MT) |
| $1.70\times10^{5}$ | $5.91\times10^{5}$ | $1.79\times10^{5}$ | $2.08\times10^{5}$ | FLG (C/PM) |
| $4.75\times10^{4}$ | $1.47\times10^{5}$ | $4.02\times10^{4}$ | $3.40\times10^{4}$ | FSCN1 (C/CS) |
| $4.88\times10^{4}$ | $1.24\times10^{5}$ | $5.69\times10^{4}$ | $5.99\times10^{4}$ | GANAB (GA/ER) |
| $1.13\times10^{6}$ | $2.97\times10^{6}$ | $1.52\times10^{6}$ | $1.54\times10^{6}$ | GAPDH (C/NU/CS) |
| $3.64\times10^{2}$ | $4.55\times10^{3}$ | $8.28\times10^{2}$ | $7.52\times10^{2}$ | GCLC (C) |
| $4.45\times10^{2}$ | $3.12\times10^{4}$ | $3.11\times10^{2}$ | $1.96\times10^{2}$ | GCLM (C) |
| $3.79\times10^{4}$ | $9.38\times10^{4}$ | $4.50\times10^{4}$ | $3.63\times10^{4}$ | GDI1 (C/GA) |
| $1.56\times10^{2}$ | $4.75\times10^{2}$ | $5.36\times10^{2}$ | $3.40\times10^{3}$ | GFAP (C) |
| $3.28\times10^{5}$ | $1.36\times10^{6}$ | $4.58\times10^{5}$ | $6.45\times10^{5}$ | GLO1 (C/NU/PM/EC) |
| $3.50\times10^{4}$ | $1.76\times10^{5}$ | $6.26\times10^{3}$ | $9.67\times10^{3}$ | GLOD4 (MT) |
| $1.54\times10^{4}$ | $2.03\times10^{5}$ | $1.75\times10^{4}$ | $2.55\times10^{4}$ | GPX4 (C/MT) |
| $1.74\times10^{4}$ | $3.87\times10^{4}$ | $1.19\times10^{4}$ | $1.43\times10^{4}$ | GRPEL1 (MT) |
| $6.42\times10^{4}$ | $4.91\times10^{6}$ | $5.61\times10^{4}$ | $7.09\times10^{4}$ | GSR (C/MT) |
| $1.05\times10^{4}$ | $8.54\times10^{4}$ | $9.64\times10^{3}$ | $1.21\times10^{4}$ | GSTO1 (C) |

### Table S7 (page 3)

#### c-DD-DAO (10 min): no chase (NC) vs. chase (C)

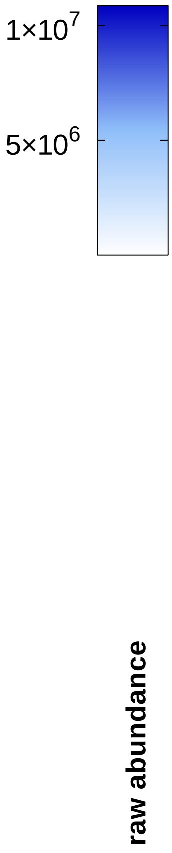


| NC (L-Ala) | NC (D-Ala) | C (L-Ala) | C (D-Ala) | |
|---|---|---|---|---|
| $3.49\times10^{4}$ | $2.68\times10^{5}$ | $4.87\times10^{4}$ | $5.56\times10^{4}$ | GSTP1 (C/NU/MT) |
| $1.04\times10^{4}$ | $8.97\times10^{4}$ | $1.14\times10^{4}$ | $7.03\times10^{3}$ | H2AC6 (NU) |
| $1.60\times10^{4}$ | $3.53\times10^{5}$ | $1.81\times10^{4}$ | $1.19\times10^{4}$ | H2BC11 (NU) |
| $3.82\times10^{4}$ | $9.23\times10^{4}$ | $3.55\times10^{4}$ | $3.66\times10^{4}$ | HNRNPA1 (NU/C) |
| $7.03\times10^{3}$ | $2.41\times10^{4}$ | $8.01\times10^{3}$ | $9.15\times10^{3}$ | HNRNPA2B1 (C/NU/EC) |
| $4.07\times10^{4}$ | $1.73\times10^{5}$ | $4.55\times10^{4}$ | $4.71\times10^{4}$ | HNRNPDL (C/NU) |
| $2.11\times10^{4}$ | $2.02\times10^{6}$ | $3.01\times10^{4}$ | $1.82\times10^{4}$ | HPRT1 (C) |
| $2.73\times10^{4}$ | $6.46\times10^{5}$ | $3.82\times10^{4}$ | $5.33\times10^{4}$ | HSD17B10 (MT) |
| $4.67\times10^{3}$ | $1.33\times10^{4}$ | $4.75\times10^{3}$ | $3.69\times10^{3}$ | HSD17B4 (PO) |
| $1.20\times10^{6}$ | $3.28\times10^{6}$ | $1.62\times10^{6}$ | $1.42\times10^{6}$ | HSP90AA1 (C/NU/MT/PM) |
| $1.44\times10^{6}$ | $4.72\times10^{6}$ | $2.31\times10^{6}$ | $2.04\times10^{6}$ | HSP90AB1 (C/NU/PM/EC) |
| $1.86\times10^{6}$ | $5.46\times10^{6}$ | $2.23\times10^{6}$ | $2.23\times10^{6}$ | HSP90B1 (ER) |
| $2.97\times10^{6}$ | $8.14\times10^{6}$ | $4.00\times10^{6}$ | $3.82\times10^{6}$ | HSPA1B (C/CS) |
| $3.34\times10^{5}$ | $8.07\times10^{5}$ | $4.09\times10^{5}$ | $3.61\times10^{5}$ | HSPA4 (C) |
| $9.59\times10^{3}$ | $5.35\times10^{4}$ | $2.15\times10^{4}$ | $1.93\times10^{4}$ | HSPA4L (C/NU) |
| $1.76\times10^{7}$ | $4.14\times10^{7}$ | $1.46\times10^{7}$ | $1.51\times10^{7}$ | HSPA5 (C/ER) |
| $5.71\times10^{5}$ | $1.88\times10^{6}$ | $6.96\times10^{5}$ | $9.73\times10^{5}$ | HSPA8 (C/PM/NU) |
| $1.07\times10^{7}$ | $2.50\times10^{7}$ | $9.49\times10^{6}$ | $9.45\times10^{6}$ | HSPA9 (MT) |
| $8.04\times10^{4}$ | $5.97\times10^{5}$ | $6.40\times10^{4}$ | $8.38\times10^{4}$ | HSPB1 (C/NU/CS) |
| $1.07\times10^{6}$ | $2.63\times10^{6}$ | $7.61\times10^{5}$ | $8.11\times10^{5}$ | HSPD1 (MT) |
| $4.41\times10^{4}$ | $1.60\times10^{5}$ | $6.42\times10^{4}$ | $5.42\times10^{4}$ | HSPH1 (C) |
| $6.88\times10^{3}$ | $1.78\times10^{4}$ | $8.43\times10^{3}$ | $6.82\times10^{3}$ | IMPA1 (C) |
| $2.83\times10^{4}$ | $9.14\times10^{4}$ | $3.30\times10^{4}$ | $4.35\times10^{4}$ | ISOC1 (C/PO) |
| $6.20\times10^{3}$ | $4.88\times10^{5}$ | $9.61\times10^{3}$ | $9.91\times10^{3}$ | ISYNA1 (C) |
| $1.60\times10^{5}$ | $4.50\times10^{5}$ | $1.97\times10^{5}$ | $1.81\times10^{5}$ | KCTD12 (PM) |
| $3.68\times10^{3}$ | $7.76\times10^{4}$ | $7.49\times10^{2}$ | $2.05\times10^{3}$ | LCP1 (PM/CS) |
| $1.66\times10^{5}$ | $4.39\times10^{5}$ | $1.72\times10^{5}$ | $1.65\times10^{5}$ | LDHA (C) |
| $6.57\times10^{5}$ | $1.57\times10^{6}$ | $9.52\times10^{5}$ | $8.57\times10^{5}$ | LDHB (C/MT) |
| $1.41\times10^{5}$ | $8.75\times10^{5}$ | $1.26\times10^{5}$ | $1.42\times10^{5}$ | LGALS7 (C/NU/EC) |
| $1.57\times10^{5}$ | $4.03\times10^{5}$ | $1.52\times10^{5}$ | $1.43\times10^{5}$ | LMAN2 (GA/ER) |
| $1.43\times10^{4}$ | $7.80\times10^{4}$ | $7.02\times10^{3}$ | $9.98\times10^{3}$ | LMNA (NU) |
| $1.12\times10^{5}$ | $2.97\times10^{5}$ | $1.63\times10^{5}$ | $1.26\times10^{5}$ | LTA4H (C) |
| $2.16\times10^{4}$ | $4.53\times10^{4}$ | $3.78\times10^{4}$ | $2.58\times10^{4}$ | LTF (C/EC) |
| $3.63\times10^{3}$ | $1.74\times10^{4}$ | $7.45\times10^{3}$ | $7.10\times10^{3}$ | LUC7L2 (NU) |
| $1.67\times10^{4}$ | $4.81\times10^{4}$ | $1.94\times10^{4}$ | $1.21\times10^{4}$ | LXN (C) |
| $8.33\times10^{3}$ | $3.75\times10^{4}$ | $1.22\times10^{4}$ | $1.51\times10^{4}$ | MAD2L1 (C/CS/NU) |
| $1.93\times10^{4}$ | $3.33\times10^{5}$ | $1.53\times10^{4}$ | $1.57\times10^{4}$ | MARCKS (CS/PM) |
| $6.39\times10^{3}$ | $5.89\times10^{4}$ | $1.16\times10^{3}$ | $2.61\times10^{3}$ | MARCKSL1 (CS/PM) |
| $1.16\times10^{5}$ | $3.90\times10^{5}$ | $1.52\times10^{5}$ | $1.20\times10^{5}$ | MAT2A (C) |
| $4.43\times10^{4}$ | $1.06\times10^{5}$ | $4.22\times10^{4}$ | $4.35\times10^{4}$ | MDH2 (MT) |
| $4.65\times10^{4}$ | $3.99\times10^{5}$ | $6.96\times10^{4}$ | $5.58\times10^{4}$ | MIF (C/EC) |
| $1.17\times10^{5}$ | $2.40\times10^{5}$ | $6.33\times10^{4}$ | $6.34\times10^{4}$ | MTPN (C/NU) |
| $3.34\times10^{4}$ | $1.13\times10^{5}$ | $3.41\times10^{4}$ | $3.05\times10^{4}$ | MYG1 (NU/MT) |
| $1.11\times10^{4}$ | $4.49\times10^{4}$ | $5.43\times10^{3}$ | $6.52\times10^{3}$ | MYH9 (CS) |
| $1.12\times10^{4}$ | $3.36\times10^{4}$ | $8.61\times10^{3}$ | $8.27\times10^{3}$ | MYL6 (C/CS/EC) |
| $1.68\times10^{5}$ | $5.93\times10^{5}$ | $1.98\times10^{5}$ | $1.89\times10^{5}$ | NASP (C/NU) |
| $4.36\times10^{5}$ | $1.82\times10^{6}$ | $4.78\times10^{5}$ | $6.74\times10^{5}$ | NME1-NME2 (C) |
| $2.07\times10^{5}$ | $6.99\times10^{5}$ | $2.28\times10^{5}$ | $1.81\times10^{5}$ | NPEPPS (C/NU) |
| $1.96\times10^{5}$ | $8.80\times10^{5}$ | $2.70\times10^{5}$ | $3.67\times10^{5}$ | NPM1 (NU) |
| $7.28\times10^{3}$ | $5.09\times10^{5}$ | $5.27\times10^{3}$ | $6.97\times10^{3}$ | NUDC (CS/NU) |

#### c-DD-DAO (10 min): no chase (NC) vs. chase (C)

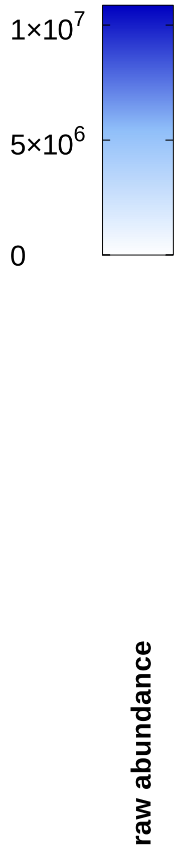


| | NC (L-Ala) | NC (D-Ala) | C (L-Ala) | C (D-Ala) |
|---|---|---|---|---|
| NUDCD2 (CS) | $6.14\times10^{4}$ | $1.90\times10^{5}$ | $7.68\times10^{4}$ | $9.37\times10^{4}$ |
| NUDT5 (NU) | $2.38\times10^{4}$ | $7.30\times10^{4}$ | $2.32\times10^{4}$ | $2.49\times10^{4}$ |
| P4HB (PM/ER) | $2.61\times10^{6}$ | $6.49\times10^{6}$ | $2.28\times10^{6}$ | $2.51\times10^{6}$ |
| PAFAH1B2 (C) | $2.41\times10^{3}$ | $5.53\times10^{3}$ | $2.30\times10^{3}$ | $2.64\times10^{3}$ |
| PAFAH1B3 (C) | $7.36\times10^{3}$ | $4.47\times10^{4}$ | $1.82\times10^{4}$ | $1.51\times10^{4}$ |
| PAICS (C/EC) | $8.95\times10^{3}$ | $1.15\times10^{5}$ | $6.61\times10^{3}$ | $6.62\times10^{3}$ |
| PARK7 (C/ER/NU/PM/MT) | $4.66\times10^{5}$ | $1.28\times10^{6}$ | $4.97\times10^{5}$ | $5.62\times10^{5}$ |
| PBDC1 (???) | $1.29\times10^{5}$ | $3.87\times10^{5}$ | $1.81\times10^{5}$ | $2.01\times10^{5}$ |
| PCCA (MT) | $9.07\times10^{4}$ | $2.49\times10^{5}$ | $1.09\times10^{5}$ | $1.14\times10^{5}$ |
| PCMT1 (C) | $1.35\times10^{4}$ | $5.63\times10^{5}$ | $4.01\times10^{4}$ | $6.33\times10^{4}$ |
| PDIA3 (ER) | $4.54\times10^{5}$ | $2.02\times10^{6}$ | $5.67\times10^{5}$ | $5.57\times10^{5}$ |
| PDIA4 (ER) | $3.91\times10^{5}$ | $1.43\times10^{6}$ | $4.85\times10^{5}$ | $5.37\times10^{5}$ |
| PDIA6 (ER/PM) | $1.05\times10^{5}$ | $3.93\times10^{5}$ | $1.11\times10^{5}$ | $1.33\times10^{5}$ |
| PEBP1 (C) | $3.04\times10^{5}$ | $1.01\times10^{6}$ | $3.71\times10^{5}$ | $3.54\times10^{5}$ |
| PFDN2 (C/NU/MT) | $1.44\times10^{3}$ | $7.06\times10^{4}$ | $7.15\times10^{2}$ | $1.31\times10^{3}$ |
| PFDN4 (C/NU/MT) | 0 | $1.08\times10^{5}$ | 0 | 0 |
| PFN1 (CS) | $5.52\times10^{4}$ | $1.90\times10^{5}$ | $3.71\times10^{4}$ | $3.52\times10^{4}$ |
| PGAM1 (C/EC) | $2.59\times10^{5}$ | $5.55\times10^{5}$ | $2.78\times10^{5}$ | $2.75\times10^{5}$ |
| PGD (C) | $1.67\times10^{4}$ | $4.39\times10^{4}$ | $1.94\times10^{4}$ | $1.68\times10^{4}$ |
| PGK1 (C) | $8.00\times10^{4}$ | $2.58\times10^{5}$ | $1.10\times10^{5}$ | $1.01\times10^{5}$ |
| PHPT1 (C) | $1.55\times10^{5}$ | $6.93\times10^{5}$ | $1.41\times10^{5}$ | $2.13\times10^{5}$ |
| PITHD1 (C) | $1.96\times10^{3}$ | $4.59\times10^{5}$ | $1.78\times10^{3}$ | $1.58\times10^{3}$ |
| PKM (C/NU) | $6.52\times10^{4}$ | $1.85\times10^{5}$ | $5.80\times10^{4}$ | $7.03\times10^{4}$ |
| PKP1 (NU) | $2.39\times10^{5}$ | $8.33\times10^{5}$ | $1.46\times10^{5}$ | $2.78\times10^{5}$ |
| PLS1 (C) | $1.16\times10^{5}$ | $3.59\times10^{5}$ | $9.29\times10^{4}$ | $1.26\times10^{5}$ |
| PLS3 (C) | $2.77\times10^{4}$ | $2.63\times10^{5}$ | $3.34\times10^{4}$ | $3.14\times10^{4}$ |
| PNP (C) | $7.90\times10^{4}$ | $1.65\times10^{5}$ | $7.26\times10^{4}$ | $7.34\times10^{4}$ |
| POF1B (C/CS) | $5.60\times10^{4}$ | $1.66\times10^{5}$ | $3.04\times10^{4}$ | $5.25\times10^{4}$ |
| POLR2H (NU) | $1.01\times10^{5}$ | $2.46\times10^{5}$ | $1.36\times10^{5}$ | $1.01\times10^{5}$ |
| PPA1 (C) | $3.49\times10^{5}$ | $1.16\times10^{6}$ | $5.21\times10^{5}$ | $4.26\times10^{5}$ |
| PPIA (C/NU/EC) | $2.71\times10^{6}$ | $8.10\times10^{6}$ | $3.07\times10^{6}$ | $2.62\times10^{6}$ |
| PPP2R2A (C/NU) | $6.68\times10^{3}$ | $3.91\times10^{5}$ | $3.08\times10^{3}$ | $9.72\times10^{3}$ |
| PRDX1 (C) | $3.10\times10^{6}$ | $4.38\times10^{7}$ | $3.16\times10^{6}$ | $2.17\times10^{6}$ |
| PRDX2 (C) | $8.53\times10^{5}$ | $1.07\times10^{7}$ | $9.45\times10^{5}$ | $7.50\times10^{5}$ |
| PRDX3 (MT/C/EE) | $7.06\times10^{5}$ | $1.33\times10^{7}$ | $5.48\times10^{5}$ | $6.64\times10^{5}$ |
| PRDX4 (C/ER) | $4.44\times10^{5}$ | $3.49\times10^{6}$ | $4.55\times10^{5}$ | $5.13\times10^{5}$ |
| PRDX5 (C/NU/MT/PO) | $3.20\times10^{3}$ | $1.93\times10^{5}$ | $8.86\times10^{3}$ | $2.13\times10^{4}$ |
| PRDX6 (C/LY) | $9.81\times10^{4}$ | $1.50\times10^{6}$ | $7.27\times10^{4}$ | $1.47\times10^{5}$ |
| PRMT5 (C/NU/GA) | $1.85\times10^{4}$ | $3.15\times10^{5}$ | $1.69\times10^{4}$ | $1.49\times10^{4}$ |
| PSMA6 (C/NU) | $1.10\times10^{4}$ | $2.66\times10^{4}$ | $1.21\times10^{4}$ | $1.34\times10^{4}$ |
| PSMA7 (C/NU) | $1.57\times10^{4}$ | $4.68\times10^{4}$ | $1.82\times10^{4}$ | $1.80\times10^{4}$ |
| PSMD1 (C/NU/LY/EC) | $6.42\times10^{4}$ | $1.71\times10^{5}$ | $9.83\times10^{4}$ | $9.34\times10^{4}$ |
| PSMD4 (C/NU) | $4.57\times10^{4}$ | $1.76\times10^{5}$ | $7.92\times10^{4}$ | $8.14\times10^{4}$ |
| RAB11A (PM/ES) | $5.52\times10^{3}$ | $2.43\times10^{4}$ | $3.34\times10^{3}$ | $4.24\times10^{3}$ |
| RAB1A (C/ES/GA/PM) | $3.52\times10^{4}$ | $8.72\times10^{4}$ | $5.68\times10^{4}$ | $4.42\times10^{4}$ |
| RAB2A (GA/ER) | $1.12\times10^{4}$ | $4.55\times10^{4}$ | $7.33\times10^{3}$ | $1.02\times10^{4}$ |
| RAB35 (PM/ES) | $3.08\times10^{4}$ | $8.20\times10^{4}$ | $3.67\times10^{4}$ | $3.45\times10^{4}$ |
| RACK1 (C/NU/PM) | $1.12\times10^{4}$ | $4.21\times10^{4}$ | $1.53\times10^{4}$ | $1.39\times10^{4}$ |
| RANBP1 (C/NU/CS) | $7.65\times10^{5}$ | $2.69\times10^{6}$ | $7.34\times10^{5}$ | $8.58\times10^{5}$ |
| RAP1A (PM/C/ES) | $1.50\times10^{4}$ | $5.44\times10^{4}$ | $1.63\times10^{4}$ | $1.63\times10^{4}$ |

### Table S7 (page 5)

**c-DD-DAO (10 min): no chase (NC) vs. chase (C)**

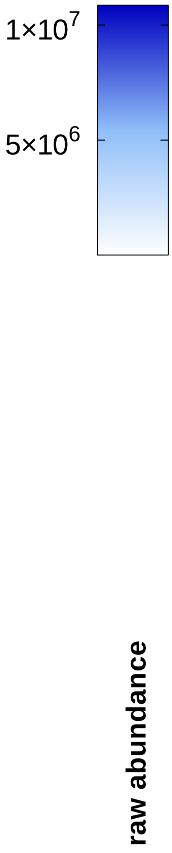


| | NC (L-Ala) | NC (D-Ala) | C (L-Ala) | C (D-Ala) |
|---|---|---|---|---|
| RBBP4 (NU) | $2.27\times10^{4}$ | $5.59\times10^{4}$ | $2.85\times10^{4}$ | $2.37\times10^{4}$ |
| RBBP7 (NU) | $2.80\times10^{4}$ | $6.53\times10^{5}$ | $4.41\times10^{4}$ | $3.43\times10^{4}$ |
| RCN1 (ER) | $3.72\times10^{5}$ | $9.41\times10^{5}$ | $3.66\times10^{5}$ | $4.32\times10^{5}$ |
| RPA3 (NU) | $1.12\times10^{4}$ | $3.43\times10^{4}$ | $1.00\times10^{4}$ | $9.76\times10^{3}$ |
| RPL8 (C) | $3.61\times10^{2}$ | $2.83\times10^{3}$ | $7.04\times10^{2}$ | $3.55\times10^{2}$ |
| RPLP1 (C) | $1.37\times10^{4}$ | $3.21\times10^{4}$ | $1.15\times10^{4}$ | $7.32\times10^{3}$ |
| RPLP2 (C/EC) | $2.89\times10^{4}$ | $9.70\times10^{4}$ | $2.20\times10^{4}$ | $2.08\times10^{4}$ |
| RPN1 (ER) | $4.98\times10^{4}$ | $1.76\times10^{5}$ | $6.74\times10^{4}$ | $6.47\times10^{4}$ |
| RPN2 (ER) | $3.36\times10^{4}$ | $1.07\times10^{5}$ | $4.90\times10^{4}$ | $4.74\times10^{4}$ |
| RPS12 (C) | $8.22\times10^{4}$ | $2.42\times10^{5}$ | $8.98\times10^{4}$ | $6.59\times10^{4}$ |
| RPS14 (C/NU/EC) | $3.48\times10^{2}$ | $1.01\times10^{3}$ | $1.07\times10^{3}$ | $6.95\times10^{2}$ |
| RPS21 (C/ER) | $7.82\times10^{4}$ | $5.42\times10^{5}$ | $8.64\times10^{4}$ | $1.86\times10^{5}$ |
| RPS27A (C/NU) | $9.37\times10^{5}$ | $2.94\times10^{6}$ | $8.61\times10^{5}$ | $8.84\times10^{5}$ |
| RPSA (NU/C/PM) | $6.75\times10^{4}$ | $2.19\times10^{5}$ | $8.03\times10^{4}$ | $5.41\times10^{4}$ |
| RUVBL1 (C/NU/CS) | $2.18\times10^{4}$ | $5.07\times10^{4}$ | $2.37\times10^{4}$ | $1.99\times10^{4}$ |
| RUVBL2 (C/NU) | $1.90\times10^{4}$ | $6.92\times10^{4}$ | $3.63\times10^{4}$ | $2.69\times10^{4}$ |
| S100A14 (C) | $1.00\times10^{5}$ | $2.08\times10^{5}$ | $6.74\times10^{4}$ | $1.09\times10^{5}$ |
| S100A16 (C/NU) | $1.93\times10^{4}$ | $3.85\times10^{4}$ | $1.13\times10^{4}$ | $2.14\times10^{4}$ |
| S100A7 (C/EC) | $7.28\times10^{4}$ | $2.32\times10^{5}$ | $3.37\times10^{4}$ | $3.17\times10^{4}$ |
| S100A8 (C/PM/EC/CS) | $2.92\times10^{5}$ | $5.94\times10^{5}$ | $1.96\times10^{5}$ | $2.71\times10^{5}$ |
| SAR1A (GA/ER) | $4.57\times10^{4}$ | $1.07\times10^{5}$ | $1.47\times10^{5}$ | $4.69\times10^{4}$ |
| SCO2 (MT) | $1.20\times10^{4}$ | $3.02\times10^{4}$ | $9.65\times10^{3}$ | $9.12\times10^{3}$ |
| SERPINB1 (C/ES/LY/EC) | $1.57\times10^{3}$ | $1.36\times10^{4}$ | $7.54\times10^{1}$ | $5.84\times10^{1}$ |
| SERPINB7 (C) | $9.10\times10^{3}$ | $2.55\times10^{4}$ | $4.53\times10^{3}$ | $5.64\times10^{3}$ |
| SFN (C/NU/EC) | $1.74\times10^{4}$ | $1.15\times10^{5}$ | $8.53\times10^{3}$ | $5.19\times10^{3}$ |
| SKP1 (C/NU) | $9.37\times10^{3}$ | $1.16\times10^{6}$ | $1.09\times10^{4}$ | $1.64\times10^{4}$ |
| SLC1A5 (PM) | $6.08\times10^{4}$ | $1.37\times10^{5}$ | $6.03\times10^{4}$ | $5.16\times10^{4}$ |
| SLC3A2 (LY/PM) | $2.04\times10^{4}$ | $4.78\times10^{4}$ | $1.95\times10^{4}$ | $2.22\times10^{4}$ |
| SNRPF (C/NU) | $2.08\times10^{5}$ | $8.20\times10^{5}$ | $2.45\times10^{5}$ | $3.59\times10^{5}$ |
| SOD1 (C/NU/MT) | $5.73\times10^{3}$ | $4.27\times10^{4}$ | $5.21\times10^{3}$ | $8.96\times10^{3}$ |
| SSR1 (ER) | $8.87\times10^{3}$ | $3.20\times10^{4}$ | $2.15\times10^{4}$ | $2.29\times10^{4}$ |
| SSR4 (ER) | $4.18\times10^{4}$ | $1.21\times10^{5}$ | $4.54\times10^{4}$ | $4.31\times10^{4}$ |
| STRAP (C/NU) | $6.08\times10^{4}$ | $4.44\times10^{5}$ | $8.08\times10^{4}$ | $6.51\times10^{4}$ |
| SUMO3 (C/NU) | $6.80\times10^{4}$ | $1.64\times10^{5}$ | $6.04\times10^{4}$ | $7.41\times10^{4}$ |
| SYPL1 (VE) | $9.95\times10^{3}$ | $3.20\times10^{4}$ | $1.37\times10^{4}$ | $1.38\times10^{4}$ |
| TALDO1 (C) | $2.65\times10^{5}$ | $5.96\times10^{5}$ | $2.28\times10^{5}$ | $2.30\times10^{5}$ |
| TGM1 (C/PM) | $2.96\times10^{5}$ | $6.32\times10^{5}$ | $2.18\times10^{5}$ | $3.35\times10^{5}$ |
| THOP1 (C) | $1.27\times10^{3}$ | $9.93\times10^{3}$ | $1.16\times10^{3}$ | $2.43\times10^{2}$ |
| TIPRL (C) | $4.90\times10^{3}$ | $3.42\times10^{4}$ | $8.60\times10^{3}$ | $9.32\times10^{3}$ |
| TKT (C/NU/ER/PO/EC) | $3.39\times10^{4}$ | $1.21\times10^{5}$ | $4.49\times10^{4}$ | $5.46\times10^{4}$ |
| TMED10 (PM/ER/GA) | $1.96\times10^{4}$ | $6.52\times10^{4}$ | $2.26\times10^{4}$ | $2.18\times10^{4}$ |
| TMED7 (ER/GA) | $8.32\times10^{4}$ | $3.11\times10^{5}$ | $6.90\times10^{4}$ | $6.17\times10^{4}$ |
| TOMM22 (MT) | $2.90\times10^{4}$ | $8.01\times10^{4}$ | $3.61\times10^{4}$ | $3.34\times10^{4}$ |
| TOMM40 (MT) | $7.82\times10^{3}$ | $2.58\times10^{4}$ | $1.21\times10^{4}$ | $9.26\times10^{3}$ |
| TPM3 (CS) | $8.81\times10^{4}$ | $2.59\times10^{6}$ | $4.17\times10^{4}$ | $6.24\times10^{4}$ |
| TPM4 (CS) | $8.61\times10^{3}$ | $3.26\times10^{5}$ | $2.95\times10^{3}$ | $5.54\times10^{3}$ |
| TPT1 (C) | $8.14\times10^{3}$ | $1.89\times10^{4}$ | $1.32\times10^{4}$ | $1.18\times10^{4}$ |
| TRA2A (NU) | $4.44\times10^{3}$ | $1.23\times10^{4}$ | $5.23\times10^{3}$ | $6.66\times10^{3}$ |
| TRAP1 (MT) | $7.76\times10^{4}$ | $2.48\times10^{5}$ | $9.52\times10^{4}$ | $9.71\times10^{4}$ |
| TRMT112 (NU) | $1.14\times10^{4}$ | $4.15\times10^{4}$ | $1.25\times10^{4}$ | $1.39\times10^{4}$ |

### Table S7
### (page 6)

#### c-DD-DAO (10 min): no chase (NC) vs. chase (C)

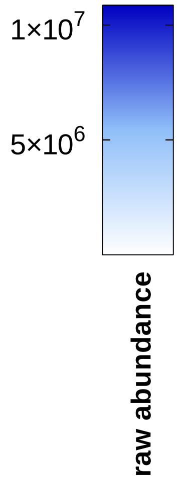


| | NC (L-Ala) | NC (D-Ala) | C (L-Ala) | C (D-Ala) |
|---|---|---|---|---|
| TUBA1C (CS) | $5.27\times10^{4}$ | $2.09\times10^{5}$ | $5.48\times10^{4}$ | $4.05\times10^{4}$ |
| TUBB (CS) | $2.05\times10^{3}$ | $3.72\times10^{4}$ | $3.48\times10^{3}$ | $3.43\times10^{3}$ |
| TXN (C/NU/EC) | $9.06\times10^{5}$ | $2.55\times10^{7}$ | $5.90\times10^{5}$ | $7.39\times10^{5}$ |
| TXNDC17 (C) | $4.75\times10^{3}$ | $8.02\times10^{5}$ | $5.20\times10^{3}$ | $1.16\times10^{4}$ |
| TXNDC5 (ER) | $8.56\times10^{4}$ | $2.44\times10^{5}$ | $5.00\times10^{4}$ | $5.99\times10^{4}$ |
| UBA1 (C/NU/MT) | $3.68\times10^{5}$ | $1.06\times10^{6}$ | $5.46\times10^{5}$ | $4.70\times10^{5}$ |
| UBE2K (C) | $9.95\times10^{3}$ | $9.96\times10^{4}$ | $2.05\times10^{4}$ | $3.27\times10^{4}$ |
| UBE2L3 (C/NU) | $7.14\times10^{2}$ | $1.46\times10^{5}$ | $1.44\times10^{3}$ | $1.93\times10^{3}$ |
| UCHL1 (C/ER) | $4.68\times10^{6}$ | $1.32\times10^{7}$ | $6.00\times10^{6}$ | $5.47\times10^{6}$ |
| UCHL3 (C) | $8.28\times10^{4}$ | $1.80\times10^{5}$ | $7.72\times10^{4}$ | $8.70\times10^{4}$ |
| UGGT1 (ER) | $6.53\times10^{4}$ | $2.22\times10^{5}$ | $5.58\times10^{4}$ | $9.13\times10^{4}$ |
| UQCRFS1 (MT) | $1.73\times10^{5}$ | $7.28\times10^{5}$ | $1.72\times10^{5}$ | $2.81\times10^{5}$ |
| VCP (C/NU/ER) | $7.08\times10^{4}$ | $3.12\times10^{5}$ | $9.75\times10^{4}$ | $1.12\times10^{5}$ |
| VDAC3 (MT) | $2.15\times10^{4}$ | $5.33\times10^{4}$ | $3.24\times10^{4}$ | $2.55\times10^{4}$ |
| VIM (C/NU/PM) | $1.59\times10^{4}$ | $4.46\times10^{4}$ | $1.01\times10^{4}$ | $8.35\times10^{3}$ |
| WDR77 (C/NU) | $4.58\times10^{3}$ | $1.86\times10^{5}$ | $8.53\times10^{3}$ | $7.06\times10^{3}$ |
| YARS1 (C) | $4.65\times10^{4}$ | $1.04\times10^{5}$ | $5.25\times10^{4}$ | $4.16\times10^{4}$ |
| YWHAB (C) | $2.53\times10^{5}$ | $6.64\times10^{5}$ | $2.50\times10^{5}$ | $2.91\times10^{5}$ |
| YWHAG (C) | $2.18\times10^{4}$ | $8.51\times10^{4}$ | $2.38\times10^{4}$ | $2.21\times10^{4}$ |
| YWHAH (C/PM/MT/EC) | $3.23\times10^{4}$ | $7.12\times10^{4}$ | $3.29\times10^{4}$ | $2.99\times10^{4}$ |
| YWHAQ (C) | $1.93\times10^{5}$ | $1.51\times10^{6}$ | $2.93\times10^{5}$ | $3.04\times10^{5}$ |
| YWHAZ (C) | $2.78\times10^{5}$ | $1.05\times10^{6}$ | $3.54\times10^{5}$ | $3.69\times10^{5}$ |
| ZNF598 (???) | $1.29\times10^{6}$ | $4.15\times10^{6}$ | $1.27\times10^{6}$ | $1.33\times10^{6}$ |
| ZPR1 (C/NU) | $2.48\times10^{3}$ | $7.48\times10^{3}$ | $4.41\times10^{3}$ | $3.23\times10^{3}$ |

#### Table S8
#### (page 1)

**External $H_2O_2$ (1 mM, 10 min)**

protein abundance (percentage distribution) — colour scale: 0, 20, 40, 60, 80, 100

| NaCl eluate | DTT eluate | |
|---|---|---|
| 19.94 | 80.06 | AAMDC (C) |
| 4.67 | 95.33 | AASDHPPT (C) |
| 15.82 | 84.18 | ACP1 (C) |
| 0.63 | 99.37 | ACP6 (MT) |
| 17.54 | 82.46 | ACT (???) |
| 42.07 | 57.93 | ACTN1 (C/PM) |
| 1.01 | 98.99 | ACYP2 (?) |
| 46.27 | 53.73 | AHCY (C) |
| 1.38 | 98.62 | AIF1L (CS/PM) |
| 6.95 | 93.05 | AK2 (MT) |
| 16.23 | 83.77 | AKAP12 (CS/PM) |
| 25.30 | 74.70 | ANXA1 (C/NU/PM/EC) |
| 8.92 | 91.08 | ANXA2 (PM/EC) |
| 31.62 | 68.38 | APOA1 (C/PM/EC) |
| 24.90 | 75.10 | ARL6IP4 (NU) |
| 4.38 | 95.62 | ASF1A (NU) |
| 7.15 | 92.85 | ASF1B (NU) |
| 4.92 | 95.08 | ASNS (C) |
| 33.17 | 66.83 | ASRGL1 (C) |
| 0.44 | 99.56 | ATOX1 (C) |
| 6.54 | 93.46 | BAG3 (C/NU) |
| 7.56 | 92.44 | BAG4 (C) |
| 2.56 | 97.44 | BAG5 (C/NU/MT) |
| 6.26 | 93.74 | BICD2 (C/NU/CS/GA) |
| 4.03 | 95.97 | BID (C/MT) |
| 45.25 | 54.75 | BLMH (C) |
| 5.48 | 94.52 | BOLA1 (MT) |
| 4.78 | 95.22 | BOLA2B (C/NU) |
| 2.13 | 97.87 | C11orf54 (NU) |
| 5.54 | 94.46 | C11orf58 (???) |
| 1.93 | 98.07 | C12orf29 (???) |
| 0 | 100.00 | C1orf174 (NU) |
| 19.81 | 80.19 | C1orf50 (???) |
| 1.05 | 98.95 | C1orf52 (NU) |
| 13.66 | 86.34 | CALD1 (CS) |
| 12.79 | 87.21 | CAPZA2 (C/CS/EC) |
| 16.33 | 83.67 | CARHSP1 (C) |
| 18.89 | 81.11 | CASP3 (C) |
| 4.39 | 95.61 | CAST (C/ER) |
| 3.92 | 96.08 | CCDC50 (C) |
| 8.99 | 91.01 | CCDC6 (C/CS) |
| 10.81 | 89.19 | CCS (C) |
| 9.55 | 90.45 | CDK4 (C/NU) |
| 40.85 | 59.15 | CDK6(C/NU/CS) |
| 9.13 | 90.87 | CHAC2 (C) |
| 4.64 | 95.36 | CHMP5 (C/ES) |
| 47.82 | 52.18 | CIAO2A (C/NU) |
| 5.74 | 94.26 | CIAPIN1 (C/NU/MT) |
| 6.60 | 93.40 | CKMT1A (MT) |
| 31.61 | 68.39 | CLIC1 (C/NU/PM) |

**Table S8 (page 2)**

**External $H_2O_2$ (1 mM, 10 min)**

Color scale: protein abundance (percentage distribution), 0 – 20 – 40 – 60 – 80 – 100

| Protein | NaCl eluate | DTT eluate |
|---|---|---|
| COX17 (C/MT) | 0.17 | 99.83 |
| CPOX (MT) | 5.10 | 94.90 |
| CPPED1 (C) | 2.70 | 97.30 |
| CRKL (C/NU/PM) | 4.86 | 95.14 |
| CRMP1 (C/CS) | 21.09 | 78.91 |
| CRYZL1 (C) | 8.46 | 91.54 |
| CSTB (C/NU) | 1.72 | 98.28 |
| CTH (C) | 2.49 | 97.51 |
| CWC15 (NU) | 3.41 | 96.59 |
| DCPS (C/NU) | 5.78 | 94.22 |
| DCTPP1 (C/NU/MT) | 5.99 | 94.01 |
| DENR (???) | 5.92 | 94.08 |
| DHPS (C) | 3.02 | 96.98 |
| DLAT (MT) | 14.54 | 85.46 |
| DNPEP (C) | 2.44 | 97.56 |
| DPYSL2 (C/CS/PM) | 25.25 | 74.75 |
| DPYSL3 (C) | 18.38 | 81.62 |
| DPYSL5 (C) | 11.58 | 88.42 |
| DRG1 (C/NU) | 76.00 | 24.00 |
| DSTN (C/CS/EC) | 30.89 | 69.11 |
| DUSP3 (NU) | 19.04 | 80.96 |
| DYNLL1 (NU/CS/MT) | 6.02 | 93.98 |
| EFHD2 (PM) | 7.19 | 92.81 |
| EIF1AD (NU) | 15.07 | 84.93 |
| EIF3B (C) | 40.98 | 59.02 |
| EIF4B (C) | 4.46 | 95.54 |
| EIF4EBP1 (C/NU) | 0.06 | 99.94 |
| EIF6 (C/NU) | 28.55 | 71.45 |
| EPS15L1 (NU/PM) | 8.58 | 91.42 |
| ERICH5 (???) | 6.78 | 93.22 |
| EVPL (CS) | 28.01 | 71.99 |
| FAH (C/EC) | 0.58 | 99.42 |
| FLG (C/PM) | 10.08 | 89.92 |
| FLYWCH2 (???) | 6.09 | 93.91 |
| FN3KRP (C) | 14.65 | 85.35 |
| FNTA (C/PM/CS) | 3.99 | 96.01 |
| FNTB (C/CS) | 1.66 | 98.34 |
| FTH1 (C/NU/LY/EC) | 9.76 | 90.24 |
| GCLC (C) | 4.24 | 95.76 |
| GCLM (C) | 6.54 | 93.46 |
| GEMIN6 (C/NU) | 12.15 | 87.85 |
| GFER (C/MT/EC) | 2.04 | 97.96 |
| GLDC (MT) | 15.13 | 84.87 |
| GLOD4 (MT) | 9.54 | 90.46 |
| GLRX3 (C) | 28.42 | 71.58 |
| GPHN (C/PM) | 7.88 | 92.12 |
| GPX1 (C) | 28.91 | 71.09 |
| GPX4 (C/MT) | 15.35 | 84.65 |
| GRB2 (C/MT/ES/GA) | 6.85 | 93.15 |
| GRIPAP1 (ES) | 6.11 | 93.89 |

**Table S8**
**(page 3)**

**External $H_2O_2$ (1 mM, 10 min)**

protein abundance (percentage distribution): color scale 0, 20, 40, 60, 80, 100

| Protein | NaCl eluate | DTT eluate |
|---|---|---|
| GSR (C/MT) | 2.67 | 97.33 |
| HEXIM1 (C/NU) | 10.33 | 89.67 |
| HIRIP3 (NU) | 15.23 | 84.77 |
| HPRT1 (C) | 5.72 | 94.28 |
| HSD17B10 (MT) | 7.28 | 92.72 |
| HSD17B8 (MT) | 1.38 | 98.62 |
| HSPA4L (C/NU) | 23.26 | 76.74 |
| IAH1 (???) | 13.89 | 86.11 |
| IBA57 (MT) | 1.95 | 98.05 |
| IMPDH1 (C/NU) | 10.95 | 89.05 |
| IMPDH2 (C/NU) | 17.72 | 82.28 |
| ISCA2 (MT) | 14.05 | 85.95 |
| ISCU (C/NU/MT) | 3.25 | 96.75 |
| ISYNA1 (C) | 4.83 | 95.17 |
| JMJD6 (C/NU) | 63.52 | 36.48 |
| JPT2 (C/NU) | 2.90 | 97.10 |
| KCTD5 (C/NU) | 4.23 | 95.77 |
| KHDRBS1 (C/NU/PM) | 66.12 | 33.88 |
| KHSRP (C/NU) | 8.30 | 91.70 |
| KIF3A (CS) | 19.01 | 80.99 |
| LARP7 (NU) | 86.53 | 13.47 |
| LBH (C/NU) | 0.36 | 99.64 |
| LCP1 (PM/CS) | 10.56 | 89.44 |
| LRRFIP1rC/NU) | 7.96 | 92.04 |
| LTF (C/EC) | 93.21 | 6.79 |
| LYRM7 (MT) | 13.02 | 86.98 |
| MAP4 (CS) | 8.67 | 91.33 |
| MAPK1(C/CS/NU) | 25.07 | 74.93 |
| MAPK14 (C/NU) | 12.62 | 87.38 |
| MARCKS (CS/PM) | 11.65 | 88.35 |
| MARCKSL1(CS/PM) | 1.03 | 98.97 |
| MCMBP (NU) | 9.50 | 90.50 |
| MDH1 (C) | 11.37 | 88.63 |
| METTL26 (???) | 9.62 | 90.38 |
| MINDY3 (NU) | 27.76 | 72.24 |
| MMAB (MT) | 15.54 | 84.46 |
| MOCS2 (C) | 5.15 | 94.85 |
| MPP1 (PM) | 17.81 | 82.19 |
| MSRB3 (MT/ER) | 6.29 | 93.71 |
| MTAP (C/NU) | 4.53 | 95.47 |
| NASP (C/NU) | 29.27 | 70.73 |
| NDUFAB1 (MT) | 24.98 | 75.02 |
| NFU1 (C/MT) | 10.13 | 89.87 |
| NPEPPS (C/NU) | 22.34 | 77.66 |
| NT5DC2 (???) | 6.39 | 93.61 |
| NUDC (CS/NU) | 3.42 | 96.58 |
| OGFR (C/NU) | 4.78 | 95.22 |
| OPTN (C/GA/ES) | 9.61 | 90.39 |
| OTUB1 (C) | 18.88 | 81.12 |
| PABPC4 (C) | 48.42 | 51.58 |

**Table S8**
**(page 4)**

**External $H_2O_2$ (1 mM, 10 min)**

protein abundance (percentage distribution): 100, 80, 60, 40, 20, 0

| | NaCl eluate | DTT eluate |
|---|---|---|
| PAICS (C/EC) | 14.61 | 85.39 |
| PAWR (C/NU) | 2.83 | 97.17 |
| PBK (NU) | 15.56 | 84.44 |
| PCBP1 (C/NU) | 20.71 | 79.29 |
| PCMT1 (C) | 6.64 | 93.36 |
| PDCD6IP (C/CS/EC) | 38.25 | 61.75 |
| PDHB (MT) | 7.29 | 92.71 |
| PFDN1 (C) | 24.15 | 75.85 |
| PFDN2 (C/NU/MT) | 10.45 | 89.55 |
| PFDN4 (C/NU/MT) | 7.73 | 92.27 |
| PFDN5 (C/NU) | 3.65 | 96.35 |
| PFN2 (PFN2) | 10.29 | 89.71 |
| PGLS (C) | 43.32 | 56.68 |
| PHAX (C/NU) | 8.25 | 91.75 |
| PIN1 (C/NU) | 5.66 | 94.34 |
| PITHD1 (C) | 2.78 | 97.22 |
| PITRM1 (MT) | 21.04 | 78.96 |
| PLS3 (C) | 23.28 | 76.72 |
| PM20D2 (NU) | 13.85 | 86.15 |
| PMPCB (MT) | 86.37 | 13.63 |
| PNPO (C) | 4.91 | 95.09 |
| POLDIP2 (NU/MT) | 23.16 | 76.84 |
| PPAT (C) | 18.71 | 81.29 |
| PPCDC (C) | 4.18 | 95.82 |
| PPL (C/CS/PM) | 0.87 | 99.13 |
| PPP1R11 (C/NU) | 2.97 | 97.03 |
| PPP1R2 (???) | 0 | 100.00 |
| PPP1R7 (NU) | 21.84 | 78.16 |
| PPP2R2D (C) | 1.76 | 98.24 |
| PRDX1 (C) | 4.06 | 95.94 |
| PRDX2 (C) | 9.14 | 90.86 |
| PRDX3 (MT/C/EC) | 10.13 | 89.87 |
| PRDX4 (C/ER) | 2.20 | 97.80 |
| PRDX5 (C/NU/MT/PO) | 8.32 | 91.68 |
| PRDX6 (C/LY) | 5.18 | 94.82 |
| PREP (C) | 30.63 | 69.37 |
| PRMT1 (C/NU) | 58.72 | 41.28 |
| PRMT5 (C/NU/GA) | 15.94 | 84.06 |
| PSMD9 (C/NU) | 6.77 | 93.23 |
| PSME3 (C/NU) | 13.91 | 86.09 |
| PSME3IP1 (NU) | 5.56 | 94.44 |
| RABEP2 (C/CS/ES) | 15.81 | 84.19 |
| RANBP3 (C/NU) | 5.46 | 94.54 |
| RBBP7 (NU) | 19.23 | 80.77 |
| RBM8A (C/NU) | 34.45 | 65.55 |
| RCC2 (NU/PM/CS) | 10.00 | 90.00 |
| RILPL1 (C/CS) | 8.88 | 91.12 |
| RPIA(C) | 20.88 | 79.12 |
| RRM2 (C) | 17.97 | 82.03 |
| S100A9 (C/CS/PM/EC) | 64.85 | 35.15 |

**Table S8 (page 5)**

**External $H_2O_2$ (1 mM, 10 min)**

Color scale: protein abundance (percentage distribution), 0–100

| NaCl eluate | DTT eluate | |
|---|---|---|
| 4.12 | 95.88 | SDE2 (NU) |
| 5.59 | 94.41 | SELENBP1 (C/NU) |
| 6.31 | 93.69 | SERPINA1 (ER/EC) |
| 5.58 | 94.42 | SERPINB1 (C/ES/LY/EC) |
| 5.47 | 94.53 | SFN (C/NU/EC) |
| 0 | 100.00 | SH3BP5 (MT/ES) |
| 27.45 | 72.55 | SHTN1 (CS) |
| 3.65 | 96.35 | SKP1 (C/NU) |
| 6.08 | 93.92 | SLC9A3R (NU/PM) |
| 2.11 | 97.89 | SMAP1 (PM) |
| 25.24 | 74.76 | SNAP23 (PM) |
| 9.15 | 90.85 | SNAPIN (C/LY/GA/VE) |
| 14.25 | 85.75 | SOD1 (C/NU/MT) |
| 10.13 | 89.87 | SOD2 (MT) |
| 3.98 | 96.02 | SORBS3 (NU/CS) |
| 78.42 | 21.58 | SRSF10 (C/NU) |
| 23.09 | 76.91 | SRXN1 (C) |
| 13.52 | 86.48 | STRAP (C/NU) |
| 11.37 | 88.63 | TACC3 (C/CS) |
| 7.00 | 93.00 | TBCA (CS) |
| 10.63 | 89.37 | TCEAL4 (NU) |
| 8.53 | 91.47 | THOP1 (C) |
| 16.26 | 83.74 | TIPRL (C) |
| 2.48 | 97.52 | TK1 (C) |
| 11.33 | 88.67 | TPM1 (CS) |
| 4.90 | 95.10 | TPM3 (CS) |
| 2.67 | 97.33 | TPM4 (CS) |
| 14.14 | 85.86 | TSN (C/NU) |
| 16.90 | 83.10 | TSNAX (NU/GA/C) |
| 0.96 | 99.04 | TSR2 (???) |
| 2.51 | 97.49 | TTC1 (C/PO) |
| 1.70 | 98.30 | TTR (C/EC) |
| 11.00 | 89.00 | TUBA4A (CS) |
| 26.71 | 73.29 | TWF1 (C/CS) |
| 12.82 | 87.18 | TWF2 (C/CS) |
| 0 | 100.00 | TXLNA (C/EC) |
| 2.96 | 97.04 | TXN (C/NU/EC) |
| 37.68 | 62.32 | TXNDC17 (C) |
| 56.22 | 43.78 | TXNL1 (C/NU) |
| 3.08 | 96.92 | UBAP2 (C/NU) |
| 3.61 | 96.39 | UBE2A (C/NU) |
| 2.94 | 97.06 | UBE2B (NU/PM) |
| 7.15 | 92.85 | UBE2G1 (C/EC) |
| 3.62 | 96.38 | UBE2H (C/NU) |
| 2.75 | 97.25 | UBE2I (C/NU) |
| 10.10 | 89.90 | UBE2K (C) |
| 2.76 | 97.24 | UBE2L3 (C/NU) |
| 7.26 | 92.74 | UBE2N (C/NU) |
| 17.42 | 82.58 | UBFD1 (???) |
| 4.46 | 95.54 | UQCRH (MT) |

##### External $H_2O_2$ (1 mM, 10 min)

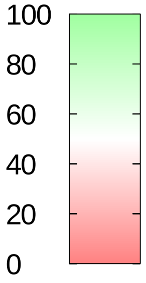


protein abundance (percentage distribution)

| | NaCl eluate | DTT eluate |
|---|---|---|
| USP5 (C/NU/LY) | 46.99 | 53.01 |
| VBP1 (C/NU) | 9.00 | 91.00 |
| WBP2 (C/NU) | 9.92 | 90.08 |
| WDR44 (C/GA/ES) | 14.31 | 85.69 |
| WDR48 (C/NU/LY/ES) | 66.21 | 33.79 |
| WDR77 (C/NU) | 16.87 | 83.13 |
| XRCC4 (NU) | 2.09 | 97.91 |
| YWHAE (C/NU) | 27.91 | 72.09 |
| YWHAG (C) | 45.04 | 54.96 |
| YWHAQ (C) | 10.31 | 89.69 |
| ZFAND5 (C) | 71.55 | 28.45 |
| ZFYVE16 (C/ES) | 0 | 100.00 |

**Table S1**

| Protein | Cysteine position | | |
|---|---|---|---|
| | External $H_2O_2$ | Peroxisomal $H_2O_2$ | Cytosolic $H_2O_2$ |
| ACP1 | 13, 18 | | |
| ACP6 | 416 | | |
| AHCY | 195 | | |
| AIFM1 | | | 317, 441 |
| AK2 | 92 | | |
| AKAP12 | 1139, 1399, 1407, 1521 | | |
| ALDOA | 339 | 339 | 73, 339 |
| ANXA2 | 133 | | |
| ARG1 | | 168 | |
| ASF1B | 172, 178 | | |
| ASNS | 158 | | |
| BAG3 | 151 | | |
| BLMH | 326 | 326 | |
| BSG | | | 242 |
| CACYBP | 154 | | |
| CALR | | 105 | |
| CANX | | | 360, 366 |
| CAPZB | 206 | | 206 |
| CASP14 | | 24 | |
| CAST | 241, 328, 381, 408, 413, 661 | | 661 |
| CBX3 | | | 177 |
| CDK4 | 78 | 78 | |
| CHCHD4 | 87 | | |
| CHMP5 | 20 | | |
| CHORDC1 | 24, 59, 211 | | 59 |
| CIAPIN1 | 285, 288 | | 274, 277 |
| CNN3 | 173 | | |
| COX17 | 23, 24, 26, 45 | | 36, 45 |
| CPD | | | 195 |
| CPOX | 319 | | |
| CRKL | 44, 249 | | 44 |
| CTH | 109 | | |
| CTSB | 93, 105, 108 | | 105, 108 |
| CTSD | | 329 | |
| DCTPP1 | 162 | | |
| DFFA | 38 | | |
| DHPS | 142, 177 | | |
| DNAJB11 | 193, 196 | | 193, 196 |
| DNPEP | 48, 337, 457 | | |
| DPYSL2 | 248 | | 248 |
| DPYSL3 | 39 (alternative sequence), 248, 448, 471 | | 248 |

| Protein | Cysteine position | | |
|---|---|---|---|
| | External $H_2O_2$ | Peroxisomal $H_2O_2$ | Cytosolic $H_2O_2$ |
| DSG1 | | 253, 255, 589, 669 | |
| DSP | 1069 | 57, 135, 174, 482, 798, 1069, 1805, 1957, 1983, 2259, 2656 | 135, 174, 482, 798, 1069 |
| DYNLL1 | 56 | | |
| EEF2 | | | 591, 693 |
| EIF3G | 139 | | |
| EIF3I | 76 | | |
| EIF4EBP1 | 62 | | |
| EIF5A | 73 | | |
| ENO1 | 337, 339 | 337, 339 | |
| ENSG00000286022 | | | 386 |
| EPDR1 | 42 | | |
| ERICH5 | 35 | | |
| ERP44 | | 318 | 92, 189, 301, 318 |
| FABP5 | | 120, 127 | |
| FLG2 | 317 | 25, 317, 329, 338, 346, 366 | |
| FN3KRP | 24 | | |
| GAPDH | 247 | 247 | 152, 156, 247 |
| GGCT | | 42 | |
| GLOD4 | 41 (isoform 2), 197, 221 | | 197, 221 |
| GLRX3 | 229 | | |
| GPC4 | 60, 66, 67, 341 | | |
| GPHN | 212, 284, 419 | | |
| GRB2 | 32 | | |
| GRN | 164, 165, 171, 215, 221, 222, 284, 290, 296, 297 | | |
| GSR | | 377 | 102, 107, 467 |
| HPRT1 | 66, 106, 206 | 106, 206 | 66, 206 |
| HSD17B10 | 58, 91, 214 | 58, 91 | 91 |
| HSPA1B | | | 306 |
| HSPA4L | 540 | | |
| HSPA8 | | 17 | 17 |
| HSPA9 | | | 66, 366 |
| IMPDH2 | 468 | | |
| ISCA2 | 79 | | |
| ISCU | 69, 95 | | |
| ISYNA1 | 485 | | |
| JMJD6 | 101, 207 | | |
| JPT2 | 118 | | |
| JUP | | 511, 457 | |
| LCP1 | 101, 140 | | 140 |
| LDHA | | 163 | 163 |

| Protein | Cysteine position | | |
|---|---|---|---|
| | External $H_2O_2$ | Peroxisomal $H_2O_2$ | Cytosolic $H_2O_2$ |
| LDHB | 164 | 164 | 164 |
| LMAN2 | | | 202, 239 |
| LRRFIP1 | 644 | | |
| LTF | 28, 64, 189, 200, 367, 424, 512, 523, 526, 608, 668, 696, 705 | 64, 668 | |
| LYRM7 | 97 | | |
| MALSU1 | 110 | | |
| MAP4 | 535 | | |
| MARCKSL1 | 134 | | |
| MCMBP | 287 | | |
| MDH1 | 137, 154 | | |
| NAA10 | 194 | | |
| NASP | 254, 708 | | 254, 708 |
| NDUFAB1 | 140 | | |
| NDUFS6 | 87, 112, 115 | | |
| NFU1 | 210, 213 | | |
| NME1-NME2 | | | 109 |
| OTUB1 | 23 | | |
| P3H1 | 645 | | |
| PAWR | 173 | | |
| PBK | 22, 70, 300 | | |
| PCBP1 | 54, 109, 158, 194 | | |
| PCCA | 111 | 111 | |
| PCMT1 | 95 | 95 | |
| PDIA3 | 85, 92 | | 85, 92 |
| PDLIM1 | 73 | | |
| PHAX | 51 | | |
| PIN1 | 57, 113 | | |
| PITHD1 | 14, 42, 187 | | 42, 187 |
| PKP1 | | | 161, 409 |
| PLS3 | 104, 167 | 167 | 104, 143 |
| PM20D2 | 303 | | |
| PNPO | 156 | | |
| PPCDC | 7 | | |
| PPIA | | 62 | |
| PPP1CC | 62 | | |
| PPP1R11 | 60, 61, 62 | | |
| PRDX1 | 173 | 173 | 71, 83, 173 |
| PRDX2 | 172 | | |
| PRDX3 | 229 | | 229 |
| PRDX4 | 51, 245 | 51 | 51, 245 |

| Protein | Cysteine position | | |
|---|---|---|---|
| | External $H_2O_2$ | Peroxisomal $H_2O_2$ | Cytosolic $H_2O_2$ |
| PRDX5 | 100 | | |
| PRDX6 | 47, 91 | 47, 91 | 47 |
| PRMT5 | 196 | | |
| PSMD9 | 59 | | |
| PSME3 | 92 | | |
| PSME3IP1 | 187 | | |
| PXN | 108 | | |
| RANBP3 | 228, 249 | | |
| RBBP7 | 97 | | 97 |
| RCC2 | 209, 337 | | |
| RPL7A | | 199 | |
| S100A14 | | 74 | 74 |
| S100A7 | | | 47 |
| S100A8 | | 42 | 42 |
| SELENBP1 | 31, 57, 268 | | |
| SHTN1 | 565 | | |
| SKP1 | 120, 160 | 160 | 120 |
| SMAP1 | 33, 36 | | |
| SNAP23 | 112 | | |
| SOD1 | 58, 147 | | 147 |
| SRXN1 | 99 | | |
| STRAP | 152, 305 | | 305 |
| TACC3 | 20, 206, 242, 426 | | |
| TBCA | 67 | | |
| TIPRL | 14, 87 | | |
| TK1 | 66, 153, 156, 206, 230 | | |
| TMED7 | | | 48, 109 |
| TPI1 | | 218 | |
| TPM3 | 170, 226, 233 (all in isoform 2) | | 170, 233 (all in isoform 2) |
| TSR2 | 114 | | |
| TTC1 | 28 | | |
| TWF2 | 67, 141 | | |
| TXN | 73 | 73 | 32, 35, 73 |
| TXNDC17 | 43, 46 | | |
| TXNDC5 | 121, 128, 381 | | 121, 128, 381 |
| UBE2I | 43 | | |
| UBE2K | 170 | | |
| UBE2L3 | 86 | | 86 |
| UBE2N | 87 | | |
| UCHL1 | | | 90, 152, 220 |
| UQCRH | 37, 53 | | |

| **Protein** | **Cysteine position** | | |
|---|---|---|---|
| | **External $H_2O_2$** | **Peroxisomal $H_2O_2$** | **Cytosolic $H_2O_2$** |
| VBP1 | 113 | | |
| WDR48 | 225 | | |
| YWHAQ | 94, 134, 237 | | 134 |
| YWHAZ | 94 | | |
| ZFAND5 | 76 | | |
